# Extensive binding of nebulous human transcription factors to genomic dark matter

**DOI:** 10.1101/2024.11.11.622123

**Authors:** Rozita Razavi, Ali Fathi, Isaac Yellan, Alexander Brechalov, Kaitlin U. Laverty, Arttu Jolma, Aldo Hernandez-Corchado, Hong Zheng, Ally W.H. Yang, Mihai Albu, Marjan Barazandeh, Chun Hu, Ilya E. Vorontsov, Zain M. Patel, The Codebook Consortium, Ivan V. Kulakovskiy, Philipp Bucher, Quaid Morris, Hamed S. Najafabadi, Timothy R. Hughes

## Abstract

The functional impact of a large portion of the human genome known as “dark matter DNA”, which is composed mainly of repeat sequences, remains enigmatic. The genome also encodes hundreds of putative and poorly characterized transcription factors (TFs). Here, we determined genomic binding locations of 166 poorly characterized human TFs in living cells. Nearly half of them associate strongly with known regulatory regions such as promoters and enhancers, frequently co-localizing with each other at conserved motif matches. The other half often associate with genomic dark matter, however, at largely non-overlapping (i.e., unique) sites, via intrinsic sequence recognition. Fifty-four of the latter half, which we term “Dark TFs”, mainly bind within regions of closed chromatin, with each recognizing a unique set of repeat sequences. The Dark TFs include many KZNFs, which are known to bind and silence TEs, and other TFs with apparent repressive functions. By contrast, some may be pioneers: we find that induction of TPRX1, a known regulator of zygotic preimplantation, leads to chromatin opening at many of its binding sites in the dark matter genome. Altogether, our results shed light on a large fraction of poorly characterized human TFs and simultaneously illuminate the diversity of function within the dark matter genome.

## INTRODUCTION

For decades, the term “dark matter”, borrowed from astronomy and cosmology, has been applied to describe the excess of seemingly non-functional regions of many genomes^1^. These regions are dominated by repetitive sequences such as transposable elements (TEs) and endogenous retroelements (EREs), which largely account for the extreme differences in genome size across eukaryotes. There are many potential roles for genomic dark matter^1,2^, but it is also conceivable that it is mainly inert or simply provides spacing. An initial proposal for the function of the dark matter was gene regulation^3^, and indeed, while some so-called “gene deserts” appear to be dispensable, others contain transcriptional enhancers^4–6^. Function of genomic dark matter in gene regulation is also supported by the existence of transcription factors (TFs) that are capable of binding TE and ERE sequences^7^.

It is well-established that TEs can rewire regulatory circuitry by introducing transcription factor binding sites, thus spawning novel cis-regulatory elements^8–10^. In some cases, these elements derive from the promoter of the TE (e.g., endogenous retrovirus long terminal repeats (ERV LTRs))^11,12^. Other cases may represent inadvertent matches to a TF motif within a TE^13^; over evolutionary time, regulatory sites tend to turn over rapidly, arising seemingly at random, and TEs/EREs are a known contributor to this process^14^. In addition, mammalian genomes encode a large family of TFs, the KRAB-containing C2H2 zinc-finger (KZNF) class (∼350 members in human), which has evolved rapidly in parallel to ERE classes bound by its members^15^. Different KZNFs typically bind to different specific subclasses of ERE in ChIP-seq^7,16^. The KRAB domain is best known for recruiting KAP1/TRIM28, which associates physically with both readers (e.g., HP1/CBX proteins) and writers (SETDB1)^17,18^ of the H3K9me3 mark that defines constitutive heterochromatin^19,22^. Silencing EREs, presumably to limit their spread, appears to be the major driver of KZNF evolution, but the fact that so many KZNFs have been retained, often tens of millions of years after the EREs have been reduced to molecular fossils, suggest that the KZNFs often contribute to adaptive evolution^20,21^. At least some have clearly taken on regulatory roles that benefit the genome in which they reside^21,22^.

There are also many poorly characterized human TFs that, like the dark matter genome, remain nebulous in regard to DNA-binding, which is the defining property of TFs: the human genome encodes over 1,600 apparent TFs, but hundreds of them have been identified as such only on the basis of conserved protein domain structures and have no known DNA binding motif^23^. TFs overall are best known for their association with promoters and/or enhancers^24,25^, although the largest class in human is the KZNFs, which appear primarily to act as repressors. TFs also vary greatly in their ability to bind nucleosomal DNA^26^, and only a subset of TFs, termed “pioneers”, can access binding sequences within closed chromatin and recruit or allow access to other TFs^27^. There are many possible mechanisms of cooperation between TFs and TF binding sites, of which pioneering is just one example^28^. Presumably as a consequence, decoding gene regulation is a long-standing challenge in molecular biology and computational genomics, which is compounded by the fact that the sequence preferences of many TFs remain unknown.

In addition, even for TFs with known motifs, it is uncertain whether the motif is an accurate representation of the TF’s sequence preferences. The KZNFs, for example, are known for the potential to have very long binding sites, enabled by their long C2H2-zf domain arrays^29^, and the DNA binding preferences of these proteins have proven difficult to characterize precisely due to generally poor functionality in biochemical assays. ChIP-seq reveals what families of EREs are bound by C2H2-zf proteins, but it is difficult to accurately determine their precise sequence specificity because the repeat elements are related by common descent, which confounds motif discovery and testing^30^. In addition, *in vivo* measurements include the effects of indirect recruitment, nonspecific binding, and chromatin^31,32^. For KZNFs, it has not been proven that the sequence specificity of the C2H2-zf array on its own is sufficient to identify specific ERE classes within the genome: the predictions from scans with existing motifs yield an excess of matches outside EREs^33^.

As part of an international initiative termed the “Codebook consortium”, aimed at obtaining binding motifs for all human TFs^34^, we analyzed 315 human TFs (i.e., with unknown DNA sequence specificity) by ChIP-seq in HEK293 cells, together with 58 controls. Roughly 60% of these TFs (n=217) yielded reliable results in our study (i.e., reproducible and/or enriched for TF motif matches). We evaluated the data in conjunction with other data from the Codebook project, which includes binding to the human genome *in vitro*^35^, and use of multiple *in vitro* data types to define binding motifs^34,36^, thus allowing base-level identification of direct binding sites in ChIP-seq. Previous ChIP-seq analyses have focused mainly on preferential association of TFs with promoters vs. enhancers^24^, and consistent with this expectation, roughly half of the TFs with successful data in our study recognize such sites. Surprisingly, however, the other half of the Codebook TFs, including most KZNFs as well as other TF families, bound to apparently unique sites dispersed across the genome that lack active chromatin marks prior to introduction of the TF. Fifty-four of these TFs bind mainly to sequences located in closed chromatin. We refer to this subset as “Dark TFs” and show that most bind preferentially to specific subfamilies of TEs and EREs. Our further investigation into the binding sites for three Dark TFs revealed significant reconfiguration of chromatin architecture at these genomic locations. Multiple lines of evidence suggest diverse biochemical, cellular, and physiological functions of the Dark TFs, and by extension, shed light on potential functions of the dark matter genome.

## RESULTS

### Generation of ChIP-seq data for hundreds of putative TFs

We surveyed the genomic binding sites of 315 poorly characterized, putative human TFs (the “Codebook” set, derived from Lambert 2018^37^, and described in detail elsewhere^34^), and 58 previously characterized TFs as controls (selected from Isakova 2017^38^ and Schmitges 2016^33^), using ChIP-seq in HEK293 cells (**Fig. 1a**, **Supplementary Table 1**). We used an established inducible eGFP-tagged transgene system (**Fig. 1b**) previously employed in ChIP-seq and protein-protein interaction studies^16,33,39^. Of particular relevance here, this system was effective at revealing specific binding of KZNFs to classes of retroelements, confirming that the system can readily assess binding to inactive or repressed regions of the genome^16,33^. The present study includes biological ChIP replicates for each TF performed by different experimentalists, such that the resulting dataset includes 678 ChIP-seq experiments for Codebook TFs, and 112 experiments for control TFs (**Supplementary Table 1**). A full list of experiments is given in **Supplementary Table 2**. Representative motifs obtained from control TFs of various families illustrate that the assay recovers known sequence-binding preferences, as expected (**Fig. 1c** and **Supplementary Fig. 1**).

**Figure 1.**
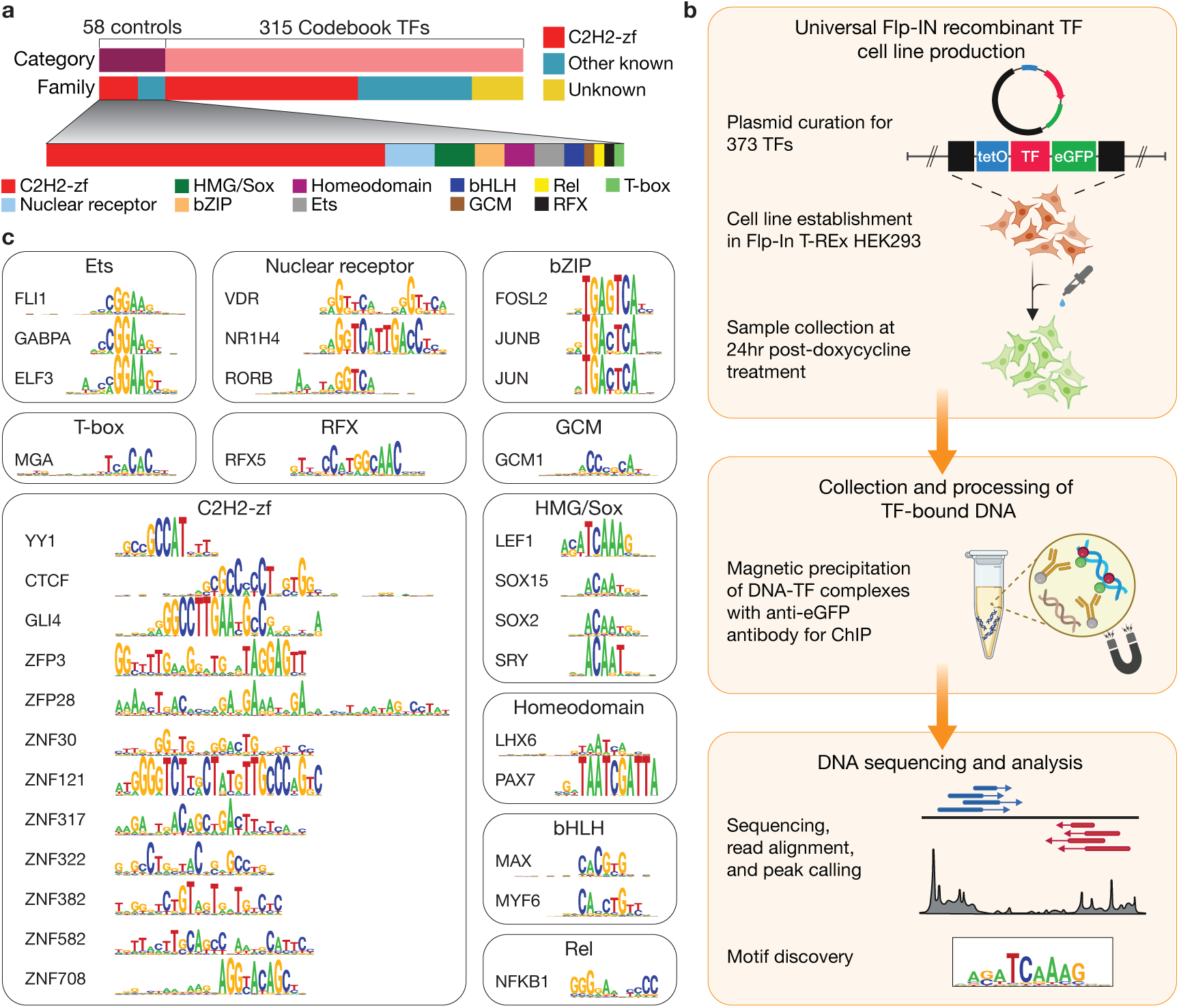
Overview of ChIP-seq strategy to identify genomic binding sites for 315 nebulous human TFs. (**a**) Summary of human TF categories (38 control and 315 Codebook) and protein families (with known and unknown classifications) that were selected for ChIP-seq. The “other known” category includes non-C2H2-zf protein families ten of which are included in the control TFs. (**b**) A schematic of the experimental pipeline for production of 373 inducible EGFP-labelled TF cell lines, universal Chromatin immunoprecipitation with eGFP antibody, and derivation of TF binding sites for each line. (**c**) Samples of representative motifs obtained for each protein family of the control TFs show expected matches.

We used two criteria to determine which experiments were successful. First, the data were analyzed as part of a larger Codebook benchmarking effort, which is described in more detail in accompanying manuscripts^34,40^. The Codebook benchmarking included expert curation that relied mainly on obtaining similar motifs for the same TF from different *in vivo* and *in vitro* data types (ChIP-seq, Protein Binding Microarrays^41^, SMiLE-seq^38^, and several variants of HT-SELEX^42^) as evidence of direct, sequence-specific DNA binding. This Codebook motif benchmarking identified 130 Codebook TFs and 49 controls with successful, motif-enriched ChIP-seq data, meaning that sequence-specific DNA binding observed in ChIP-seq (*in vivo*) is supported in almost all cases by *in vitro* experimental data (**Supplementary Table 1**).

Second, we identified experiments in which the peak overlaps of biological replicates exceeded what is expected at random (i.e., with TF identities permuted). ChIP-seq experiments that were classified as successful based on the motif similarity analysis described above displayed a higher overlap between TF replicates relative to mismatched TFs (median Kulczynski II coefficient of 0.57 vs. 0.034; **Supplementary Fig. 2a**; this statistic is a modified Jaccard value that compensates for class imbalance). A Kulczynski II coefficient threshold of 0.4 captures 79% of approved experiments with replicates, and 90% of controls, but eliminates 94.5% of mismatched experiments; thus, we used this value as a threshold for success of replicate experiments, independent of motifs. By this measure, an additional 36 putative TFs (and two controls) were considered technically successful, despite not displaying motif enrichment (**Supplementary Tables 1** and **2**). The DNA binding in these cases may be indirect, i.e., the proteins may be DNA-associated chromatin factors, rather than TFs. Alternatively, they may recognize properties of the DNA sequence that are not captured by common motif models. Nevertheless, we included these 38 TFs in subsequent analyses. Cautioning that these proteins may be chromatin factors, for simplicity, we refer to the entire set of 217 successful proteins (130 putative + 49 control TFs successful in the first round, and 36 putative + 2 controls with matching replicates; **Supplementary Fig. 3a** and **Supplementary Table 3**) as “TFs” in the text.

We merged the peaks from TF replicates among the 427 ChIP-seq experiments deemed successful to produce a dataset for downstream analysis (**Supplementary Table 3**). This dataset encompasses 217 proteins (166 Codebook putative TFs, and 51 controls), with a median of 12,200 peaks per protein (range 7-145,470) (using a MACS threshold of P<10^-10^; see **Methods** for explanation of threshold choice)^43^.

### ChIP-seq data illustrate that half of the Codebook TFs bind genomic dark matter

To begin characterizing the ChIP-seq data, we surveyed for preferential association of the putative TFs with promoters and/or enhancers. **Fig. 2a** shows the fraction of peaks from each protein that overlaps with protein-coding promoters (defined by RefSeq^44^) and enhancers (defined by HEK293 chromatin state^45^, and mainly carrying H3K4me1 signal; see **Methods** and **Supplementary Fig. 4a**). Indeed, many proteins are highly associated with promoters, and a smaller number with enhancers. On average, the HEK293-derived enhancer set yielded higher overlaps than the larger, more universal “GeneHancer” set^46^ (**Supplementary Fig. 4b**), indicating that this lower number of enhancer-favoring TFs (relative to promoter-favoring) is not due to incomplete enhancer annotations. Moreover, for control TFs with publicly available endogenous data in multiple cell types^47^, the fraction of peaks overlapping GeneHancer enhancers, HEK293 enhancers, or promoters shows a consistent trend with Codebook data, suggesting that the low number of enhancer-enriched TFs is not particularly due to HEK293-specific chromatinization (**Supplementary Fig. 4c-e**). Surprisingly, a large fraction of proteins did not associate with either promoters or enhancers (**Fig. 2a**).

**Figure 2.**
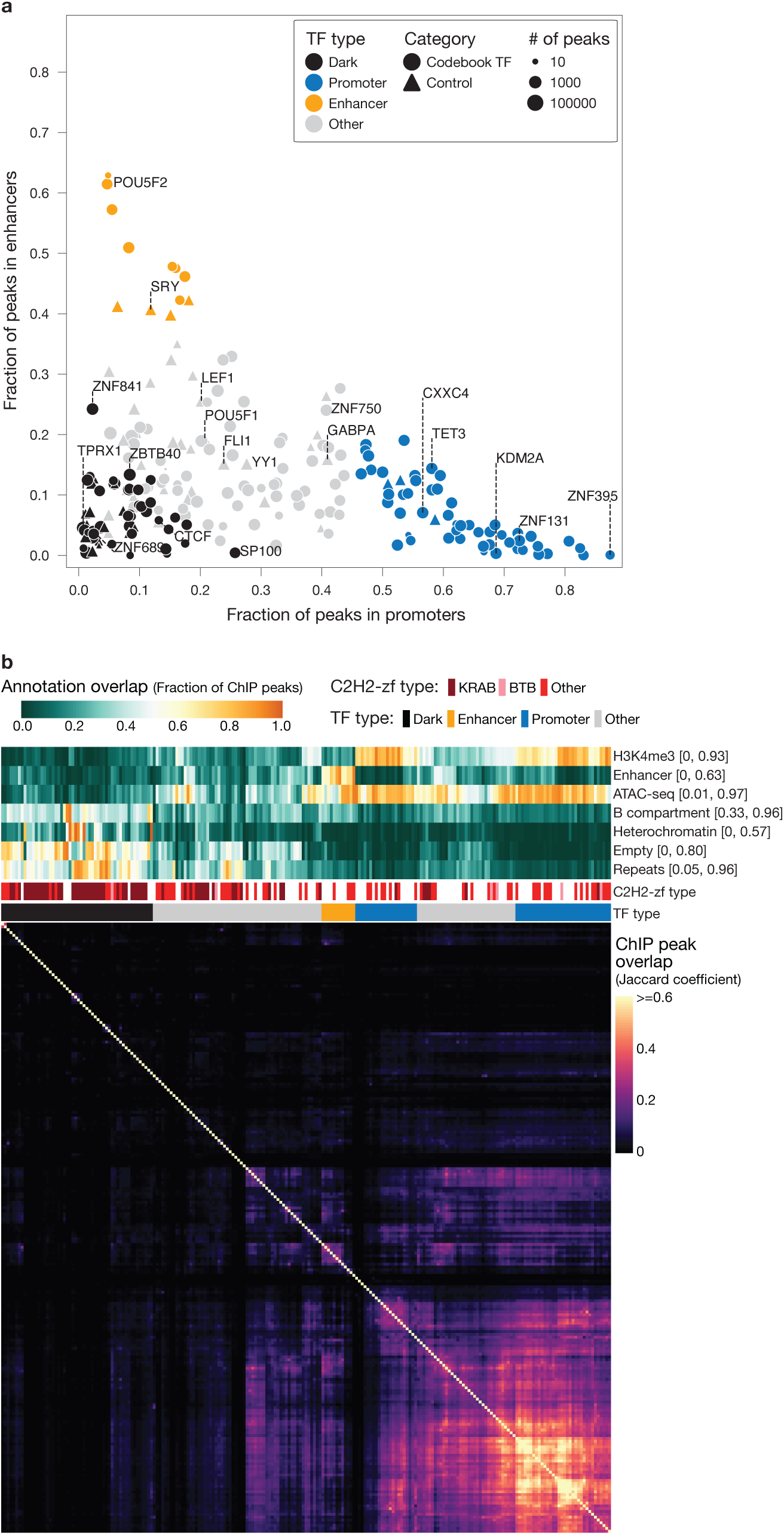
Distinct genomic binding patterns identifies four different TF types with varying preferences for accessible or closed chromatin. (**a**) distribution of ChIP-seq peak fractions for 217 TFs in protein-coding promoters (x-axis) and HEK293 enhancers (y-axis) used for discrimination of Promoter and Enhancer TFs. Dark TFs and “Other” categories are also labelled here. Point sizes are proportional to the number of peaks for each TF (log scale). (**b**) *Bottom:* (square) heatmap of Jaccard similarity coefficient between ChIP-seq peaks of all TF pairs showcasing the colocalization patterns of binding sites. The bright corner aligns with Promoter TFs. *Top:* heatmap of ChIP-seq peak fractions that fall within various annotated genomic tracks of chromatin accessibility as indicated, as well as protein properties of the TFs. Fractions are scaled to fit in [min, max] range across the TFs for better visualization, as indicated in the right. TF ordering is determined by hierarchical clustering with Ward linkage and Euclidean distance, using the tracks for ‘H3K4me3’, ‘ATAC-seq’, ‘B compartment’, ‘Empty’ + ‘Heterochromatin’, ‘Repeats’, ‘CpG’, ‘Protein-coding promoters’, ‘H3K27ac’ (the last three not shown), along with the one-hot encoded ‘TF type’ to aid in illustration.

To gain further insight into the properties of the ChIP-seq binding sites, we compared the peak sets for each of the proteins to all other proteins in the dataset, along with an extensive panel of genome annotations, protein type, and expression levels (**Fig. 2b** and **Supplementary Fig. 5)**. **Fig. 2b** (bottom) shows a symmetric heatmap of Jaccard values (intersection/union) between the peak sets of the 217 successful TFs, providing an overview of similarities in genomic localization. The heatmap at the top of **Fig. 2b** shows the fraction of each corresponding TF peak set that overlaps with each type of genome annotation. The chromatin states were derived mainly from public-domain data for unperturbed HEK293 cells; we therefore expect them to reflect the state of the chromosomes prior to induction of the tagged TF transgene. This state could be involved in recruiting the TF, but it could also result from endogenous expression of the native, untagged protein, as most of the studied TFs are already expressed in HEK293 cells^33^ (**Supplementary Fig. 5**).

**Fig. 2b** reflects and expands upon trends observed in **Fig. 2a**. The large bright square in the lower right quadrant of the bottom heatmap corresponds to TFs that associate primarily with open chromatin (ATAC-seq) and H3K4me3. These TFs also often associate with many promoters (median 8,813 coding gene promoters; see below), leading to high overlap between the peak sets. The observation that many TFs co-bind promoters and/or enhancers is prevalent in the literature (e.g.^24^); we note, however, that the Codebook proteins were not previously known to bind these sequences, and therefore it appears that even well-known regulatory elements often contain previously unidentified TF binding sites.

A second main feature of **Fig. 2b** is the diagonal line in the upper left quadrant of the heatmap. These are cases in which each peak set has very little overlap with any other peak set. In addition, the individual peaks bound by each TF are often unique, i.e., they do not overlap with any peak from any other protein in the Codebook ChIP-seq dataset (**Supplementary Fig. 2b**). The unique binding profiles are not explained by experimental error or random events since there is strong peak overlap between replicates for the same protein (**Supplementary Fig. 2c**), and these proteins often bind to the same unique sequences both *in vivo* and *in vitro* (see below).

The ChIP-seq peaks for these proteins also tend to be outside open chromatin and outside of either promoters or apparent enhancers in HEK293 cells (**Fig. 2b**, *top*). In fact, roughly half of these TFs’ peak sets are either enriched for marks that characterize heterochromatin or lack any of the diagnostic marks of promoter or enhancer activity (i.e., “empty” ChromHMM regions, **Supplementary Fig. 4a**). Peaks for a subset of these TFs (n=87) are mainly associated with the Hi-C “B” compartment, and many associate with specific classes of repeats, particularly TEs (**Fig. 2b** and see below). This is an expected property of KZNFs, which constitute more than half (48/87) of these TFs^16^ (bar in the middle of **Fig. 2b**).

### Classification of TFs based on genomic annotations of binding sites

We sought to gain a better understanding of the properties and characteristics of TFs (and their binding sites) that represent the major patterns shown in **Fig. 2**. As a starting point, we defined four mutually exclusive groups (see **Supplementary Table 3** for TF labels) based on the dominant annotation types that overlap with their genomic binding sites (i.e., ChIP-seq peaks). One group we named “Promoter TFs” (56 proteins); for these proteins, more than 45% of peaks exclusively overlap with promoters (this threshold captures the most prominent features in **Fig. 2a**). Another group was designated “Enhancer TFs” (12 proteins); for these proteins, >37% of peaks overlap with enhancers (this threshold corresponds to the visual separation of data points on the vertical axis in **Fig. 2a**). A third group we named “Dark TFs”, after the genomic “dark matter” (54 proteins); for these, most peaks lie within either the “empty”, “constitutive heterochromatin”, or “facultative heterochromatin” states (i.e. in HEK293 cells, they are outside of the states that represent promoters, enhancers, insulators, or gene bodies), and fewer than half of the peaks overlap with ATAC-seq peaks in unperturbed HEK293 cells. The remaining ∼40% of TFs we labeled as “Other” (i.e., not Promoter, Enhancer, or Dark TFs); they include 30 control TFs and 65 Codebook proteins, which display diverse combinations of binding sites within the defined categories that do not reach our classification thresholds. The “Other TFs” thus present a rich landscape for additional exploration, but we did not attempt to further subclassify them here.

The choice of features and thresholds to define the categories of TFs was to readily draw distinctions between TFs via interpretable binding behaviors. Unbiased clustering of the TFs using all the tracks presented on the top in **Fig. 2b** results in the nearly same groups, but with less stringency (i.e. some “Other” TFs being labeled as Promoter TFs, Enhancer TFs, or Dark TFs, **Supplementary Fig. 6**). Our thresholds do exclude some TFs that may associate significantly with specific regions but also bind many other locations; for example, the control TF YY1 bound 58% of all promoters in human, but it also had 40,764 additional binding sites (72% of all its binding sites) outside promoters. Similarly, some KZNFs were not classified as Dark TFs because they had many binding sites within open chromatin.

Since the Promoter TFs and Dark TFs bind distinct and prominent genomic features and contain most of the Codebook TFs, we focused mainly on contrasting these two groups for the remainder of our analyses. It is important to note that the Promoter and Dark TF classifications could not be accounted for by technical artifacts despite the known bias of ChIP-seq towards open chromatin^32^: the fraction of reads in peaks (FRiP) for Dark TFs is comparable to those of Promoter TFs (**Supplementary Fig. 7a**), indicating that there is no systematic difference between the quality of the peaks called within promoter regions or elsewhere. Moreover, the absence of peaks within empty or heterochromatin regions for Promoter TFs or other non-Dark TFs is not due to the lack of read coverage in those regions: for all Promoter TFs (except CXXC4), >85% of ChIP-seq reads are outside promoters (**Supplementary Fig. 7b**). In addition, the reproducibility of the Dark TF peaks is comparable to that of the Promoter TF peaks (median optimized Jaccard between the replicates is 0.59 for Dark TFs, and 0.55 for Promoter TFs) (**Supplementary Table 3**, **Supplementary Fig. 2a**). **Supplementary Fig. 7c** illustrates that, in many cases, the imperfect overlap between peak sets is at least partially accounted for by peaks near the threshold, which pass the cutoff in one experiment but not the other. Altogether, both Promoter and Dark TF peak sets are technically robust by conventional measures.

### Recognition of repeat families by Dark TFs is direct and sequence-specific

Dark TF ChIP-seq peaks in general tend to overlap genomic repeats, and 36 of the 54 Dark TFs are KZNFs, which are expected to bind to specific ERE classes (**Fig. 2b**). We further interrogated Dark TFs’ peaks to ask whether each TF displays preferences for specific repeat types. Indeed, in addition to a large fraction of peaks overlapping repeats, there is a clear tendency for the ChIP-seq peak sets of each Dark TF to display strong statistical enrichment with at least one specific and unique repeat family, mainly corresponding to TEs and EREs (**Fig. 3a**). In contrast, many fewer Promoter TF peaks overlap repeats, and the most enriched repeat is typically “G-rich”, which itself is enriched at promoters (2,425 in promoters out of 13,493 in the genome).

**Figure 3.**
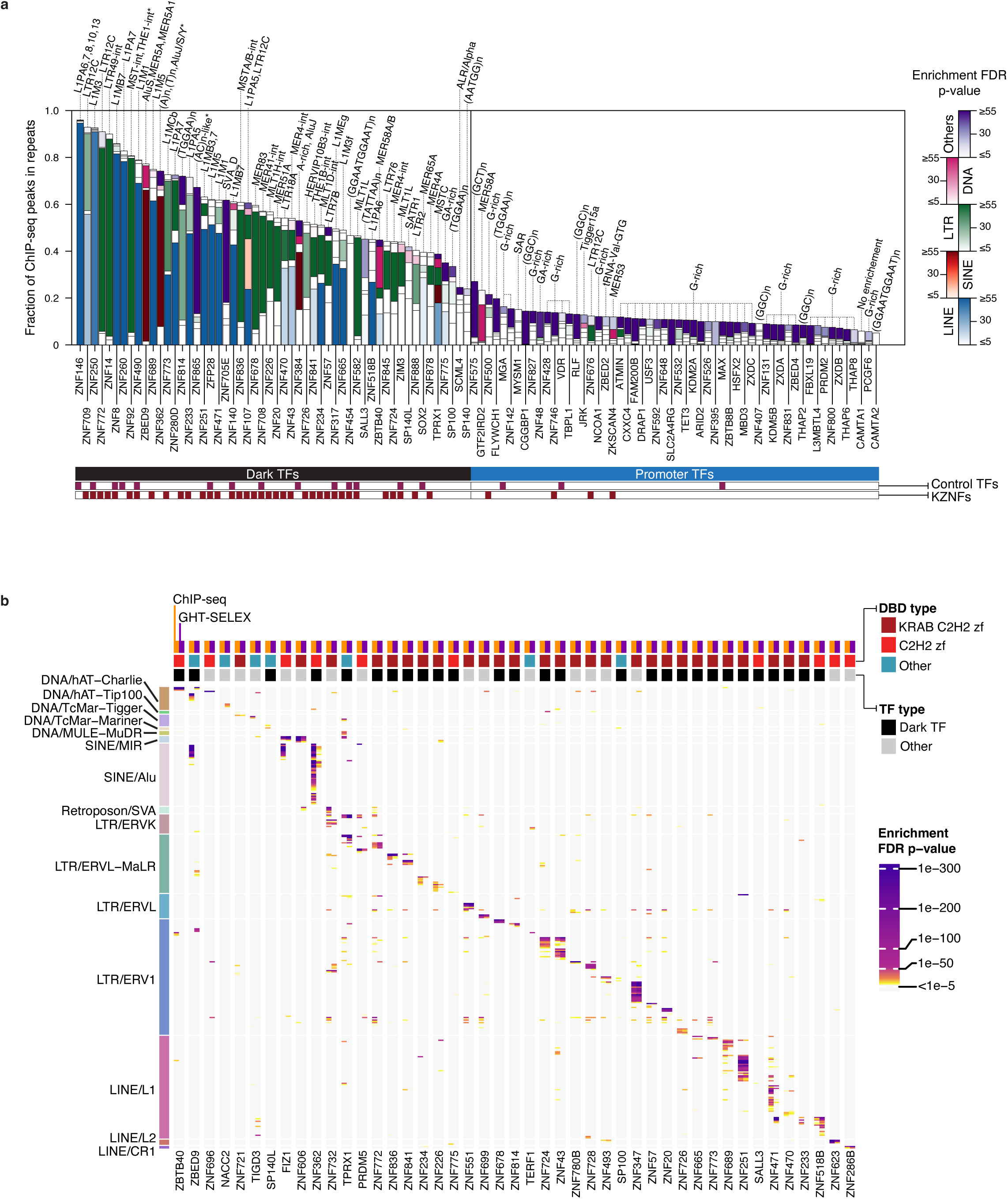
Dark TFs recognize specific families of transposable elements. (**a**) Stacked bar chart showing the fraction of ChIP-seq peaks (at MACS2.0 P<10^-10^ threshold) overlapping with repeats, partitioned into 5 superclasses: LINEs (blue), SINEs (red), LTRs (green), DNA transposons (orange), and other types (purple), for all Dark TFs and Promoter TFs. Color intensities represent the highest enrichment (–log10 FDR-corrected p-values) between the peak sets of a TF and the repeat families within that corresponding superclass. The most enriched repeat family is labeled above each bar. (**b**) Heatmap showing the enrichment (–log10 FDR-corrected p-values) of ChIP-seq (yellow) and GHT-SELEX (purple) peaks for binding to specific TE families (y-axis). Only Codebook TFs (i.e. controls excluded) having both datasets and with >30% of their ChIP-seq peaks overlapping any repeat are shown, using universal thresholds (MACS2.0 P<10^-10^ for ChIP-seq and knee specificity ≥30 for GHT-SELEX; see **Methods**). Labels on the left indicate TE superfamilies and families.

ChIP-seq can detect both direct binding (i.e., the TF intrinsically recognizes the bound DNA sequences) and indirect binding (e.g., recruitment by another factor)^31^. ChIP-seq also readily detects non-specific DNA-binding (e.g., by histones), and is biased towards open chromatin since the sonication step preferentially releases these regions^32^. To determine whether the ChIP-seq peaks are due to direct sequence-specific binding of the TFs to the genome, and to accurately identify these direct binding sites, we employed data from a novel assay, GHT-SELEX (Genomic HT-SELEX; described in detail in ^48^), which was available from the Codebook project. GHT-SELEX surveys the binding of synthetic TFs to fragmented, purified, and unmodified genomic DNA *in vitro*, yielding peaks that resemble those from ChIP-seq. The GHT-SELEX data differs from ChIP-seq in several ways, however: 1) it has greater resolution, due to the smaller DNA fragment lengths (∼64 bases), and 2) GHT-SELEX represents only the intrinsic binding of the TF to naked genomic DNA, devoid of chromatin and other cellular cofactors, which can modulate binding *in vivo*. As a result, only a subset of the ChIP-seq and GHT-SELEX peaks typically overlap (see below). Nonetheless, the general properties of binding sites across the genome can be readily compared.

**Fig. 3b** provides an overview of the statistical enrichment of specific TE families for all Codebook TFs (i.e., excluding control TFs) for which >30% of the ChIP-seq peaks overlap any kind of repeat (**Supplementary Document 1** shows an expanded version with all rows labelled), and for which GHT-SELEX data was obtained^35^. Strikingly, the enriched TEs in the GHT-SELEX data closely correspond to those in the ChIP-seq data (shown adjacent to each other in **Fig. 3b**), even though one assay is *in vivo* and one is *in vitro*. Thus, specific binding to TEs is an intrinsic and independent property of individual TFs, including many KZNFs. Most of these associations are also supported by the presence of binding motif matches (see below). Previous motif models derived from other KZNFs’ ChIP-seq data were unable to specify precise direct genomic binding to their target elements^33^. To our knowledge, this is the first experimental demonstration that KZNFs possess autonomous and sufficient sequence specificity to discriminate ERE subfamilies from the rest of the genome.

**Fig. 3b** also illustrates that many TFs in the “Other” class bind to specific repeat families, of which most (9/17) are KZNFs. Thus, the high association of KZNFs with TEs is not limited to Dark TFs: among the 28 KZNFs binding to specific TE classes, 19 are Dark TFs, and nine are Other TFs. In addition, four of the Promoter TFs in the Codebook set are KZNFs. Their peaks overlap with fewer repeats, but they are nonetheless enriched for specific classes: ZKSCAN4 tends to bind MER53; ZNF676 significantly recognizes LTR12C; and ZNF746 and ZNF500 are biased towards G-rich repeats (**Fig. 3a**). Thus, while the Dark TFs are enriched for KZNFs, and both Dark TFs and KZNFs tend to bind to TEs, these associations are far from absolute.

### Direct and sequence-specific binding of Promoter TFs to promoter regions

We next asked whether the GHT-SELEX data displayed enrichment for the same broad genome annotations observed in ChIP-seq. The fraction of intrinsic (i.e., GHT-SELEX) and cellular (i.e., ChIP-seq) sites for each TF that overlap with protein-coding gene promoters, repeat sequences (of any kind), and the combination of the “empty” and “heterochromatin” states are shown in **Figs. 4a, b**, and **c**, respectively. In each case, there is preferential binding *in vitro* that corresponds to what is observed in cells, with Promoter TFs having much higher intrinsic preference for promoter DNA, and Dark TFs having higher preference for repeats and empty/heterochromatin (as expected from their binding to specific repeat classes). We note that many Promoter and Enhancer TFs have a greater tendency to bind the “empty” and “heterochromatin” state DNA *in vitro* than in cells (**Fig. 4c**). This could be due to (a) a *bona fide* preference for open chromatin (and/or inability to access “empty” and “heterochromatin” regions), (b) functional binding at these loci in other cell types (but not HEK293), and/or (c) preferential extraction of these proteins at open chromatin in ChIP-seq experiments.

**Figure 4.**
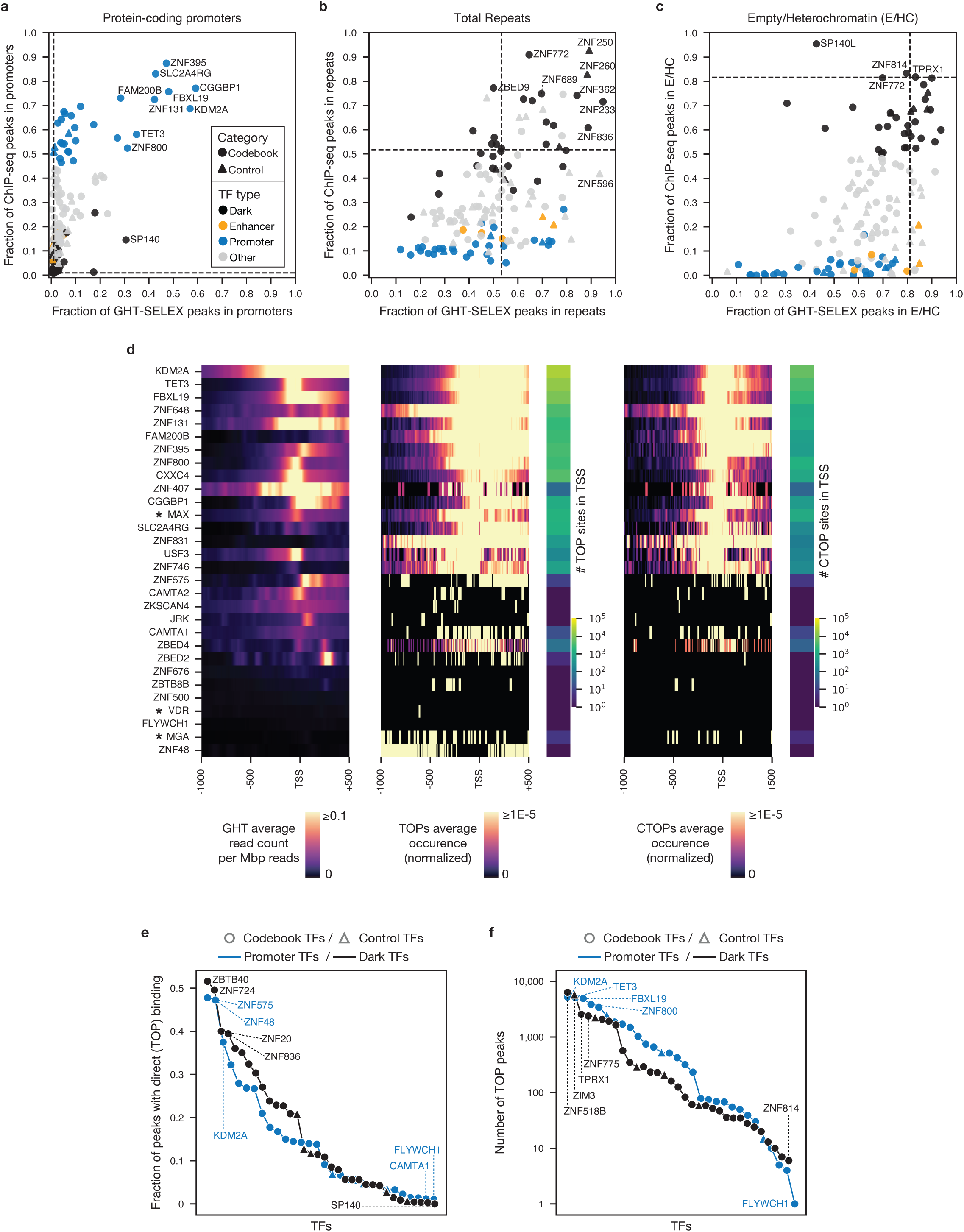
Evaluation of the genomic binding sites that are recognized by Promoter TFs, Enhancer TFs, and Dark TFs. (**a, b, c**) Fraction of GHT-SELEX (x-axis) and ChIP-seq (y-axis) peaks falling in the specified genomic regions; protein-coding promoters (**a**), repeats (**b**), and empty or heterochromatin (**c**), using the peaks at the universal threshold (MACS2.0 P<10^10^ for ChIP-seq and knee specificity of 30 for GHT-SELEX, see **Methods**), highlighting the extent of overlaps accounted by sequence specificity. Dashed lines show the expected fraction if peaks were distributed at random. (**d**) density of GHT-SELEX signal (left), TOP sites (middle), and CTOP sites (right) by position relative to TSS of protein-coding promoters, for 29 Promoter TFs that have available GHT-SELEX data. Intensity of heatmaps for TOPs (middle) and CTOPs (right) have been normalized by the total number of PWM hits (of TOPs and CTOPs, respectively) in promoters, shown by sidebars at the right of each heatmap. Control TFs are marked by asterisks. (**e**) and (**f**) compare the fraction and absolute number of peaks with direct binding (i.e. TOP sites) for Promoter TFs and Dark TFs respectively. In (**e**), for each TF, the number of TOP (i.e. triple optimized overlap) ChIP-seq peaks is divided by the total number of ChIP-seq peaks at the same optimized threshold (i.e. no overlapping). SP140 with no TOPs has been removed from (**f**).

In addition, dozens of Promoter TFs specifically displayed intrinsic preference for the regions that overlap or are just upstream of transcription start sites (TSS), by multiple measures (**Fig. 4d**, *left*), similar to that described for TFs in a variety of genomes^49–51^. This observation is consistent with functional roles for these TFs in promoter definition, delineation of TSS location, and/or gene regulation.

### Distinct conservation patterns of Dark TF vs Promoter TF binding sites

To examine the potential functionality of direct genomic binding sites, we examined the primary sequence conservation of sites that are bound directly. This analysis first requires the identification of high-resolution, high confidence binding sites, enabling conservation patterns to be attributed to individual bases within the binding site while excluding inconclusive relations arising from indirect TF recruitment. To make a conservative assessment of direct binding, we considered a ChIP-seq peak to be bound directly by a TF if the peak overlapped with a GHT-SELEX peak for the same TF as well as a high-scoring motif match. We obtained genomic scores for TF DNA binding motifs, modeled as Position Weight Matrices (PWMs), from Codebook consortium data; PWM derivation and benchmarking are described in more detail in accompanying manuscripts^34,40^. To further increase confidence in direct binding, we adjusted the significance thresholds for all three (ChIP-seq and GHT-SELEX peaks, and PWM hits treated as peaks) to maximize the Jaccard value between all three peak sets; we refer to these as “triple overlap” or TOP sites (see **Methods** for details). This approach maximizes the probability that ChIP-seq peaks are due to direct, sequence-specific binding; however, false negatives will arise due to experimental error of any type or inaccuracy of motif models. Thus, the numbers of direct binding sites obtained are likely to be underestimates. In addition, this analysis could be conducted only on 137 of the 217 TFs (101 Codebook TFs and 36 controls; see **Supplementary Table 3**) because 37% of the Codebook proteins lacked GHT-SELEX data, and/or did not yield motifs (**Supplementary Fig. 3a, b**).

For these 137 TFs, the fraction of ChIP-seq peaks that could be conservatively attributed to direct binding ranged from 0% for SP140 (indicating that its ChIP-seq sites are dominated by indirect recruitment) to 62.55% for ZNF728 (which has a unique 21-base motif), with a median of 8.5% (at the optimized threshold). The fraction is similar for the Promoter TFs (11.5%) and Dark TFs (11.1%) (**Fig. 4e**), and the absolute number of direct binding sites is similarly high for a subset of both groups (**Fig. 4f**). We conclude that there is no systematic difference between Promoter TFs and Dark TFs in direct binding characteristics, and that many of the observed TF binding sites are direct. The Promoter TF TOPs tend to be strongly aligned with TSS (**Fig. 4d**, *middle*).

We then examined the conservation of the TOP sites, producing an estimate of whether each site is under purifying selection. In essence, for many TFs, the TOP sites in aggregate display conservation patterns that mimic the selectivity of each base position in the TF’s PWM. **Fig. 5a** shows a graphic demonstration: when TOP sites are aligned to the PWM hit, and displayed as heatmaps that show base-level conservation scores (here, phyloP^52^), there are often vertical blue lines. These lines represent positions in the PWM hits that are preferentially conserved across many TOP sites. Similar to previous observations made with well-characterized TFs^52^, the positions with the highest conservation often correspond to tall letters in the sequence logo (i.e., high information content), indicating selection on the binding site to match the sequence preferences of the TF.

**Figure 5.**
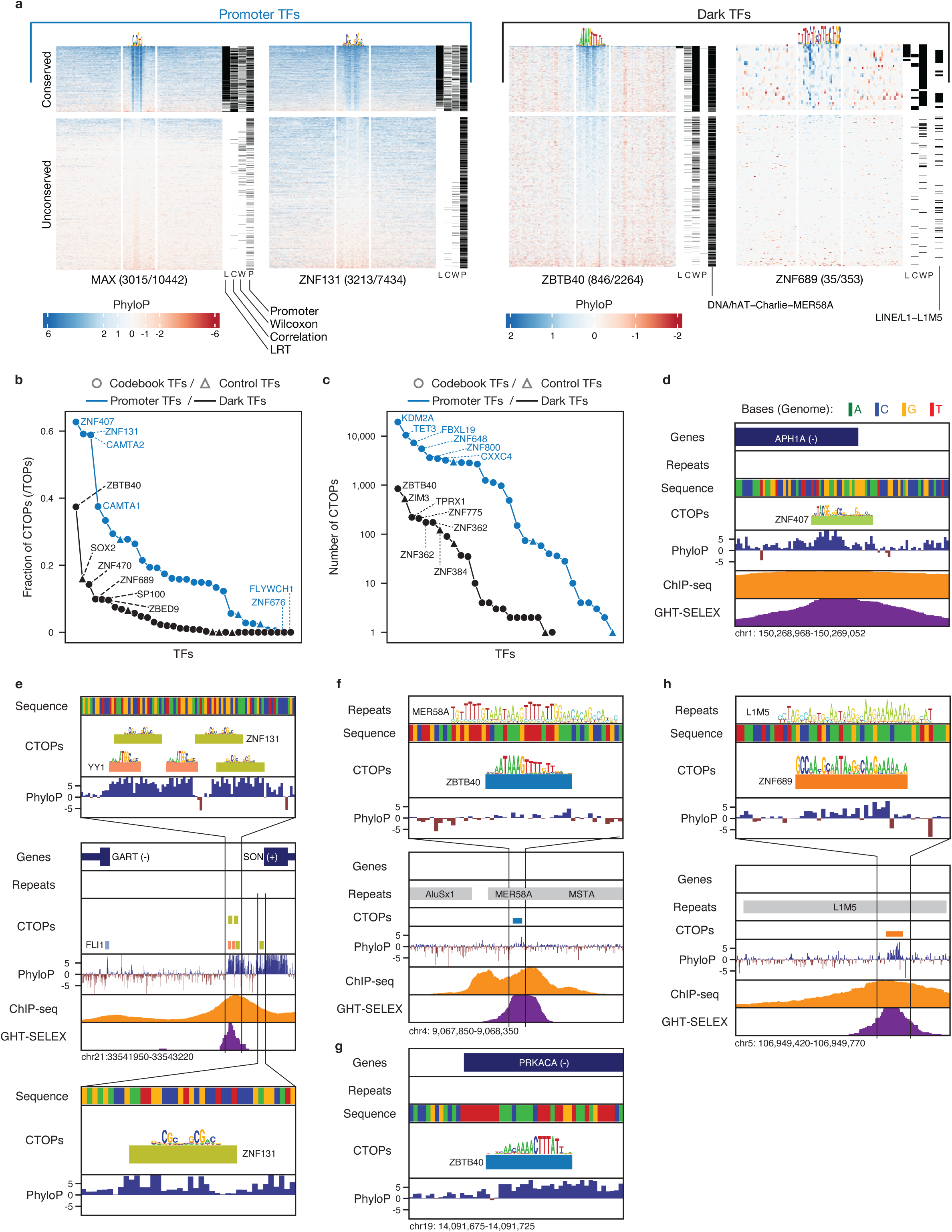
Conservation patterns of binding sites of Promoter TFs versus Dark TFs. (**a**) heatmaps of FDR-corrected phyloP scores across the TOP sites (rows) of two Promoter (MAX, ZNF131) and two Dark TFs (ZBTB40, ZNF689), split into top and bottom segments that contain conserved and unconserved sites. 100 bp segments are shown with the PWM hit in the middle. Blue/positive phyloP indicates purifying selection, and red/negative phyloP values represent diversifying selection. Bars to the right indicate which tests of conservation are satisfied (Likelihood-ratio, Correlation, Wilcoxon), along with overlaps with promoters (P) and specific repeat families if applicable. Comparison of fraction (**b**) and absolute number (**c**) of conserved TOPs (CTOPs), for Promoter TFs and Dark TFs. TFs with no CTOPs are not shown in (**c**). (**d, e, f, g, h**) Genome track alignments (genes, sequence, PhyloP, ChIP-seq and GHT-SELEX signals) of CTOPs for ZND407 (**d**), ZNF131 and YY1 (**e**), ZBTB40 (**f, g**) and ZNF689 (**h**). (**d**) is an example of a Promoter TF CTOP (ZNF407, which has the highest fraction of CTOPs) residing within the promoter of APH1A gene. (**e**) shows the four CTOPs of Promoter TF, ZNF131, colocalizing with YY1 in the prompter region of SON. (**f**) shows a CTOP of the Dark TF, ZBTB40, located within a *hAT/Charlie* (MER58A) element. ZBTB40 most-conserved TOP resides at the PRKACA promoter (**g**). (**h**) shows for another DARK TF, ZNF689, its CTOP is located within an L1M5 element. The repeat sequence logos shown in MER58A (**f**) and L1M1 (**h**) were produced in Dfam^91^.

We developed three heuristics to discriminate conserved vs. unconserved TOP sites (see **Methods**). Two of them test for a relationship between the information content at each base position of the PWM and the conservation score, while the third tests for higher overall conservation at the PWM hit than in the immediate flanking sequence. As shown in **Fig. 5a**, and in similar diagrams for all 137 TFs for which these tests could be run (**Supplementary Document 2**), these tests together detected sites that appear plausible by visual inspection (i.e., apparent conservation signal relative to flanks). We considered a site to be conserved if any of the three criteria were met, and at least one nucleotide in the PWM hit had an FDR-corrected phyloP score ≥1.

By these criteria, conservation of TOPs is observed for both Promoter TFs and Dark TFs (**Fig. 5a** and **Supplementary Fig. 8**), but the fraction and absolute number of conserved TOP sites for Promoter TFs are much higher (**Fig. 5b, c** and **Supplementary Table 3**). This outcome suggests that many Promoter TF binding sites are functional, and that the corresponding TFs have conserved functions at promoters. An individual conserved TOP site (hereafter, referred to as “CTOP” sites, **Supplementary Fig. 3c**), for the Promoter TF ZNF407, is shown in **Fig. 5d**; like many Promoter TF TOPs and CTOPs (**Fig. 4d**, *right*), it overlaps with a transcription start site. Promoter CTOPs are also often found adjacent to other CTOPs; an example of multiple sites for ZNF131 and YY1 is shown in **Fig. 5e**.

Despite their lower numbers, there are still thousands of CTOPs for Dark TFs: in aggregate, the criteria used here yielded 2,546 (**Supplementary Fig. 8** and **Supplementary Table 3**). They tend to be distant from promoters, or each other (e.g., 1,556 are > 1000 bp away from any other CTOP), and they tend to have lower phyloP scores than CTOPs for Promoter TFs. It is conceivable that there is lower statistical power to detect Dark TF binding sites because they typically overlap with TEs and are thus only present in species that contain the ancestral insertion. Nonetheless, striking examples emerged. The Dark TF ZBTB40 recognized 846 CTOP sites, the vast majority of which correspond to remnants of *hAT/Charlie* DNA transposons (**Fig. 5a, f**). Its most strongly conserved CTOP falls outside of a transposon, however, and instead is within the PRKACA 3’ UTR (**Fig. 5g**), which may be relevant to its apparent function in spermatogenesis (see below and **Supplementary Document 3**). The Dark TF ZNF689 has 35 CTOP sites (out of 353 TOPs) and is enriched for binding L1M5 elements (**Fig. 5a**; example CTOP shown in **Fig. 5h**). Overall, these analyses indicate that Dark TFs occupy a unique and expansive fraction of the genome, and thousands of their direct binding sites show indications of conserved function.

### Functions of Dark TFs

Finally, we considered potential functions of the Dark TFs. As a starting point, for three of the Dark TFs (SOX2, TPRX1, ZNF836), we examined their direct functional impact on chromatin by ChIP sequencing of H3K9me3 and ATAC-seq following ectopic induction in HEK293 cells (using the same transgene cell lines described earlier). SOX2 was included as a positive control. It is a well-known pioneer, but only 5% of its TOP sites overlap with ATAC-seq peaks in unperturbed HEK293 cells, while 66% of its TOP sites are in “empty” chromatin, which we reasoned might be impacted upon induction. TPRX1 and ZNF836 were chosen because of their relatively low endogenous expression in HEK293, yet like SOX2, most of their ChIP-seq binding sites are in “empty” or heterochromatin. In addition, TPRX1 has recently been described as a master regulator in zygotic genome activation^53^, consistent with potential pioneer activity, while ZNF836 is a KZNF that binds the MaLR LTR both *in vitro* and *in vivo* (see **Fig. 3b**).

These experiments revealed widespread impact of these TFs on chromatin at their genomic binding sites (TOPs). The ATAC signal at SOX2 TOPs increased by 2.5-fold on average, with most TOP sites clearly impacted (**Fig. 6a**). TPRX1 increased ATAC signal at its TOPs by 1.4-fold, overall, although the impact is concentrated on ∼10% of its 3,882 TOPs (**Fig. 6b**), suggesting that additional factors are required for TPRX1 to initiate chromatin opening. ZNF836, in contrast, strongly increased the H3K9me3 signal over its binding sites, with less impact at sites that already reside in heterochromatin prior to the induction (**Fig. 6c**). We conclude that at least a subset of Dark TFs can readily influence the chromatin state at their binding loci.

**Figure 6.**
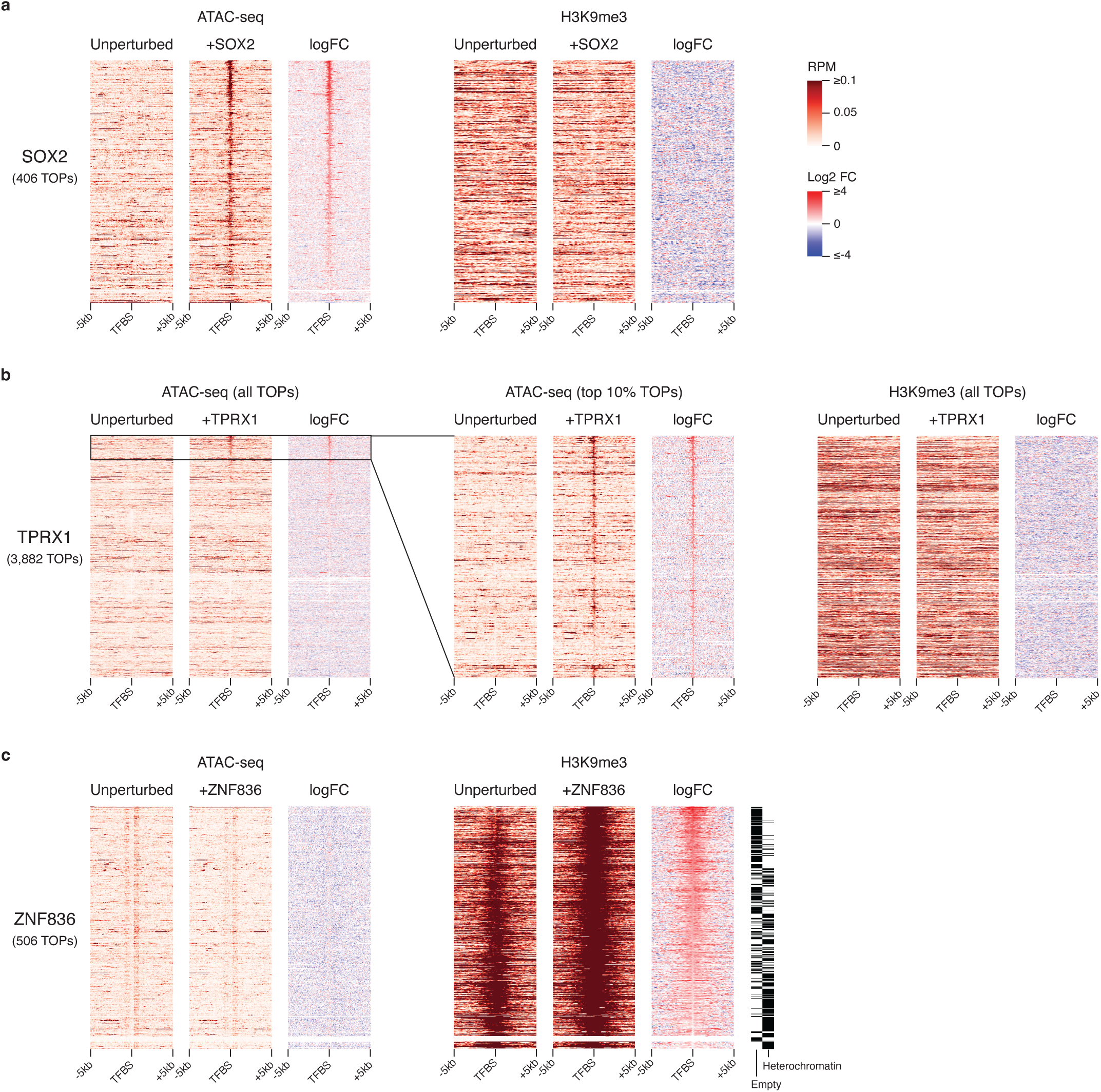
Chromatin reorganization by Dark TFs and their pioneer activity at TOP sites. Comparison of ATAC-seq and H3K9me3 ChIP-seq signals in unperturbed HEK293 cells with three over-expressed Dark TFs SOX2 (**a**), TPRX1 (**b**), and ZNF836 (**c**), measured in a window of 10 kb around their TOPs. For ATAC-seq (left panels), heatmaps show the pileup of Tn5 tag insertion sites (reads per million; RPM) extended to 50 bp on each side. Right panels show the heatmaps for the pileup of H3K9me3 ChIP-seq fragments (RPM). The logFC heatmap illustrates log₂(TF overexpressed/unperturbed HEK293). For TPRX1 ATAC-seq, the heatmap of its dominant 10% TOPs is enlarged in the middle panel for clarification. For better visualization, SOX2 and TPRX1 TOPs in all panels are sorted by their ATAC-seq log₂FC at the TFBS center, while ZNF836 TOPs are sorted by H3K9me3 signal.

We also examined existing literature and databases to survey known and potential functions for the Dark TFs in relation to our new finding **Fig. 7** provides a graphical overview of various properties of these proteins such as their interactions with chromatin remodelers and cofactors, properties of their binding sites, and previously-described biological functions (see **Supplementary Document 3** for a detailed description, and **Supplementary Table 4** for a table of known protein-protein interactions and other information). In summary, most Dark TFs have apparent roles in repression of transcription, via at least six different mechanisms that are different from recruitment of KAP1^33,54^. Intriguingly, 22 of the 47 C2H2-zf proteins, including both KRAB and non-KRAB C2H2-zf proteins (as well as transposon-derived ZBED9, SOX2, and SP100), interact with other C2H2-zf proteins, often extensively^55^, suggesting a potentially widespread role in the organization of chromosome topology. Not all Dark TFs are apparent repressors, however. In addition to TPRX1, SALL3 is a known master regulator, controlling the differentiation of hiPSCs into cardiomyocytes vs neural cells^56^. Two other Dark TFs have been described as impacting DNA metabolism; ZNF384 binds Ku and recruits non-homologous end joining factors to double-strand breaks^57^, while ZNF146 depletion slows the replication fork^58^.

**Figure 7.**
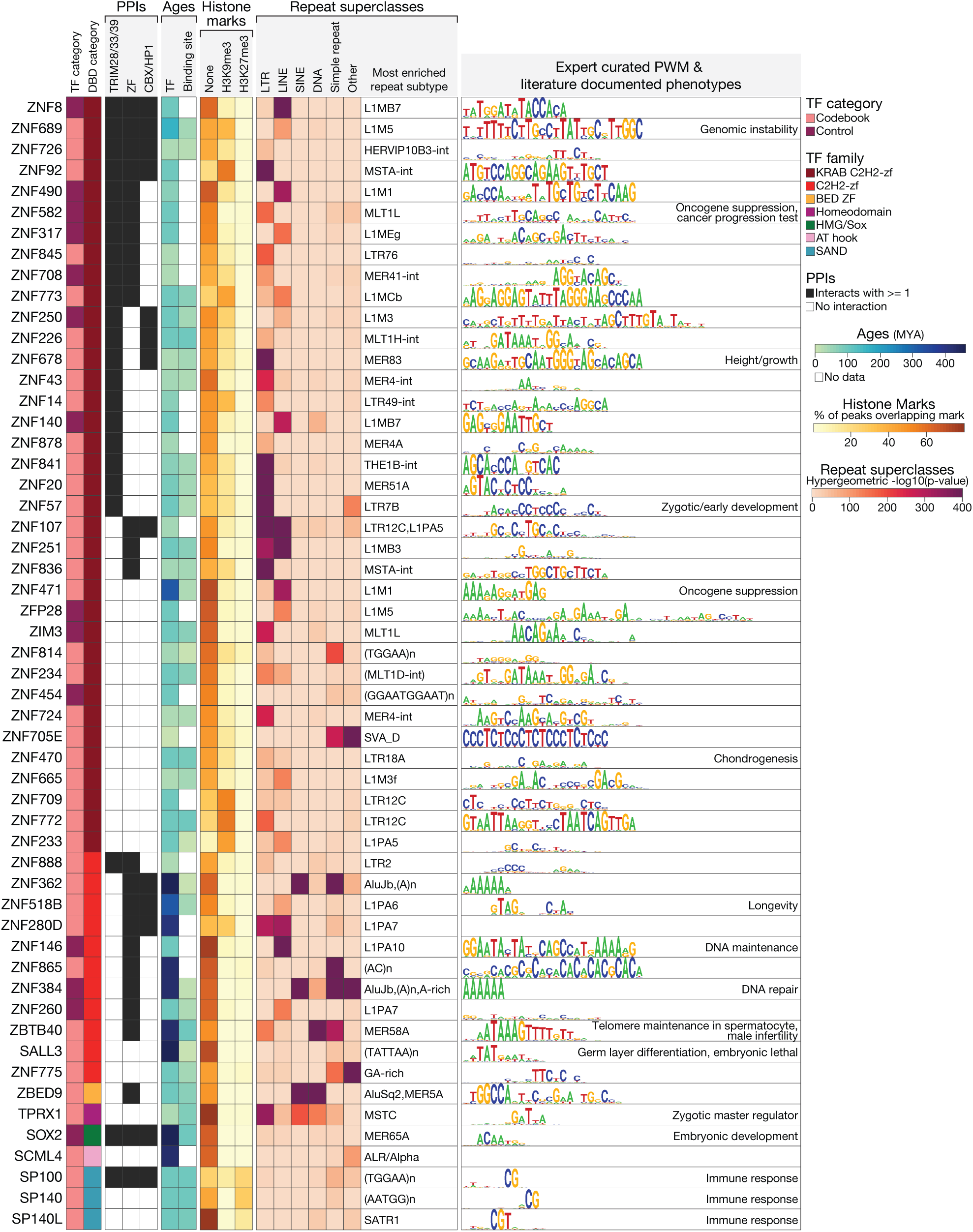
Consolidated information for functions and other properties of Dark TFs. Compiled protein-protein interactions(PPIs)^92^ information obtained from available studies and grouped into three categories with potential regulatory functions: TRIM28/33/39 interactions, zinc-finger (ZF) protein interactions, and CBX/HP1 interactions are shown at left. Median binding site age was calculated for TOP sites, only for the TFs with available GHT-SELEX data, shown along with the age of the TF. The fraction of ChIP-seq peaks (using the universal threshold) overlapping with H3K9me3 and H3K27me3 histone marks and with the ChromHMM “empty” (or None) state are shown in the middle. For the repeats, in each superclass, the enrichment score (−log(p-value) hypergeometric test) for the most enriched repeat element within that superclass is plotted as a heatmap, and the most enriched repeat subtype across all the superfamilies is mentioned beside. The expert-curated sequence logos are displayed to the right (except for ZNF280D and SCML4 which did not produce any approved PWMs), along with the corresponding phenotype for any TF with known biological function through literature review (in the same block).

The distinct binding sites and diversity of apparent effector mechanisms and cellular roles of the Dark TFs suggest that they may each regulate specific biological functions, and that they may also be multifunctional. Indeed, physiological consequences that have been reported for perturbation of the Dark TFs vary widely (**Fig. 7**, right column; **Supplementary Table 4** provides the values and sources), ranging from basic cellular processes to development. For example, the KZNF ZNF689, shown in **Fig. 5a** to bind 35 conserved sites enriched for L1M5 (**Fig. 5g**), also binds promoters of various L1 subtypes, preventing genomic instability conferred by L1 retrotransposition^59^. ZBTB40, which we observe almost exclusively at DNA *hAT/Charlie* transposons (**Fig. 5e**), and which is one of the oldest Dark TFs (**Fig. 7**), was recently shown to bind telomeric dsDNA breaks and maintain telomeric length^60^. In mouse, Zbtb40 deficiency impacts spermatogenesis through disrupted telomeric lengthening and maintenance in spermatocytes^61^. *hAT/Charlie* transposons are enriched in telomeric regions of human DNA^62^, suggesting that this function may be conserved.

### Complex patterns of coevolution among Dark TFs and their binding sites

Dark TF binding sites typically feature specific classes of TEs, which are the youngest insertions in the genome. The Dark TFs themselves are dominated by KZNFs, which are the youngest class of TFs in human. TPRX1, a homeodomain protein, is also relatively recent (it appears to be a eutherian-specific duplicate of the adjacent CRX gene^63^). KZNFs both bind and co-evolve with TEs they suppress but are also assumed to adopt functions beyond TE silencing. Given these clear observations and expected tendencies, we asked whether we could systematically derive evolutionary principles that would characterize the Dark TFs and their binding sites, relative to other TF classes. To do this, we compared the estimated ages of the TFs to the ages of their binding sites, which we defined by seeking ungapped alignment in ancestral genomes, regardless of sequence conservation, or whether the sequence is a TE (**Supplementary Fig. 9a**). Simultaneously, we considered whether the binding sites of TFs within TEs suggest either a TF-specific evolutionary history or reflect more general trends (e.g., TFs binding TEs of the same age, or specific classes of TEs being predominant).

This analysis is described in detail in **Supplementary Document 4.** No obvious systematic patterns emerged that could be described as rules, but a wealth of tendencies and specific observations were revealed. First, regardless of TE binding, or TF class, a clear trend is that TFs tend to be older than the sites they bind: Promoter TFs have a median age of 429 MYA, while Dark TFs have a median age of 99.19 MYA (**Supplementary Fig. 9b**), which is still older than a typical binding site even for Promoter TFs, and older than most human TEs (**Supplementary Fig. 9c**). These results suggest that the established phenomenon of TF binding site turnover^14^ is prevalent for the Dark TFs, just as it is for other TF classes. Second, TEs of all major classes (LINE, SINE, LTR/ERV, and DNA transposons) are recognized by specific TFs. Moreover, for all four major classes of TEs, there are cases in which greater than 10% of a TF’s TOP sites overlap one TE type and are conserved (**Supplementary Fig. 10a**). These are all apparent examples in which the corresponding TF/TE combination has systematically contributed to the regulatory function of the host genome.

Third, the encompassed TEs span a very wide age range, from human-specific AluY elements to L2, L3, and MIR, which pre-date eutherian mammals. These associations can provide insight into the evolution and molecular function(s) of specific TFs. For example, ZNF836 and ZNF841 (which are both Dark TFs and KZNFs) are paralogs that arose from a pre-simian duplication event^64^ and bind to distinct subtypes of the closely related, simian-specific MaLR LTR elements. They bind distinct motifs that specify the differing base identities at homologous positions within the diverged LTR, suggesting neofunctionalization and retention to maintain silencing of both LTR subtypes (**Supplementary Fig. 10b**).

There are also many cases in which the TFs and TEs they bind are grossly mismatched in age, however, suggesting that TE binding is fortuitous. For example, ZNF286B is a human-specific duplicate of ZNF286A which has lost its KRAB domain^65,66^, but its binding sites are enriched for LINE-3 (L3), an ancient element found across all mammals, suggestive of coincidental adaptation (**Fig. 3b**). In contrast, ZNF362 and ZNF384 (both non-KZNFs) are products of a duplication ∼429 MYA (the duplication is found across bony vertebrates), but the binding sites for both proteins are enriched for the much younger, primate-specific Alu elements, as well as poly-A repeats, consistent with their DNA binding motifs (**Supplementary Fig. 10c**). These proteins have the largest number of TOP sites within the entire Codebook dataset, and it is possible that the recently-expanded target range of these proteins is a coincidental liability, as rearrangements of both ZNF362 and ZNF384 genes (most commonly as fusion proteins with activating TFs and cofactors) are found frequently in leukemia^67,68^.

## DISCUSSION

The Codebook ChIP-seq data provide cellular binding sites for 130 putative TFs, defined as previously lacking PWMs or other models of sequence specificity. It represents a valuable resource for studying TF function and evolution in the context of regulatory genomics. For a large majority of the proteins assayed, we have also now identified a binding motif which is supported by independent assays^40^. Thirty-six of the proteins did not produce a motif and may not be *bona fide* TFs. Their ChIP-seq profiles are nonetheless informative: enrichment of ChIP-seq peaks at different types of genomic features (e.g. promoters, repeats) or chromatin states, as well as co-occurrence with peaks for other proteins (e.g. TFs), can yield clues as to potential function.

Previous large-scale ChIP-seq analyses have mainly focused on the established roles of TFs in binding to promoters and enhancers (e.g. Partridge *et al.*^24^), although studies of KZNFs typically focus on binding to TEs, and specifically EREs^7,16,33^. The analysis scheme described here considers these models of TF function as hypotheses with equal weight. The expected binding categories are clearly present, including preferences for promoters and enhancers, as well as the strong tendency for KZNFs to bind specific classes of EREs, and within constitutive heterochromatin. Binding to EREs is not the only function of KZNFs, however, and other TF classes can clearly bind genomic dark matter. Just over half (36/63; 57%) of the KZNFs we analyzed are Dark TFs - as are roughly 10% of non-KZNF control TFs (4/35). If these proportions were to hold for the 350 human KZNFs and the 781 non-KZNF TFs with a known DNA-binding motif, then there would be a total of ∼277 human Dark TFs among the ∼1,600 human TFs (i.e., 17% of all human TFs), among which roughly a third would not be KZNFs.

Overall, TF behavior with respect to chromatin states and genomic landmarks appears more varied than a simple categorization scheme would imply. We did not systematically explore the “Other” category, used here as a catch-all. However, like the Dark TFs, Other TFs appear to encompass proteins that bind both *in vitro* and *in vivo* to many regions that are labeled as inactive and/or closed chromatin in HEK293 cells prior to induction of the TF, and to satisfy some expectations of “pioneer” activity. One example is the non-KRAB protein ZSCAN2, for which we cataloged 90 CTOPs and found they are enriched for mammal-wide LINE-3 (L3) elements. ZSCAN2 is shown to be expressed only in early stages of embryogenesis as well as post-meiotic round spermatids^69^, and to affect fertility in mice (and perhaps human)^70^.

Establishing physiological functions, if any, for individual TF binding sites is a long-standing and difficult problem in regulatory genomics. It is becoming clear that even KZNFs exert specific physiological outcomes, beyond ERE repression^22^, but few are currently known. Biochemical functions of TFs may be more straightforward to identify: our observation of ATAC signal enrichment within TPRX1 TOPs upon ectopic expression of the TF suggests a straightforward approach. Biochemical function does not equate to physiological purpose, however. A particular challenge with TFs is that the level of binding site turnover observed on evolutionary timescales requires that binding sites arise at random, many of which are presumably irrelevant for gene regulation or reproductive fitness (at least initially), despite being biochemically functional. By this reasoning, we expect that many biochemically verified, direct TF binding sites should be physiologically non-functional, and indeed, we find that, overall, most TOP sites are not conserved, even for Promoter TFs. Lack of conservation cannot be taken as lack of biological purpose; nonetheless, conserved TOP sites would seem most likely to yield interpretable results in targeted laboratory studies. More generally, the Codebook TOP and CTOP catalogue will provide a rich resource for future efforts in examining genome function, e.g., employing techniques similar to GTEx^71^ to relate variants at the TOP sites to chromatin and gene expression.

The lower overall primary sequence conservation of Dark TF binding sites, relative to those of Promoter TFs, could have several explanations. One possibility is that very few of the binding sites are functional; in theory, only a single binding site that confers a modest selective advantage would be sufficient to drive retention of both the site and the TF, with all other sites arising at random (and under neutral evolutionary pressure, provided they are not detrimental). Another possibility is that the exact positioning of the sites is not critical to their function, unlike Promoter TFs, which by definition must be close to TSSs. Dark TF functions could simply require that binding sites are distributed widely across non-functional DNA and thus could be redundant over large sequence windows. Such functions would not preclude a small subset of sites being co-opted for regulation of host genes, which would become constrained (i.e., conserved). Regardless of what biochemical, cellular, and physiological functions are revealed, the Dark TFs represent a new contribution to the decades-old odyssey into the function and significance of the dark matter genome.

## METHODS

### Plasmids

The Codebook project design is described elsewhere^34^. Putative TFs were those from Lambert et al.^23^, with TFs we had already attempted as part of ENCODE removed. We attempted ChIP-seq analysis of all Codebook TFs. We designed full-length ORFs for synthesis (BioBasic.ca) and used conventional restriction cloning to insert them into Flp-In destination plasmid pTH13195 (a modified pDEST pcDNA5/FRT/TO-eGFP vector), which places ORFs under the control of a tet-on, CMV-driven promoter, with an N-terminally EGFP tag. We obtained the 58 controls independently^33,38^ and cloned them into pTH13195 by the same process. See accompanying manuscript^34^ for the sequences and other information about the inserts.

### Cell line production

We used a previously established protocol for creating individual cell lines for each TF^16^. In brief, we cultured and maintained HEK293 Flp-In T-REx cells (Invitrogen) in Dulbecco’s modified Eagle’s medium with 10% fetal bovine serum and antibiotics. We created individual cell lines for each TF (between one to five cell lines per construct as biological replicates, see **Supplementary Table 2**) by flp recombination, in which individual destination plasmids were co-transfected into Flp-In T-REx 293 cells together with the pOG44 Flp recombinase expression plasmid using FuGene (Roche, 11814443001). We then selected cells for FRT site-specific recombination into the genome using selection media containing hygromycin (200μg/ml) for 1 to 4 weeks. For each TF cell line, we confirmed expression of EGFP by fluorescent microscopy after 24 hours of Doxycycline treatment (1ug/ml), at which point 10M-20M cells were used for downstream experiments. Individual cell lines were immunoprecipitated between one to five times, in different batches. For each batch of ChIP-seq experiments, we also collected “input DNA” (i.e., non-IP sonicated chromatin), chosen at random for ∼10% of all the experiments (see **Supplementary Table 2**).

### Chromatin immunoprecipitation

We fixed ∼20M cells on 15cm plates using 1% paraformaldehyde for 10 min on ice, followed by 10 min quenching with 0.125 M glycine. We washed fixed cells twice with cold PBS, scrape-collected, pelleted, flash-froze, and stored the cells at −80 °C. Upon completion of cell collection for a panel of TFs, we thawed cell pellets on ice, lysed them as previously described^72^, and sonicated them using a BioRuptor to shear chromatin. We then trapped the protein-DNA complexes on Dynabeads using an anti-EGFP or anti-H3k9me3 (Ab290 and Ab8898, respectively, Abcam). Following wash, crosslink reversal, and elution, we assessed the size and concentration of DNA fragments with an Agilent bioanalyzer and Qubit, prior to sequencing.

### Library preparation and sequencing

DNA library preparation and sequencing were performed at three different facilities (Memorial Sloan Kettering Cancer Center, SickKids Hospital in Toronto, and the Donnelly Centre at the University of Toronto) over a period of four years. The facilities prepared DNA libraries using the NEBNext® Ultra™ II DNA Library Prep Kit for Illumina. Samples were paired-end sequenced (50-150bp) with a target depth of 20M reads (for EGFP) and 50M reads (for H3K9me3) per sample. 183 samples have two sequencing replicates (specified as D and M replicates in **Supplementary Table 2** under the Technical Seq. Rep # column).

### TF ChIP-seq data preprocessing

We mapped raw ChIP-seq reads to the human genome build hg38 with *bowtie2*^73^ (options: *--very-sensitive*, and *--no-unal*). We used Samtools^74,75^ (options: *-q 30*, and *-F 1548*) to remove reads that were unmapped, failed platform/vendor quality checks, were PCR duplicates, or had a mapping quality <30. This stringent threshold on mapping quality ensures that reads mapped to multiple locations are removed; hence, the peak locations (see below) are unambiguous, and our analysis of repeat elements is not skewed. This elimination may, however, result in missing potential peaks in young repeat families with identical or near-identical copies (e.g., L1HS). This could lead to an underestimation of the proportion of binding sites in the dark genome, as repeat elements are often found in those regions, but it would only result in false negatives.

### TF ChIP-seq background generation and peak calling

We generated sample-specific background models using a previously established approach^33^, with minor modifications. For each ChIP-seq pull-down experiment, our goal was to construct a background control that closely matched its nonspecific signal while maintaining high coverage by pooling reads from multiple “input” experiments in proportions that best reproduced the background characteristics of the given pull-down.

First, we identified genomic regions showing high background signal in at least one input DNA experiment. To do so, we performed peak calling on each input sample using MACS2 (options: p-value < 0.001, and *--nomodel*)^43,76,77^ and collected the resulting peaks as “background hotspots”. We merged hotspots whose summits were within 50 bp across experiments to produce a unified set of background regions.

Next, we quantified read counts overlapping each hotspot. For each input DNA experiment, we counted the number of reads overlapping every region in the unified hotspot set, creating a read count matrix with rows being the hotspots and columns being input experiments. For each pull-down experiment, we also calculated read counts over the same hotspot regions; these served as the response variable in the following modeling step. We modeled the pull-down read counts using a non-negative Poisson regression, with the input DNA read count matrix as predictors. This regression estimated a set of non-negative coefficients (one coefficient per input experiment) representing the weighted combination of controls that best reconstructs the background signal observed in the pull-down.

Finally, we used these coefficients to generate a synthetic background dataset. Specifically, we sampled reads from each input BAM file in proportion to its corresponding regression weight and pooled them to form a single, pulldown-specific control file. This customized control was then used as the background input for peak calling on the pull-down sample with MACS2^43,76,77^.

### ChIP peak replicate analysis and merging

For each TF with one or more replicates, we calculated the Kulczynski II similarity metric for each pair of replicates (**Supplementary Fig. 2a**). We used the Kulczynski II metric in place of Jaccard as it is less affected by the uneven size of the peak sets. We additionally calculated the Kulczynski II similarity metric for each pair of mismatched replicates (i.e., with TF identities permuted). Based on the distributions of “approved” experiment replicates and mismatch replicates, we defined a Kulczynski II threshold of 0.4 as the separating value for those two distributions (**Supplementary Fig. 2a**). For TFs with “not approved” experiments (i.e., two ChIP-seq experiments did produce a reliable motif), we retained 36 (plus two controls) that achieved a Kulczynski II value >0.4 for inclusion in downstream analyses.

To generate a single peak set for each transcription factor, we merged the peak data from all successful experiments by merging overlapping peaks from one or more replicates using BEDTools^78^ merge to generate new, wider peaks, with the sum of component peak −log(p-values) assigned as the new peak score, and the center of mass of summits as the new peak summit. By default, we employed a peak cutoff of P<10^-10^ (MACS2). Modulation of thresholds is described below.

### H3K9me3 ChIP-seq data processing

Adapter sequences were first trimmed using *cutadapt*. The resulting reads were then mapped to the human genome (hg38) using *bowtie2*^73^, followed by the creation of BAM files using *samtools*^74,75^ *view*, and sorting with *samtools sort*. Unintended or duplicated reads were removed using *sambamba view -h -f bam -F “[XS] == null and not unmapped and not duplicate and proper_pair and mapping_quality >= 30”* and *picard MarkDuplicates*. Pileups (bdg) for pulled down fragments were obtained using *macs3*^43,76,77^ *callpeak* with the options *--format BAMPE - g hs --scale-to small –bdg* using all the replicates of that experiment for option *-t*, followed by *bedGraphToBigWig*. Finally, to get the signal over TOP sites, DeepTools^79^ *computeMatrix* was used, and plotted by custom Python scripts.

### ATAC-seq experiment

We performed ATAC-seq in HEK293 cells in two separate sets. The first set included only unperturbed HEK293 cells with four replicates in order to establish the baseline status for chromatin state (see Chromatin state analysis section below). The second set included three HEK293 cell lines (SOX2, TPRX1, and ZNF836) with two biological replicates for each TF. We performed the well-established OMNI-ATAC protocol as previously described^80^. Briefly, 50,000 viable cells were pelleted (500 RCF at 4°C for 5 min) collected and palleted in 1ml cold PBS After removing the supernatant, the cells were lysed in 50 μl of cold ATAC–Lysis buffer containing 0.1% NP40, 0.1% Tween 20, and 0.01% digitonin by pipetting up and down three times followed by 3 min incubation on ice. 1 ml of cold ATAC-resuspension buffer containing 0.1% Tween 20 was then added to the lysate followed by 10 min centrifugation in 4°C at 500with to pallet the nuclei Transposition was performed using either Tagment DNA TDE1 Enzyme and Buffer Kit (Illumina) or loaded Tn5 enzyme and 2x tagmentation buffer (Diagenode, C01070012 and C01019043) in 50μl volume of transposition mixture that was prepared according to the manufacturer’s protocol. After mixing well by pipetting up and down, the mixture was incubated for 30 min at 37 °C. The tagmented DNA was then purified with DNA Clean and Concentrator kit (Zymo Research) in 21 ul of elution buffer. DNA amplification and barcoding were performed using either Nextera DNA Library Prep kit (Illumina, for unperturbed HEK293 replicates) or Diagenode’s tagmentase compatible primers “24 UDI for tagmented libraries sets (I, II, III)” (C01011034, C01011036, and C01011037, for the three TFs as well as two unperturbed HEK293 replicates) with an average number of 10 cycles per sample. Subsequent sequencing was performed at the Donnelly Center sequencing facility using 100bp paired end sequencing at 60M reads per sample (for unperturbed HEK293 replicates in the first set) or at the McMaster Genomic facility (McMaster University) using 50bp paired end sequencing at 60M reads per sample (for SOX2, TPRX1, ZNF836, and unperturbed control HEK293 in the second set).

### ATAC-seq data analysis

Adapter sequences were first trimmed using *cutadapt* to remove the Tn5 adapter CTGTCTCTTATACACATCT. The resulting reads were then mapped to the human genome (hg38) using *bowtie2*^73^, followed by the creation of BAM files using *samtools*^74,75^ *view*, and sorting with *samtools sort*. Unintended or duplicated reads were removed using *sambamba view -h -f bam -F “[XS] == null and not unmapped and not duplicate and proper_pair and mapping_quality >= 30”* and *picard MarkDuplicates*. The first set of ATAC-seq experiments went through peak calling on the sorted BAM reads by running *macs2*^43,76,77^ *callpeak* with the options *-f BAMPE, -g hs, -B*, and *-q 0.01*. Then, to generate a single peak file for HEK293 open chromatin, all the peak files were merged using *bedtools*^78^ *merge*. For the second set of experiments (SOX2, TPRX1, and ZNF836), the BAM files were first adjusted by *<u>alignmentSieve – ATACshift</u>* to eliminate the Tn5 bias by +5bp (+ strand) and −4bp (-strand). In order to make pileups (bdg) for the Tn5 tag locations, *macs3*^43,76,77^ *callpeak* was used with the options *--format BAM -g hs --nomodel --shift −50 --extsize 100 --keep-dup all --scale-to small –bdg* using all the replicates of that experiment for option *-t*, followed by *bedGraphToBigWig*. Finally, to get the signal over TOP sites, we used DeepTools^79^ *computeMatrix*, followed by a custom Python script for plotting the fold change.

### Chromatin state analysis

We obtained chromatin states by training a ChromHMM^45^ with ten states (see **Supplementary Fig. 4a**) on marks H3K4me1, H3H4me3, H3K36me3, H3K27ac, H3K9me3, and H3K27me3, collected in HEK293 cells by ENCODE^81^, plus ATAC-seq and CTCF peaks from HEK293 cells generated as part of this project. We also employed promoter regions derived from the GENCODE annotation (release 44)^82^ for our analyses (−1000 to +500). For Hi-C B compartment annotations, we labeled genomic regions with a Hi-C first eigenvector value less than 0.4 in ENCODE data for HAP1 cells^83^, comprising 65% of the genome.

### Overlap of ChIP-seq peaks between all pairs of TFs

Jaccard similarity is taken as O / (N1+N2-O), where O is the number of intersecting peaks and N1 and N2 are the sizes of each set. We utilized BEDTools^78^ to calculate overlaps. To prevent miscounting of the cases in which one peak in one set overlaps with multiple peaks in another set, we used the average of overlapping peaks (O = (O1+O2)/2 where O1 is the number of peaks in set 1 overlapping with any peak in set 2 and vice versa) to calculate the intersection in Jaccard. The same methodology was used to calculate the overlap of ChIP-seq peaks with the chromatin tracks.

### Selection of universal ChIP-seq and GHT-SELEX thresholds

We calculated the Jaccard similarity from all 137 pairs of TFs with ChIP-seq and GHT-SELEX data, using the merged ChIP-seq peaks. We performed a grid search for all TFs simultaneously, sampling ChIP-seq P-value and GHT-SELEX cutoffs (determined by selecting different “knee” values^84^ in the graph of sorted enrichment coefficients^48^), to identify a pair of thresholds that maximizes median Jaccard. Two ChIP-seq cutoffs (10^-10^ and 10^-20^) yielded an almost identical maximum; we chose 10^-10^ as it includes a larger number of peaks. A corresponding knee specificity threshold of 30 emerged for the GHT-SELEX knee-based cutoff.

### Filtering peaks in genome unmappable regions

Before comparing ChIP-seq and GHT-SELEX peak sets to define TOPs (see **Derivation of TOP sites** below), we first filtered out the peaks overlapping with genomic regions that are problematic for any of the assays. The following sources were used to make a “genomic whitelist” for our study:

1. hg38 36-bp mappable regions on human genome: Bismap^85^, downloaded from UCSC: https://genome-euro.ucsc.edu/cgi-bin/hgTables?db=hg38&hgta_group=map&hgta_track=umap&hgta_table=umap36&hgta_doSchema=describe+table+schema. 36-bp was chosen to match GHT-SELEX fragment length.
2. Codebook GHT-SELEX-covered regions: all the genomic regions with at least one read coverage in any GHT-SELEX experiment across the entire study.
3. Codebook ChIP-seq-covered regions: all the genomic regions with at least one read coverage in any ChIP-seq experiment across the entire study.
4. ENCODE blacklist v2.

The Codebook whitelist was then made by intersecting the first three tracks and the complement of the fourth. Any peak (ChIP-seq, GHT-SELEX, or PWM-predictions) with less than 25% of its length overlapping with the whitelist was removed from the further steps.

### Derivation of TOP sites

To define binding sites supported by ChIP-seq, GHT-SELEX, and PWM hits, we optimized the cut-offs of all three to maximize the overlap between all three data types. We first sorted the peaks based on their statistical scores, i.e., merged p-values for ChIP-seq peaks, cycle enrichment coefficient for GHT-SELEX peaks (see accompanying manuscript^48^), and sum-of-affinities for clusters of PWM hits with a p-value < 0.001 (from MOODS^86^), merged with neighboring hits in the case of having a distance less than 200 bp. Then, for different values of *N*, we took the top *N* peaks and calculated the overlap (measured as the Jaccard index; intersection of all three divided by the union of all) using the top *N* ChIP-seq peaks, top *N* GHT-SELEX peaks, and top *N* merged PWM hits. The *N* that maximizes the Jaccard overlap (for *N* between max(100, 0.01 x n_max) and n_max) was taken as the optimized threshold, and the overlap of all three sets at this threshold (*N*) is referred to as triple overlap or “TOP” sites.

### Analysis of purifying selection and classification as conserved and unconserved binding sites

We extracted phyloP scores^52^ for each PWM hit, and for flanking regions of equal length (for a total of 100 bp including the PWM hit and its flanks) from the 241 eutherian mammal Zoonomia alignment^87^ using DeepTools^79^. We excluded PWM hits overlapping with ENCODE Blacklist sites^88^ or protein coding sequences, due to the skew in phyloP scores caused by codons. All phyloP scores are FDR-corrected. We conducted three tests to classify PWM hits as ‘conserved’ or ‘unconserved’:

1. <u>LRT (Likelihood-Ratio Test)</u>: This test scores the likelihood that the phyloP scores are driven by the PWM information content (IC) at each base position in the PWM. For each TF, we created a scoring model that represents the relationship between the phyloP scores at a PWM hit and the information content at each base position of the PWM. This model is an l x 1 vector, where l is the length of the motif. To derive this vector, we first took the correlation of phyloP scores at each base position within the PWM hit to the IC at that position, for each PWM hit in the TOP dataset. We then selected the 100 PWM hits with the highest correlation and calculated the standard deviation (***σ***) of the phyloP score at each position of these 100 PWM hits. If a position has an invariant phyloP score (i.e. *σ* = 0), the ***σ*** at this position was replaced with a 1. As a null model, an IC value of 0 was assumed at each position, and the same ***σ*** values as the phyloP model. The LRT statistic for each PWM hit *m* was then taken according to the equation:

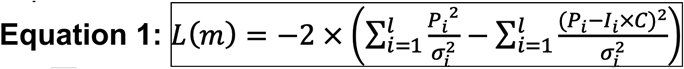 Where ***p_i_*** represents the phyloP score of position *i* in PWM hits *m*, *σ_i_* is the standard deviation of the phyloP model at position *i*, and ***I_i_*** is the IC of the PWM at position *i*. The IC value is first multiplied by a coefficient ***C***, which is the linear regression coefficient describing the relationship between the position-wise phyloP means of all of a TF’s TOPs and position-wise PWM IC. It therefore has units phyloP/bits and converts ***I_i_*** to a phyloP score. Based on manual inspection of phyloP patterns across TOPs at different test statistic thresholds, we selected a threshold of L < −10 to be considered “conserved” according to this test, which manual inspection indicated is conservative.
2. <u>Correlation Test</u>: For each TF, we permuted the position-wise IC of the PWM using the *permute* R package, up to a maximum of 1,000 unique permutations (not every PWM has 1,000 unique permutations). We then took the Pearson correlation of each of these permuted PWM IC vectors using the phyloP scores of 1,500 randomly selected PWM hits from the unfiltered PWM scan results (or fewer, if there are <1,500 total hits). This resulted in a maximum of 1,500,000 correlations per TF, dependent upon the number of unique PWM IC permutations and the number of PWM hits. We used these correlation values as a null distribution, converted to Z scores, and determined a threshold correlation value corresponding to an alpha of 0.05; this threshold was chosen manually. PWM hits with a Pearson correlation to the unpermuted motif IC values greater than this threshold were considered to have passed this test.
3. <u>Wilcoxon Test</u>: For each TOP site, we performed two one-sided Wilcoxon tests, one comparing values in the PWM hits to those in the 25bp downstream flank, and the same for the 25bp upstream flank. The alternative hypothesis was that the PWM hit phyloP values were greater than the compared flank. All p-values were FDR corrected, and an FDR-corrected p-value less than a threshold of 0.1 for both flanks was considered a positive.

A TOP PWM hit was considered conserved if it passed one of the three conservation tests above and had at least one site with an FDR-corrected phyloP score > 1 (i.e., corresponding to an FDR-corrected p-value < 0.1).

### Repeat enrichment

To calculate enrichment for each TF and repeat pair, we identified the intersections of the peak summits from ChIP-seq peaks (or TOPs) and the middle position of GHT-SELEX TOPs with the 2022-10-18 version of the UCSC Genome Browser RepeatMasker track^89^. The enrichment significance between GHT-SELEX and ChIP-seq TOPs and each repeat family was calculated using a one-sided Fisher’s Exact Test (alternative hypothesis being greater overlap), implemented in SciPy^90^. The contingency table took the form of:

|  | TF y summit + | TF y summit - |
| --- | --- | --- |
| Repeat x + | # of repeat x and TF y summit intersections | # of repeat x bases with no TF y summit intersections |
| Repeat x - | # of TF y summits with no repeat x intersections | # of repeat-annotated bases, excluding # of repeat x bases and TF y summits |

## DATA AVAILABILITY

The sequencing raw data for the experiments have been deposited into the SRA database under identifiers PRJEB78913 (ChIP-seq), PRJEB76622 (GHT-SELEX), and PRJEB61115 (HT-SELEX). Genomic interval information generated for the ChIP-seq and GHT-SELEX has been deposited into GEO under accessions GSE280248 and GSE278858, respectively. Information on constructs, experiments, analyses, processed data, comparison tracks, with many accessory files and browsable results, is available at https://codebook.ccbr.utoronto.ca. A larger collection of motifs generated for these experiments in an accompanying study^40^ can be browsed at https://mex.autosome.org and downloaded at https://doi.org/10.5281/ZENODO.8327372. Source Data are provided with this paper.

## CODE AVAILABILITY

The custom scripts for extracting phyloP scores for TOPs and classification as CTOPs by performing the three statistical tests are available at GitHub: https://github.com/imyellan/Codebook_CTOP_scripts.

## ACKNOWLEDGEMENTS

We thank the IT Group of the Institute of Computer Science at Halle University for computational resources, Maximilian Biermann for valuable technical support, and Hadi Tabarraei for aiding with ATAC-seq experiments. This work was supported by the following:

- Canadian Institutes of Health Research (CIHR) grants FDN-148403, PJT-186136, PJT-191768, and PJT-191802, and NIH grant R21HG012258 to T.R.H.
- CIHR grant PJT-191802 to T.R.H. and H.S.N.
- Natural Sciences and Engineering Research Council of Canada (NSERC) grant RGPIN-2018-05962 to H.S.N.
- MSHERF grant 075-15-2025-014 (prev. 075-15-2024-666)
- Assignment 125091010189-3 to I.V.K.
- Russian Science Foundation grant 24-14-20031 to F.A.K.
- Swiss National Science Foundation grant (no. 310030_197082) to B.D.
- Marie Skłodowska-Curie (no. 895426) and EMBO long-term (1139-2019) fellowships to J.F.K.
- NIH grants R01HG013328 and U24HG013078 to M.T.W., T.R.H., and Q.M.
- NIH grants R01AR073228, P30AR070549, and R01AI173314 to M.T.W.
- NIH grant P30CA008748 partially supported Q.M.
- Canada Research Chairs funded by CIHR to T.R.H. and H.S.N.
- Ontario Graduate Scholarships to K.U.L and I.Y.
- A.J. was supported by Vetenskapsrådet (Swedish Research Council) Postdoctoral Fellowship (2016-00158)
- The Billes Chair of Medical Research at the University of Toronto to T.R.H.
- EPFL Center for Imaging
- Institutional funding from EPFL
- Resource allocations from the Digital Research Alliance of Canada

## AUTHOR CONTRIBUTIONS

T.R.H. conceived of the study. R.R. designed the study. R.R., A.B., H.Z., M.B., and C.H. prepared cell lines and performed the ChIP-seq experiments. A.J. and A.W.H.Y. performed and analyzed the SELEX experiments. A.F., I.Y., A.B., A.H.C. and K.U.L. performed statistical and sequence analyses. I.E.V. and Z.M.P. contributed to motif discovery and assessment, and initial evaluation of experiments. K.U.L, I.Y., A.F., and R.R. prepared the illustrations. M.A. assisted with data deposition. I.V.K., P.B., Q.M., H.S.N., and H.S.N. organized the study and oversaw the data collection and data processing. T.R.H., R.R., A.F., and I.Y. wrote the paper. All authors contributed to data analysis and reviewed the manuscript.

## SUPPLEMENTAL TABLES AND DOCUMENTS

**Supplementary Table 1. Overview of the tested TFs.** This table lists the TFs that were tested in this study using ChIP-seq.

**Supplementary Table 2. List of all the ChIP-seq experiments**. This table lists the ChIP experiments, their approval status, and related produced files.

**Supplementary Table 3. Summary of 217 TFs with successful ChIP-seq data.** This table lists the TFs with successful ChIP-seq experiments either “Motif-enriched” or significant overlap between replicates, together with the list of ChIP-seq samples used in “merged” peaks for each TF and their binding category (i.e. Promoter TF, Dark TF, Enhancer TF, or Others). For the TFs with available GHT-SELEX data (hence TOP sites, the number of optimized ChIP-seq peaks (i.e. Triple peaks), number of TOP ChIP-seq peaks, fraction of direct binding sites (i.e. #TOP peaks divided by #Triple peaks), number of TOPs, number of CTOPs, fraction of conserved TOPs (i.e. #CTOPs / #TOPs), and the median age of the TOP sites are also included. Note that the number of TOP ChIP-seq peaks might be different (less) than TOPs (referring to triple-overlap PWM hits), since each peak might comprise multiple PWM hits.

**Supplementary Table 4. Consolidated functional information for Dark TFs.** This table provides the data underlying **Fig. 7** including the references in the literature.

**Supplementary Document 1. Enrichment of transposable elements in TOPs, with expanded TE family classification.** Heatmap is from **Fig. 3b**, with expanded labels for specific elements enriched in TOPs of each TF

**Supplementary Document 2. Heatmaps of conservation/phyloP score across TOPs for 137 TFs.** This document provides the same analysis as **Fig. 5a**, for all TFs of the study, heatmaps of phyloP scores in PWM hits (middle column) and flanking sequences of tops are displayed. Bars to the right indicate which tests of conservation are satisfied (Likelihood-ratio, Correlation, Wilcoxon), along with overlaps with promoters (P) and specific repeat families if applicable.

**Supplementary Document 3. Overview of Dark TFs’ function.** This document provides a summary of functional impact of Dark TFs. It is based on a survey (**Fig. 7**) of Dark TF interaction with chromatin remodelers, other zinc-finger proteins, the specific categories and age of enriched TEs within each TFs TOPs, their motifs, as well as documented functions based on literature review.

**Supplementary Document 4. Analysis of binding site ages for TFs.** This document provides a summary of evolutionary analysis performed to estimate the age of each TOP site for all the TFs in this study, as well as the age of the TFs.

---

**Supplementary Fig. 1.**
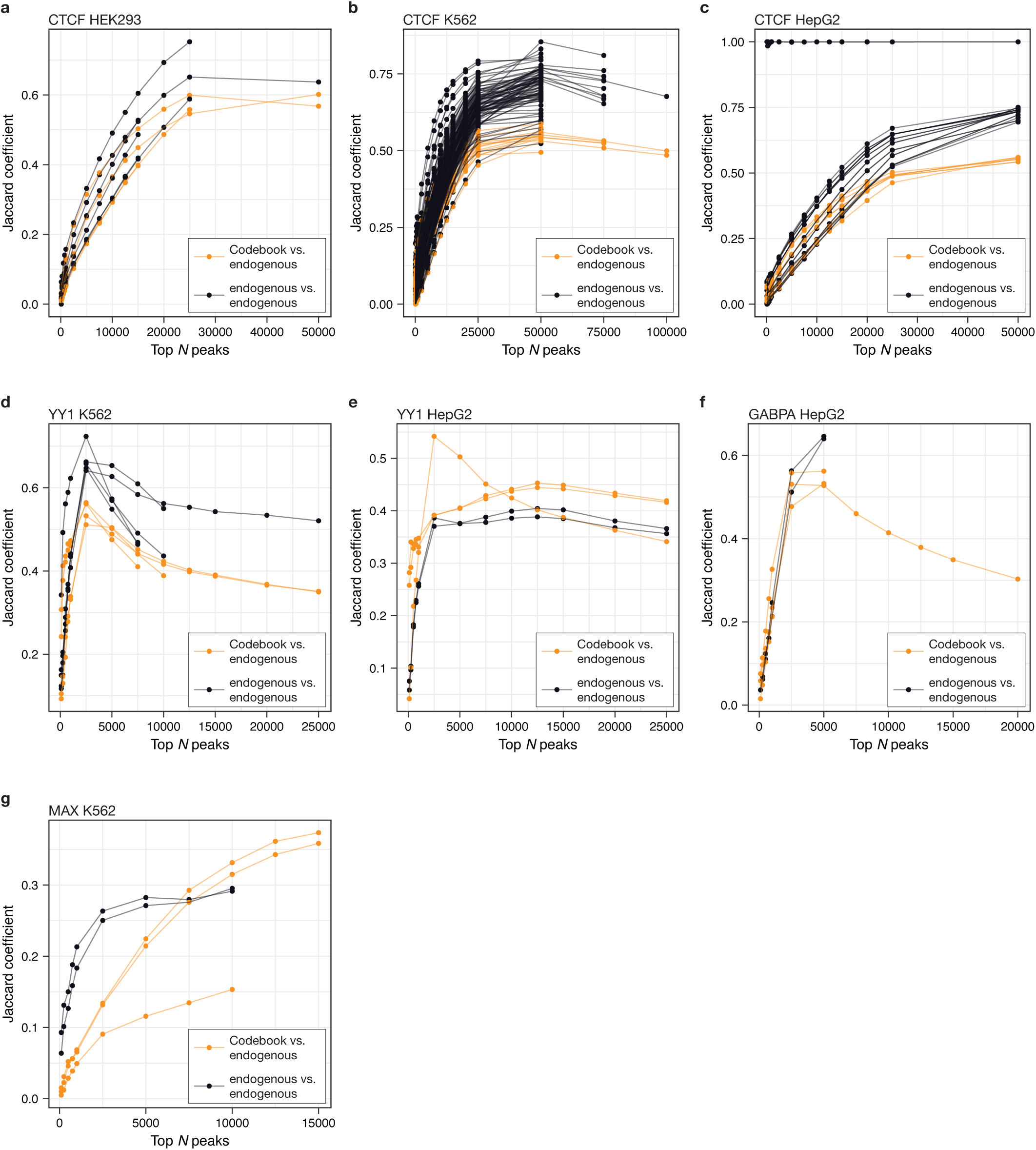
Comparison of ChIP-seq peaks obtained from ectopically expressed TF in Codebook project with ChIP-seq peaks from previous studies using endogenous TFs. Individual plots show the overlap (Jaccard coefficient) between peak sets for pairs of ChIP experiments. ChIP peaks are ordered by their scores and the Jaccard coefficient is calculated using the top *N* peaks from each experiment, with different values of *N*. Comparisons between pairs of ChIP experiments that IP the endogenous TF (endogenous vs. endogenous) are distinguished from comparisons between a Codebook ChIP experiment and a ChIP experiment that IPs the endogenous TF (Codebook vs endogenous). All Codebook ChIP experiments are performed in HEK293 cells, each panel shows comparisons for a specific TF with endogenous IP experiments performed in a specific cell line. (**a**), CTCF, HEK293 cells. (**b**) CTCF, K562 cells. (**c)** CTCF, HepG2 cells. (**d**) YY1, K562 cells. (**e**) YY1, HepG2 cells. (**f**) GABPA, HepG2 cells. (**g**) MAX, K572 cells.

**Supplementary Fig. 2.**
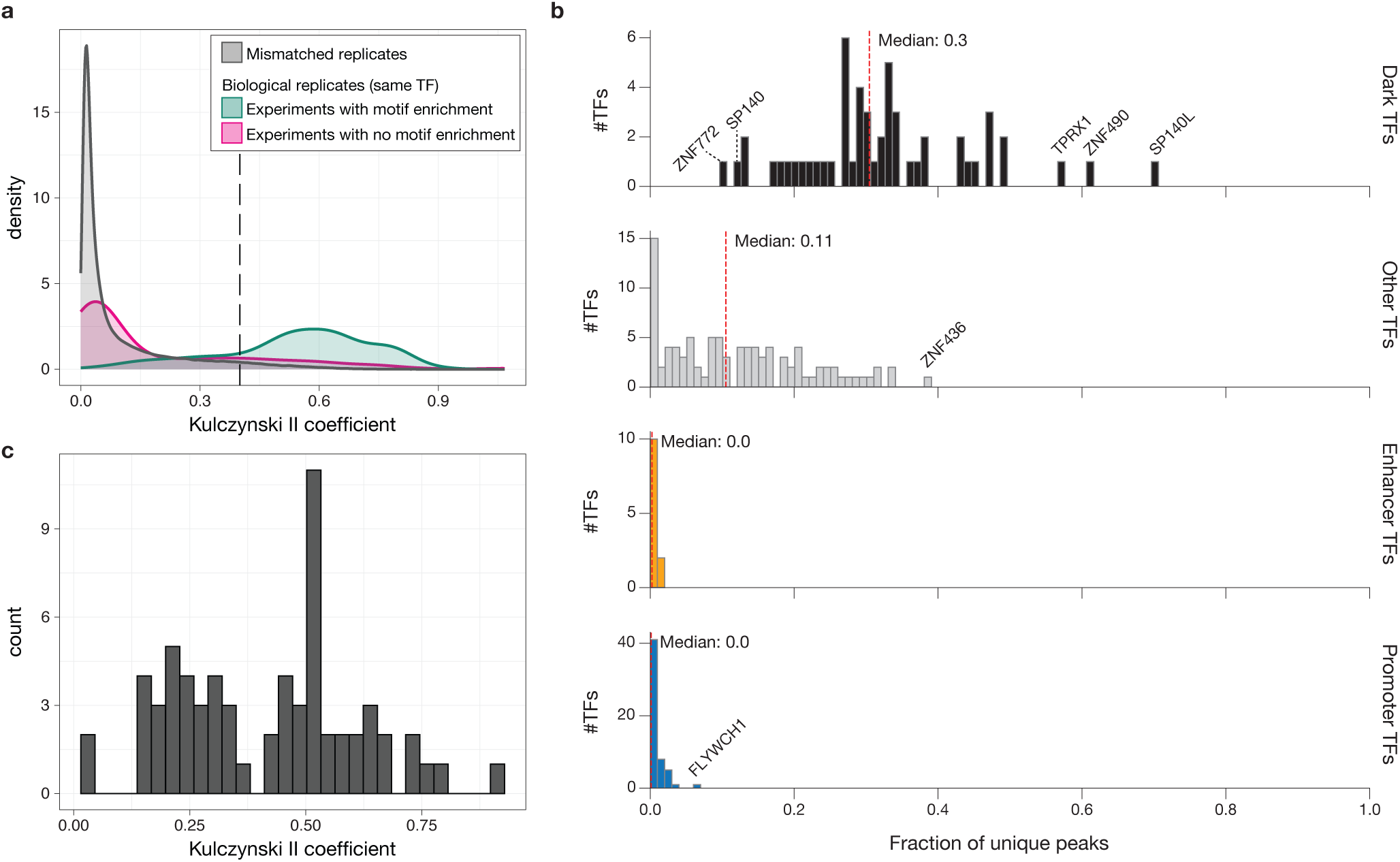
Peak overlap assessment of ChIP-seq as a measure of success. **(a)** Distribution of peak overlap between ChIP-seq replicates, for the experiments yielded a motif (green), experiments that did not yield a motif (red), and the mismatched (i.e., TF identities permuted) replicates (grey). Values are calculated Kulczynski II similarity metric (i.e. average of overlaps). Dotted line indicates the threshold at which the peak overlap of experimental replicates without motif enrichment considered successful and were included in downstream analyses. **(b)** Distribution peak uniqueness for defined TF categories (Promoter, Enhancer, Dark, and Other), measured as the fraction of ChIP-seq peaks (at the universal threshold) not overlapping with *any* peak from any other TF in this study. **(c)** Distribution of Kulczynski II similarity metric between successful ChIP-seq replicates (as in (a)), restricted to the TFs that have a low peak overlap with other TFs (specifically, the 87 TFs in the upper left darker region of the square matrix in **Fig. 2b**).

**Supplementary Fig. 3.**
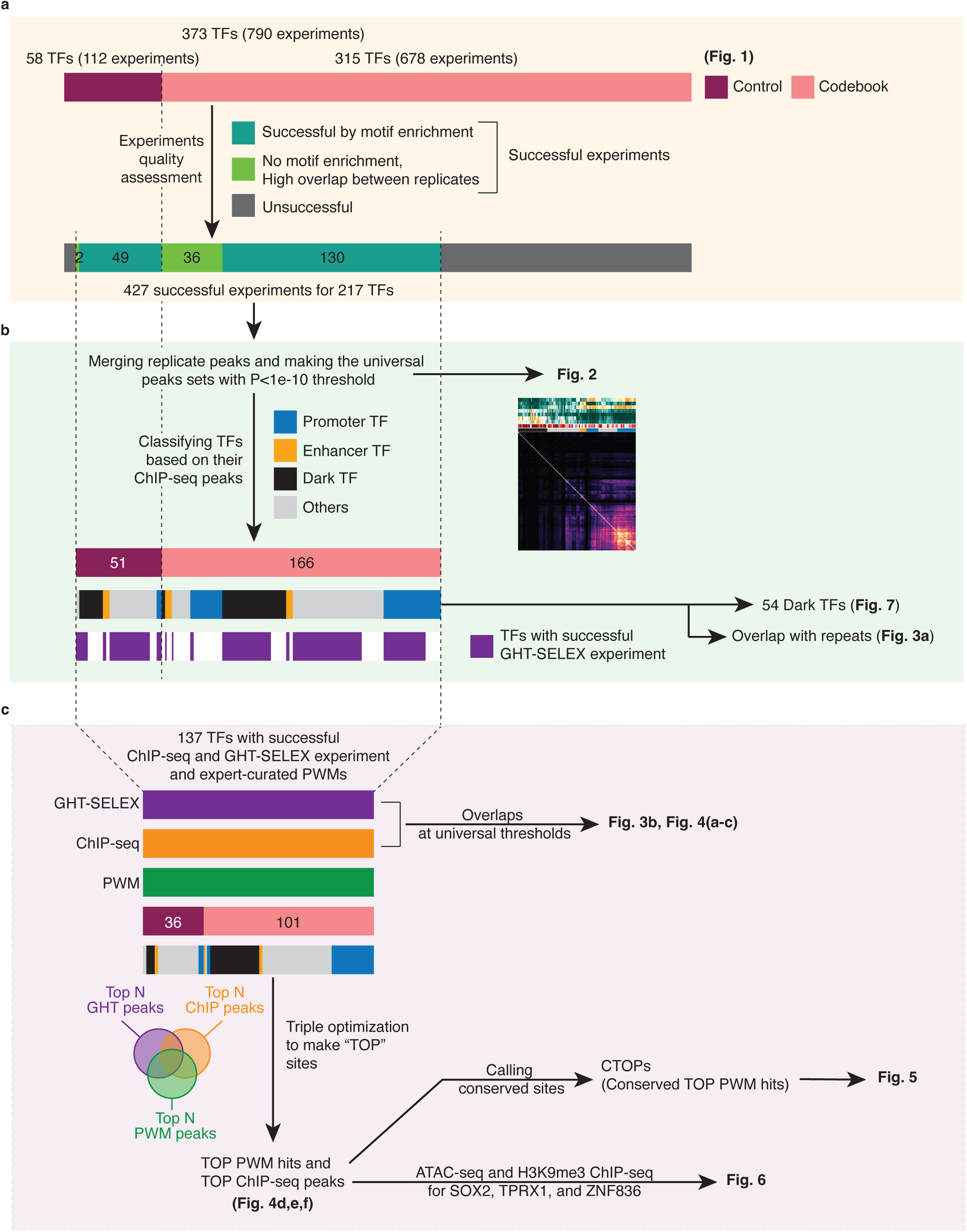
Overview of ChP-seq data processing. **(a)** selection process for successful ChIP-experiments based on replicate overlap and motif enrichment. Data for 790 ChIP experiments (including replicates for 373 TFs) were assessed and 427 (217 TFs) were considered successful. (**b**) peaks for the successful replicates were merged to create one set of peaks for each TF. The overlap of peak sets with each other and various genomic tracks were evaluated and four binding categories of TFs emerged. 137 of these TFs also had successful GHT-SELEX experiments as labelled in purple. (**c**) defining triple optimized overlap peaks (TOPs) and conserved TOPs (CTOPs). TOPs were defined based on overlap of ChIP-seq and GHT-SELEX peaks as well as PWM predicted binding sites at optimized thresholds (see **Methods**). CTOPs were defied based on phyloP conservation pattern of the PWM hit.

**Supplementary Fig. 4.**
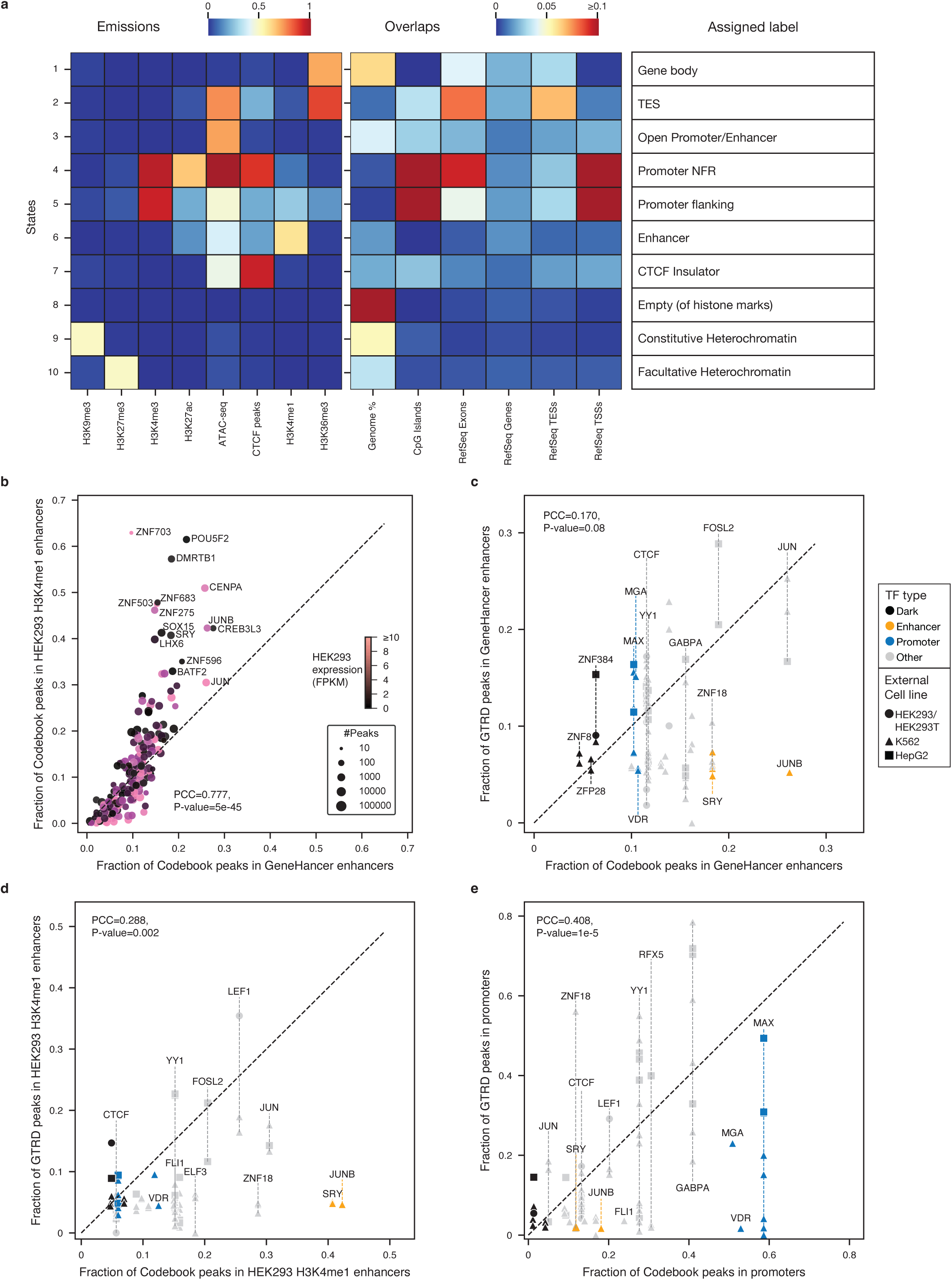
Generating HEK293-specific chromatin states. (**a**) Characterization of output tracks for the ChromHMM with 10 states trained on various HEK293-specific ChIP-seq and ATAC-seq data (i.e., ChIP-seq data for H3K9me3, H3K27me3, H3K4me1, H3K4me3, H3K36me3, and H3K27ac from ENCODE project, and ATAC-seq and CTCF peaks generated here by Codebook project). Based on the correspondence between emissions and the chromatin marks and genome annotations, states were assigned to: Gene body, Transcription end site (TES), Open Promoter/Enhancer, Promoter NFR (nucleosome-free regions), Promoter flanking region, Enhancer, CTCF Insulator, Empty (of assigned histone marks), Constitutive Heterochromatin, and Facultative Heterochromatin. (**b**) the panel shows the fraction of TF ChIP-seq peaks overlap with GeneHancer annotated enhancers (x-axis) and HEK293 enhancers (defined by H3K4me1-positive regions from ChromHMM; y-axis). Circle sizes (TFs) are scaled based on peak numbers and color gradient corresponds to endogenous expression level of TFs in HEK293 cells^33^. (**c, d, e**) Comparison of Codebook ChIP-seq peaks and publicly available ChIP-seq or ChIP-exo data on GTRD, for 21 control TFs, based on the fraction of peaks overlapping with GeneHancer enhancers (**c**), HEK293 defined enhancers (**d**), and protein-coding promoters (**e**). For the Codebook data (x-axis), the merged peak set with the universal threshold has been used.

**Supplementary Fig. 5.**
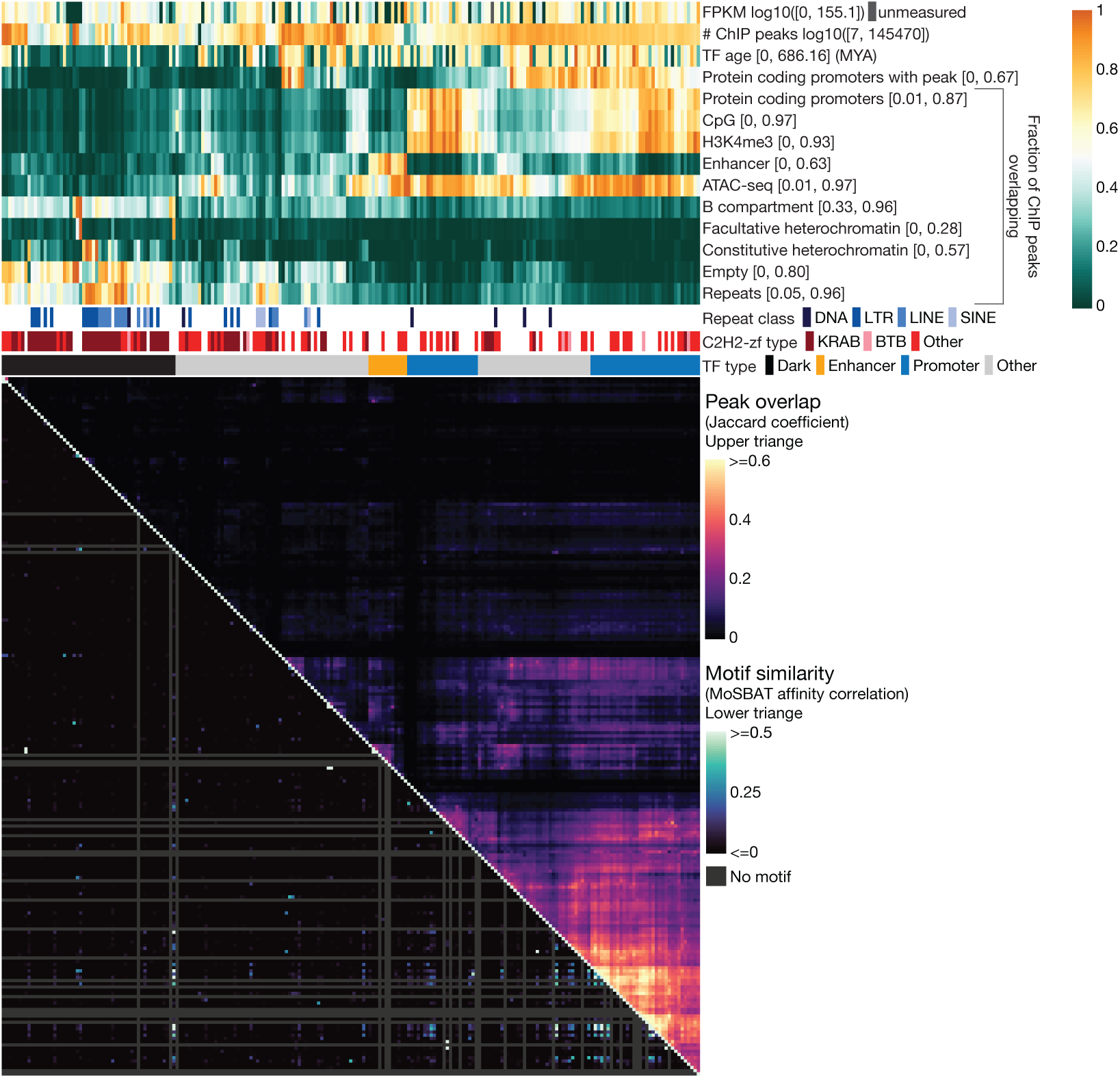
Expanded information for genomic tracks and protein expression data for 217 TF binding sites. This is a more detailed version of **Fig. 2**. Additional tracks, such as gene expression in HEK293 cells (FPKM)^33^, number of total ChIP-seq peaks (at the universal threshold of MACS2 P-value≤10^-10^), TF age, fraction of human protein-coding promoters (out of 20,052) covered by TF peaks, fraction of ChIP-seq peaks falling within: CpG islands, H3K4me3-positive regions, facultative heterochromatin, and constitutive heterochromatin, with the main repeat class bound by the TFs are included. The upper triangle in the bottom square is the same as **Fig. 2**, however, the lower triangle here shows the similarity between PWMs for each pair of TFs, calculated by MoSBAT^93^. Gray stripes correspond to the TFs without a selected PWM in the Codebook set.

**Supplementary Fig. 6.**
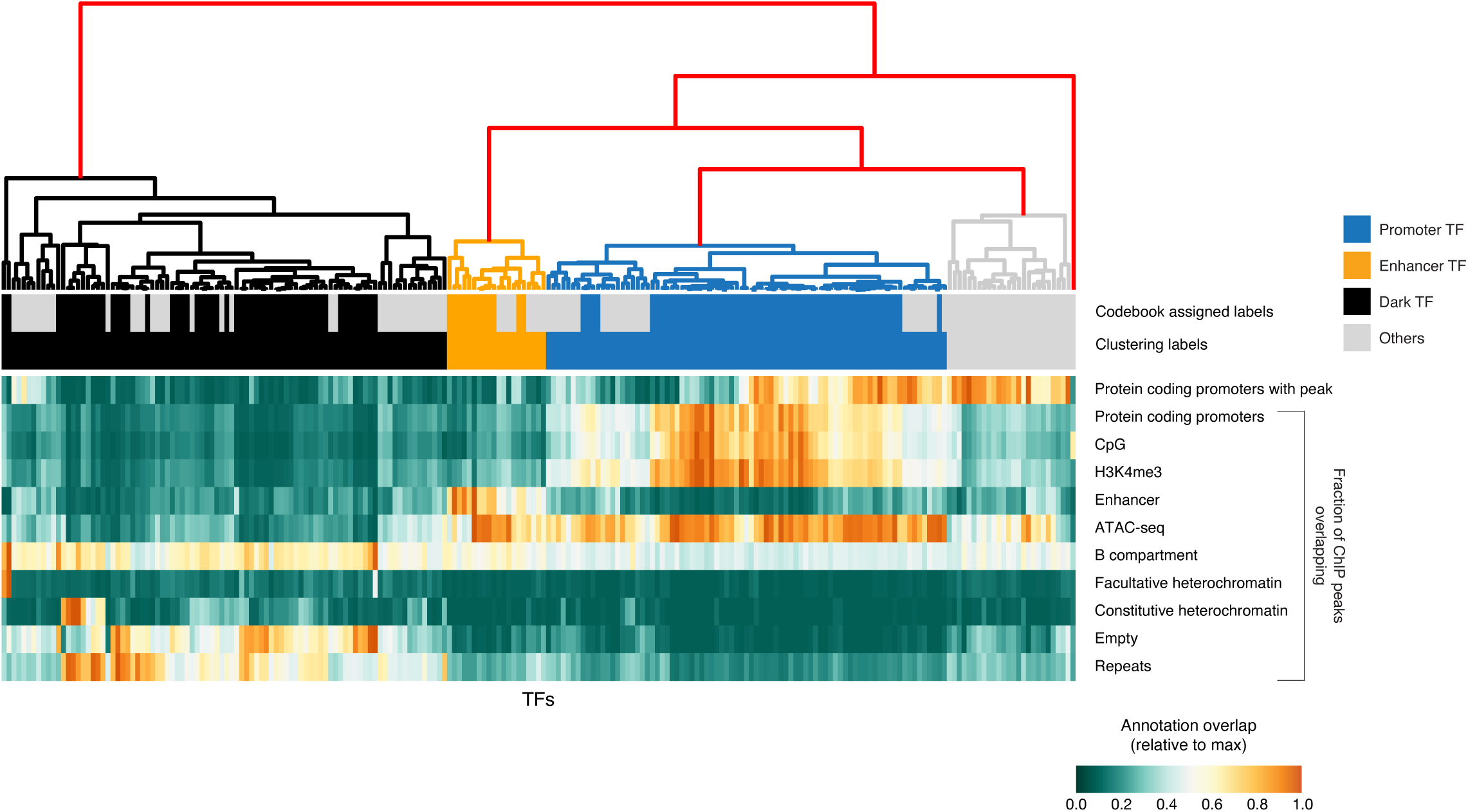
Automated clustering of the TFs and comparison with expert labeling. The 217 TFs with successful ChIP-seq data were clustered using the features presented at the top of **Fig. 2b**, applying hierarchical clustering with a correlation similarity metric and average linkage. The top five major branches (red) were then assigned to Promoter TFs, Dark TFs, and Enhancer TFs, with two branches (one containing only a single member) left as Others (see “Clustering labels”). The “Codebook assigned labels” are shown on top for comparison.

**Supplementary Fig. 7.**
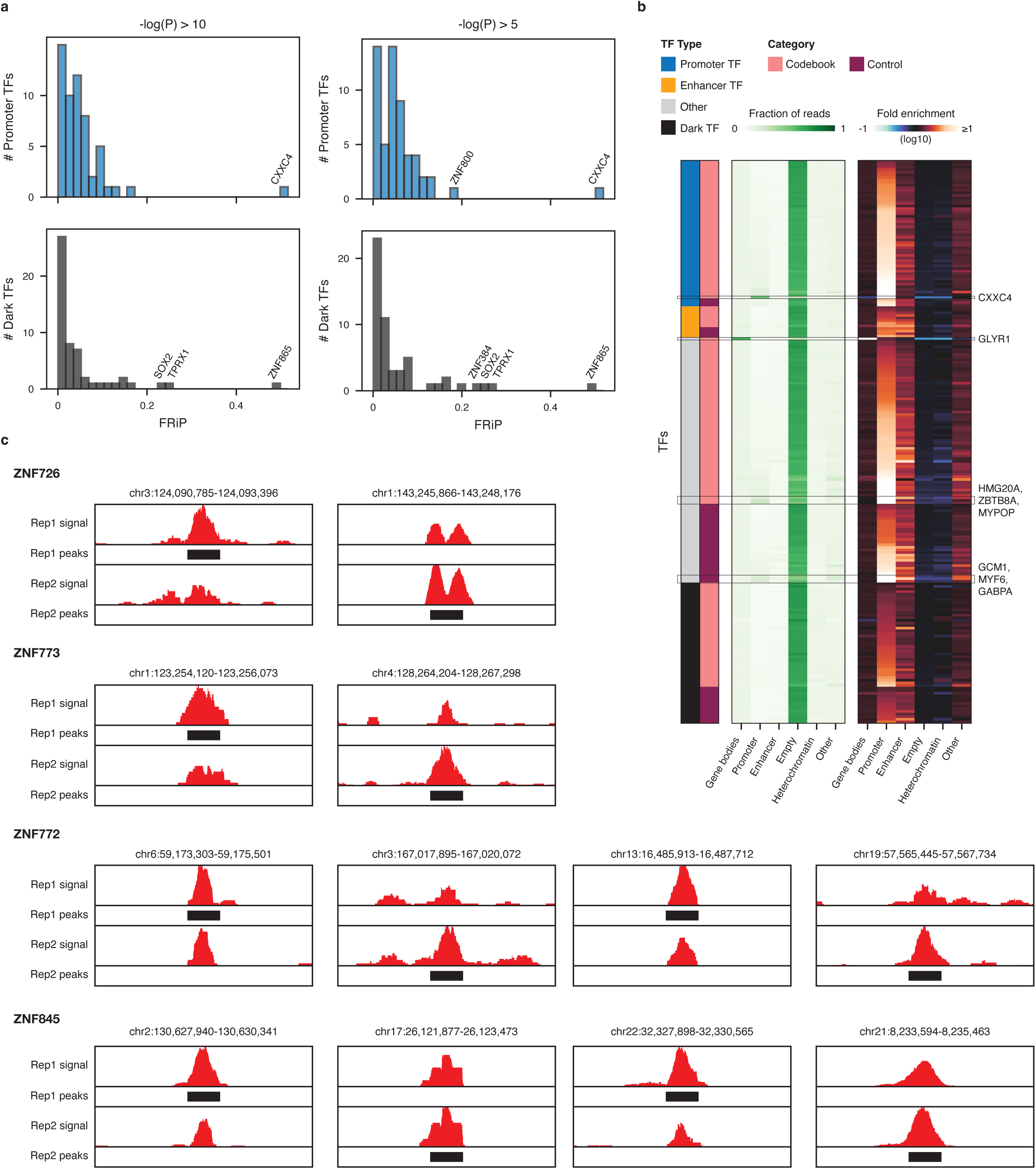
Comparative quality control for ChIP-seq peaks. (**a**) Fraction of Reads in Peaks (FRiP) for Promoter TFs and Dark TFs, at the universal threshold (MACS2 *P*-value < 1e−10) and a less stringent threshold (1e−5). (**b**) Middle heatmap: distribution of raw reads (i.e., before peak calling) in representative chromatin states (from ChromHMM) for 217 TFs with successful data. Right heatmap: log₁₀ fold enrichment of the read counts (in the middle heatmap) relative to the size of each chromatin state. Labels and categories of TFs are indicated on the left. (**c**) Genome browser examples for four Dark TF, showing read pileups (red) for two replicates, at locations where both replicates show a high read coverage, but only one replicate passed the peak calling threshold (black square).

**Supplementary Fig. 8.**
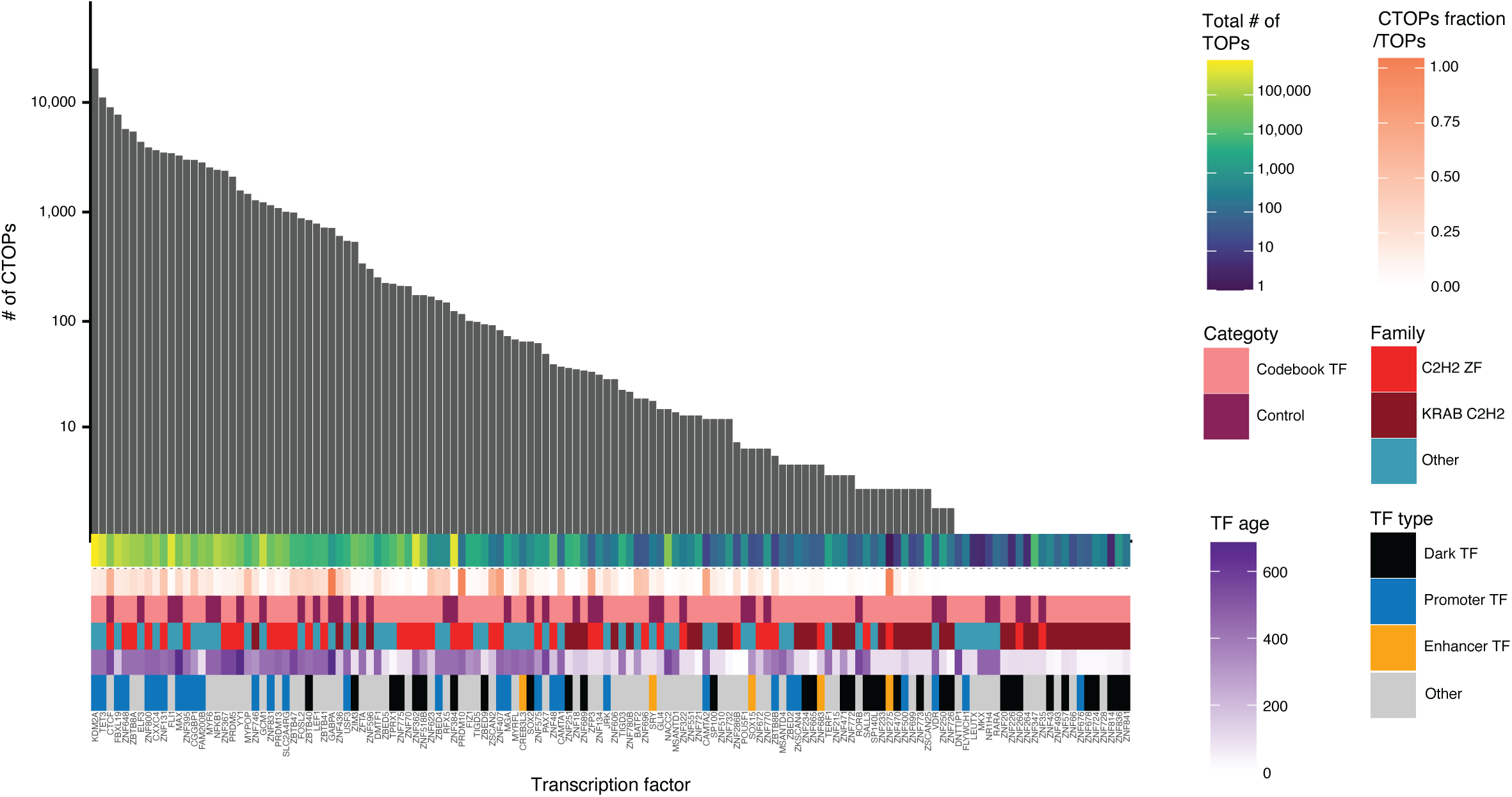
Distribution of CTOP counts for 137 Codebook TFs. Bar graph displays the number of individual CTOP sites obtained for each TF. Other properties for each TF (TOP counts, CTOPs fraction/TOPs, category, DBD family, TF type, and TF age) are annotated below the bar graph. Heatmap scales are shown on the top right.

**Supplementary Fig. 9.**
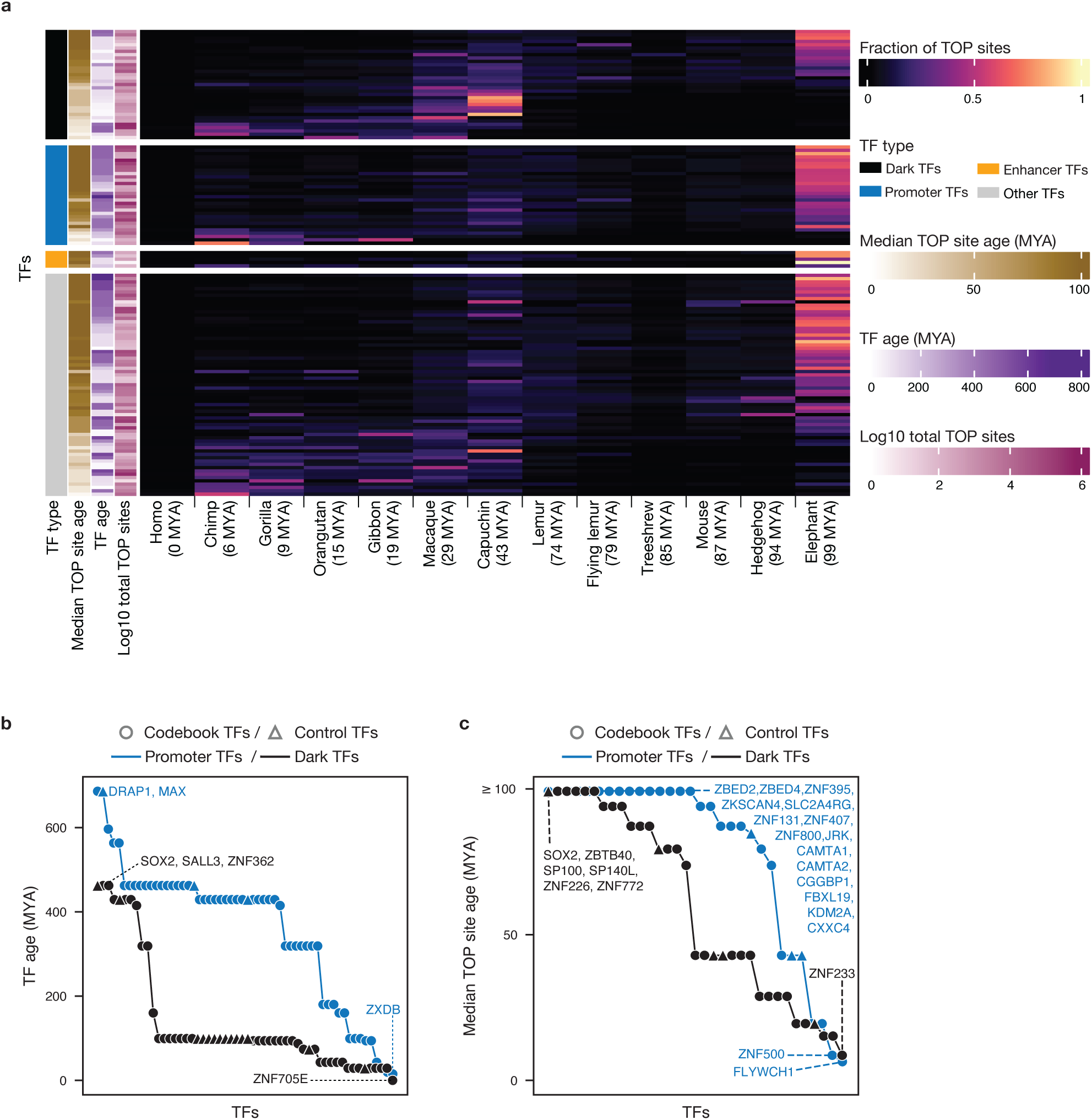
Age distribution relationship between TFs and their binding sites. (**a**) Heatmap showing the fraction of TOP sites for each TF, with an exact match within ancestral genomes available in Zoonomia (13 clades in human lineage up to 99 million years ago; MYA) ordered by age. In the left side for each TOP, the related TF types, median age of TOP sites, TF ages (MYA), and log 10 of total TOP sites are indicated. (**b**) The age comparison of Dark TFs and Promoter TFs. (**c**) Comparing the median age of the TOP sites between Promoter and Dark TFs in both control and Codebook categories.

**Supplementary Fig. 10.**
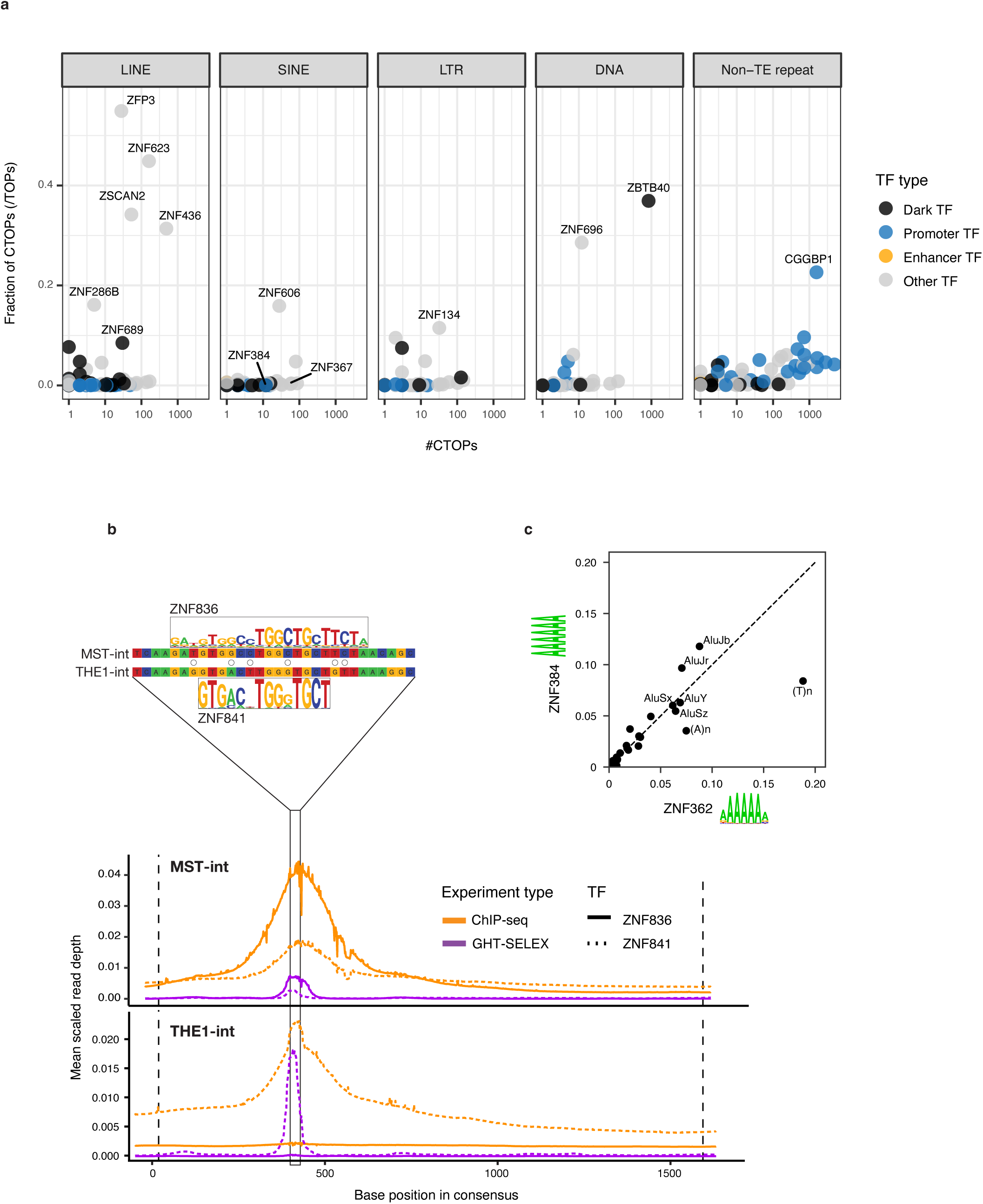
Codebook TFs exhibit diverse co-evolutionary interactions with repeats. **(a)** Plots showing the fraction of each TF’s TOPs that are conserved (i.e. CTOPs) and overlap a major class of transposable elements or non-TE repeats. The fraction of TOPs that are conserved and overlap a repeat class is shown on the y-axis, and the log10 count of these sites is shown on the x-axis. Each TF is colored according to its classification as a Dark TF, Promoter TF, Enhancer TF, and Other TFs. Only proteins with a fraction greater than 0.1 of conserved TOPs that fall in a repeat class are labeled. TFs discussed in the main text are also labeled. **(b)** Top plot shows binding motifs of two paralogous Dark TFs, ZNF836 and ZNF841, aligned to the consensus sequences of their specific TE targets, MSTA-int and THE1-int, respectively. They bind to a homologous region in the two different but related LTR families. Circles indicate nucleotide differences between the two TEs. The bottom plot shows the average ChIP-seq and GHT-SELEX signal across all the instances of MST-int and THE1-int. **(c)** compares the fractions of TOP sites for two Dark TFs, ZNF362 and ZNF384, that overlap with various repeat elements. Both bind poly-A sequences; motifs are shown adjacent to the axis labels. Alu elements are the dominant target TEs for both TFs.

**Supplementary Fig. 11.**
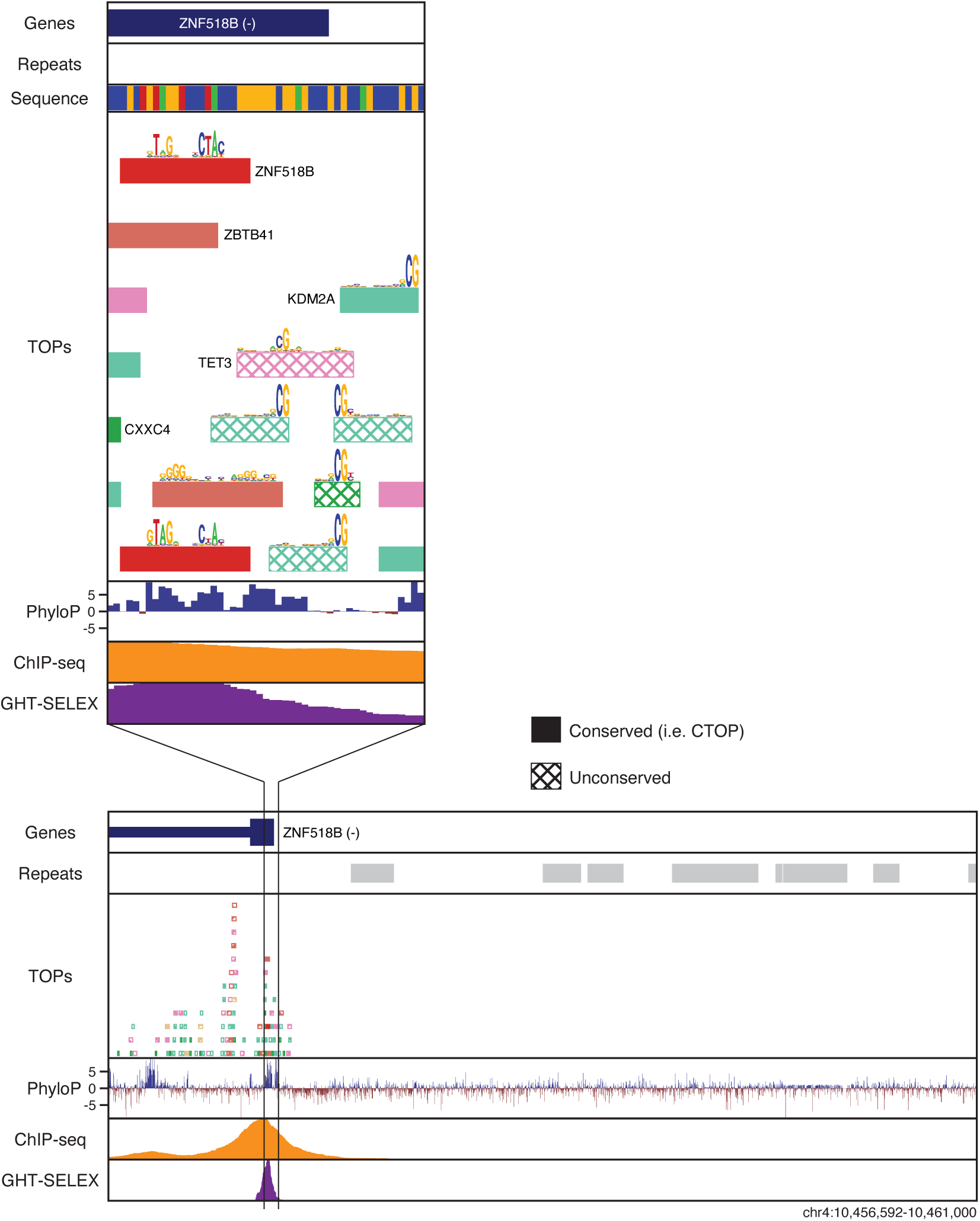
Age estimation of TOPs. Three different tests and two different thresholds were applied. Heatmaps show the proportion of each TF’s TOPs (rows) inferred to be a certain age, as in **Fig**. **5**, but with each panel utilizing a different scheme. *Top row*: Age of each TOP site inferred as that of oldest ancestral genome with a gapless alignment to the human TOP site and minimum 75% identity (left) or 100% identity (right). (**Fig. 5** shows this same analysis with a 0% identity threshold). *Middle row*: Age of each TOP site inferred as that of oldest *species* with a gapless alignment to the human TOP site and minimum 0% identity (left) or 100% identity (right). *Bottom row*: Age of each TOP site inferred as that of the oldest *clade* where 60% of the species have a gapless alignment to the human TOP with a minimum 0% identity (left) or 100% identity (right).

**Supplementary Fig. 12.**
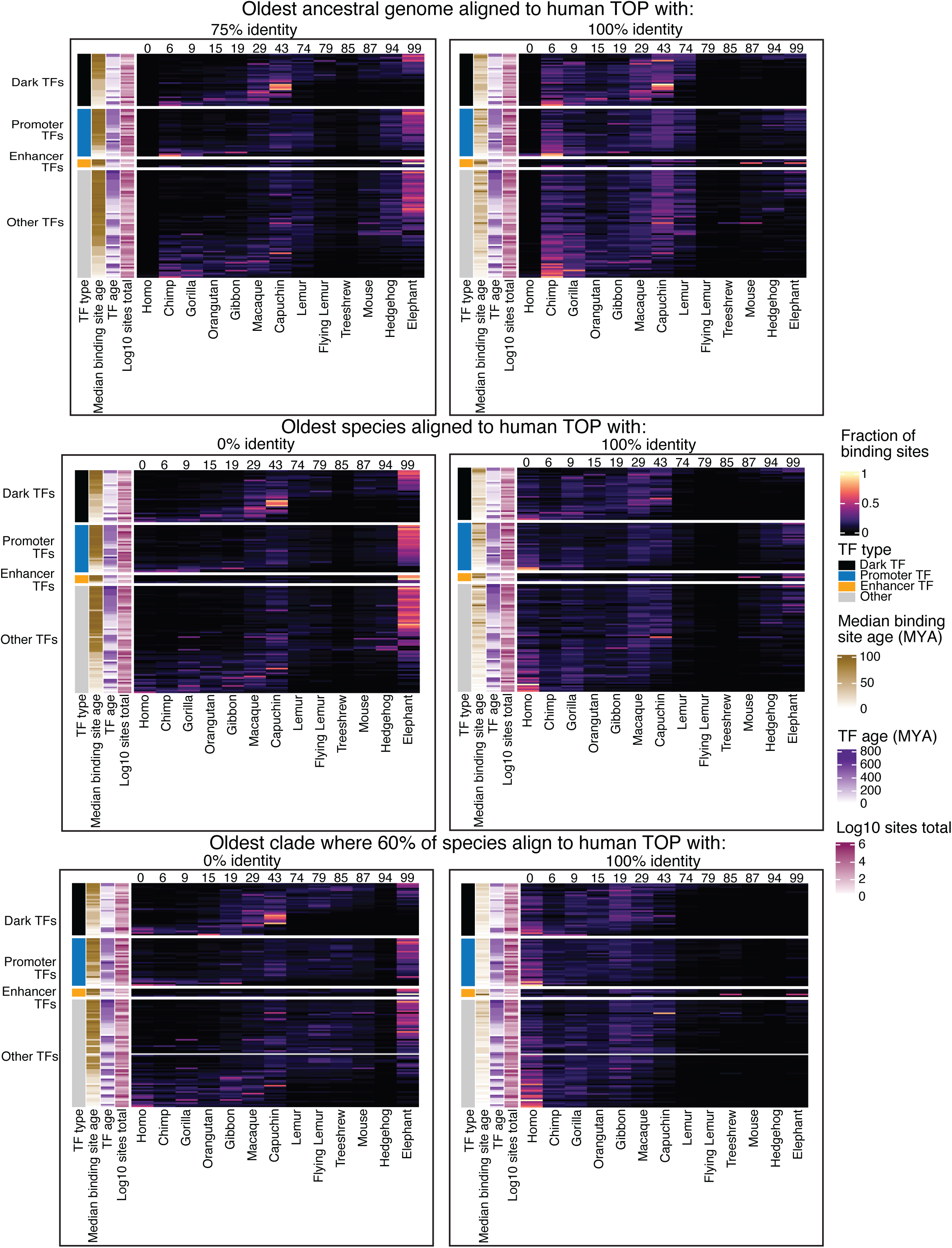
Dark TF, ZNF518B, is a potential self-regulator. Two palindromic conserved binding sites of ZNF518B (red) are in the promoter of ZNF518B itself. Binding sites of four other TFs (i.e. TOPs and CTOPs for ZBTB41, KDM2A, TET3, and CXXC4) are also present in this region.

## REFERENCES

1. Gemmell, N.J. Repetitive DNA: genomic dark matter matters. Nat Rev Genet 22, 342 (2021).

2. Alexander, R.P., Fang, G., Rozowsky, J., Snyder, M. & Gerstein, M.B. Annotating non-coding regions of the genome. Nat Rev Genet 11, 559–71 (2010).

3. Britten, R.J. & Davidson, E.H. Gene regulation for higher cells: a theory. Science 165, 349–57 (1969).

4. Nobrega, M.A., Ovcharenko, I., Afzal, V. & Rubin, E.M. Scanning human gene deserts for long-range enhancers. Science 302, 413 (2003).

5. Venter, J.C. et al. The sequence of the human genome. Science 291, 1304–51 (2001).

6. Nobrega, M.A., Zhu, Y., Plajzer-Frick, I., Afzal, V. & Rubin, E.M. Megabase deletions of gene deserts result in viable mice. Nature 431, 988–93 (2004).

7. Imbeault, M., Helleboid, P.Y. & Trono, D. KRAB zinc-finger proteins contribute to the evolution of gene regulatory networks. Nature 543, 550–554 (2017).

8. Cosby, R.L. et al. Recurrent evolution of vertebrate transcription factors by transposase capture. Science 371(2021).

9. Schmidt, D. et al. Waves of retrotransposon expansion remodel genome organization and CTCF binding in multiple mammalian lineages. Cell 148, 335–48 (2012).

10. Chuong, E.B., Elde, N.C. & Feschotte, C. Regulatory evolution of innate immunity through co-option of endogenous retroviruses. Science 351, 1083–7 (2016).

11. Cohen, C.J., Lock, W.M. & Mager, D.L. Endogenous retroviral LTRs as promoters for human genes: a critical assessment. Gene 448, 105–14 (2009).

12. Ernst, J. et al. Genome-scale high-resolution mapping of activating and repressive nucleotides in regulatory regions. Nat Biotechnol 34, 1180–1190 (2016).

13. Kunarso, G. et al. Transposable elements have rewired the core regulatory network of human embryonic stem cells. Nat Genet 42, 631–4 (2010).

14. Villar, D. et al. Enhancer evolution across 20 mammalian species. Cell 160, 554–66 (2015).

15. Jacobs, F.M. et al. An evolutionary arms race between KRAB zinc-finger genes ZNF91/93 and SVA/L1 retrotransposons. Nature 516, 242–5 (2014).

16. Najafabadi, H.S. et al. C2H2 zinc finger proteins greatly expand the human regulatory lexicon. Nat Biotechnol (2015).

17. Bannister, A.J. et al. Selective recognition of methylated lysine 9 on histone H3 by the HP1 chromo domain. Nature 410, 120–4 (2001).

18. Lachner, M., O’Carroll, D., Rea, S., Mechtler, K. & Jenuwein, T. Methylation of histone H3 lysine 9 creates a binding site for HP1 proteins. Nature 410, 116–20 (2001).

19. Ayyanathan, K. et al. Regulated recruitment of HP1 to a euchromatic gene induces mitotically heritable, epigenetic gene silencing: a mammalian cell culture model of gene variegation. Genes Dev 17, 1855–69 (2003).

20. Emerson, R.O. & Thomas, J.H. Adaptive evolution in zinc finger transcription factors. PLoS Genet 5, e1000325 (2009).

21. Senft, A.D. & Macfarlan, T.S. Transposable elements shape the evolution of mammalian development. Nat Rev Genet 22, 691–711 (2021).

22. Milovanović, D. et al. Tissue-specific restriction of TE-derived regulatory elements safeguards cell-type identity. bioRxiv, 2025.05.13.653700 (2025).

23. Lambert, S.A. et al. The Human Transcription Factors. Cell 175, 598–599 (2018).

24. Partridge, E.C. et al. Occupancy maps of 208 chromatin-associated proteins in one human cell type. Nature 583, 720–728 (2020).

25. Long, H.K., Prescott, S.L. & Wysocka, J. Ever-Changing Landscapes: Transcriptional Enhancers in Development and Evolution. Cell 167, 1170–1187 (2016).

26. Zhu, F. et al. The interaction landscape between transcription factors and the nucleosome. Nature 562, 76–81 (2018).

27. Zaret, K.S. & Carroll, J.S. Pioneer transcription factors: establishing competence for gene expression. Genes Dev 25, 2227–41 (2011).

28. Courey, A.J. Cooperativity in transcriptional control. Curr Biol 11, R250–2 (2001).

29. Stubbs, L., Sun, Y. & Caetano-Anolles, D. Function and Evolution of C2H2 Zinc Finger Arrays. Subcell Biochem 52, 75–94 (2011).

30. Najafabadi, H.S., Albu, M. & Hughes, T.R. Identification of C2H2-ZF binding preferences from ChIP-seq data using RCADE. Bioinformatics 31, 2879–81 (2015).

31. Worsley Hunt, R. & Wasserman, W.W. Non-targeted transcription factors motifs are a systemic component of ChIP-seq datasets. Genome Biol 15, 412 (2014).

32. Auerbach, R.K. et al. Mapping accessible chromatin regions using Sono-Seq. Proc Natl Acad Sci U S A 106, 14926–31 (2009).

33. Schmitges, F.W. et al. Multiparameter functional diversity of human C2H2 zinc finger proteins. Genome Res 26, 1742–1752 (2016).

34. Jolma, A. et al. Perspectives on Codebook: sequence specificity of uncharacterized human transcription factors. bioRxiv, 2024.11.11.622097 (2024).

35. Jolma, A. et al. GHT-SELEX demonstrates unexpectedly high intrinsic sequence specificity and complex DNA binding of many human transcription factors. bioRxiv, 2024.11.11.618478 (2024).

36. Vorontsov, I.E. et al. Cross-platform DNA motif discovery and benchmarking to explore binding specificities of poorly studied human transcription factors. bioRxiv, 2024.11.11.619379 (2024).

37. Lambert, S.A. et al. The Human Transcription Factors. Cell 172, 650–665 (2018).

38. Isakova, A. et al. SMiLE-seq identifies binding motifs of single and dimeric transcription factors. Nat Methods 14, 316–322 (2017).

39. Kean, M.J. et al. Structure-function analysis of core STRIPAK Proteins: a signaling complex implicated in Golgi polarization. J Biol Chem 286, 25065–75 (2011).

40. Vorontsov, I.E. et al. Cross-platform DNA motif discovery and benchmarking to explore binding specificities of poorly studied human transcription factors. bioRxiv, 2024.11.11.619379 (2024).

41. Berger, M.F. et al. Compact, universal DNA microarrays to comprehensively determine transcription-factor binding site specificities. Nat Biotechnol 24, 1429–35 (2006).

42. Jolma, A. et al. Multiplexed massively parallel SELEX for characterization of human transcription factor binding specificities. Genome Res 20, 861–73 (2010).

43. Zhang, Y. et al. Model-based analysis of ChIP-Seq (MACS). Genome Biol 9, R137 (2008).

44. O’Leary, N.A. et al. Reference sequence (RefSeq) database at NCBI: current status, taxonomic expansion, and functional annotation. Nucleic Acids Res 44, D733–45 (2016).

45. Ernst, J. & Kellis, M. ChromHMM: automating chromatin-state discovery and characterization. Nat Methods 9, 215–6 (2012).

46. Fishilevich, S. et al. GeneHancer: genome-wide integration of enhancers and target genes in GeneCards. Database (Oxford) 2017(2017).

47. Kolmykov, S. et al. GTRD: an integrated view of transcription regulation. Nucleic Acids Res 49, D104–D111 (2021).

48. Jolma, A. et al. GHT-SELEX demonstrates unexpectedly high intrinsic sequence specificity and complex DNA binding of many human transcription factors. bioRxiv, 2024.11.11.618478 (2024).

49. Badis, G. et al. A library of yeast transcription factor motifs reveals a widespread function for Rsc3 in targeting nucleosome exclusion at promoters. Mol Cell 32, 878–87 (2008).

50. Core, L.J. et al. Analysis of nascent RNA identifies a unified architecture of initiation regions at mammalian promoters and enhancers. Nat Genet 46, 1311–20 (2014).

51. FitzGerald, P.C., Shlyakhtenko, A., Mir, A.A. & Vinson, C. Clustering of DNA sequences in human promoters. Genome Res 14, 1562–74 (2004).

52. Pollard, K.S., Hubisz, M.J., Rosenbloom, K.R. & Siepel, A. Detection of nonneutral substitution rates on mammalian phylogenies. Genome Res 20, 110–21 (2010).

53. Zou, Z. et al. Translatome and transcriptome co-profiling reveals a role of TPRXs in human zygotic genome activation. Science 378, abo7923 (2022).

54. Helleboid, P.Y. et al. The interactome of KRAB zinc finger proteins reveals the evolutionary history of their functional diversification. EMBO J 38, e101220 (2019).

55. Huttlin, E.L. et al. Dual proteome-scale networks reveal cell-specific remodeling of the human interactome. Cell 184, 3022–3040 e28 (2021).

56. Kuroda, T. et al. SALL3 expression balance underlies lineage biases in human induced pluripotent stem cell differentiation. Nat Commun 10, 2175 (2019).

57. Singh, J.K. et al. Zinc finger protein ZNF384 is an adaptor of Ku to DNA during classical non-homologous end-joining. Nat Commun 12, 6560 (2021).

58. Feu, S. et al. OZF is a Claspin-interacting protein essential to maintain the replication fork progression rate under replication stress. FASEB J 34, 6907–6919 (2020).

59. Ge, L.P. et al. ZNF689 deficiency promotes intratumor heterogeneity and immunotherapy resistance in triple-negative breast cancer. Cell Res 34, 58–75 (2024).

60. Zhou, M. et al. ZBTB40 is a telomere-associated protein and protects telomeres in human ALT cells. J Biol Chem 299, 105053 (2023).

61. Cui, Y., Zhou, M., He, Q. & He, Z. Zbtb40 Deficiency Leads to Morphological and Phenotypic Abnormalities of Spermatocytes and Spermatozoa and Causes Male Infertility. Cells 12(2023).

62. Campos-Sanchez, R., Kapusta, A., Feschotte, C., Chiaromonte, F. & Makova, K.D. Genomic landscape of human, bat, and ex vivo DNA transposon integrations. Mol Biol Evol 31, 1816–32 (2014).

63. Lewin, T.D., Royall, A.H. & Holland, P.W.H. Dynamic Molecular Evolution of Mammalian Homeobox Genes: Duplication, Loss, Divergence and Gene Conversion Sculpt PRD Class Repertoires. J Mol Evol 89, 396–414 (2021).

64. Thomas, J.H. & Schneider, S. Coevolution of retroelements and tandem zinc finger genes. Genome Res 21, 1800–12 (2011).

65. Nowick, K. et al. Gain, loss and divergence in primate zinc-finger genes: a rich resource for evolution of gene regulatory differences between species. PLoS One 6, e21553 (2011).

66. Nowick, K., Hamilton, A.T., Zhang, H. & Stubbs, L. Rapid sequence and expression divergence suggest selection for novel function in primate-specific KRAB-ZNF genes. Mol Biol Evol 27, 2606–17 (2010).

67. Alexander, T.B. et al. The genetic basis and cell of origin of mixed phenotype acute leukaemia. Nature 562, 373–379 (2018).

68. Arber, D.A. et al. International Consensus Classification of Myeloid Neoplasms and Acute Leukemias: integrating morphologic, clinical, and genomic data. Blood 140, 1200–1228 (2022).

69. Denny, P. & Ashworth, A. A zinc finger protein-encoding gene expressed in the post-meiotic phase of spermatogenesis. Gene 106, 221–7 (1991).

70. Lord, T. et al. A novel high throughput screen to identify candidate molecular networks that regulate spermatogenic stem cell functionsdagger. Biol Reprod 106, 1175–1190 (2022).

71. Consortium, G.T. The Genotype-Tissue Expression (GTEx) project. Nat Genet 45, 580–5 (2013).

72. Schmidt, D. et al. ChIP-seq: using high-throughput sequencing to discover protein-DNA interactions. Methods 48, 240–8 (2009).

73. Langmead, B. & Salzberg, S.L. Fast gapped-read alignment with Bowtie 2. Nat Methods 9, 357–9 (2012).

74. Li, H. et al. The Sequence Alignment/Map format and SAMtools. Bioinformatics 25, 2078–9 (2009).

75. Danecek, P. et al. Twelve years of SAMtools and BCFtools. Gigascience 10(2021).

76. Feng, J., Liu, T., Qin, B., Zhang, Y. & Liu, X.S. Identifying ChIP-seq enrichment using MACS. Nat Protoc 7, 1728–40 (2012).

77. Liu, T. Use model-based Analysis of ChIP-Seq (MACS) to analyze short reads generated by sequencing protein-DNA interactions in embryonic stem cells. Methods Mol Biol 1150, 81–95 (2014).

78. Quinlan, A.R. & Hall, I.M. BEDTools: a flexible suite of utilities for comparing genomic features. Bioinformatics 26, 841–2 (2010).

79. Ramirez, F., Dundar, F., Diehl, S., Gruning, B.A. & Manke, T. deepTools: a flexible platform for exploring deep-sequencing data. Nucleic Acids Res 42, W187–91 (2014).

80. Grandi, F.C., Modi, H., Kampman, L. & Corces, M.R. Chromatin accessibility profiling by ATAC-seq. Nat Protoc 17, 1518–1552 (2022).

81. Consortium, E.P. et al. Perspectives on ENCODE. Nature 583, 693–698 (2020).

82. Frankish, A. et al. GENCODE: reference annotation for the human and mouse genomes in 2023. Nucleic Acids Res 51, D942–D949 (2023).

83. Dixon, J.R. et al. Topological domains in mammalian genomes identified by analysis of chromatin interactions. Nature 485, 376–80 (2012).

84. Satopaa, V., Albrecht, J., Irwin, D. & Raghavan, B. Finding a “kneedle” in a haystack: Detecting knee points in system behavior. in 2011 31st international conference on distributed computing systems workshops 166–171 (IEEE, 2011).

85. Karimzadeh, M., Ernst, C., Kundaje, A. & Hoffman, M.M. Umap and Bismap: quantifying genome and methylome mappability. Nucleic Acids Res 46, e120 (2018).

86. Korhonen, J., Martinmaki, P., Pizzi, C., Rastas, P. & Ukkonen, E. MOODS: fast search for position weight matrix matches in DNA sequences. Bioinformatics 25, 3181–2 (2009).

87. Armstrong, J. et al. Progressive Cactus is a multiple-genome aligner for the thousand-genome era. Nature 587, 246–251 (2020).

88. Amemiya, H.M., Kundaje, A. & Boyle, A.P. The ENCODE Blacklist: Identification of Problematic Regions of the Genome. Sci Rep 9, 9354 (2019).

89. Jurka, J. Repbase update: a database and an electronic journal of repetitive elements. Trends Genet 16, 418–20 (2000).

90. Virtanen, P. et al. SciPy 1.0: fundamental algorithms for scientific computing in Python. Nat Methods 17, 261–272 (2020).

91. Storer, J., Hubley, R., Rosen, J., Wheeler, T.J. & Smit, A.F. The Dfam community resource of transposable element families, sequence models, and genome annotations. Mob DNA 12, 2 (2021).

92. Oughtred, R. et al. The BioGRID database: A comprehensive biomedical resource of curated protein, genetic, and chemical interactions. Protein Sci 30, 187–200 (2021).

93. Lambert, S.A., Albu, M., Hughes, T.R. & Najafabadi, H.S. Motif comparison based on similarity of binding affinity profiles. Bioinformatics 32, 3504–3506 (2016).

