## Supplementary figures and images for "Extensive binding of nebulous human transcription factors to genomic dark matter"

### Supplementary Document 1

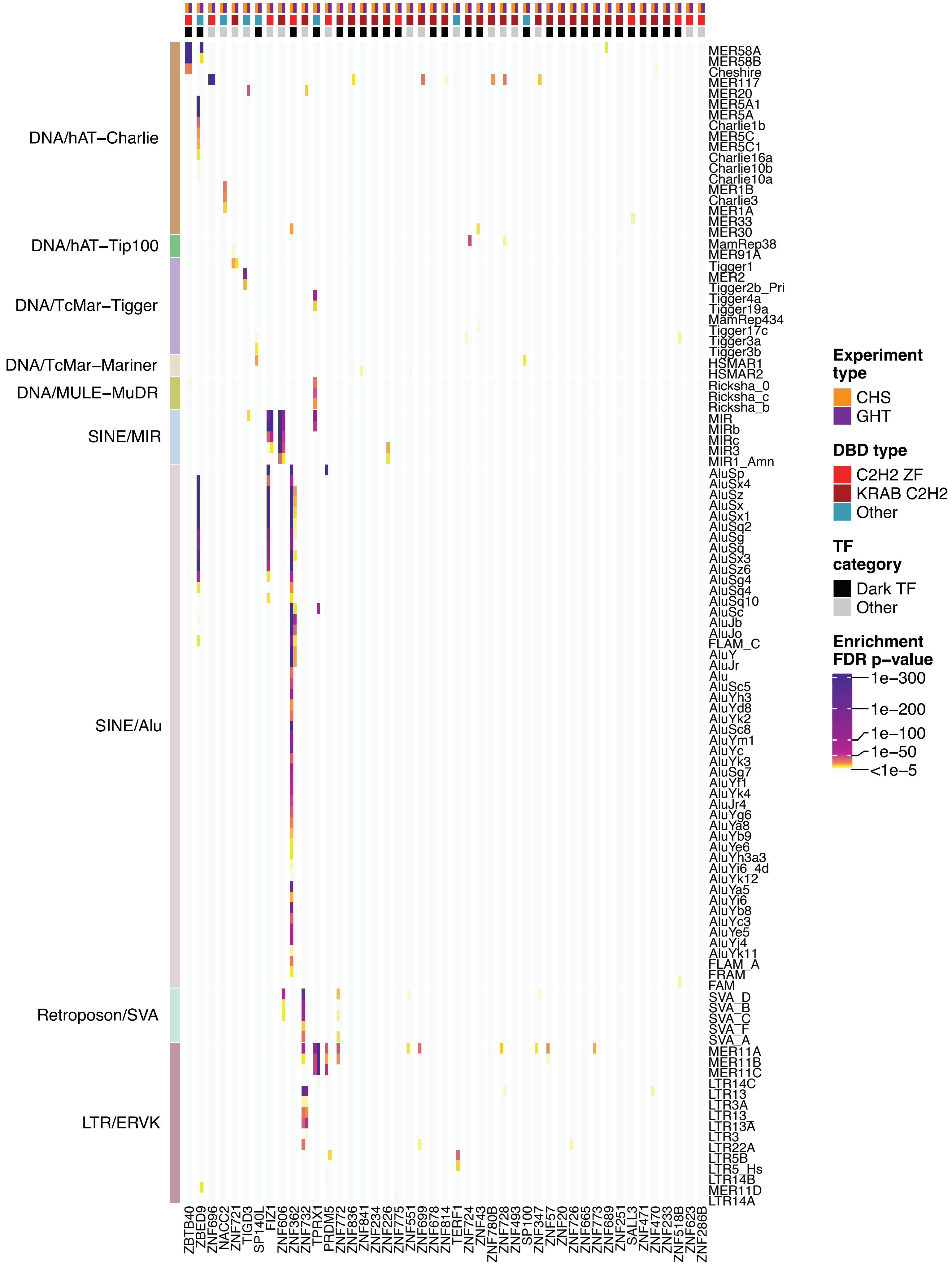

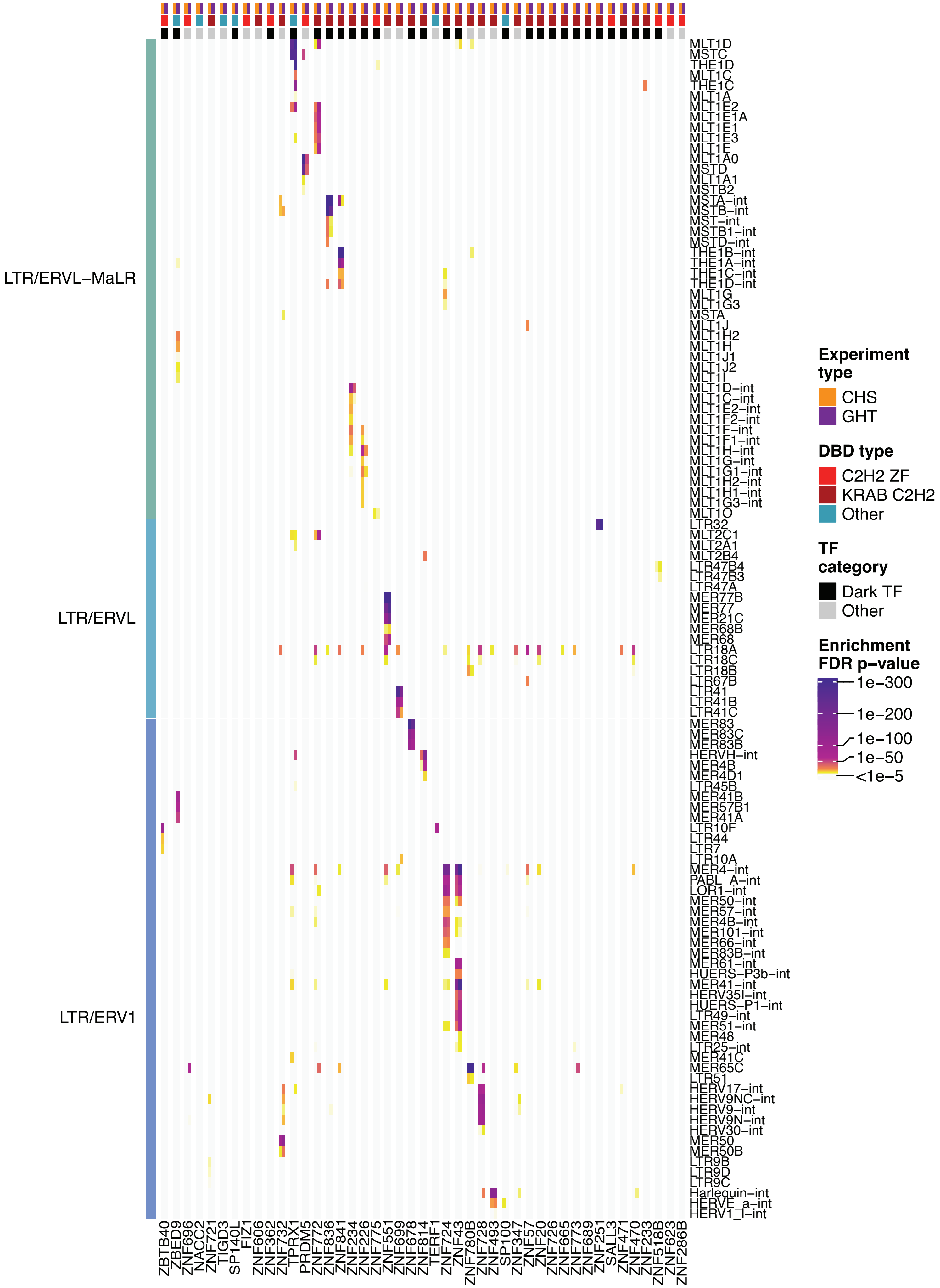

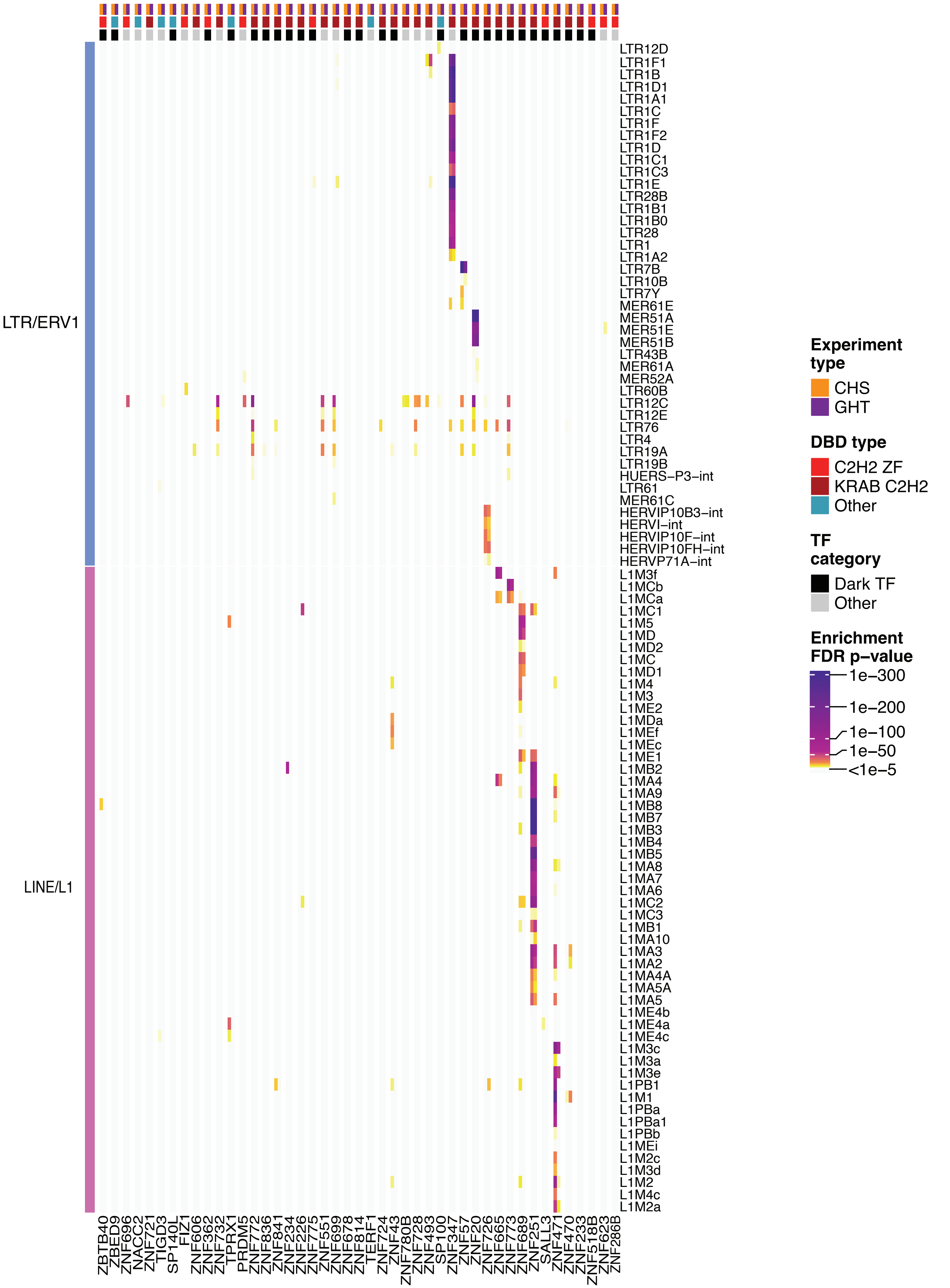

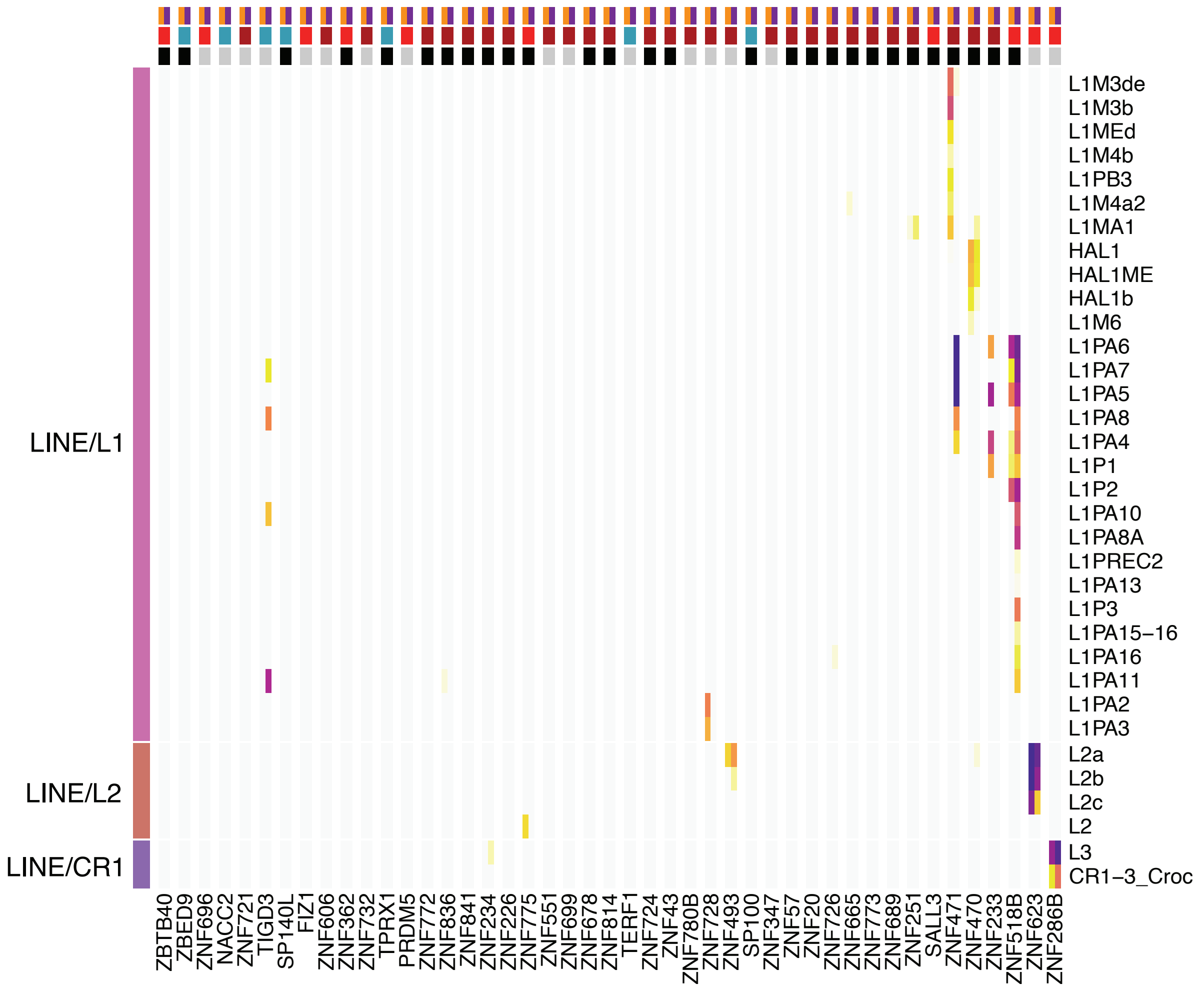
