## Supplementary Document 2 for "Extensive binding of nebulous human transcription factors to genomic dark matter"

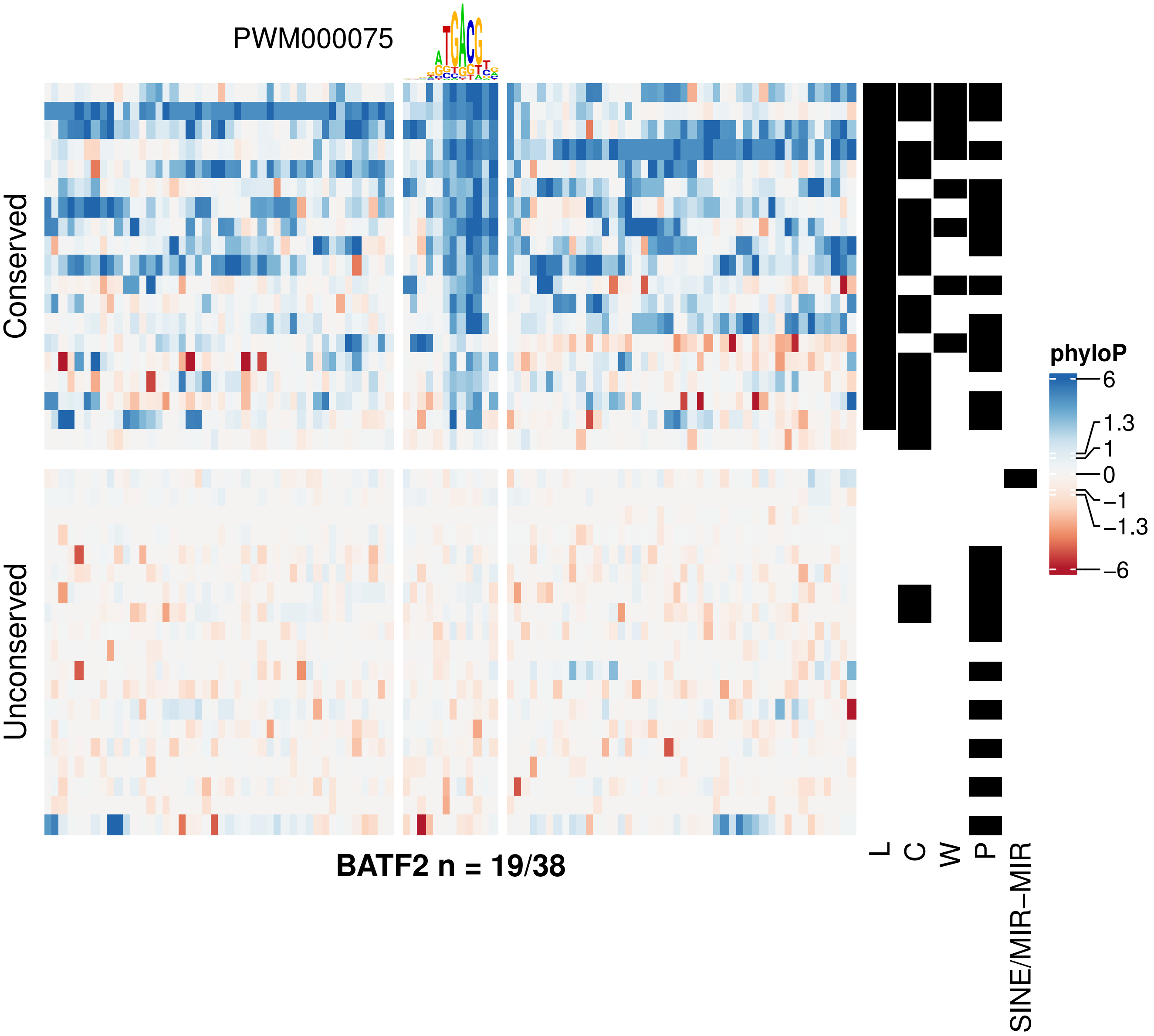
PWM000075
Conserved
Unconserved
BATF2 n = 19/38
L
C
W
P
SINE/MIR−MIR
phyloP
6
1.3
1
0
−1
−1.3
−6

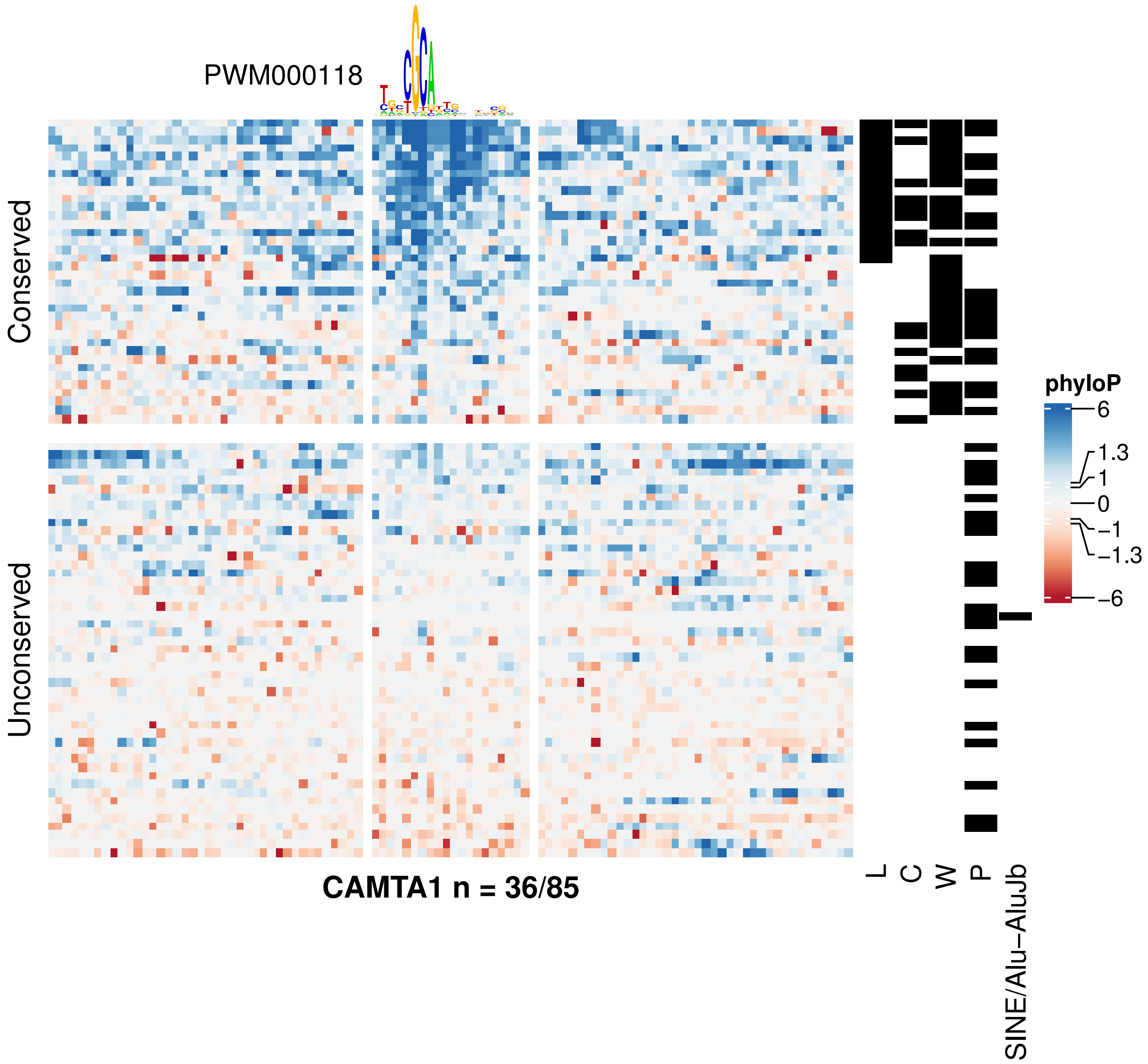
PWM000118
Conserved
Unconserved
CAMTA1 n = 36/85
L
C
W
P
SINE/Alu−AluJb
phyloP
6
1.3
1
0
−1
−1.3
−6

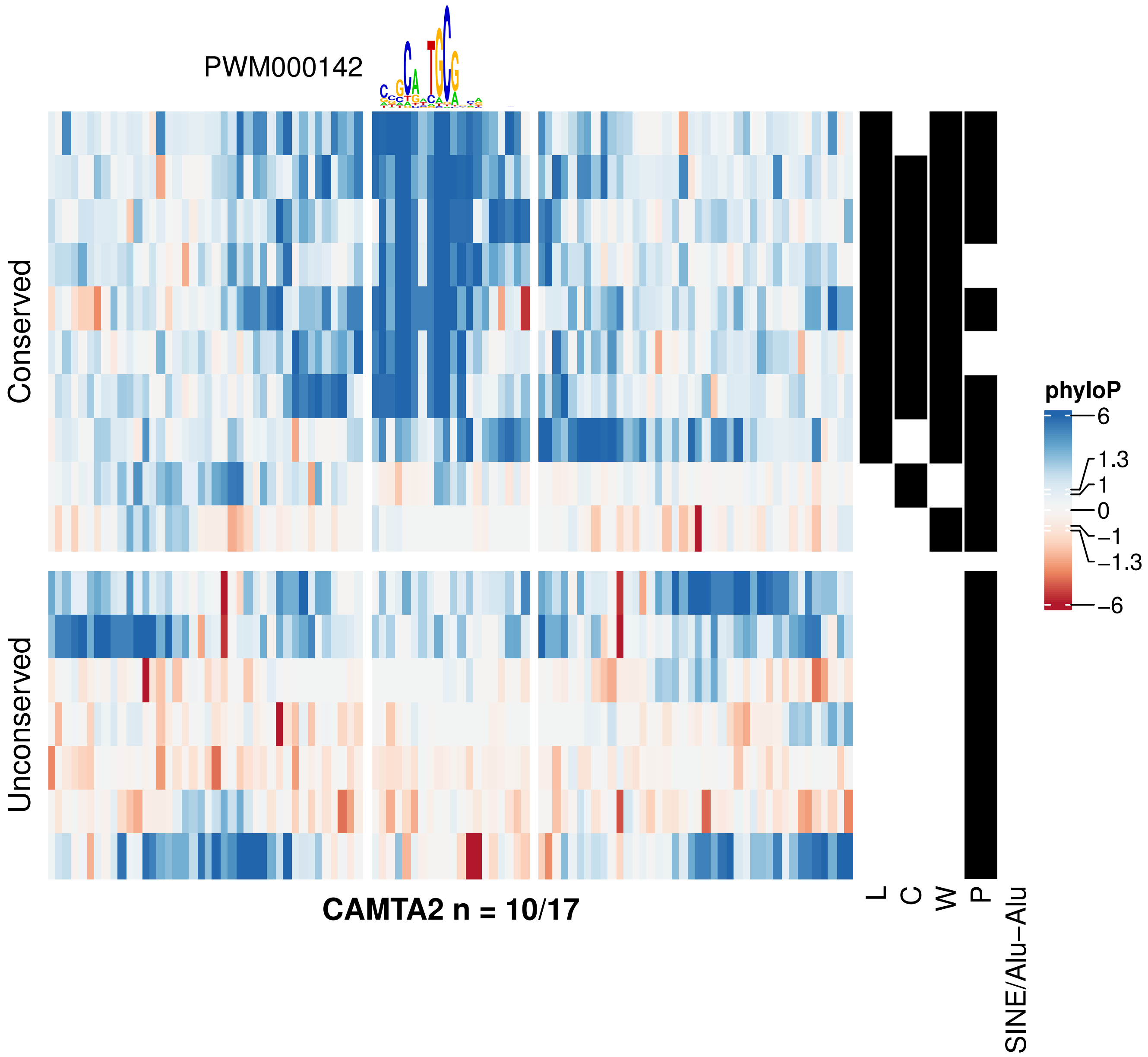

PWM000142
Conserved
Unconserved
CAMTA2 n = 10/17
L
C
W
P
SINE/Alu−Alu
phyloP
6
1.3
1
0
−1
−1.3
−6

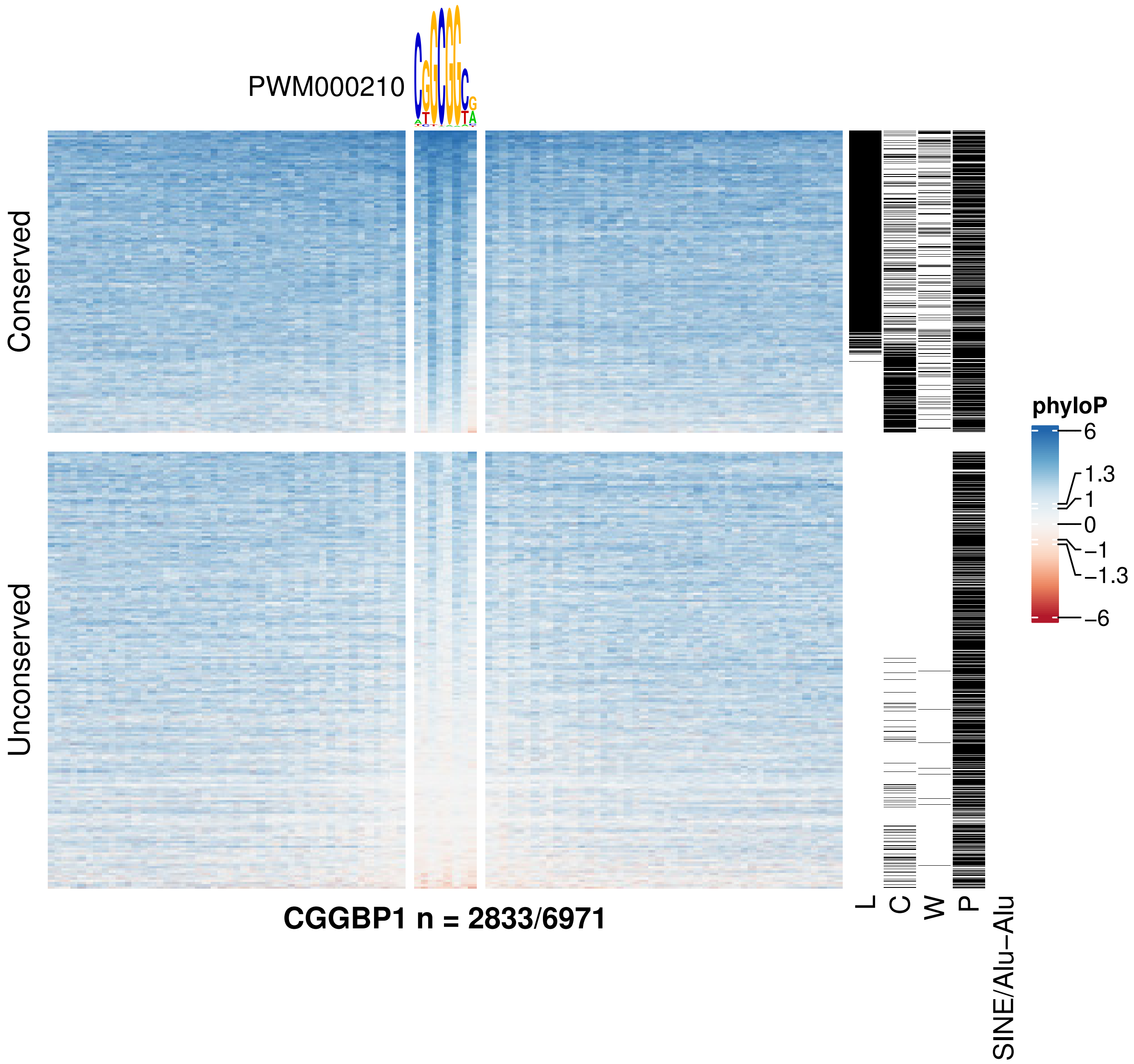
PWM000210
Conserved
Unconserved
CGGBP1 n = 2833/6971
L
C
W
P
SINE/Alu−Alu
phyloP
6
1.3
1
0
−1
−1.3
−6

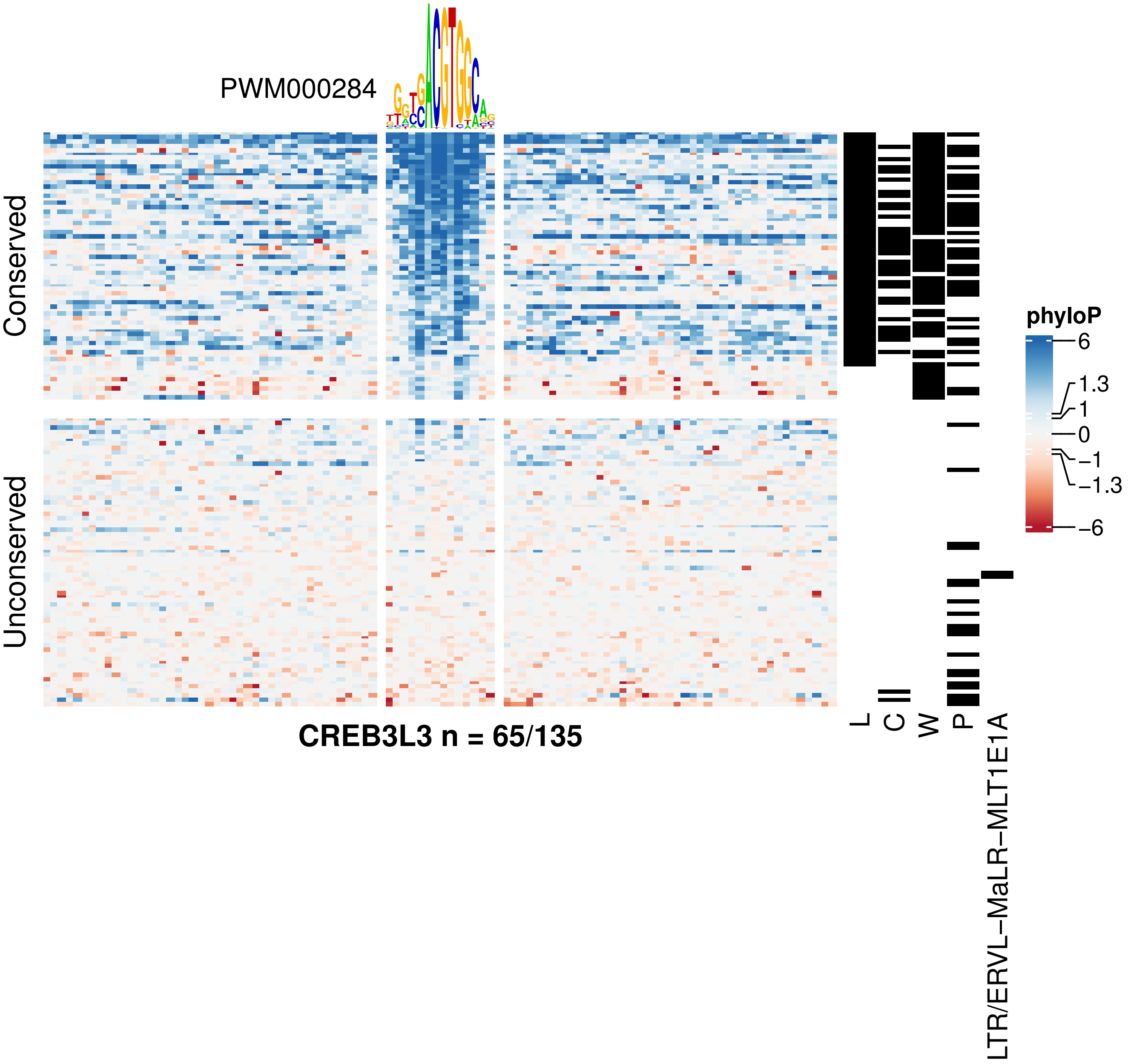

PWM000284
Conserved
Unconserved
CREB3L3 n = 65/135
L
C
W
P
LTR/ERVL−MaLR−MLT1E1A
phyloP
6
1.3
1
0
−1
−1.3
−6

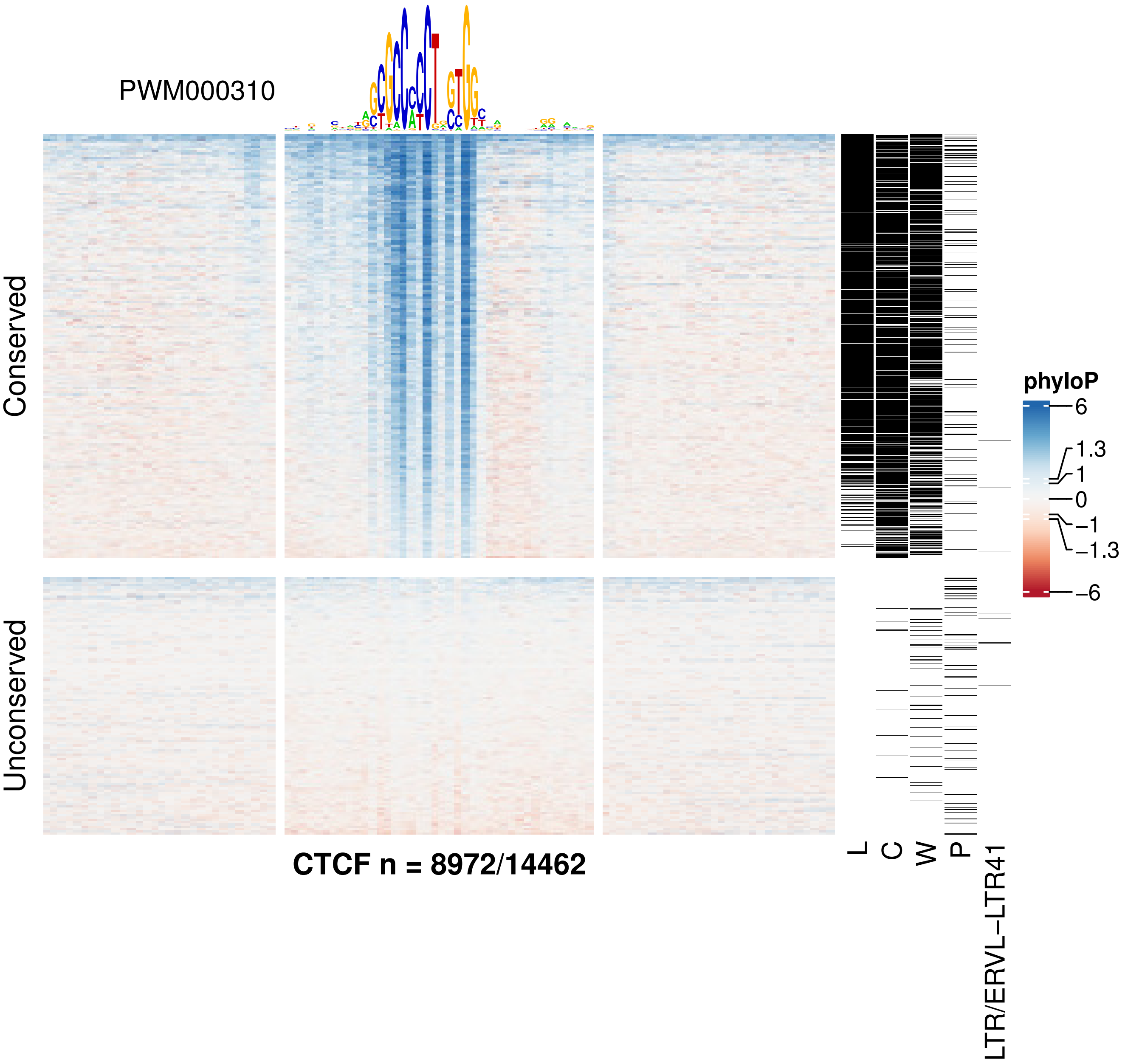

PWM000310
Conserved
Unconserved
CTCF n = 8972/14462
L
C
W
P
LTR/ERVL−LTR41
phyloP
6
1.3
1
0
−1
−1.3
−6

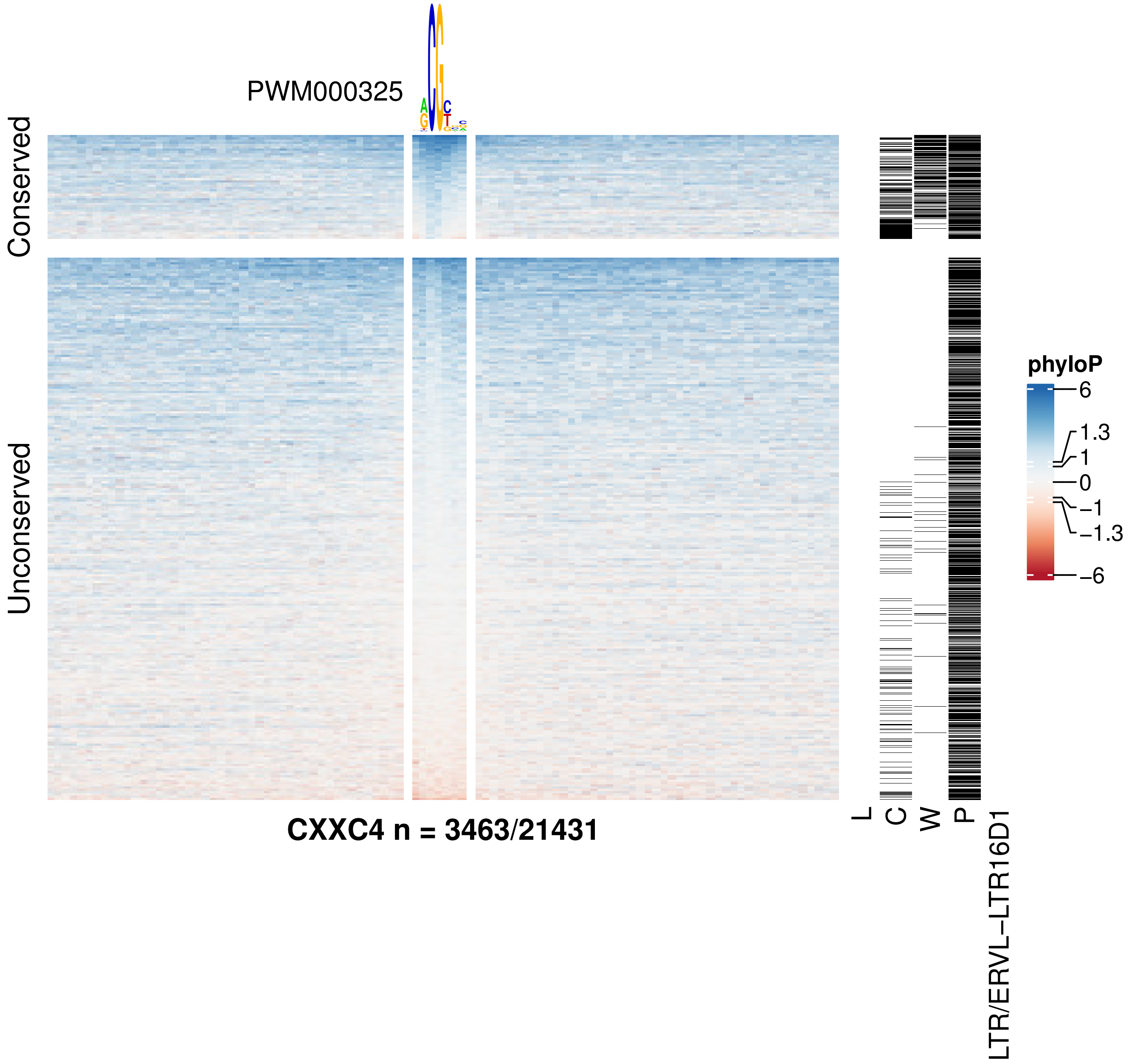
PWM000325
Conserved
Unconserved
CXXC4 n = 3463/21431
L
C
W
P
LTR/ERVL−LTR16D1
phyloP
6
1.3
1
0
−1
−1.3
−6

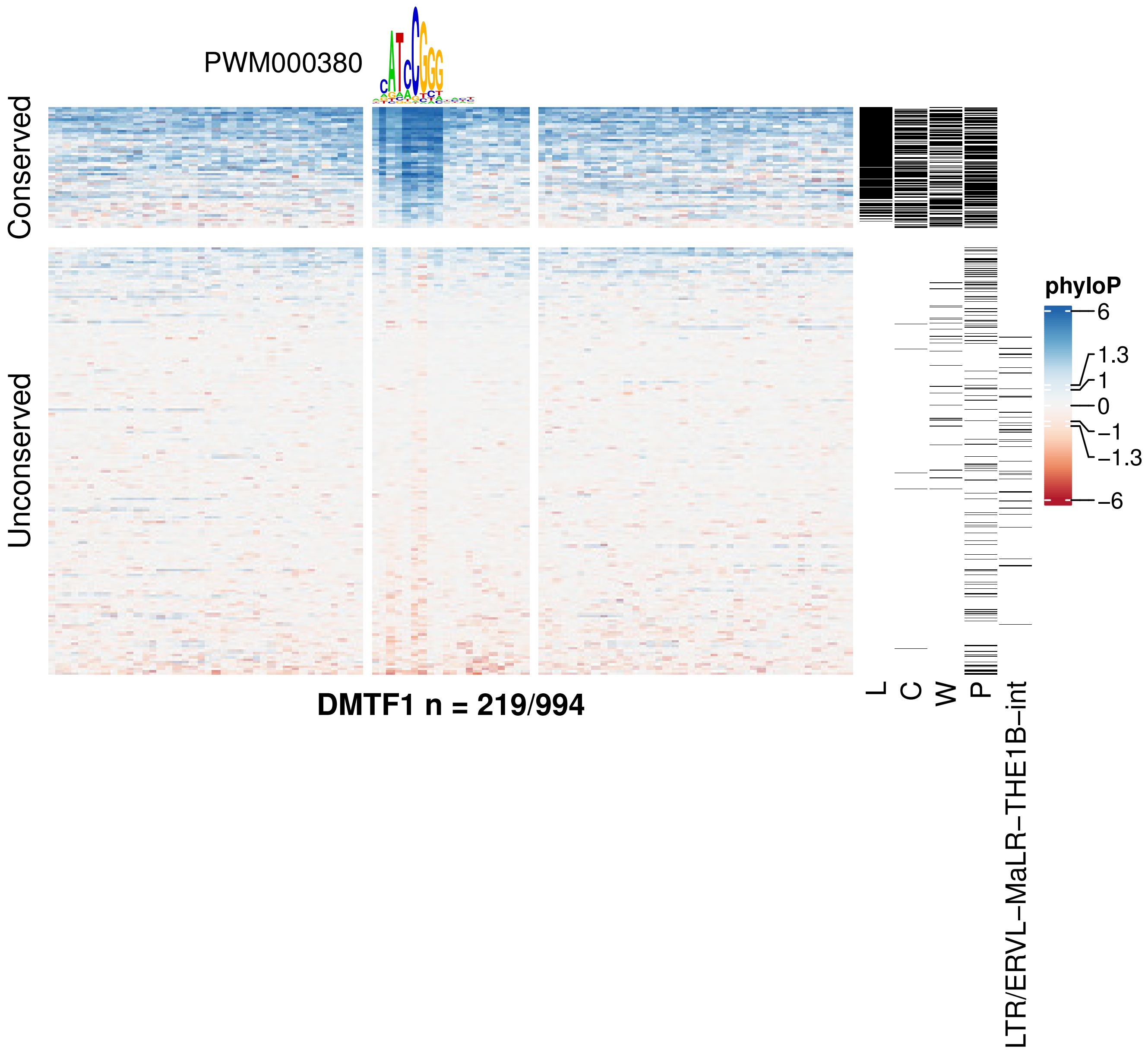
PWM000380
Conserved
Unconserved
DMTF1 n = 219/994
L
C
W
P
LTR/ERVL−MaLR−THE1B−int
phyloP
6
1.3
1
0
−1
−1.3
−6

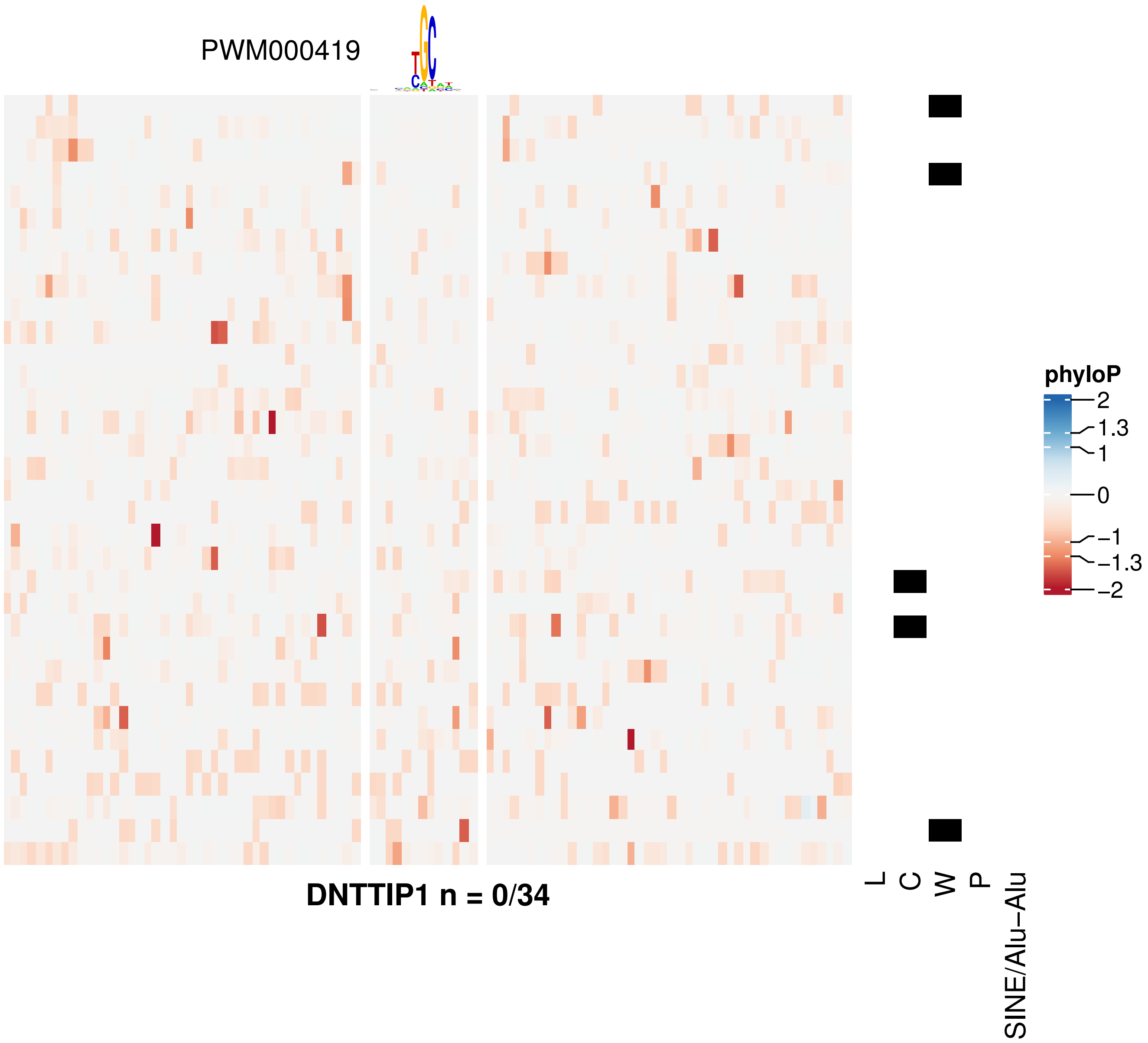
PWM000419
DNTTIP1 n = 0/34
L
C
W
P
SINE/Alu−Alu
phyloP
2
1.3
1
0
−1
−1.3
−2

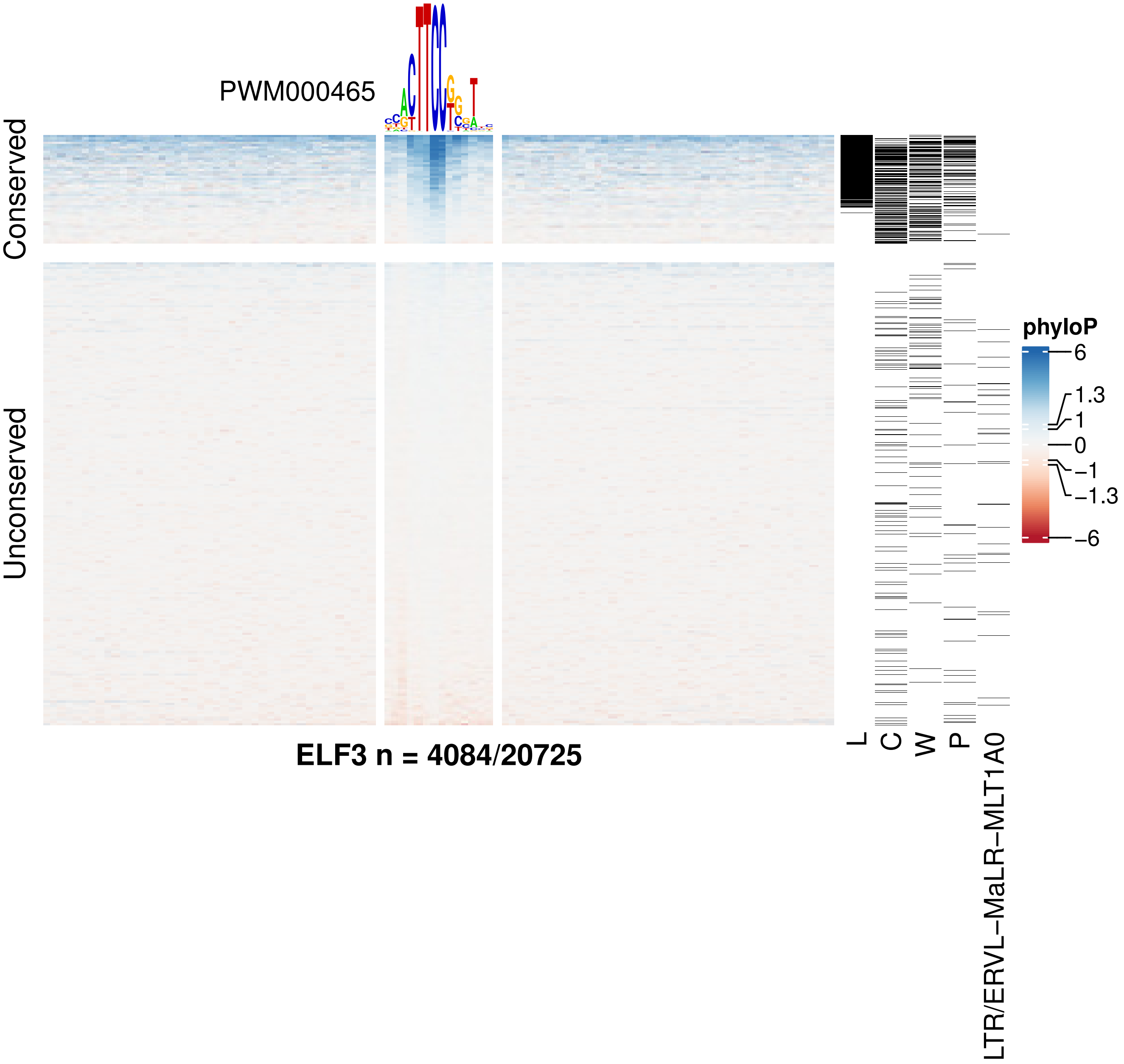
PWM000465
Conserved
Unconserved
ELF3 n = 4084/20725
L
C
W
P
LTR/ERVL−MaLR−MLT1A0
phyloP
6
1.3
1
0
−1
−1.3
−6

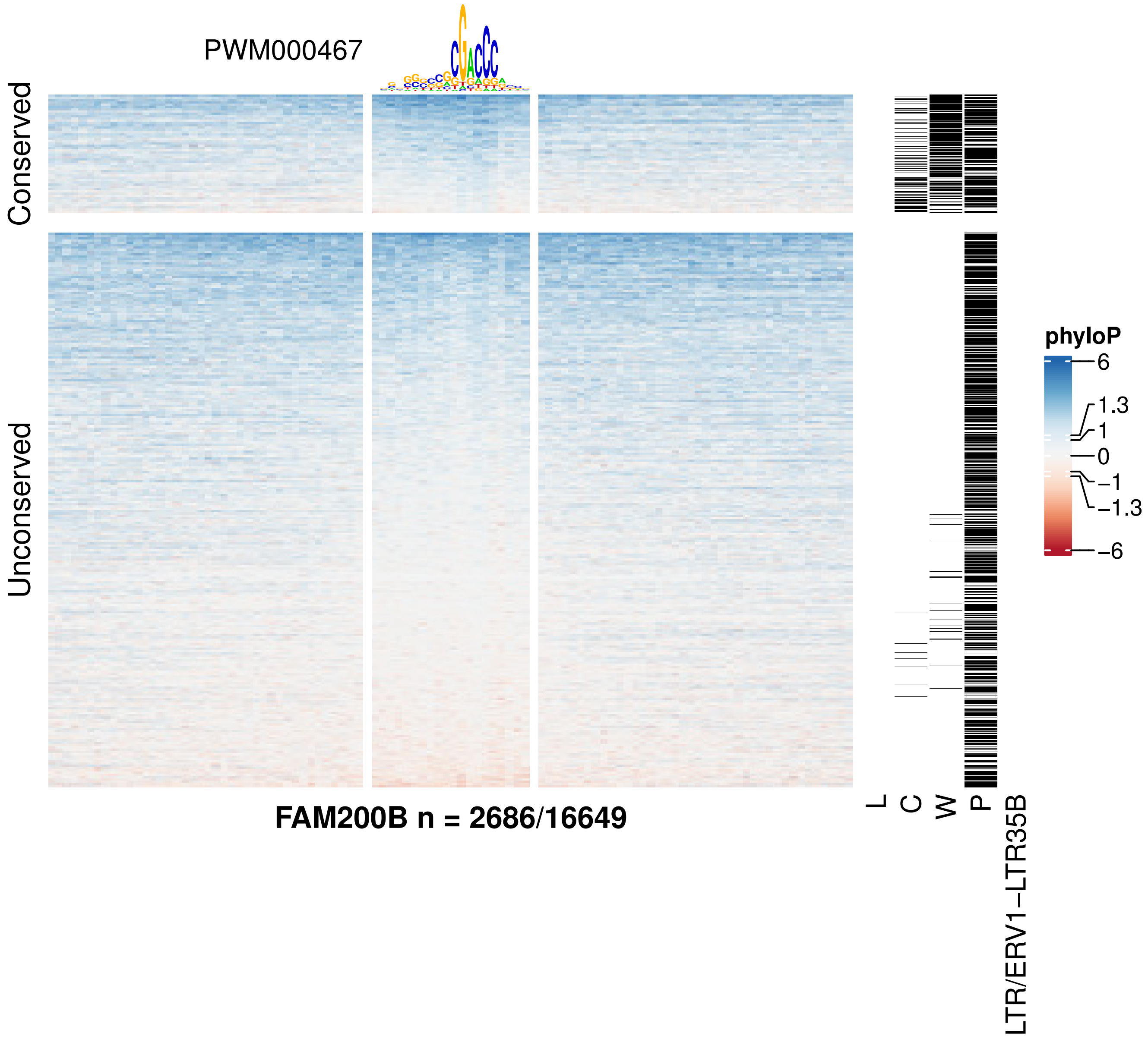

PWM000467
Conserved
Unconserved
FAM200B n = 2686/16649
L
C
W
P
LTR/ERV1−LTR35B
phyloP
6
1.3
1
0
−1
−1.3
−6

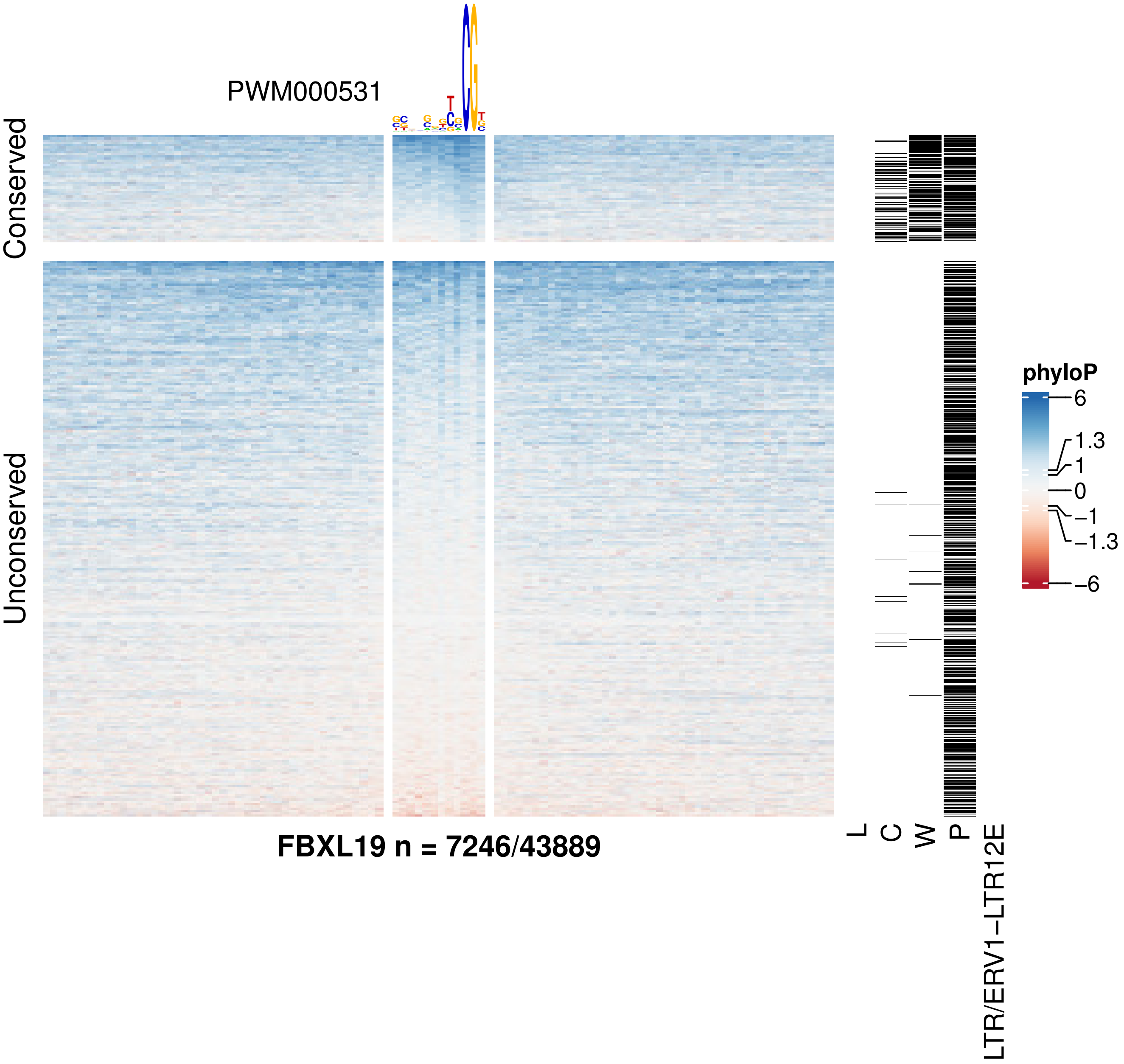
PWM000531
Conserved
Unconserved
FBXL19 n = 7246/43889
L
C
W
P
LTR/ERV1−LTR12E
phyloP
6
1.3
1
0
−1
−1.3
−6

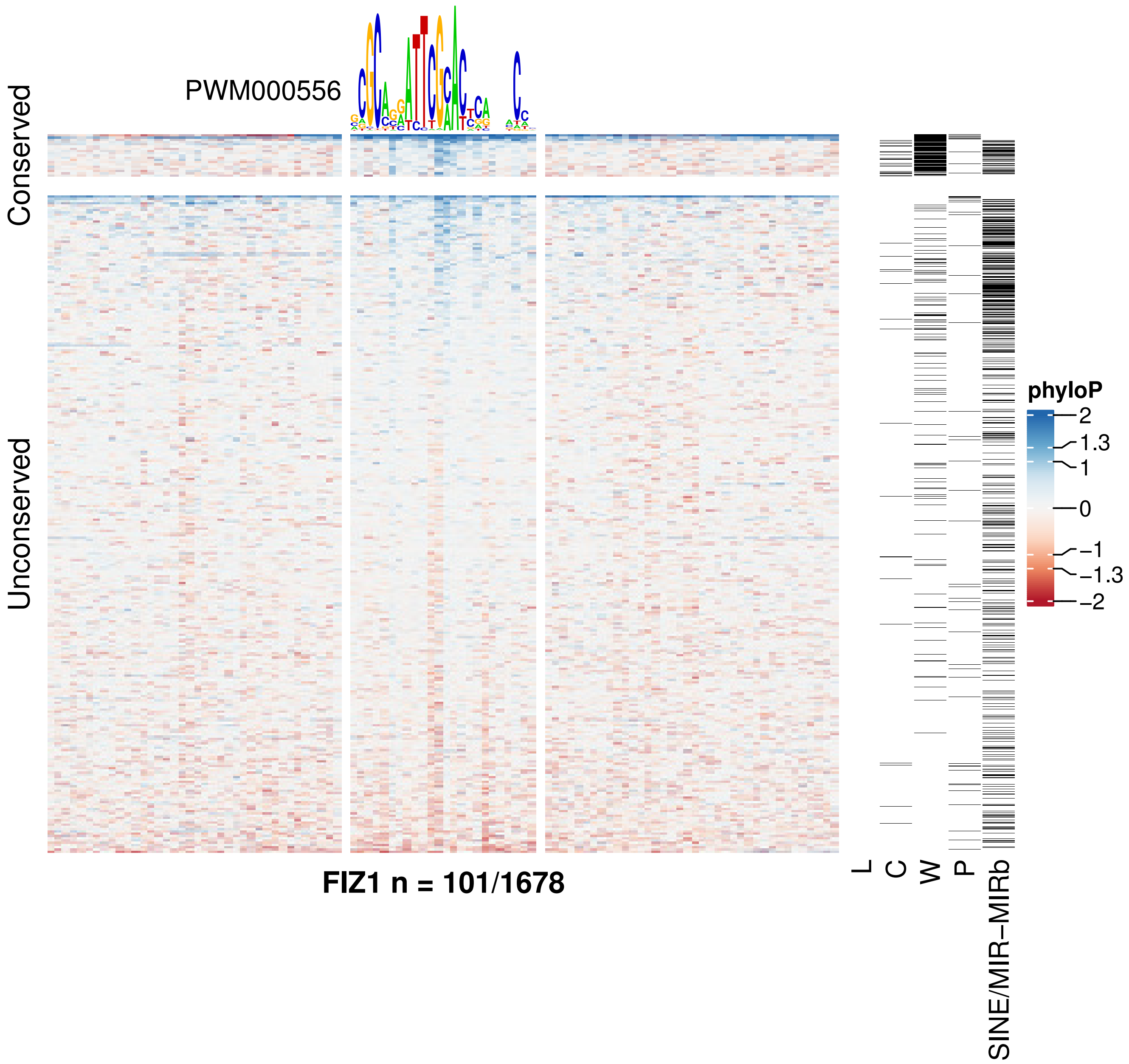

PWM000556
Conserved
Unconserved
FIZ1 n = 101/1678
L
C
W
P
SINE/MIR−MIRb
phyloP
2
1.3
1
0
−1
−1.3
−2

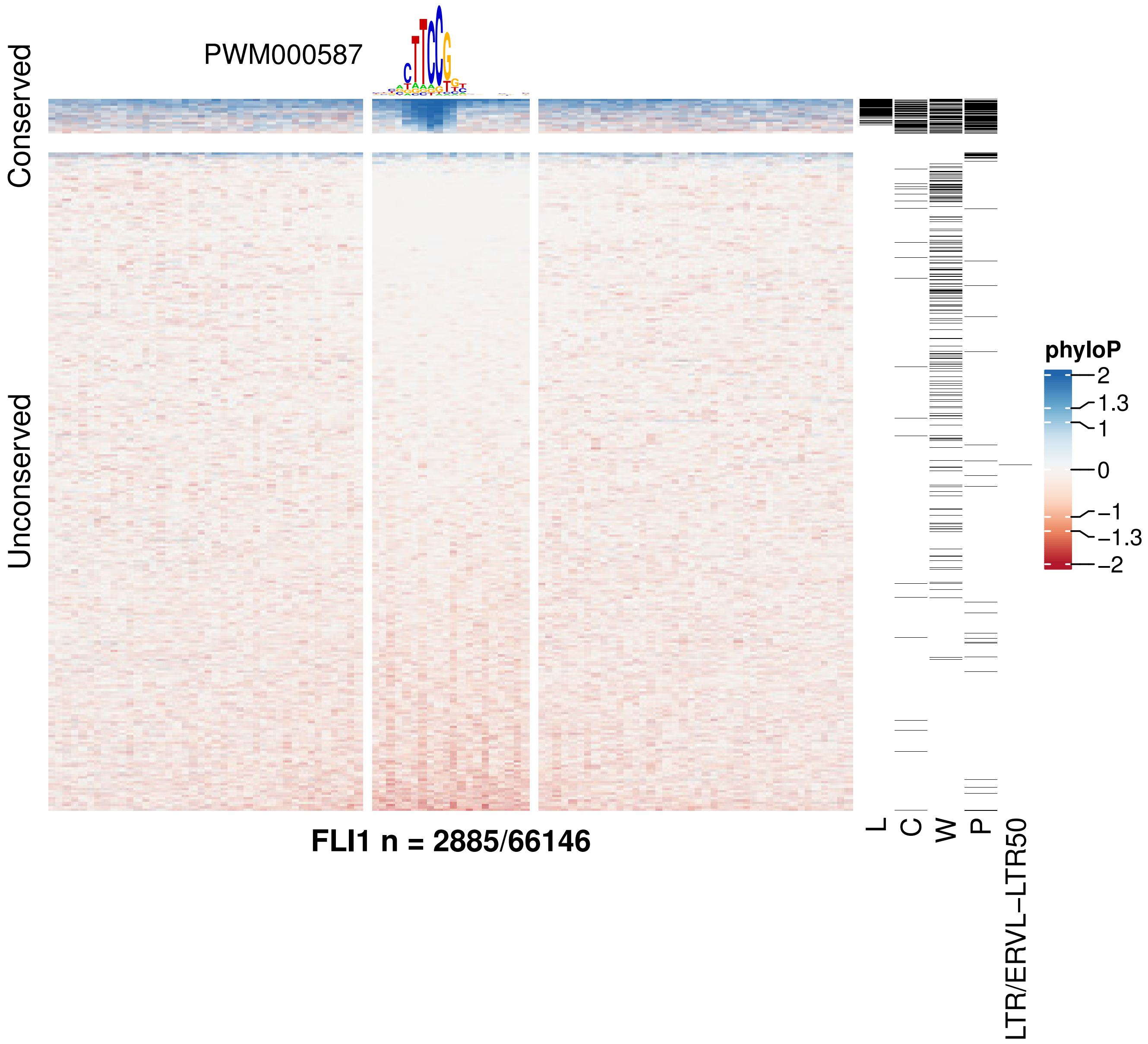

PWM000587
Conserved
Unconserved
FLI1 n = 2885/66146
L
C
W
P
LTR/ERVL−LTR50
phyloP
2
1.3
1
0
−1
−1.3
−2

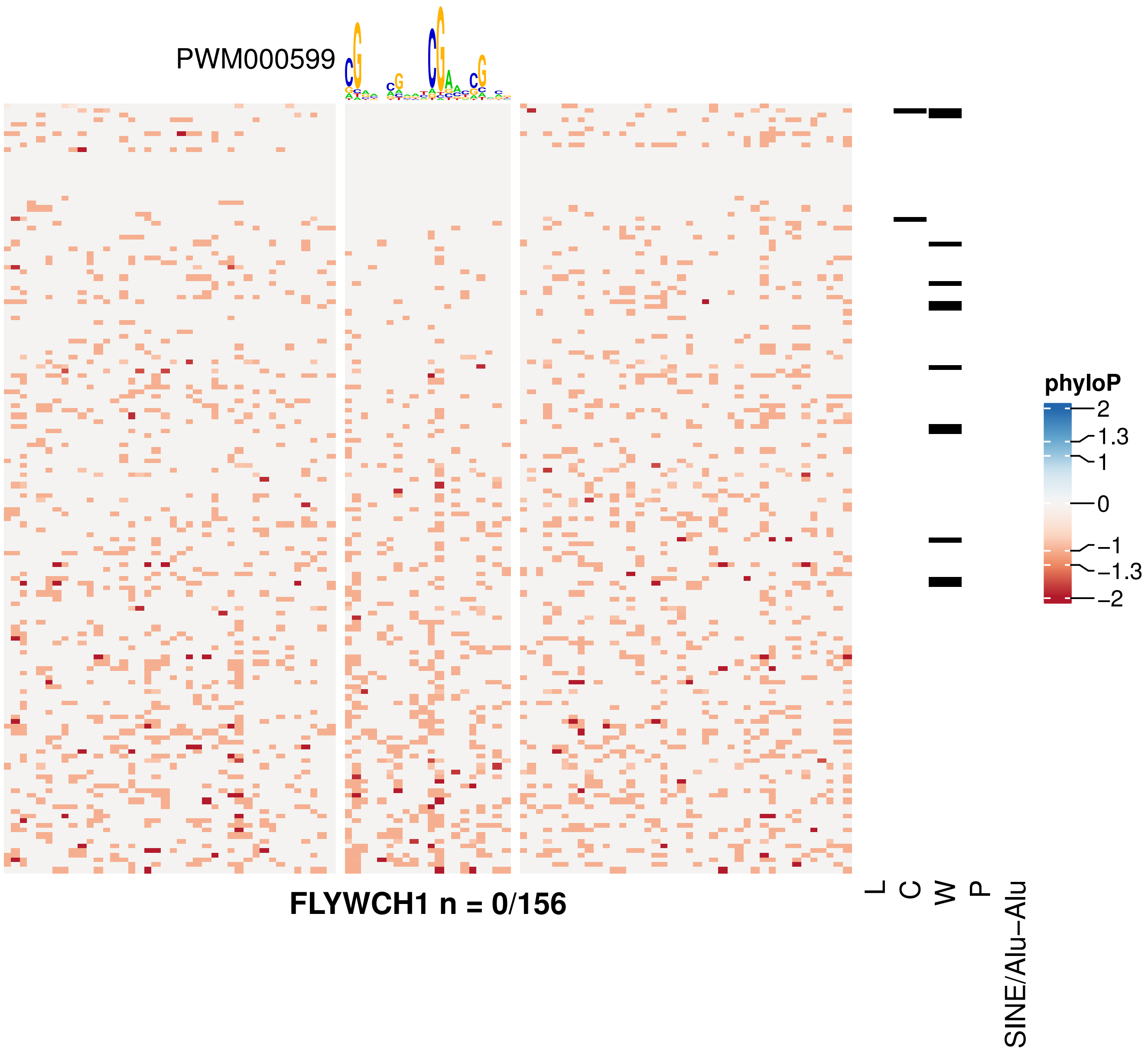

PWM000599
phyloP
2
1.3
1
0
−1
−1.3
−2
FLYWCH1 n = 0/156
L
C
W
P
SINE/Alu−Alu

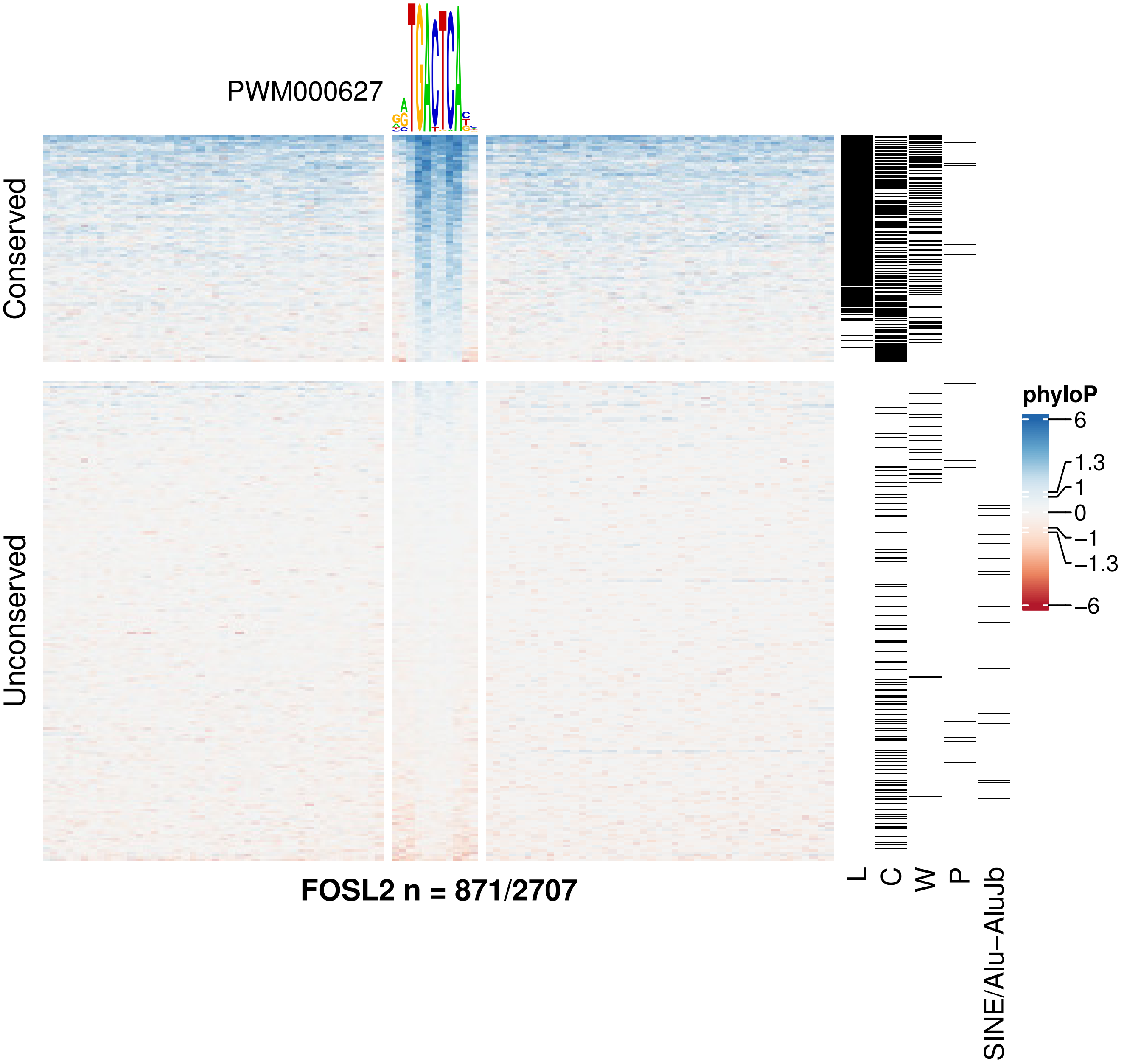
PWM000627
Conserved
Unconserved
phyloP
6
1.3
1
0
−1
−1.3
−6
L
C
W
P
SINE/Alu−AluJb
FOSL2 n = 871/2707

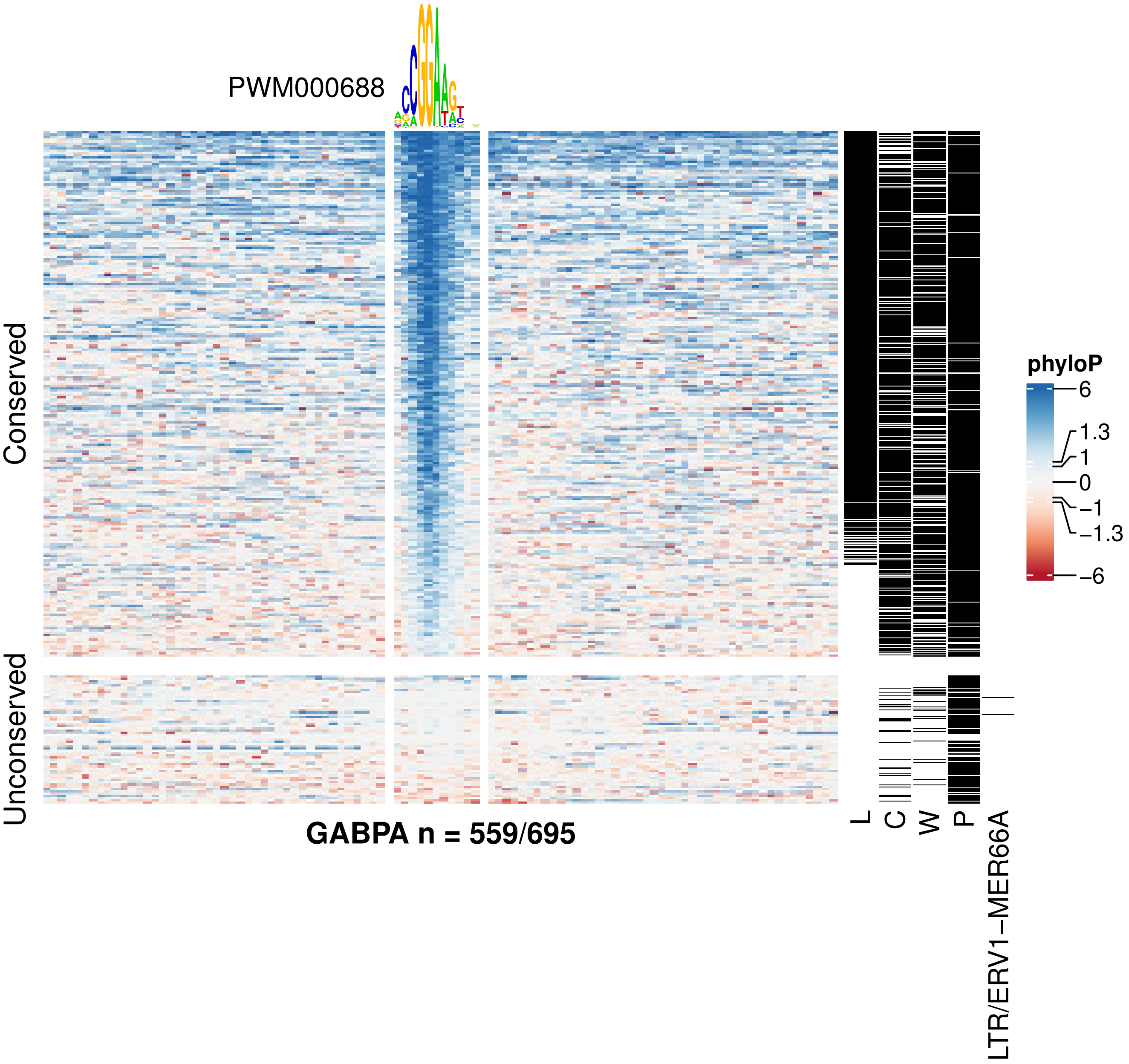
PWM000688
Conserved
Unconserved
GABPA n = 559/695
L
C
W
P
LTR/ERV1−MER66A
phyloP
6
1.3
1
0
−1
−1.3
−6

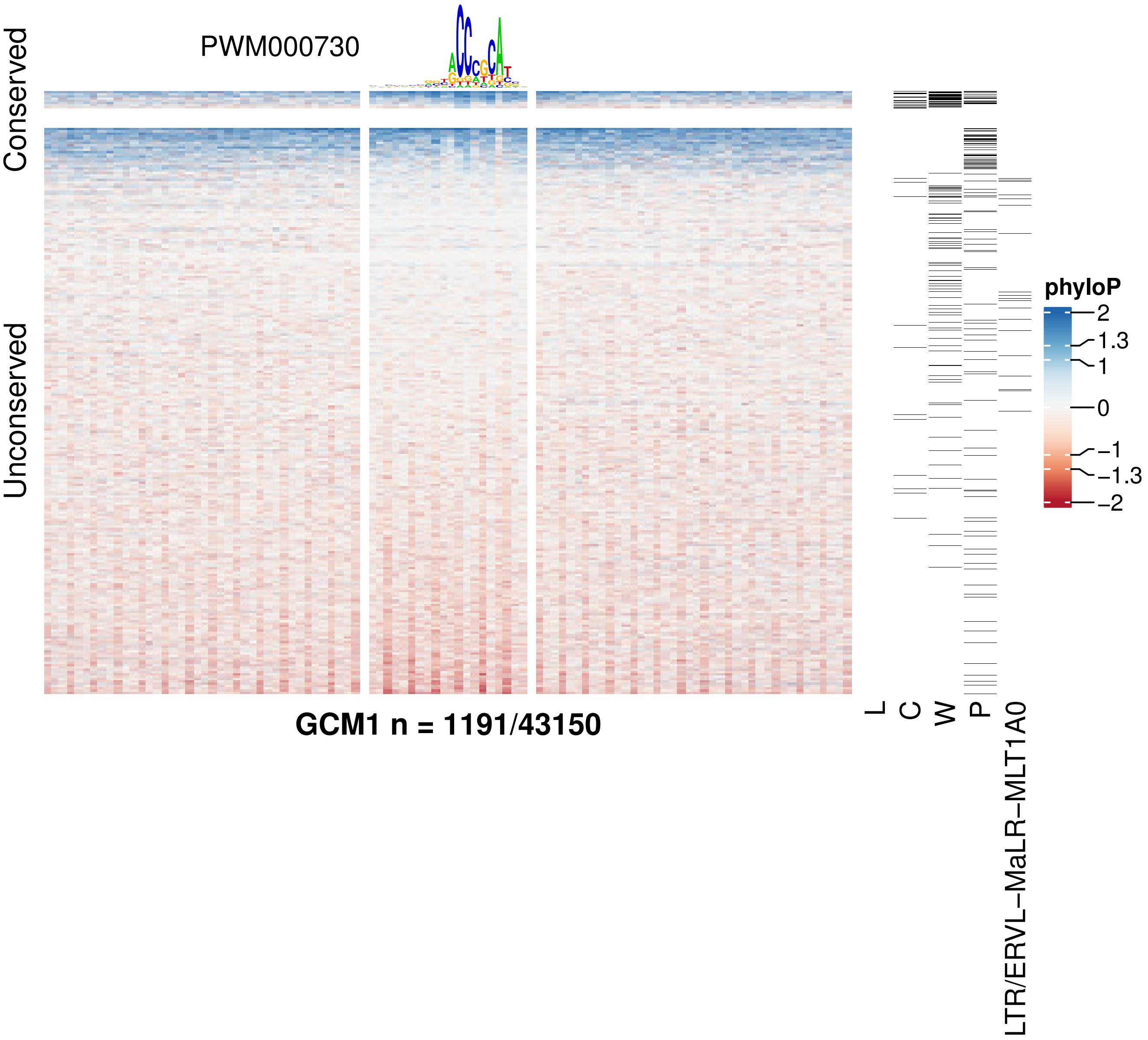

PWM000730
Conserved
Unconserved
GCM1 n = 1191/43150
L
C
W
P
LTR/ERVL−MaLR−MLT1A0
phyloP
2
1.3
1
0
−1
−1.3
−2

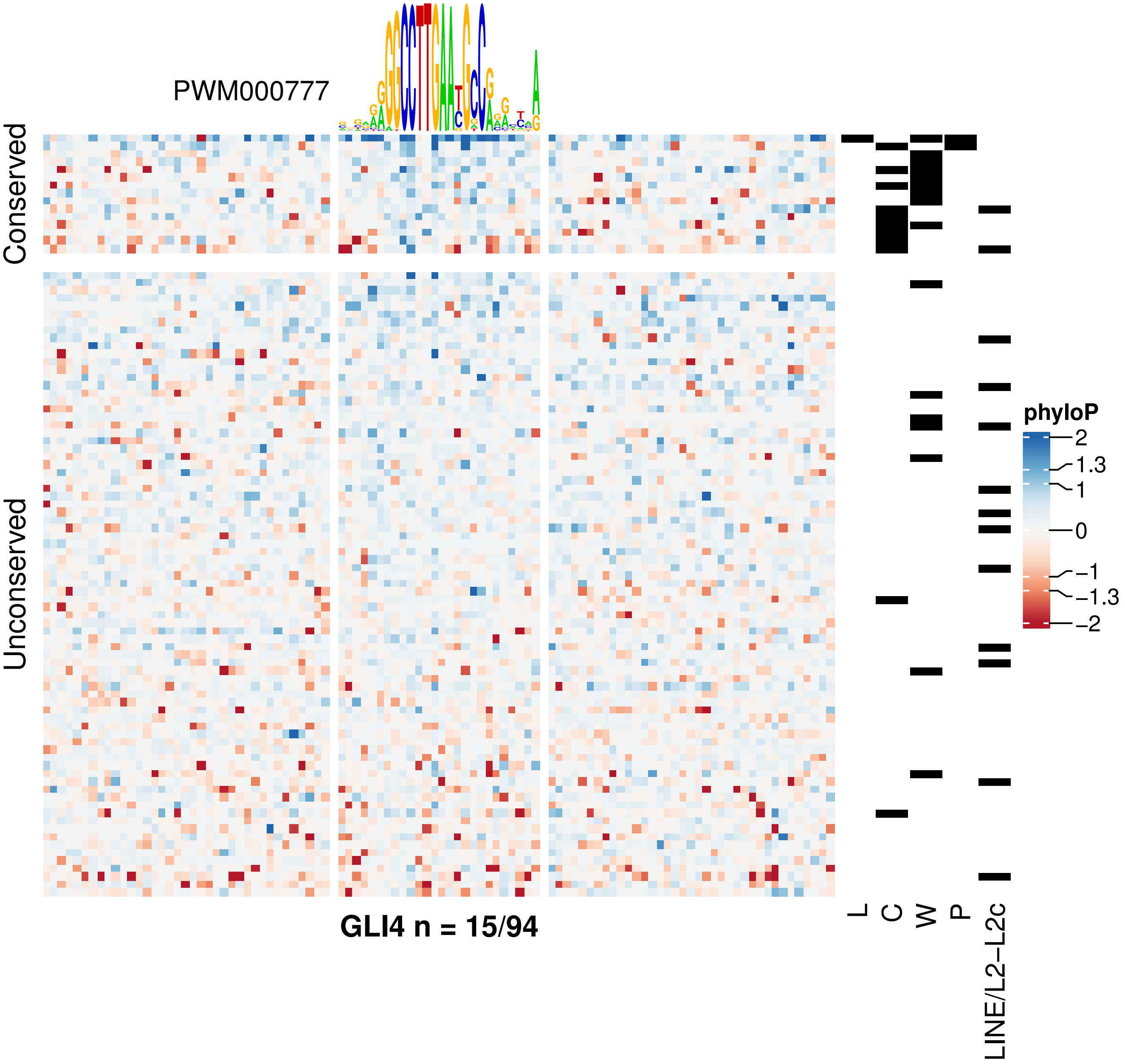
PWM000777
Conserved
Unconserved
GLI4 n = 15/94
L
C
W
P
LINE/L2−L2c
phyloP
2
1.3
1
0
−1
−1.3
−2

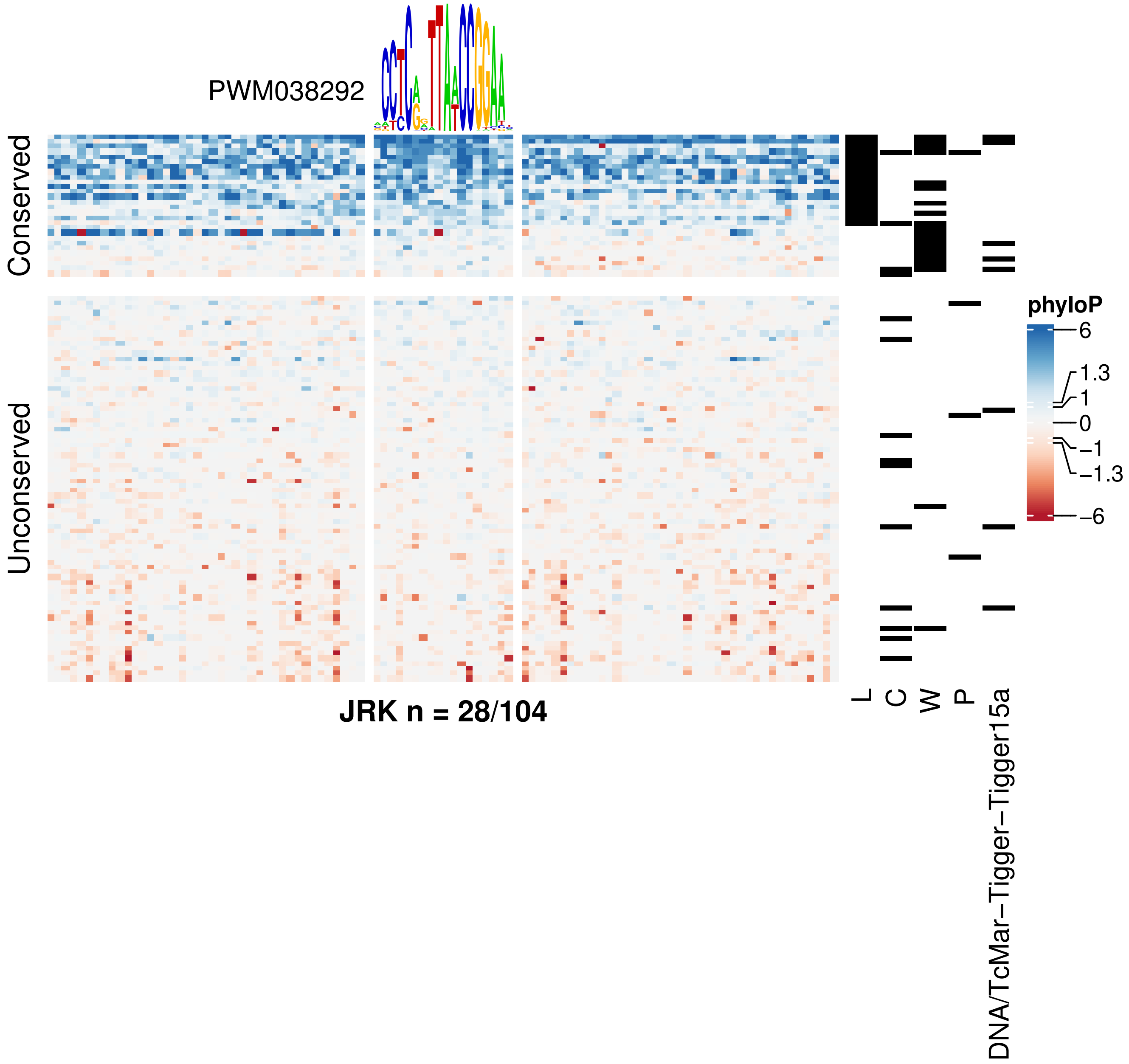
PWM038292
Conserved
Unconserved
JRK n = 28/104
L
C
W
P
DNA/TcMar−Tigger−Tigger15a
phyloP
6
1.3
1
0
−1
−1.3
−6

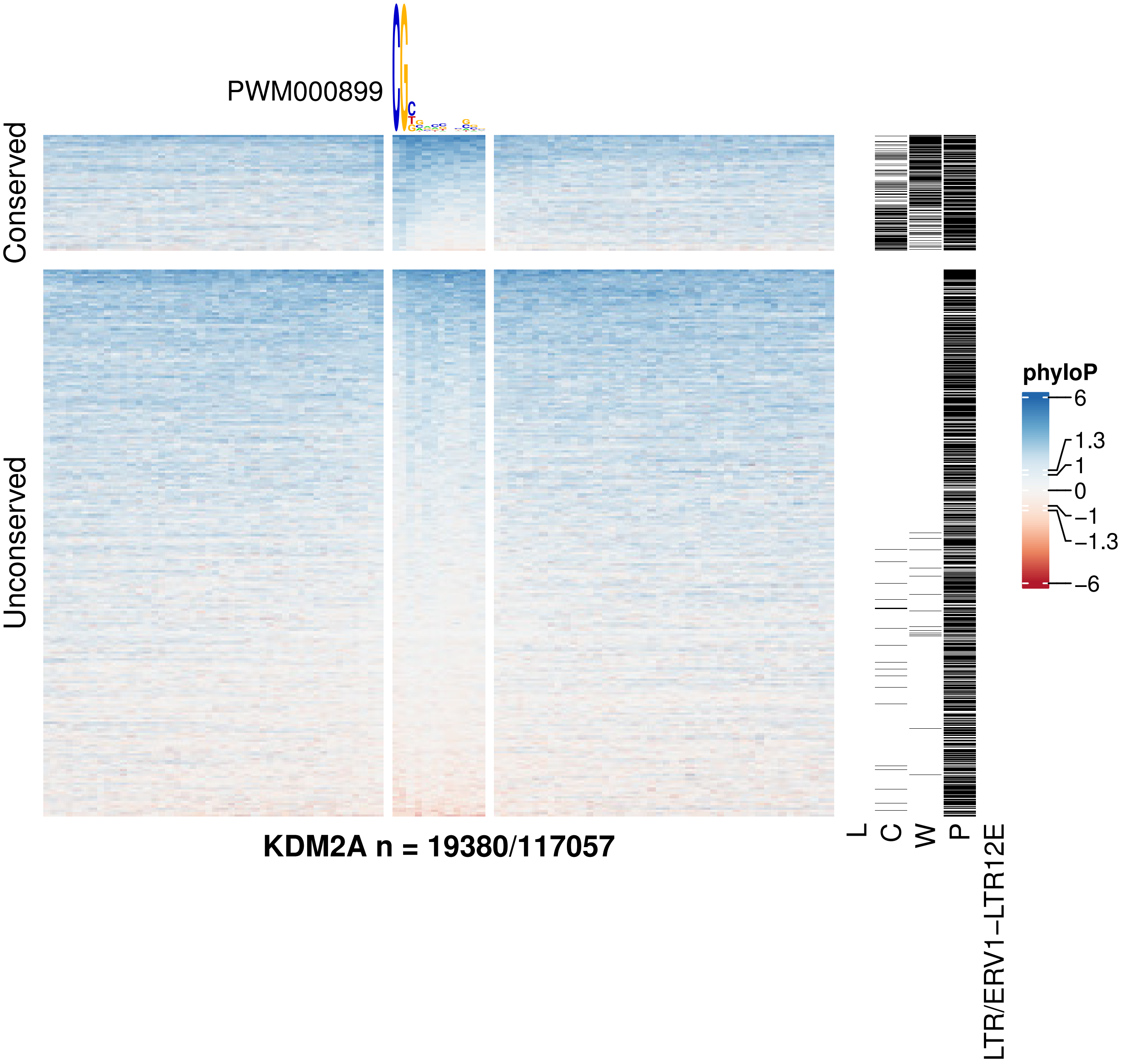
PWM000899
Conserved
Unconserved
KDM2A n = 19380/117057
L
C
W
P
LTR/ERV1−LTR12E
phyloP
6
1.3
1
0
−1
−1.3
−6

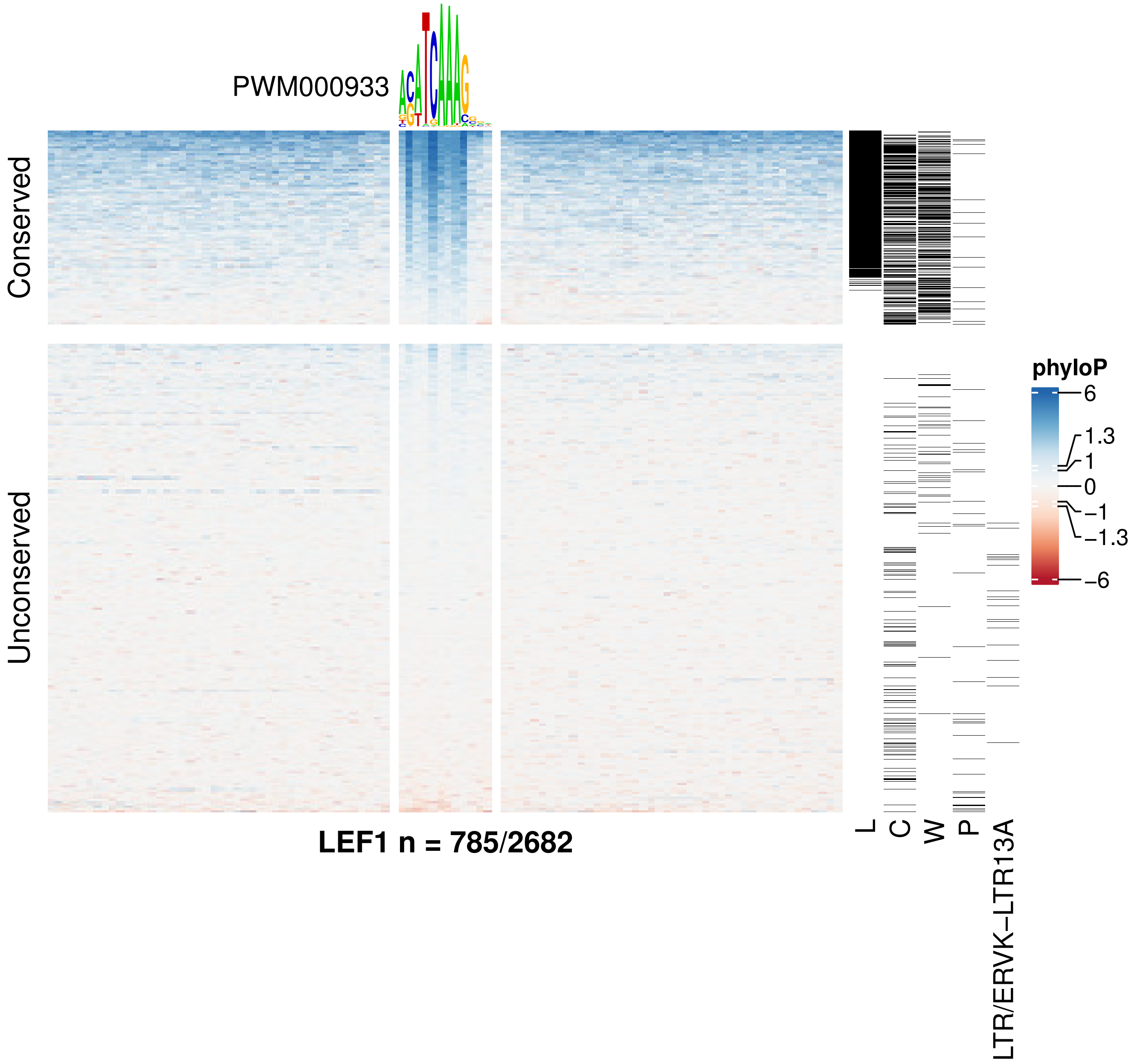

PWM000933
Conserved
Unconserved
phyloP
6
1.3
1
0
−1
−1.3
−6
L
C
W
P
LTR/ERVK−LTR13A
LEF1 n = 785/2682

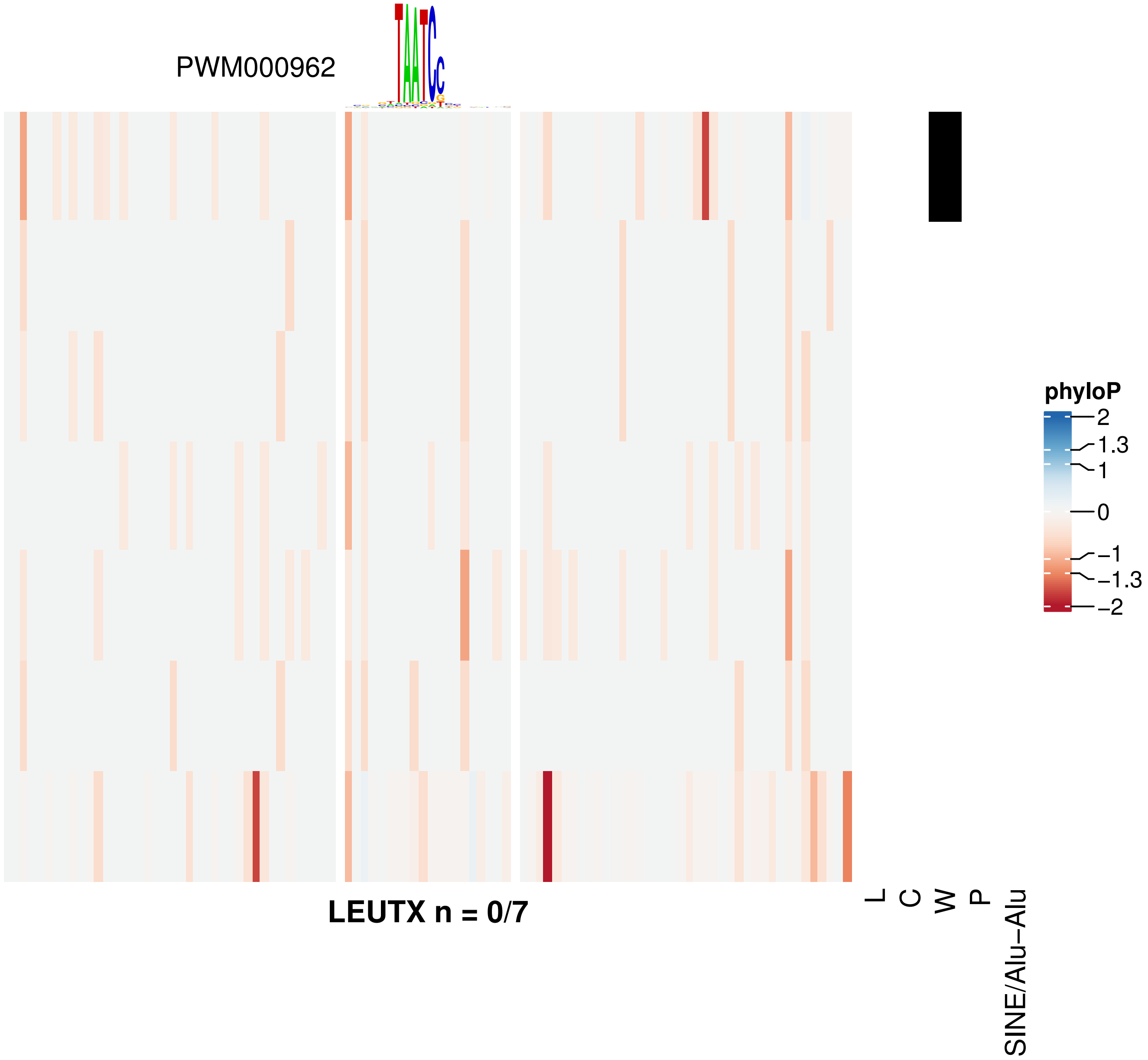

PWM000962
LEUTX n = 0/7
phyloP
2
1.3
1
0
−1
−1.3
−2
L
C
W
P
SINE/Alu−Alu

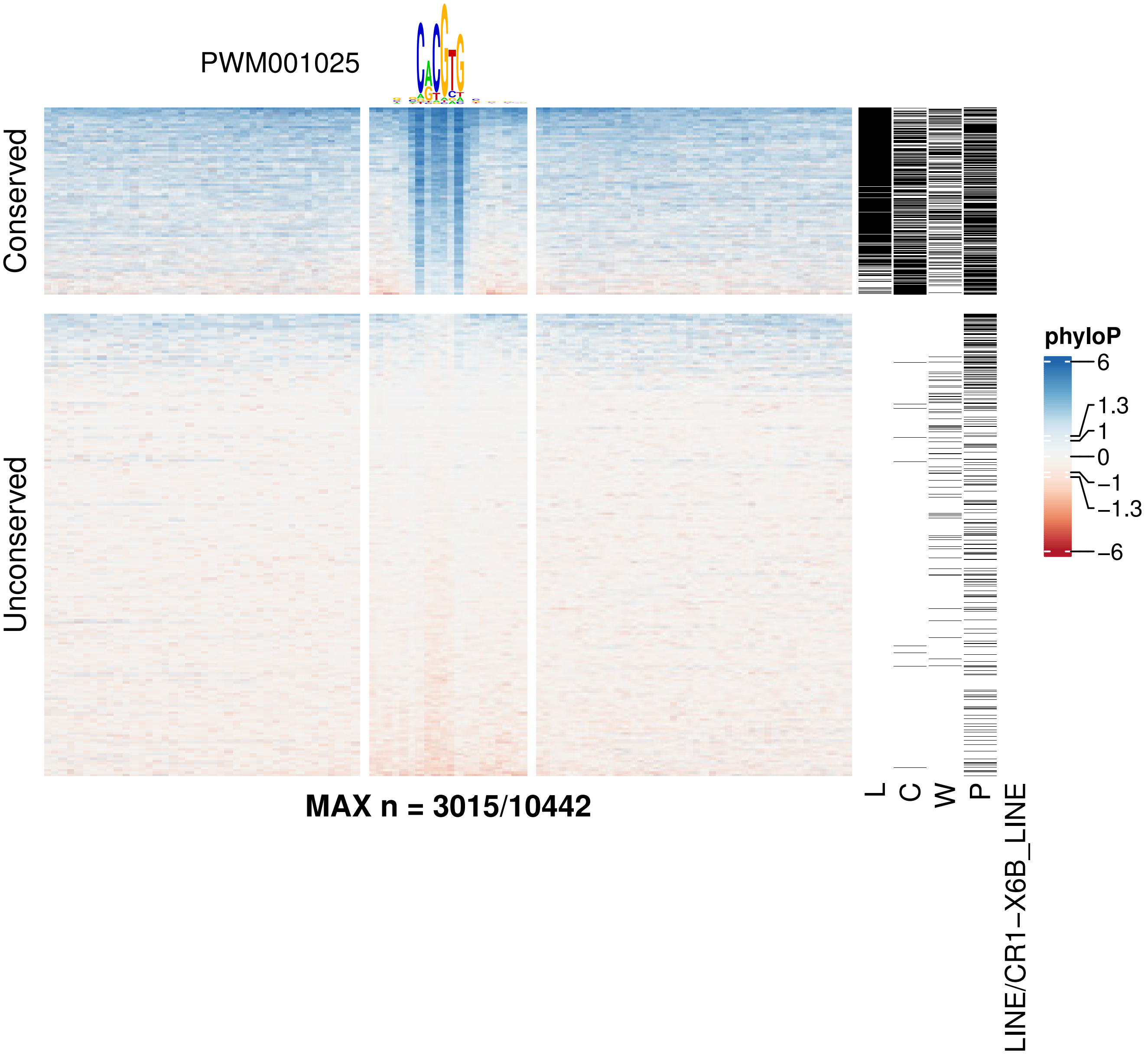

PWM001025
Conserved
Unconserved
MAX n = 3015/10442
L
C
W
P
LINE/CR1−X6B_LINE
phyloP
6
1.3
1
0
−1
−1.3
−6

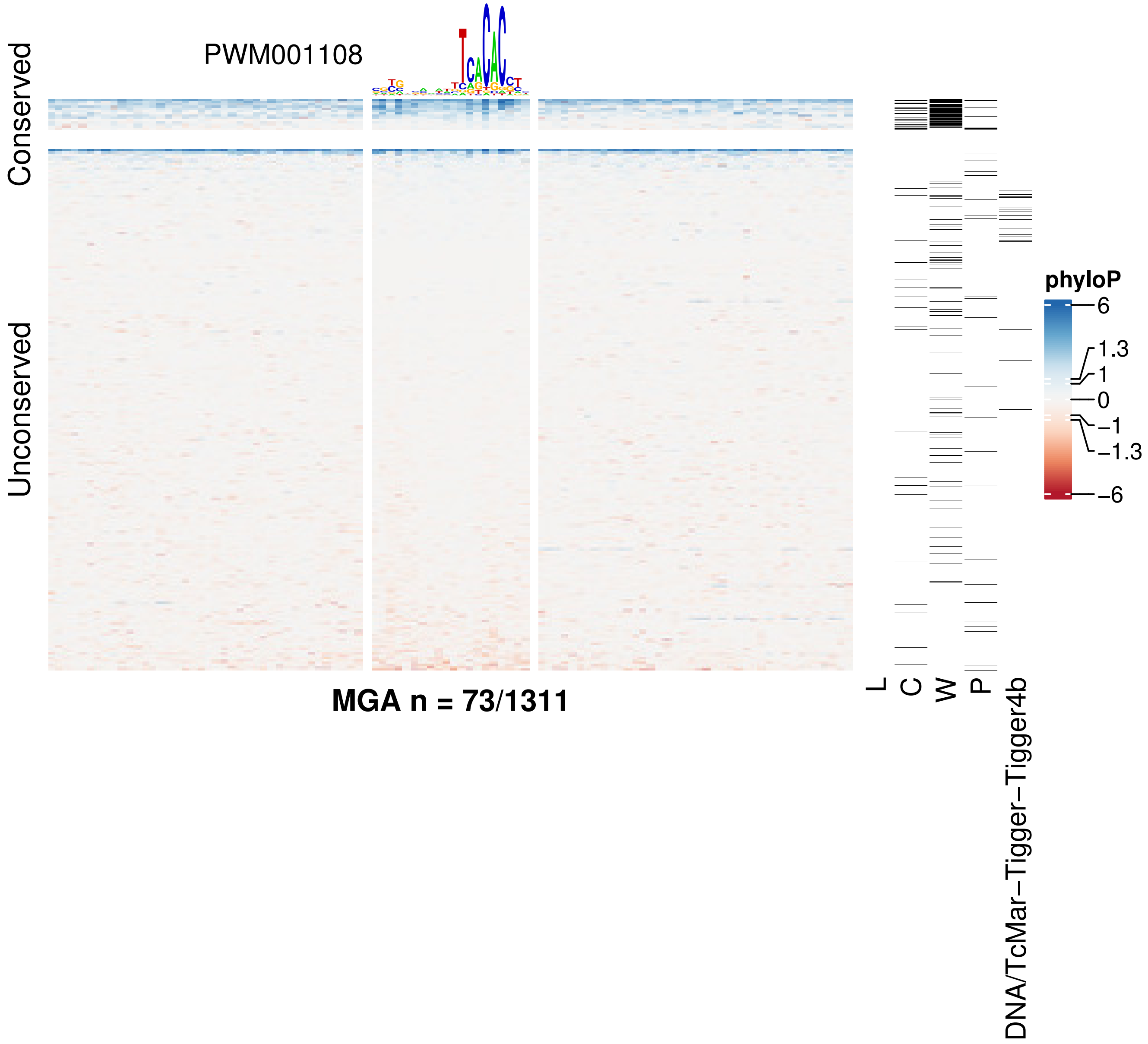

PWM001108
Conserved
Unconserved
MGA n = 73/1311
L
C
W
P
DNA/TcMar−Tigger−Tigger4b
phyloP
6
1.3
1
0
−1
−1.3
−6

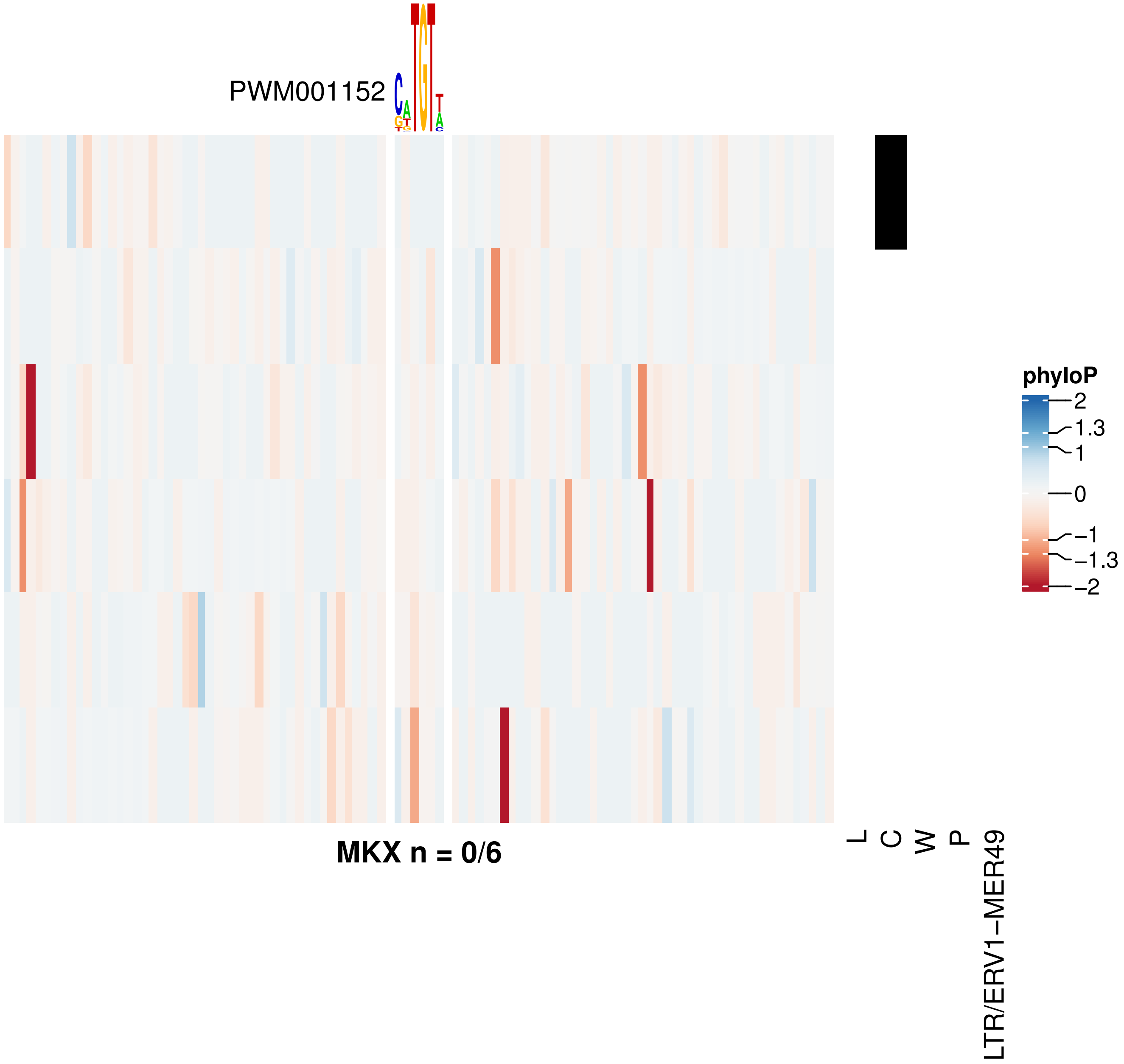
PWM001152
phyloP
2
1.3
1
0
−1
−1.3
−2
L
C
W
P
LTR/ERV1−MER49
MKX n = 0/6

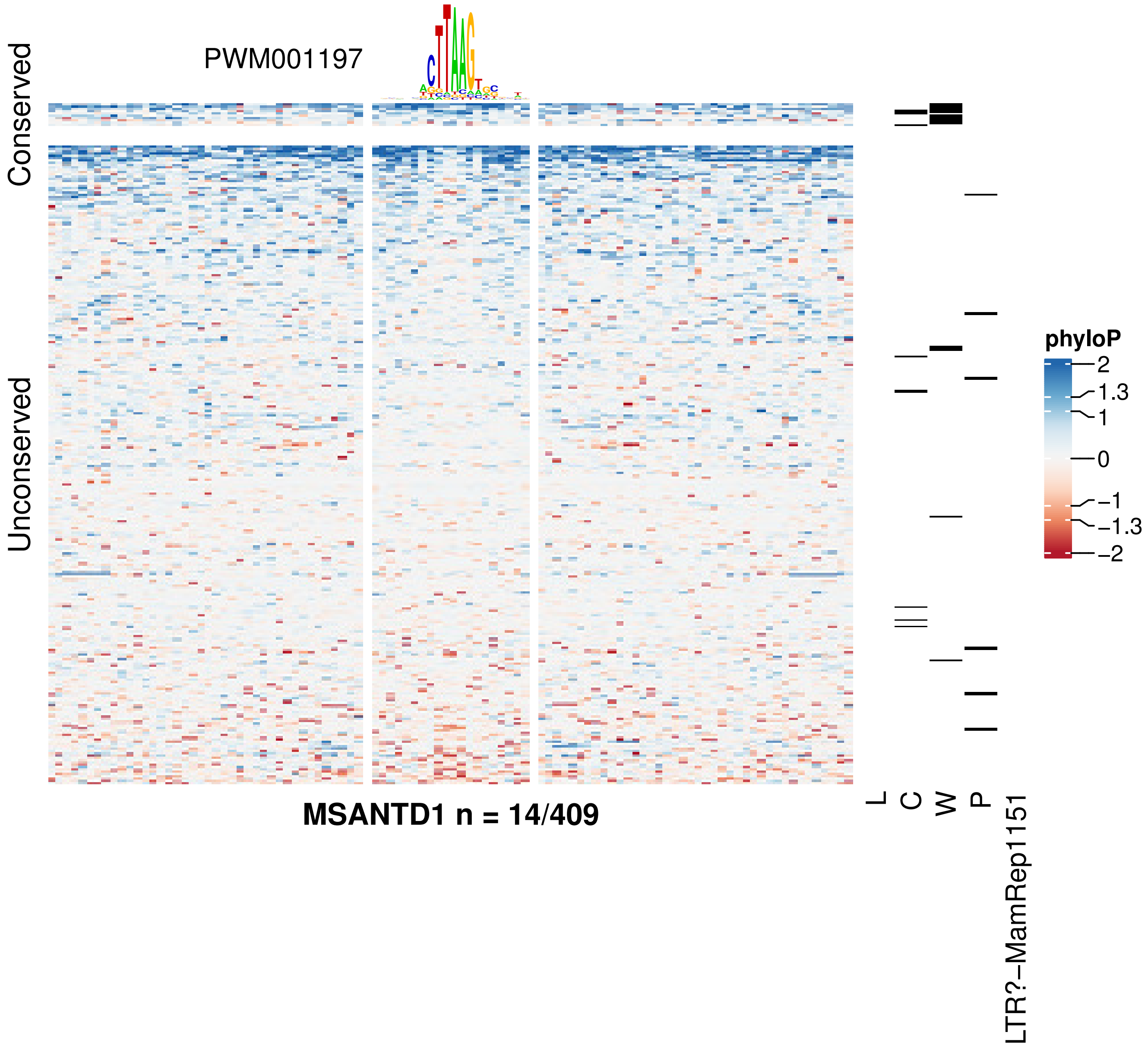

PWM001197
Conserved
Unconserved
phyloP
2
1.3
1
0
−1
−1.3
−2
L
C
W
P
LTR?−MamRep1151
MSANTD1 n = 14/409

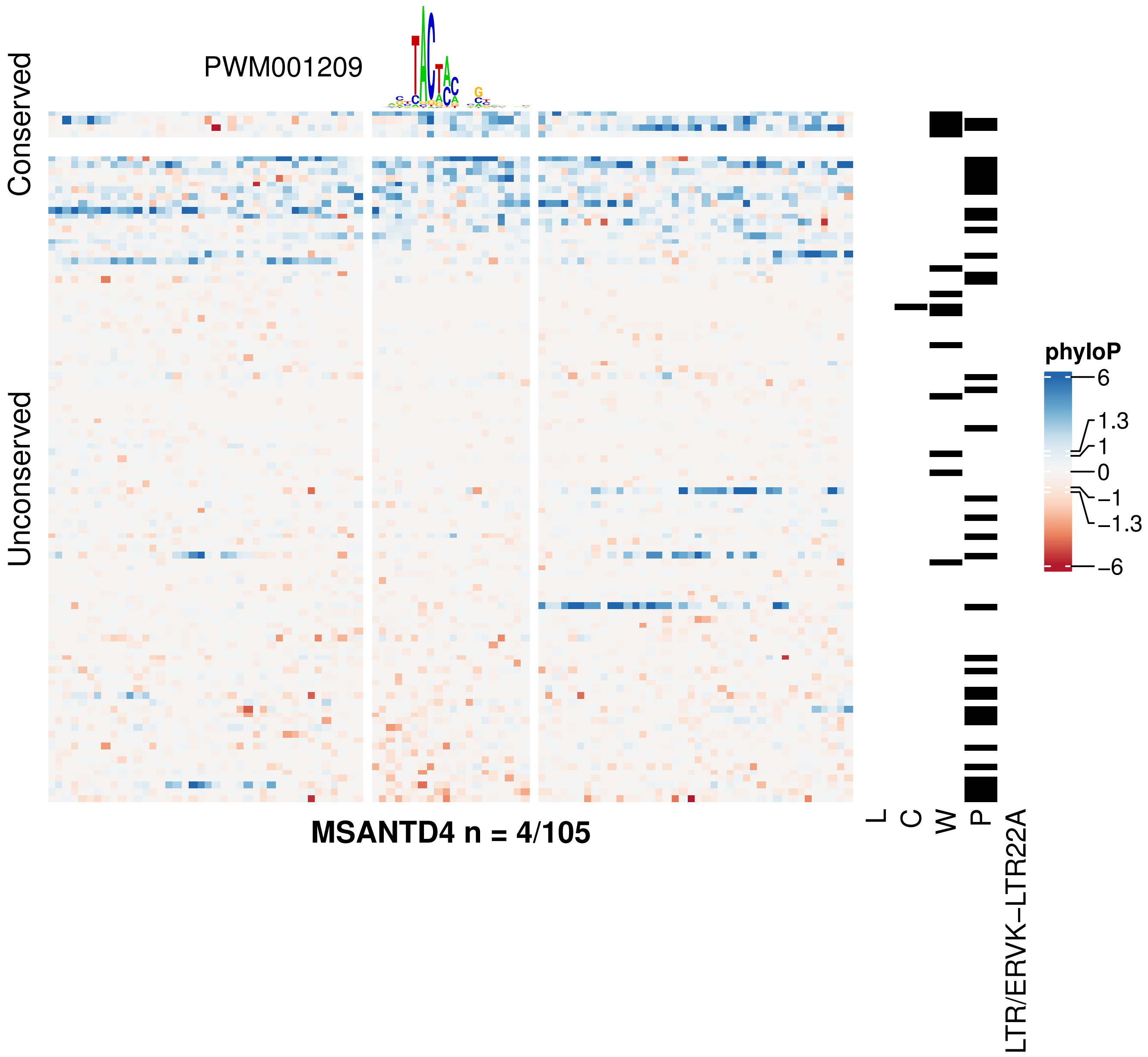

PWM001209
Conserved
Unconserved
MSANTD4 n = 4/105
L
C
W
P
LTR/ERVK-LTR22A
phyloP
6
1.3
1
0
−1
−1.3
−6

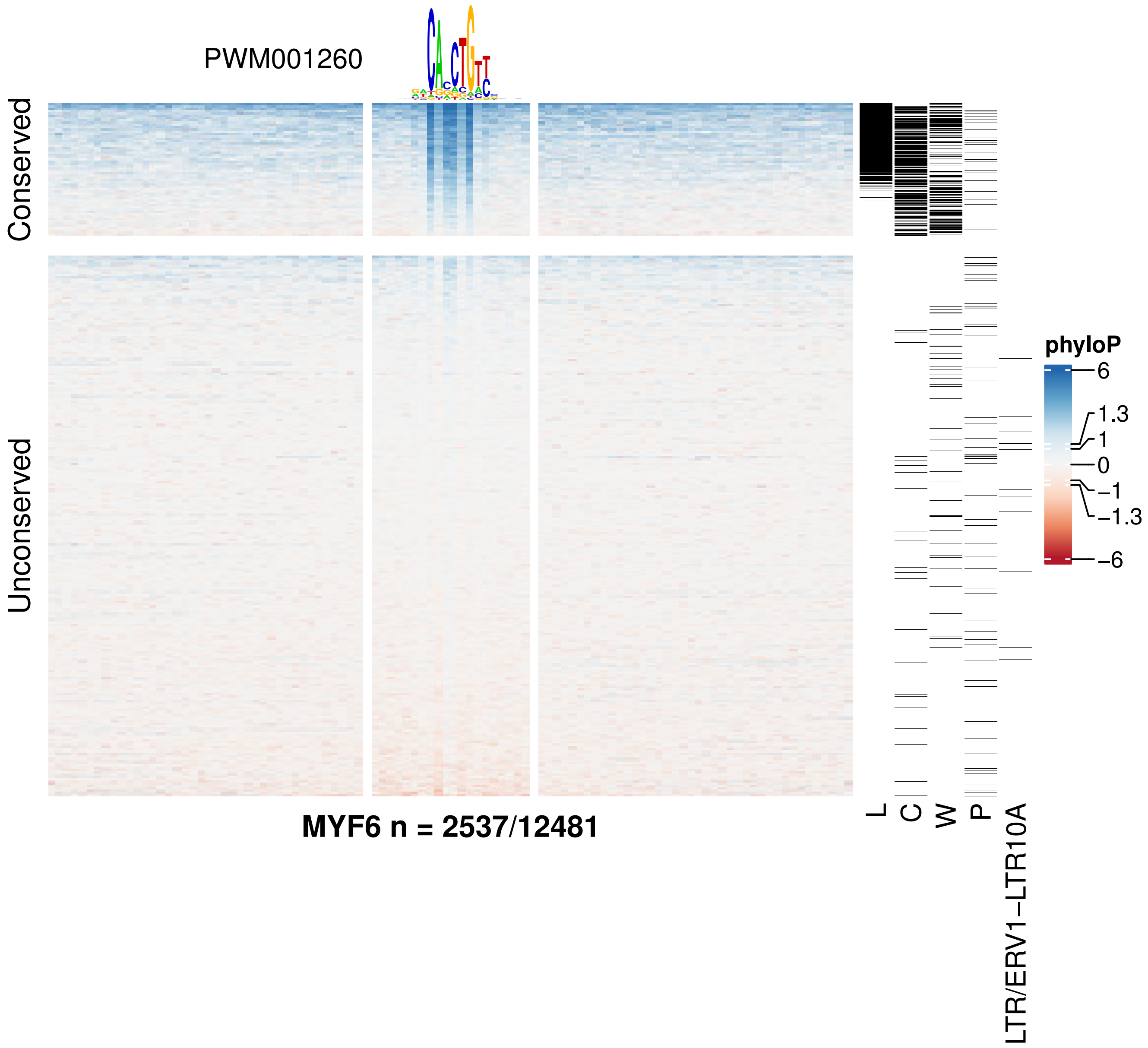
PWM001260
Conserved
Unconserved
MYF6 n = 2537/12481
L
C
W
P
LTR/ERV1−LTR10A
phyloP
6
1.3
1
0
−1
−1.3
−6

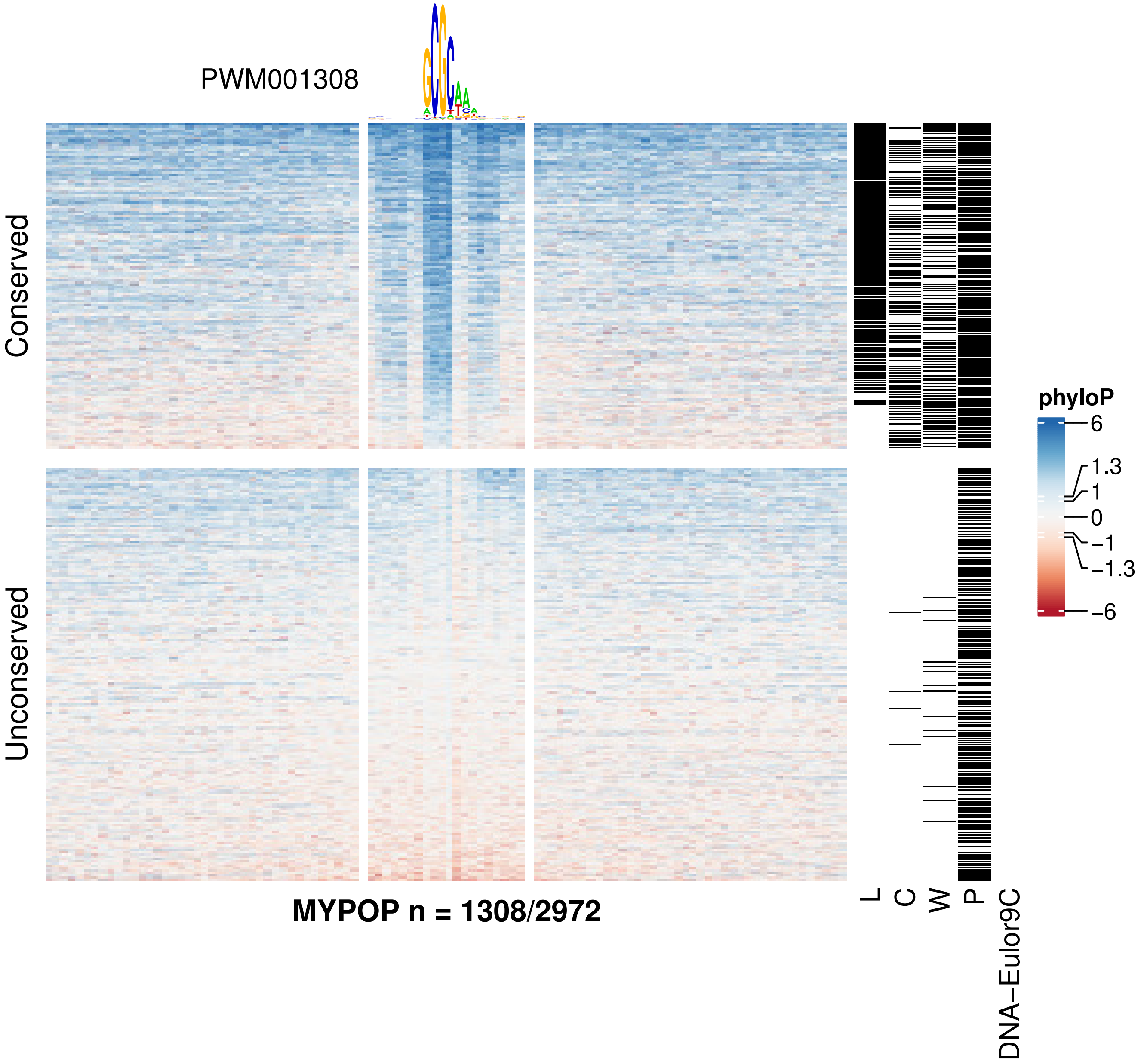
PWM001308
Conserved
Unconserved
phyloP
6
1.3
1
0
−1
−1.3
−6
MYPOP n = 1308/2972
L
C
W
P
DNA−Eulor9C

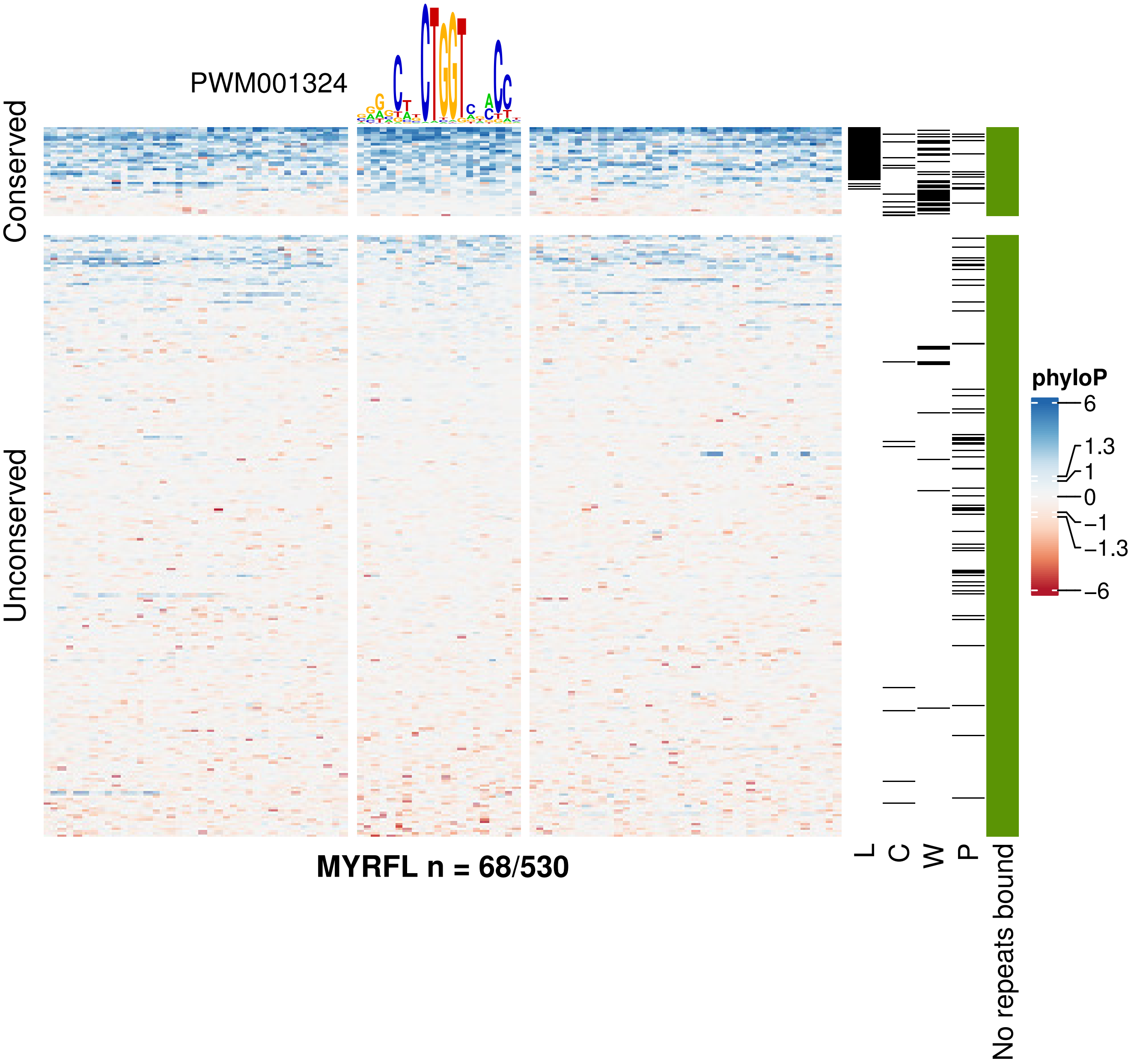
PWM001324
Conserved
Unconserved
MYRFL n = 68/530
L
C
W
P
No repeats bound
phyloP
6
1.3
1
0
−1
−1.3
−6

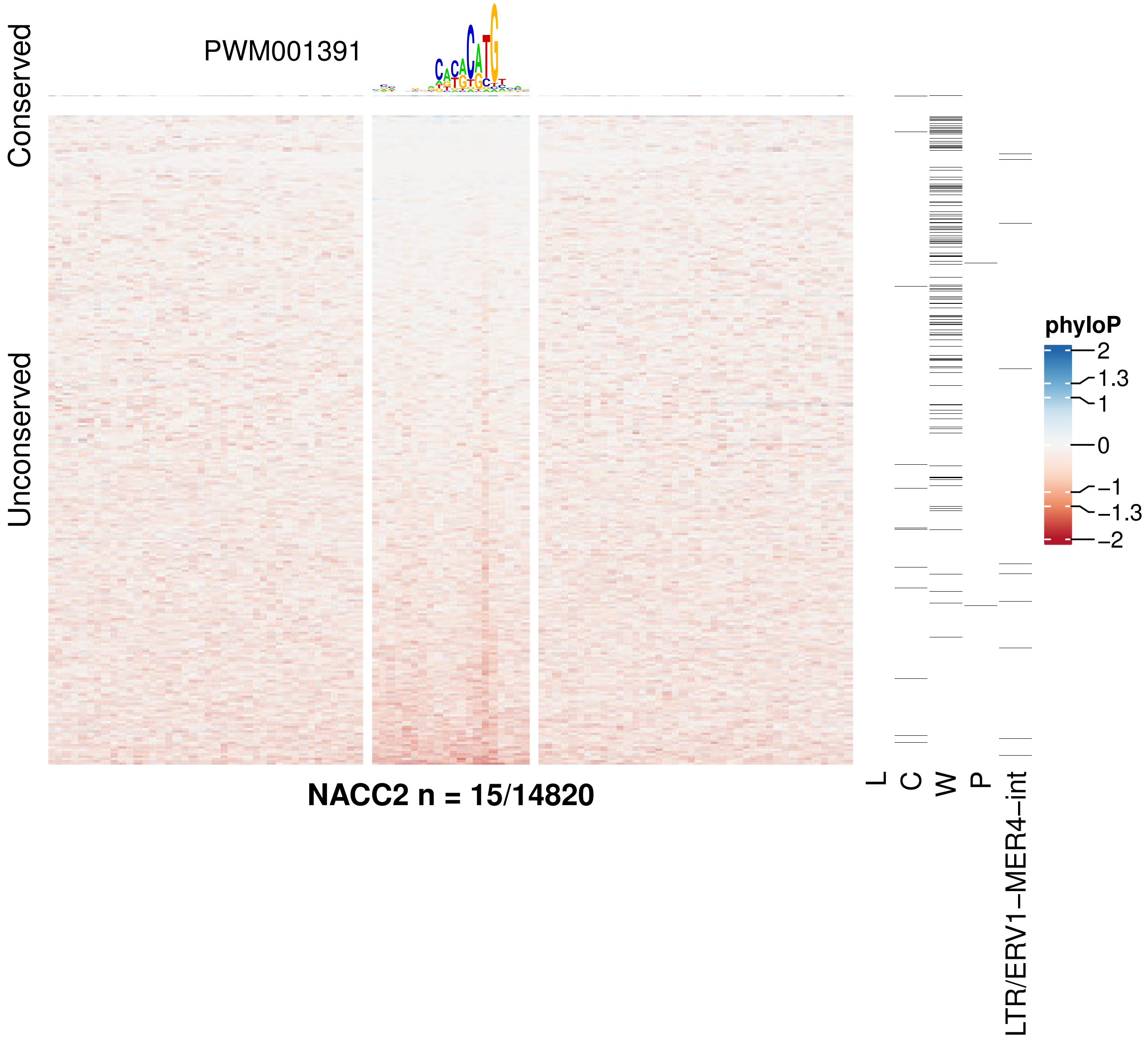

Conserved
Unconserved
PWM001391
NACC2 n = 15/14820
L
C
W
P
LTR/ERV1−MER4−int
phyloP
2
1.3
1
0
−1
−1.3
−2

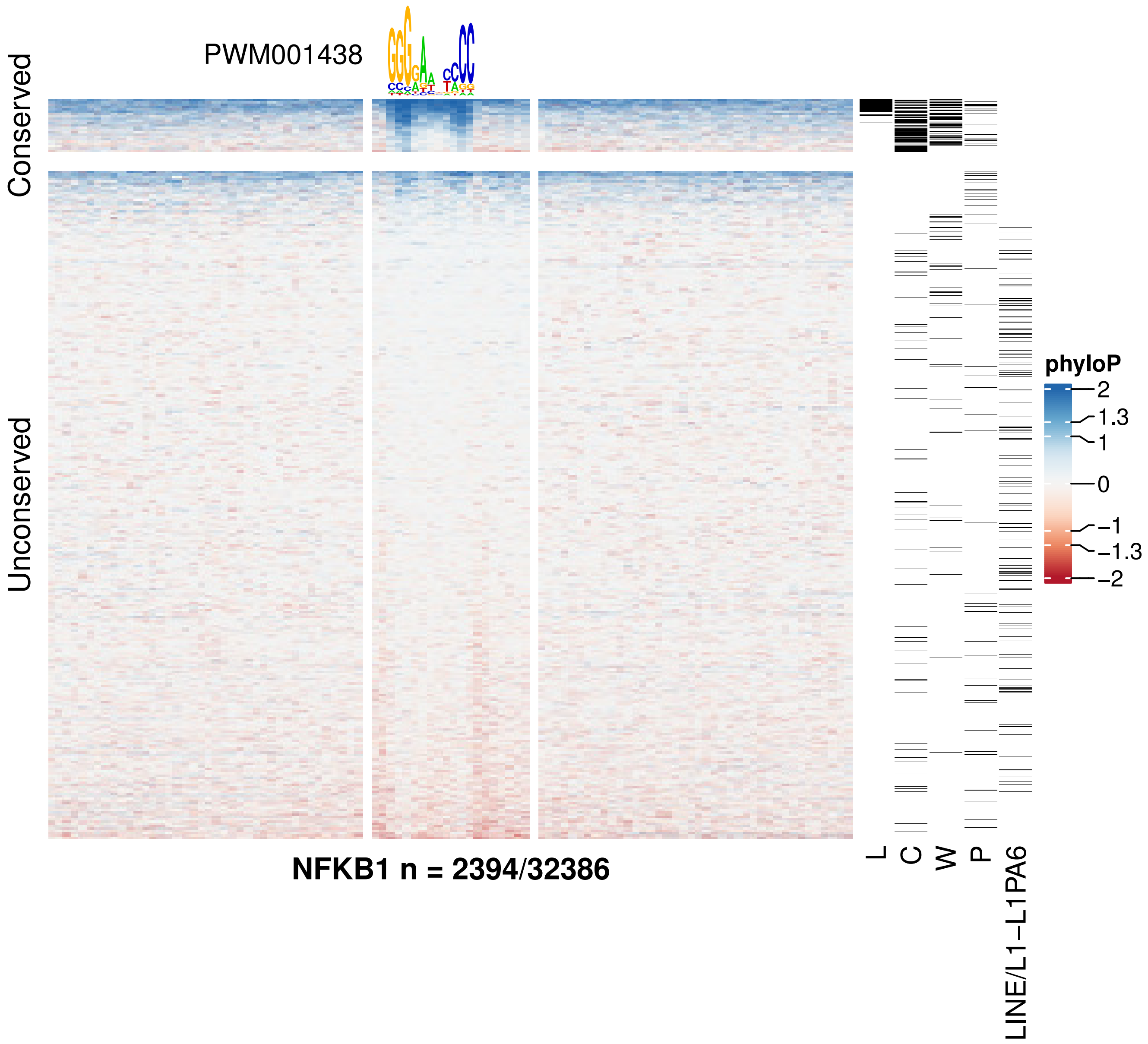

PWM001438
Conserved
Unconserved
NFKB1 n = 2394/32386
L
C
W
P
LINE/L1−L1PA6
phyloP
2
1.3
1
0
−1
−1.3
−2

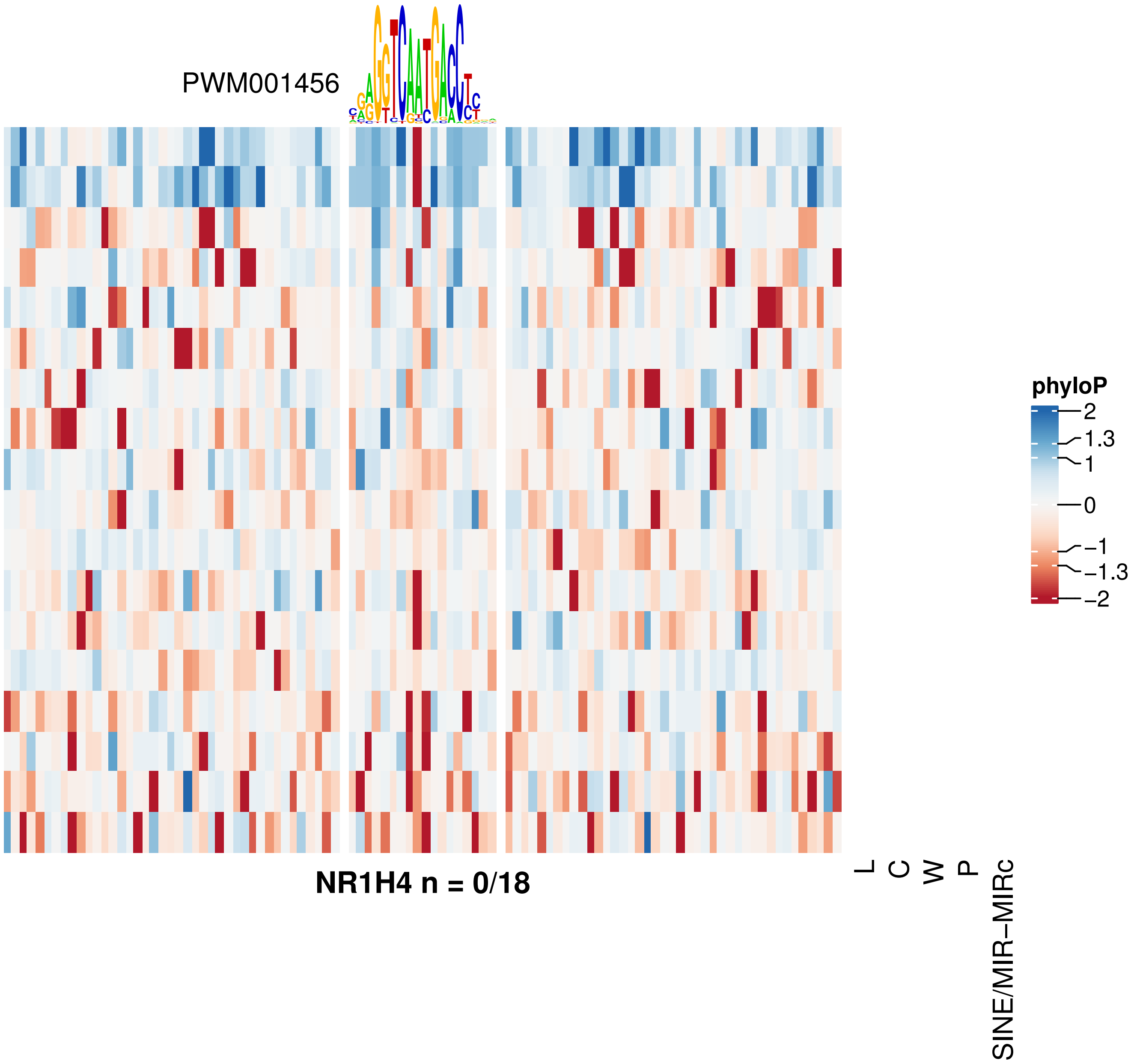


**NR1H4 n = 0/18**

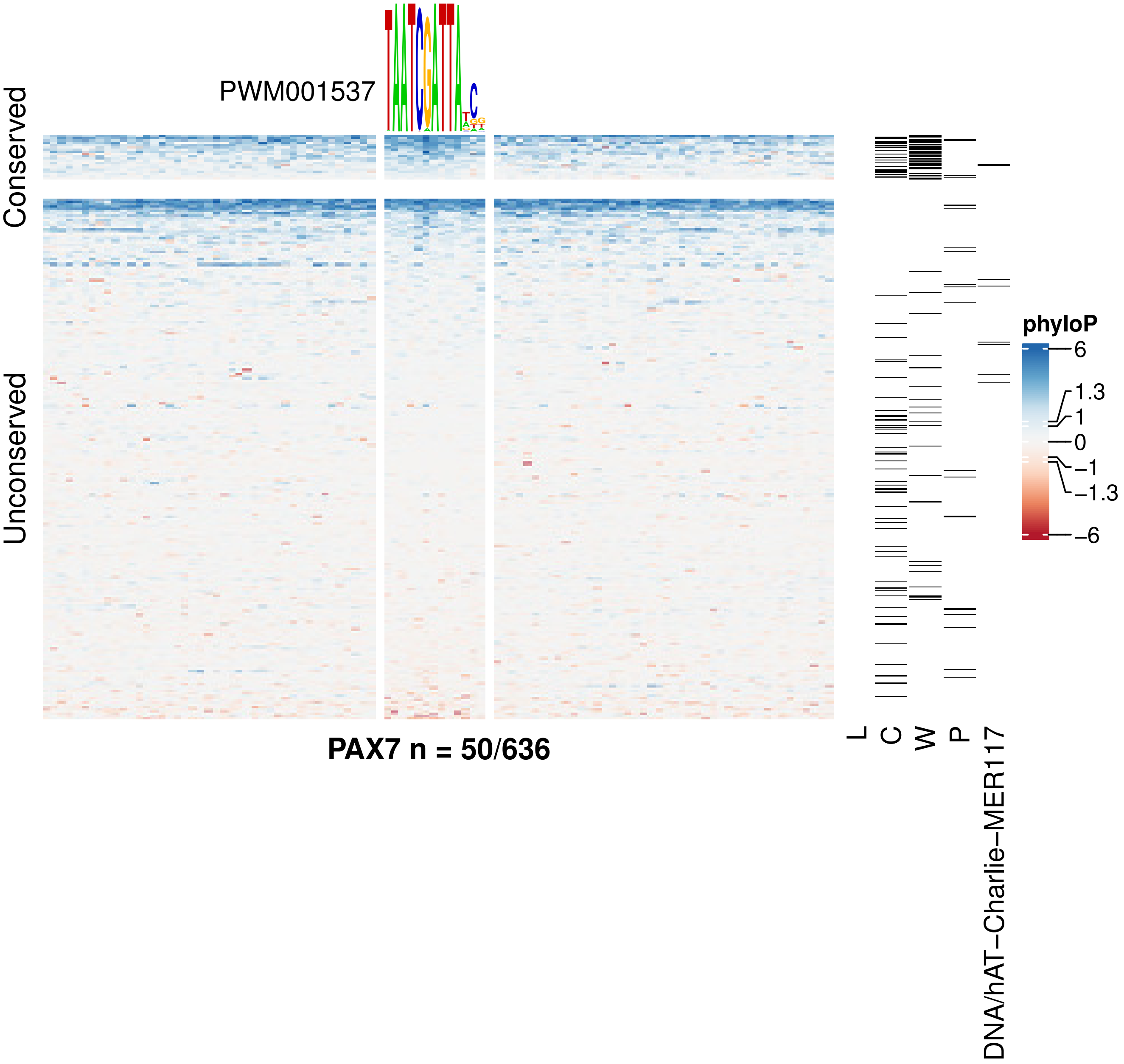
PWM001537
Conserved
Unconserved
phyloP
6
1.3
1
0
−1
−1.3
−6
PAX7 n = 50/636
L
C
W
P
DNA/hAT−Charlie−MER117

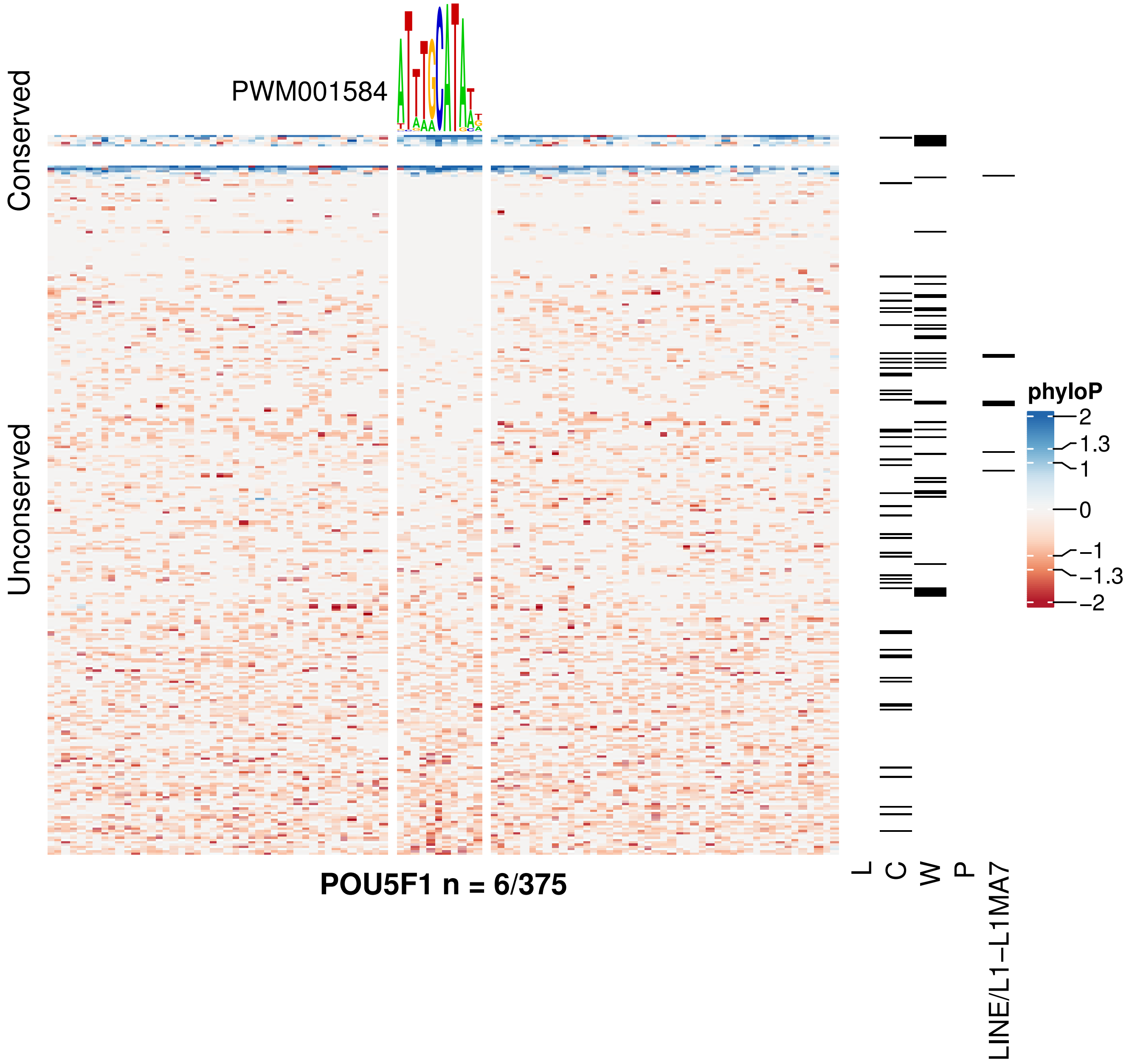
PWM001584
Conserved
Unconserved
POU5F1 n = 6/375
L
C
W
P
LINE/L1−L1MA7
phyloP
2
1.3
1
0
−1
−1.3
−2

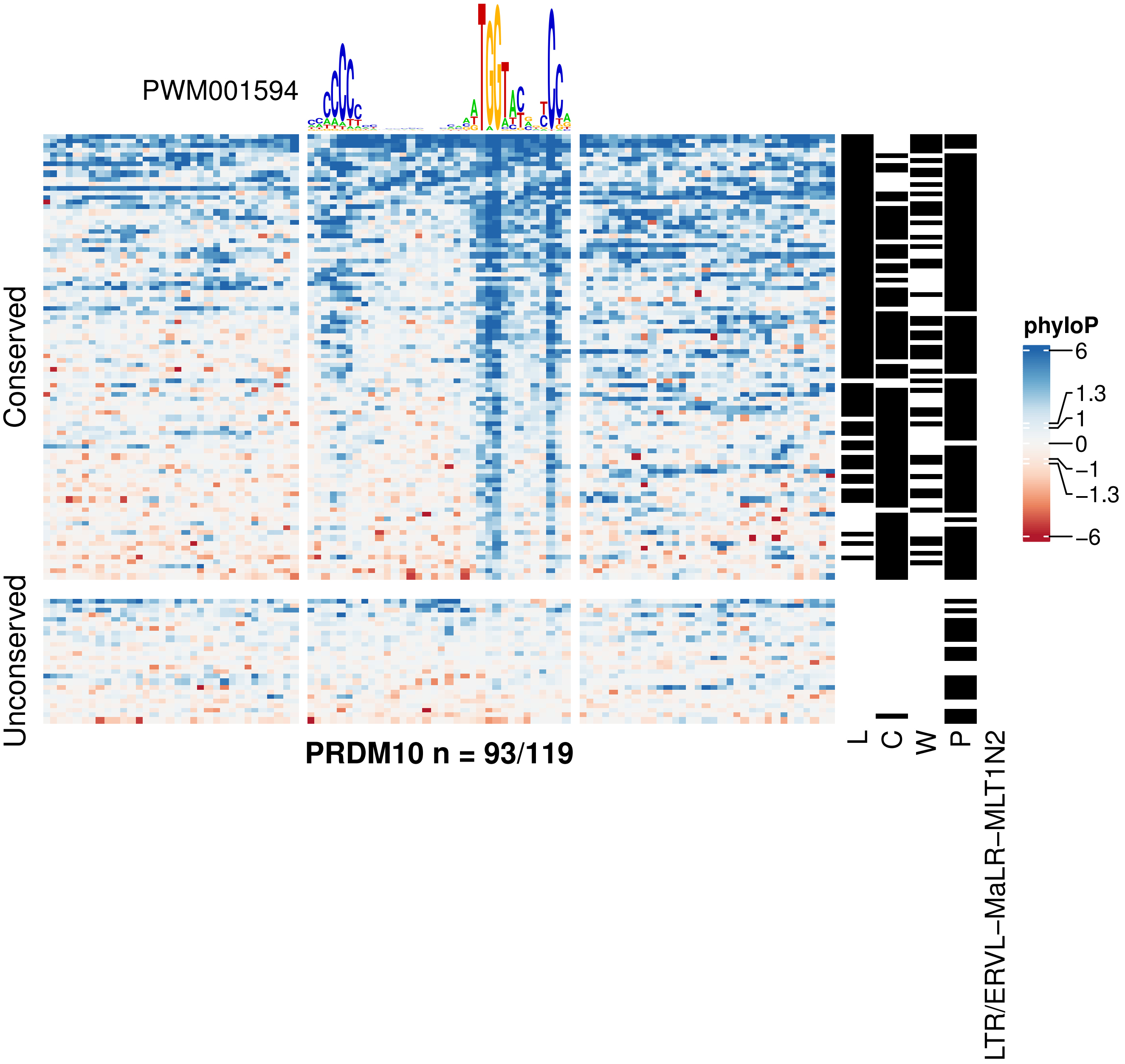

PWM001594
Conserved
Unconserved
PRDM10 n = 93/119
L
C
W
P
LTR/ERVL−MaLR−MLT1N2
phyloP
6
1.3
1
0
−1
−1.3
−6

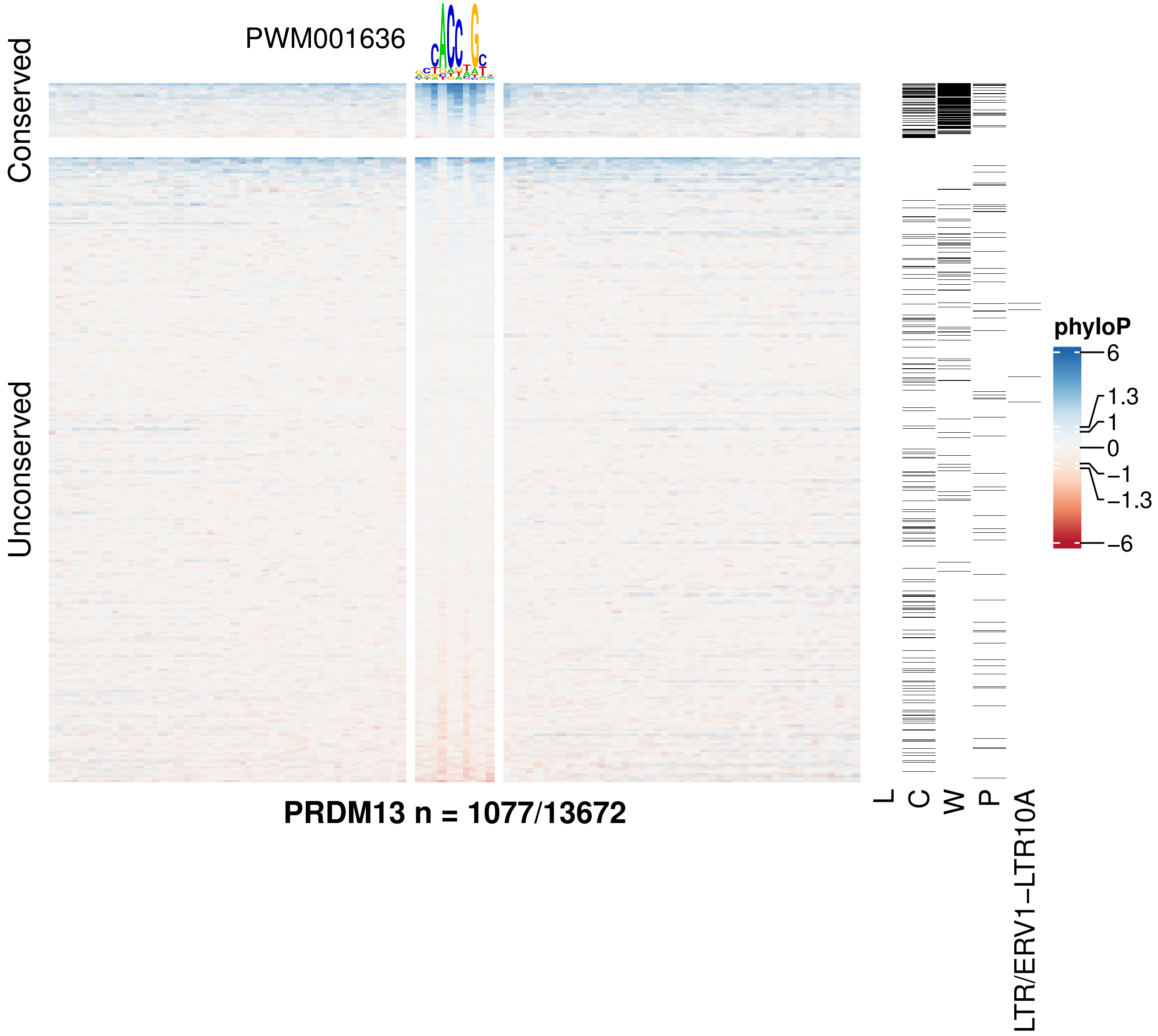

PWM001636
Conserved
Unconserved
PRDM13 n = 1077/13672
L
C
W
P
LTR/ERV1−LTR10A
phyloP
6
1.3
1
0
−1
−1.3
−6

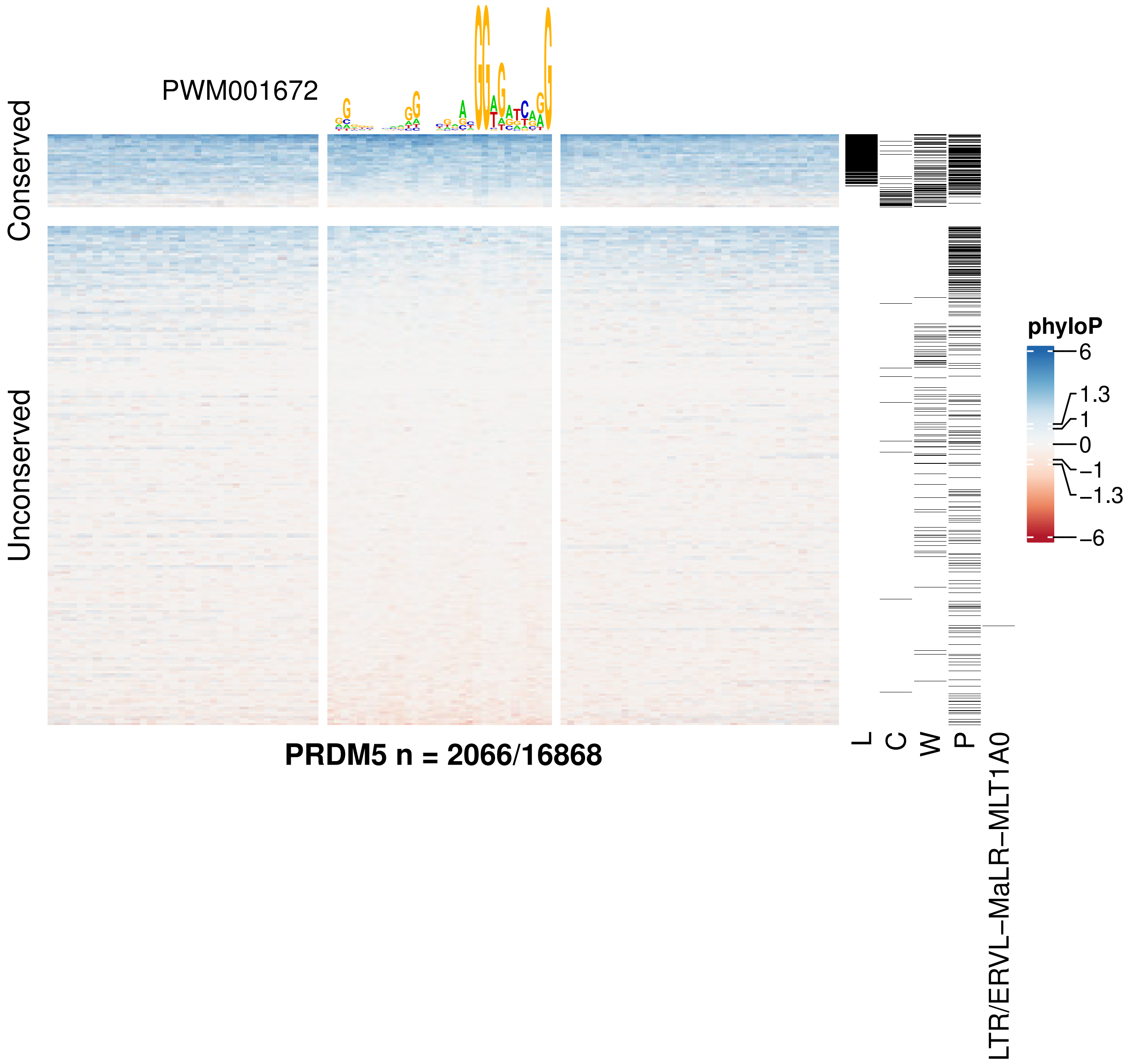
PWM001672
Conserved
Unconserved
PRDM5 n = 2066/16868
L
C
W
P
LTR/ERVL−MaLR−MLT1A0
phyloP
6
1.3
1
0
−1
−1.3
−6

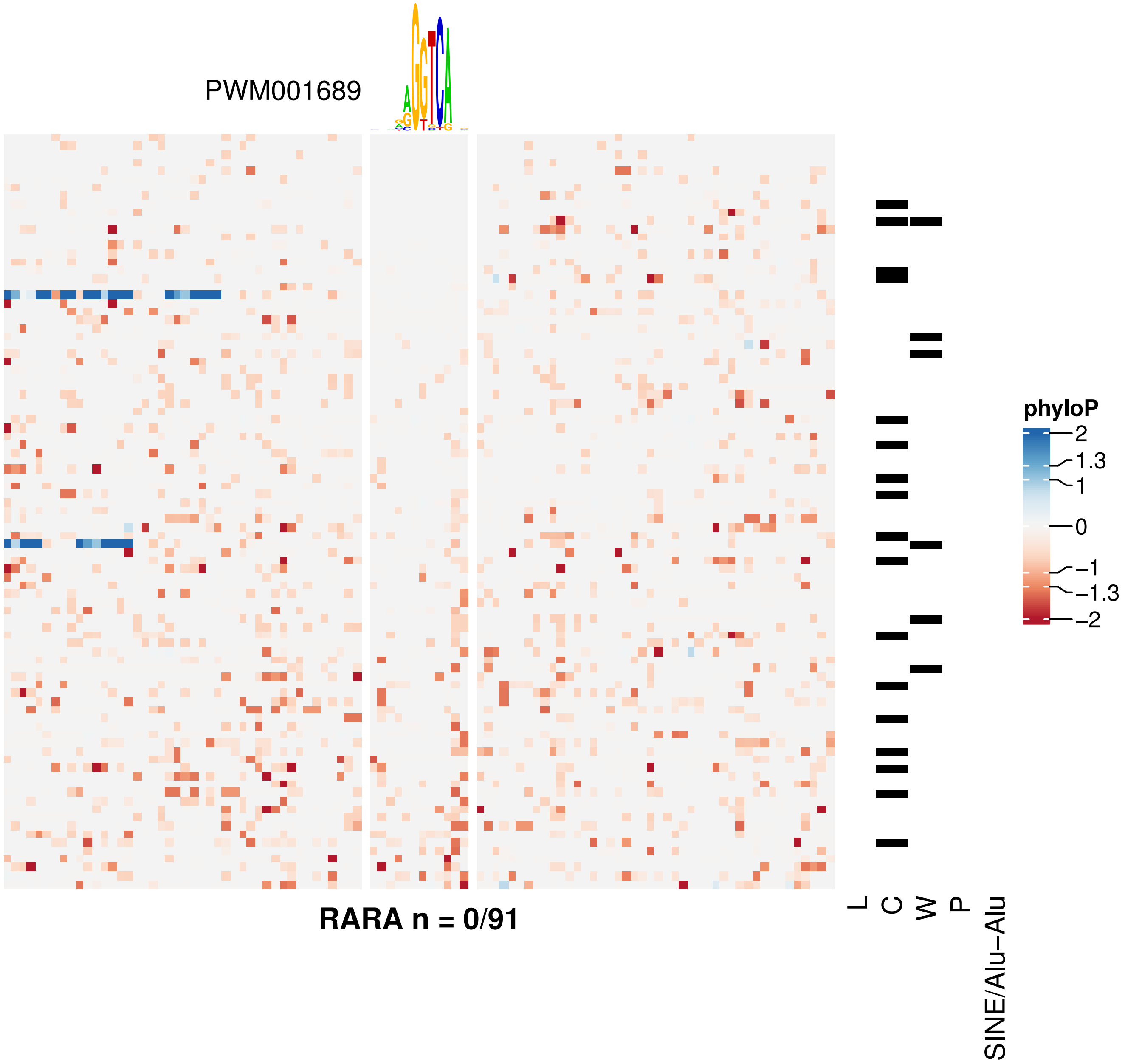

PWM001689
RARA n = 0/91
L
C
W
P
SINE/Alu−Alu
phyloP
2
1.3
1
0
−1
−1.3
−2

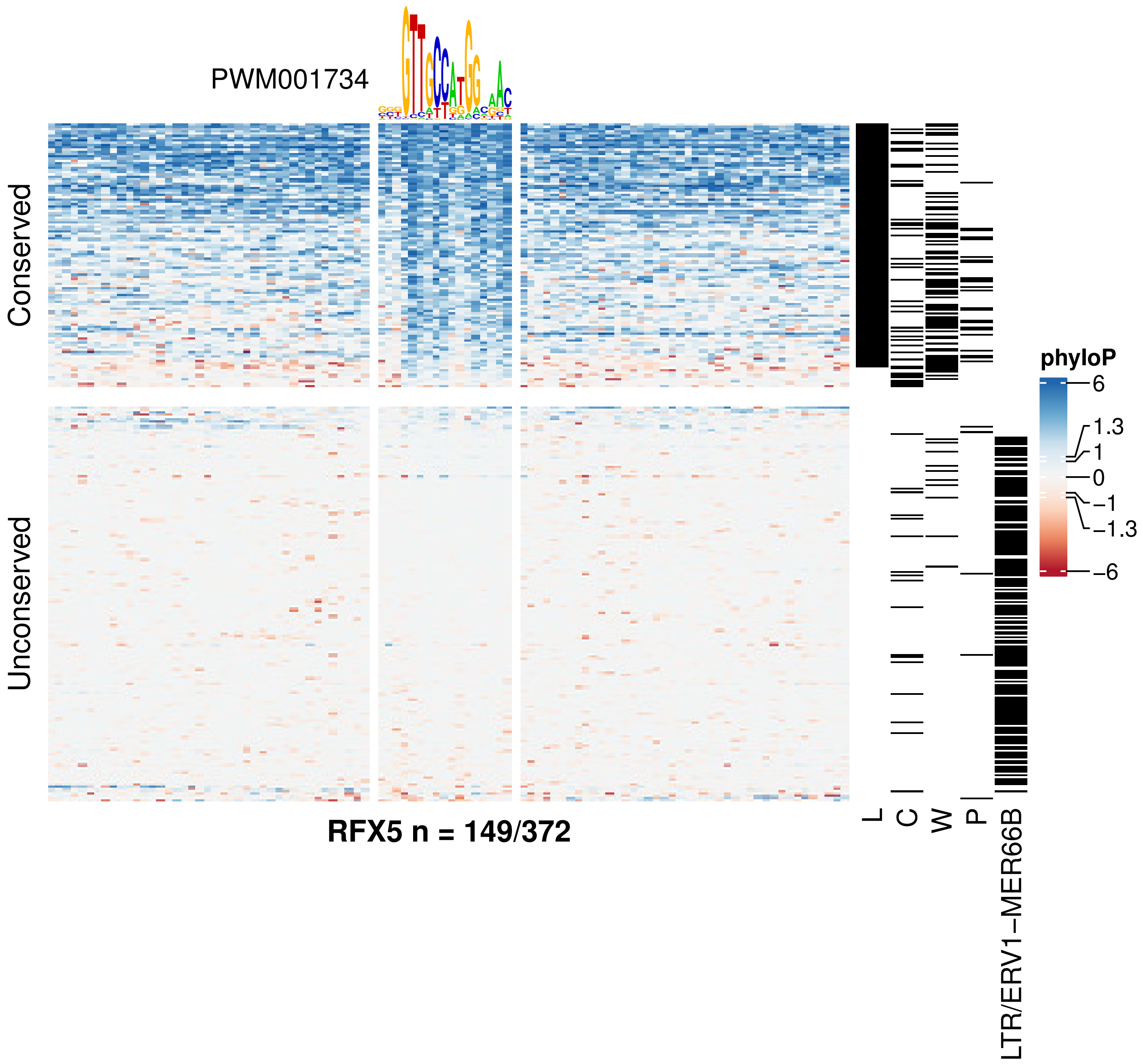

PWM001734
Conserved
Unconserved
RFX5 n = 149/372
L
C
W
P
LTR/ERV1−MER66B
phyloP
6
1.3
1
0
−1
−1.3
−6

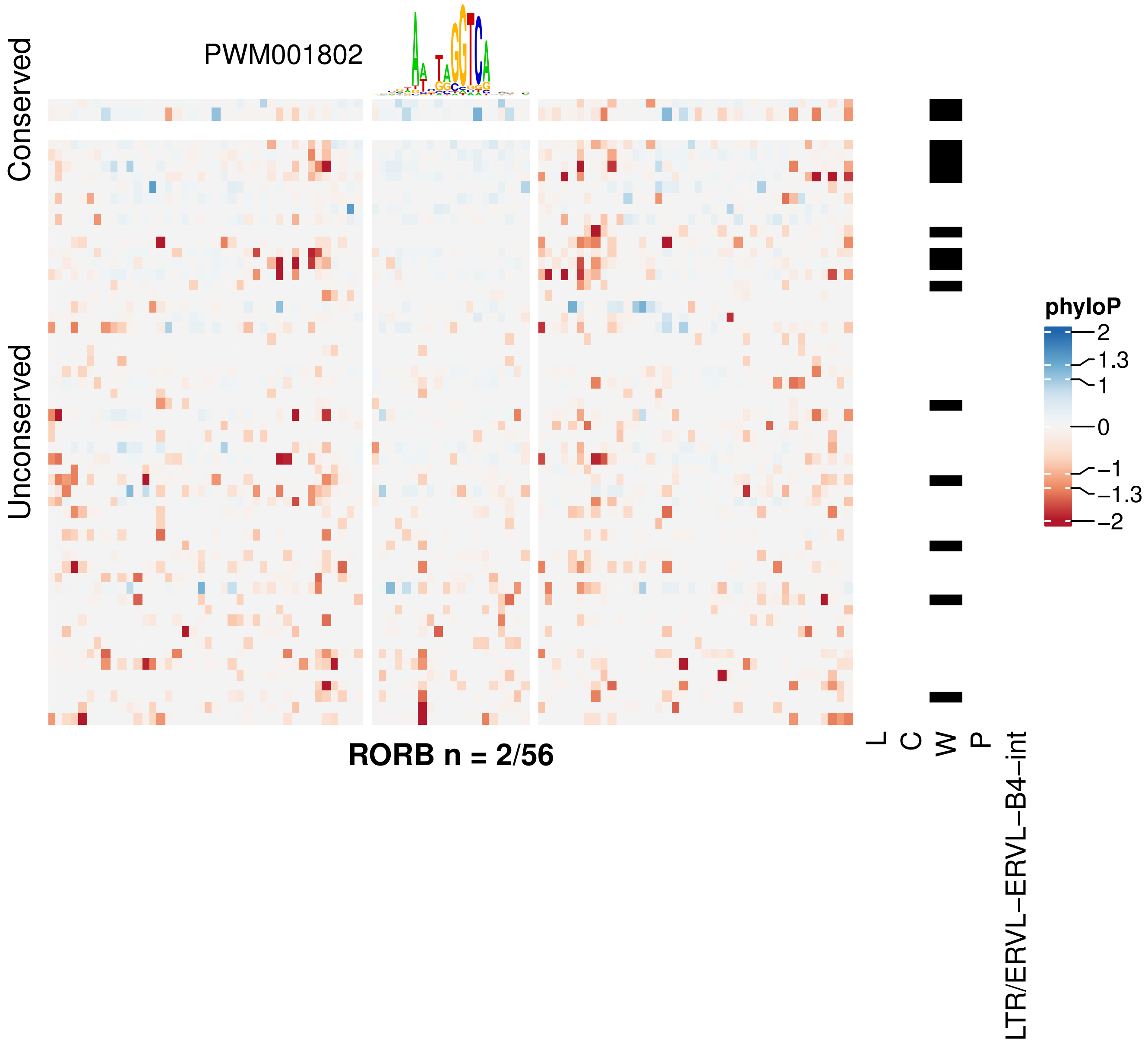

PWM001802
Conserved
Unconserved
RORB n = 2/56
L
C
W
P
LTR/ERVL−ERVL−B4−int
phyloP
2
1.3
1
0
−1
−1.3
−2

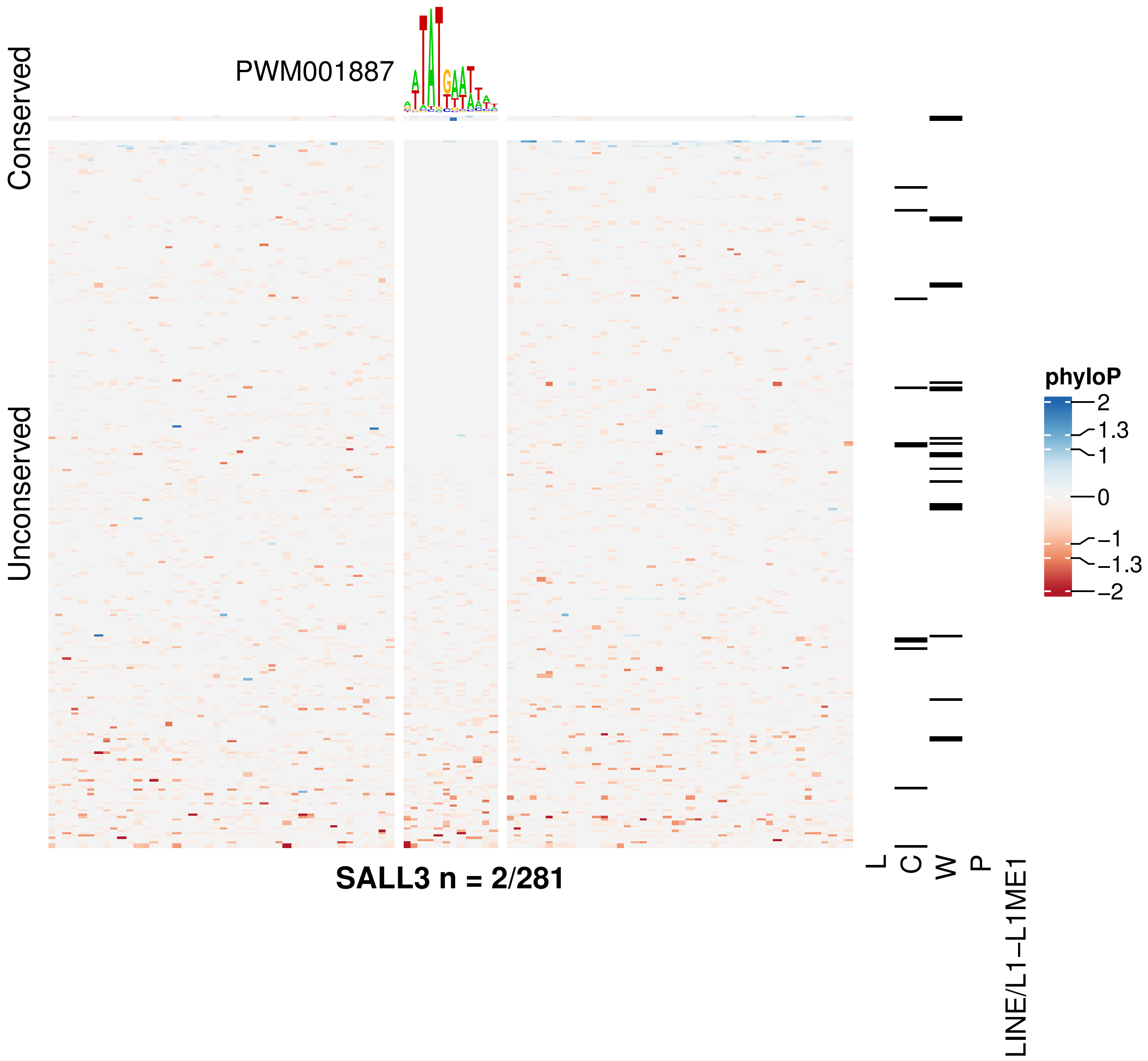

PWM001887
Conserved
Unconserved
SALL3 n = 2/281
L
C
W
P
LINE/L1−L1ME1
phyloP
2
1.3
1
0
−1
−1.3
−2

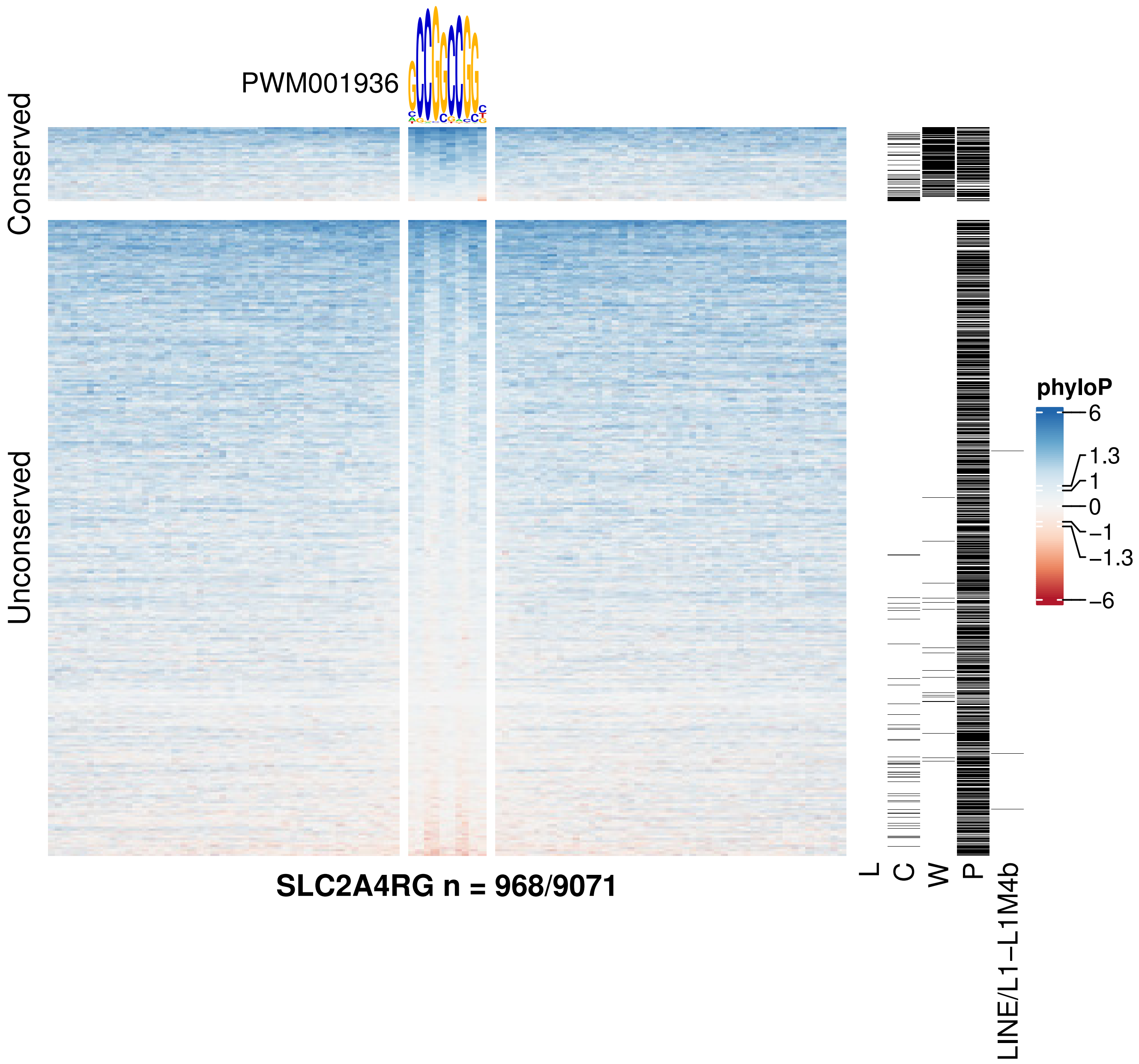
PWM001936
Conserved
Unconserved
SLC2A4RG n = 968/9071
L
C
W
P
LINE/L1−L1M4b
phyloP
6
1.3
1
0
−1
−1.3
−6

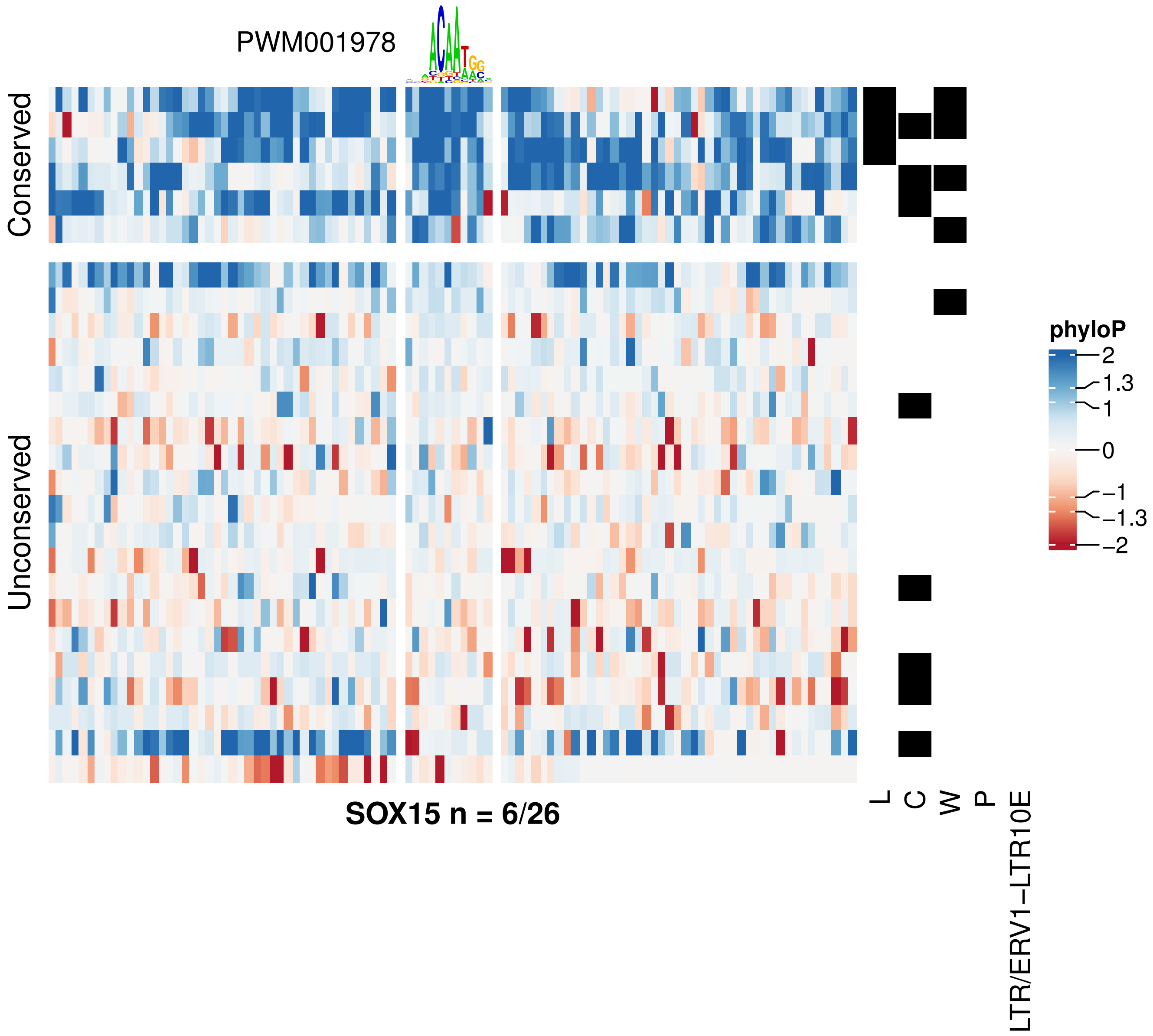

PWM001978
Conserved
Unconserved
SOX15 n = 6/26
L
C
W
P
LTR/ERV1−LTR10E
phyloP
2
1.3
1
0
−1
−1.3
−2

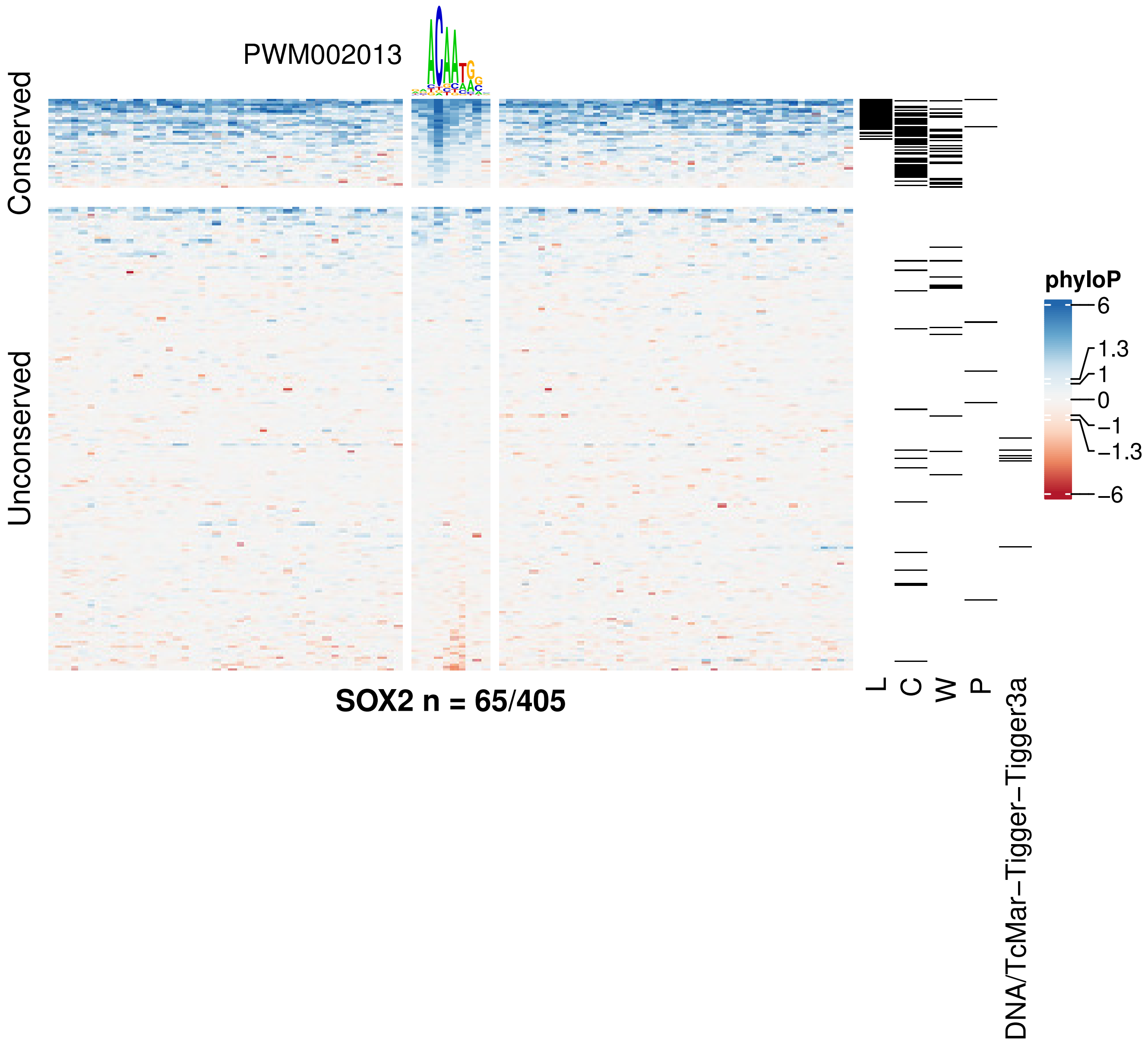
PWM002013
Conserved
Unconserved
SOX2 n = 65/405
L
C
W
P
DNA/TcMar−Tigger−Tigger3a
phyloP
6
1.3
1
0
−1
−1.3
−6

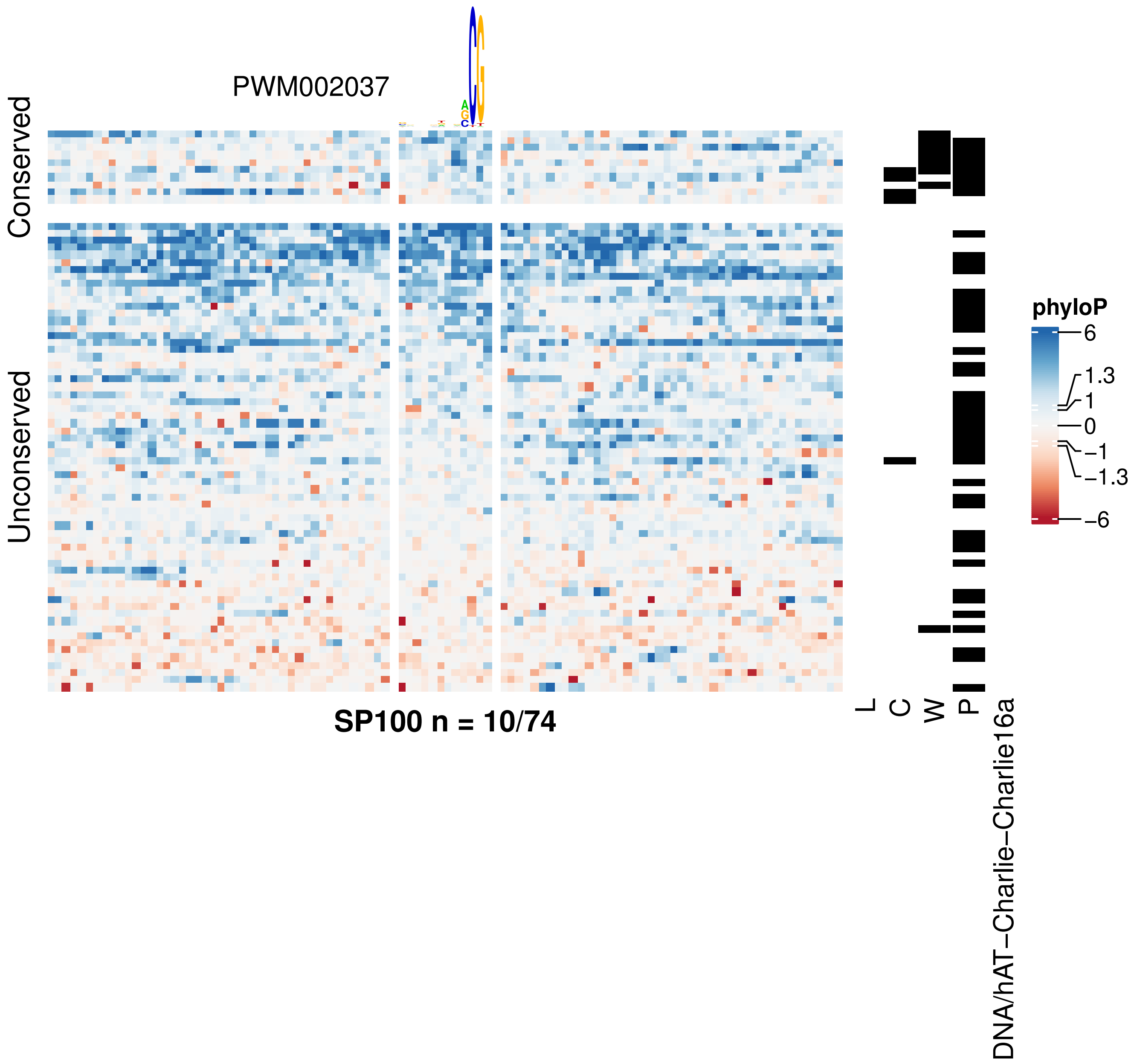
PWM002037
Conserved
Unconserved
SP100 n = 10/74
L
C
W
P
DNA/hAT−Charlie−Charlie16a
phyloP
6
1.3
1
0
−1
−1.3
−6

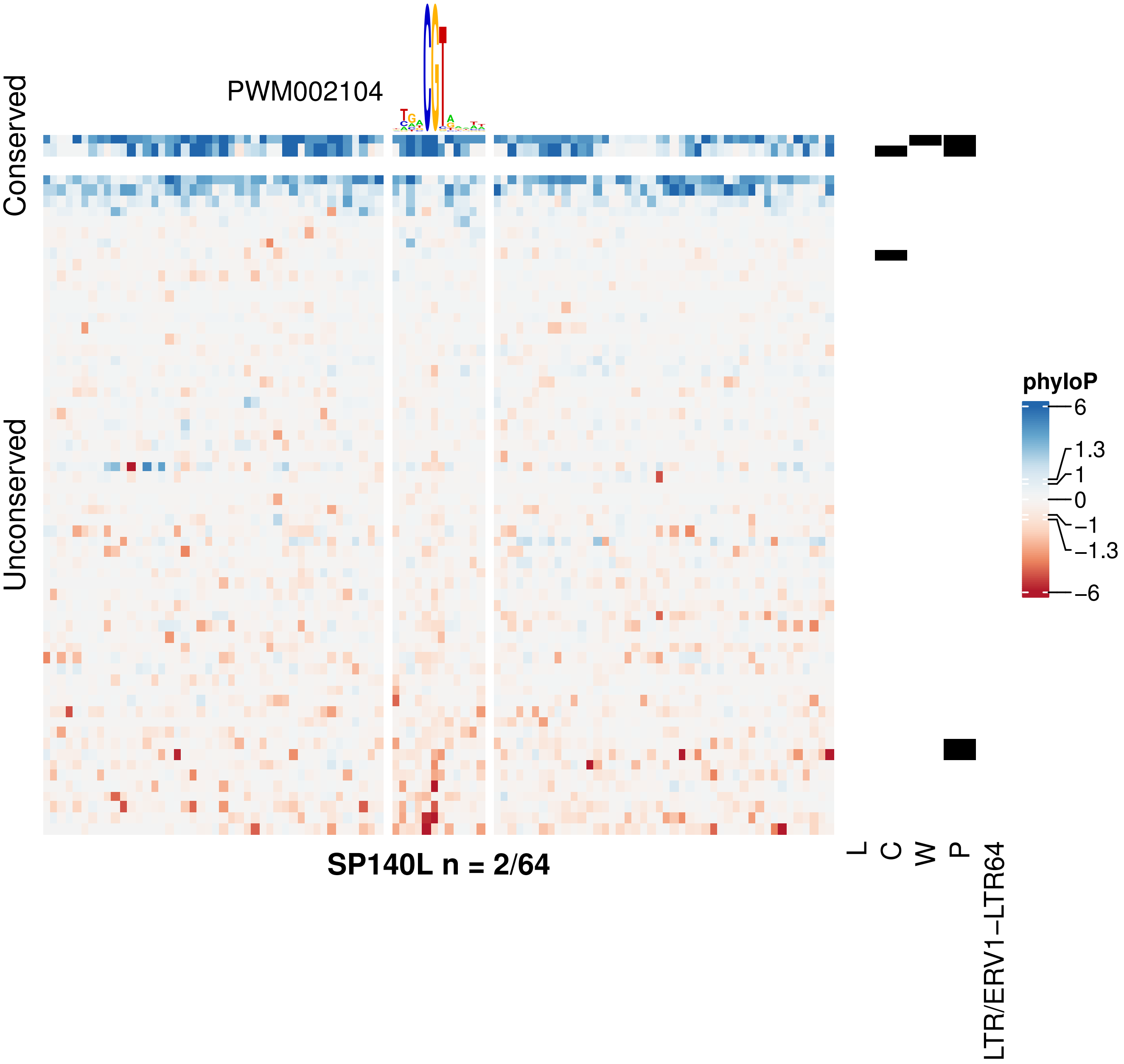

PWM002104
Conserved
Unconserved
SP140L n = 2/64
L
C
W
P
LTR/ERV1−LTR64
phyloP
6
1.3
1
0
−1
−1.3
−6

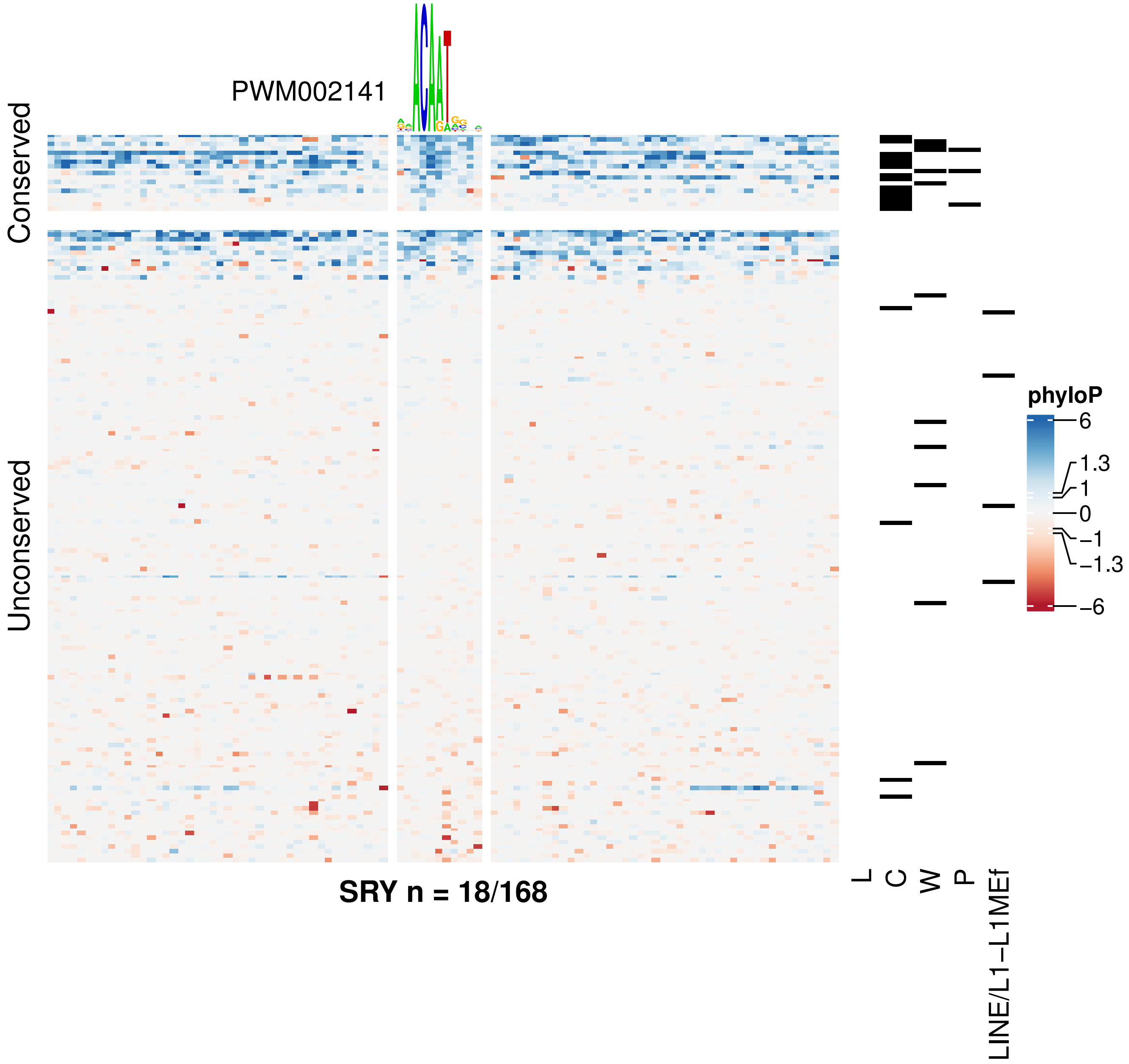

PWM002141
Conserved
Unconserved
SRY n = 18/168
L
C
W
P
LINE/L1−L1MEf
phyloP
6
1.3
1
0
−1
−1.3
−6

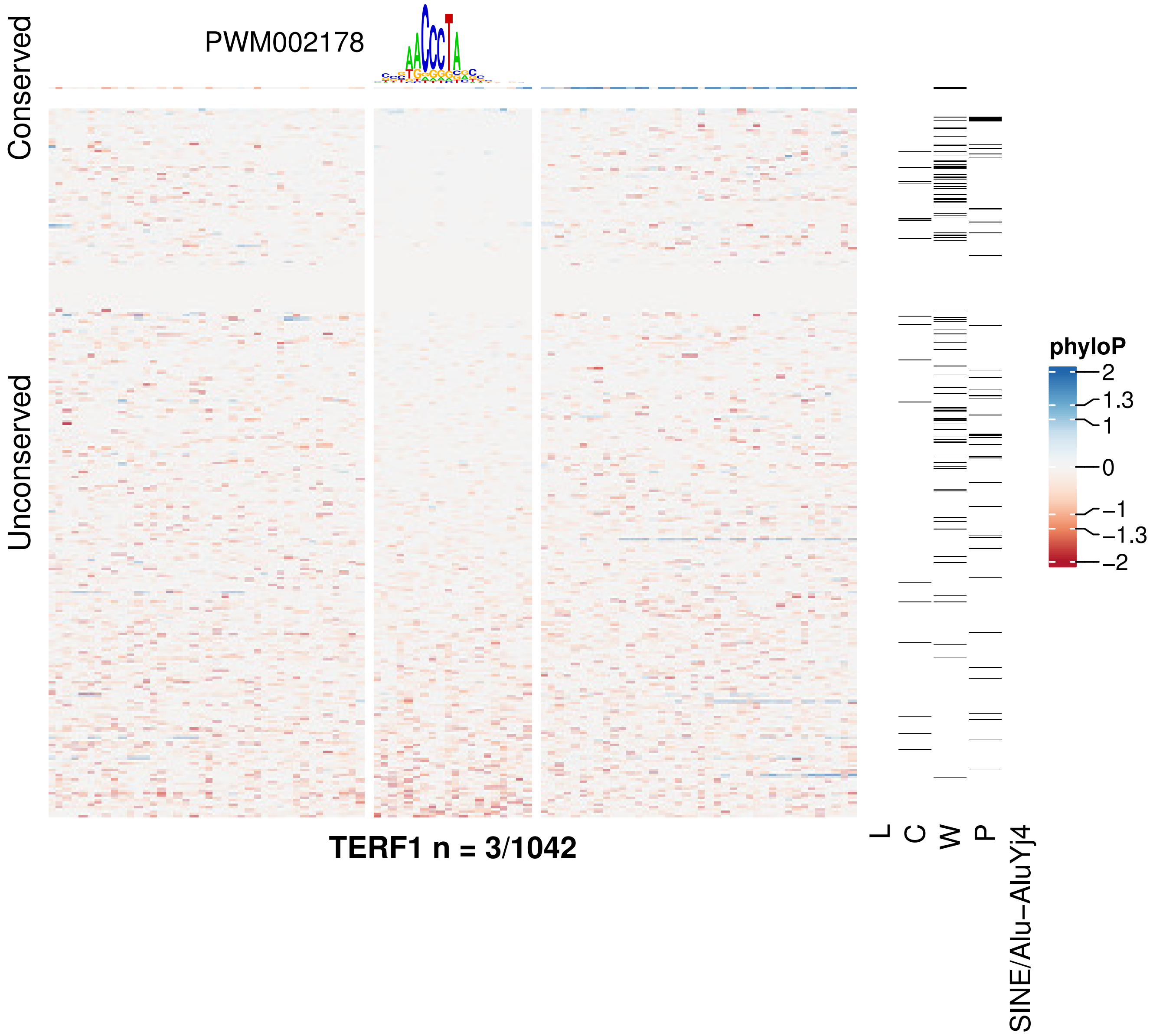

PWM002178
Conserved
Unconserved
TERF1 n = 3/1042
L
C
W
P
SINE/Alu−AluYj4
phyloP
2
1.3
1
0
−1
−1.3
−2

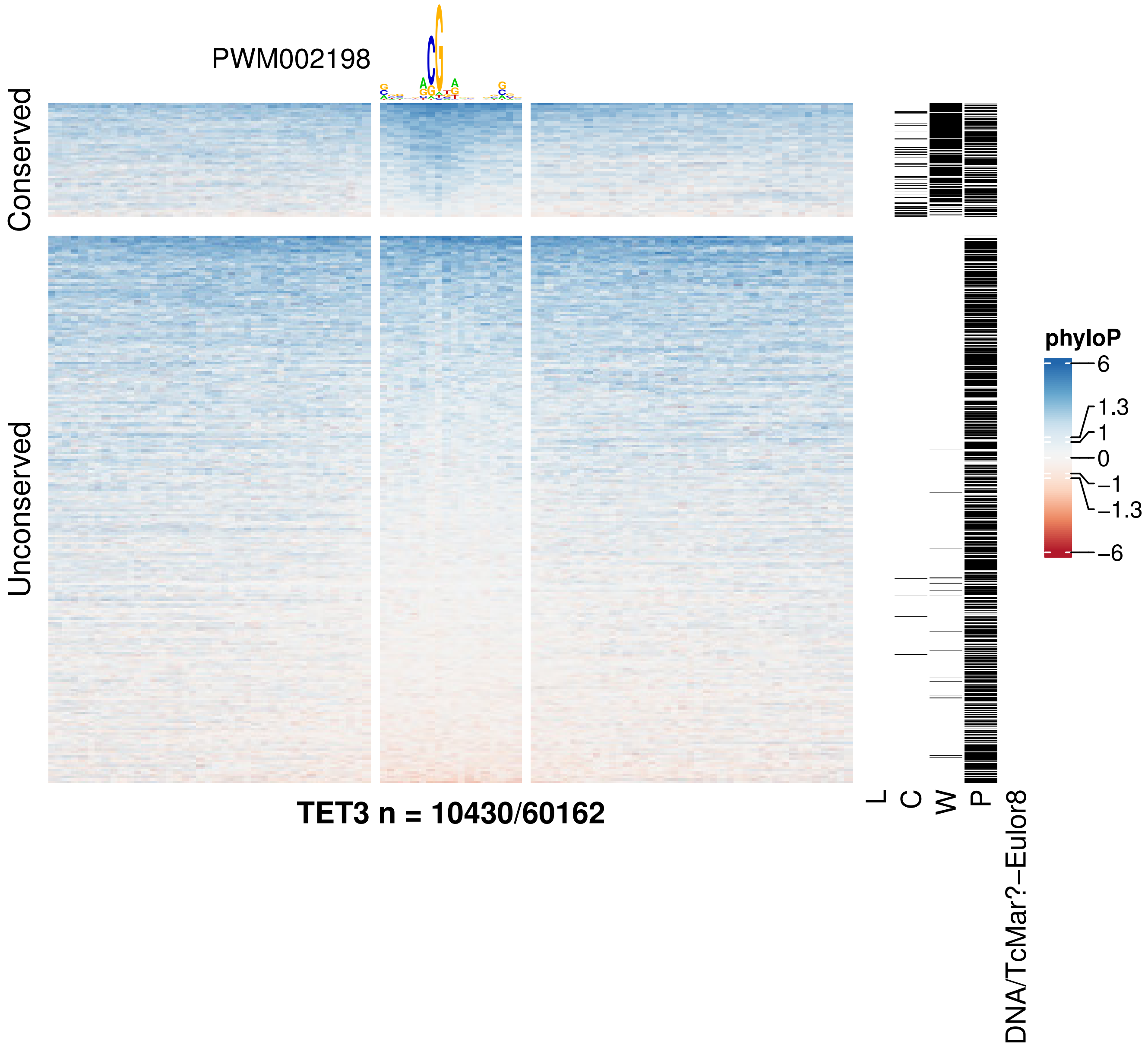

PWM002198
Conserved
Unconserved
TET3 n = 10430/60162
L
C
W
P
DNA/TcMar?−Eulor8
phyloP
6
1.3
1
0
−1
−1.3
−6

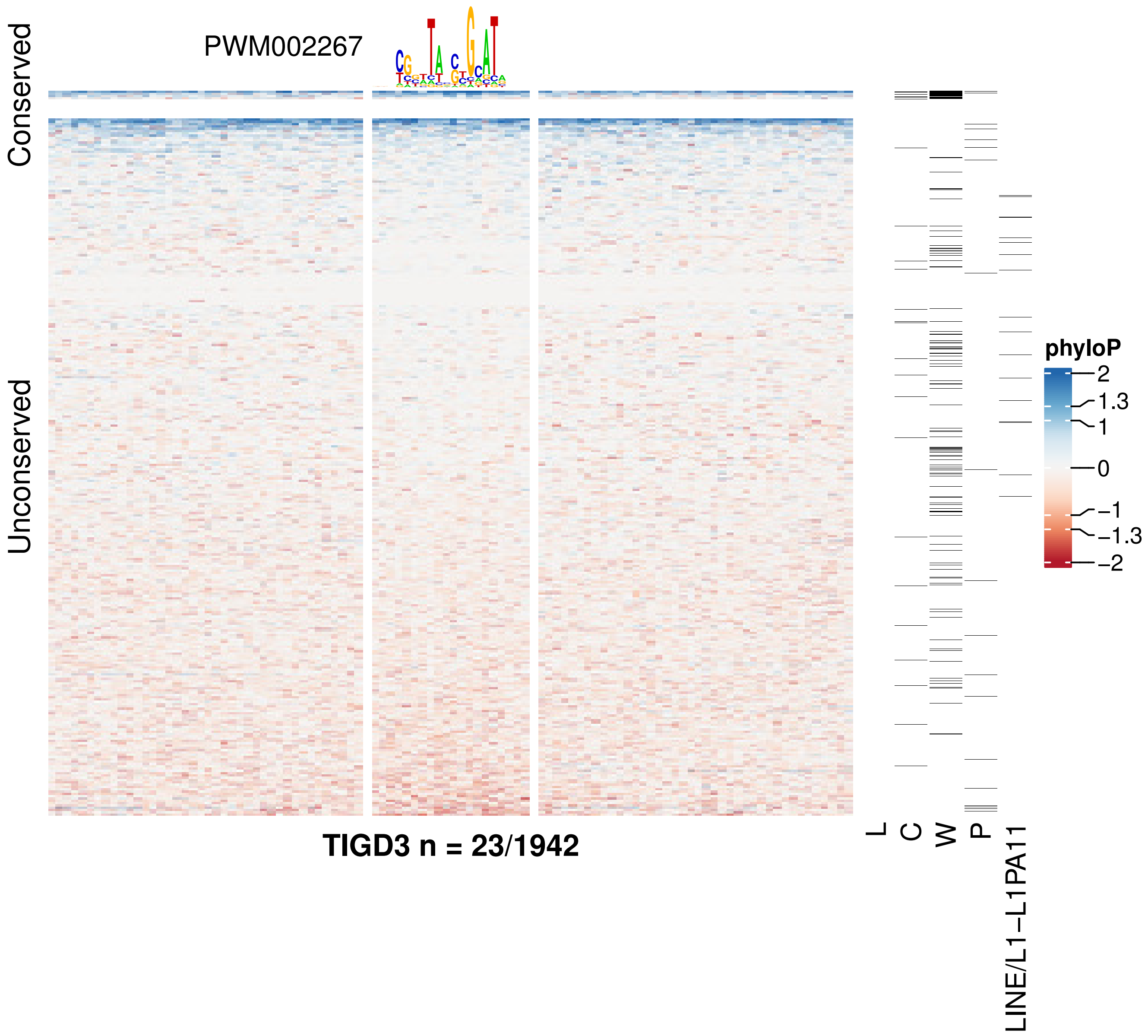

PWM002267
Conserved
Unconserved
TIGD3 n = 23/1942
L
C
W
P
LINE/L1−L1PA11
phyloP
2
1.3
1
0
−1
−1.3
−2

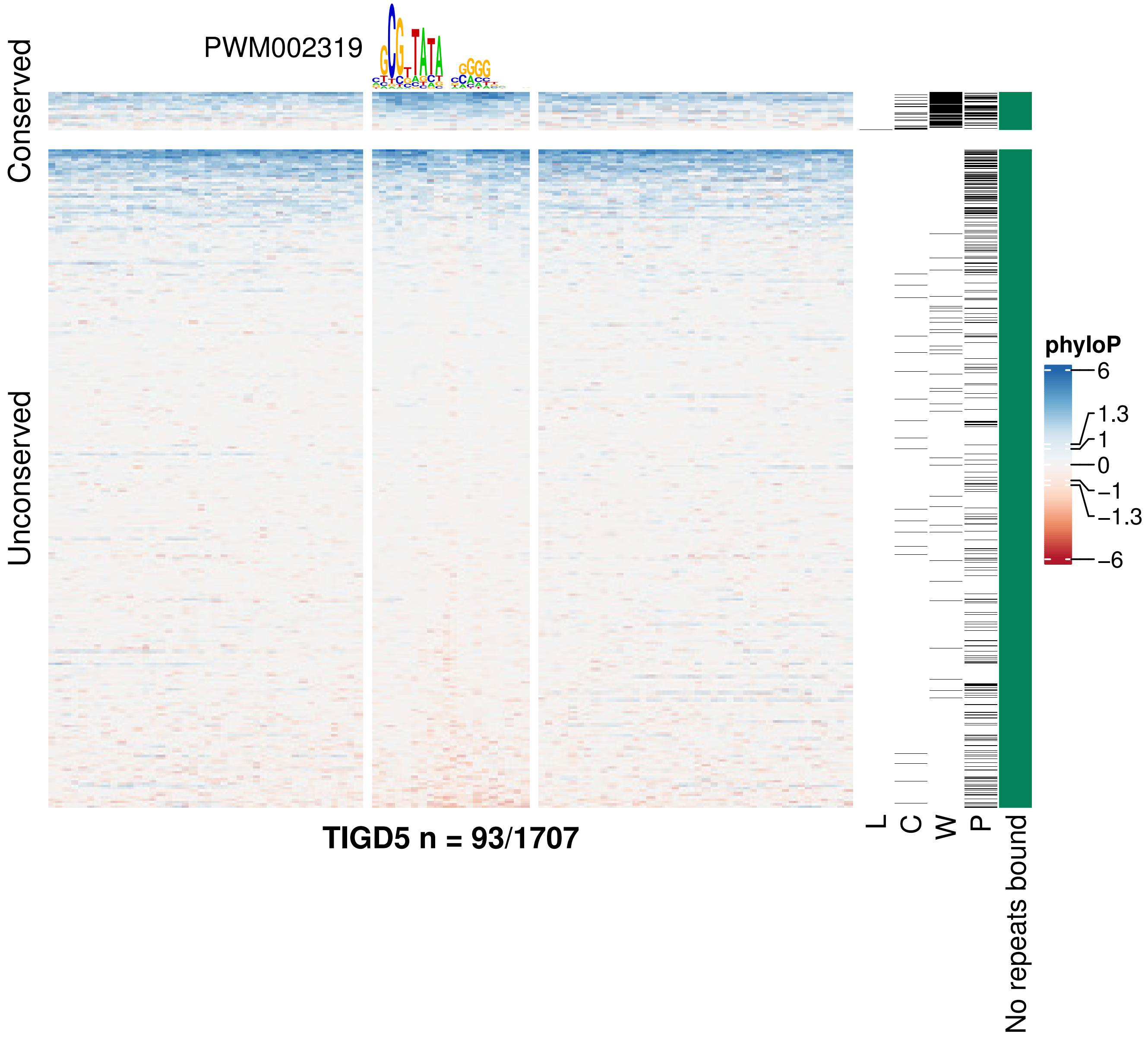

PWM002319
Conserved
Unconserved
TIGD5 n = 93/1707
L
C
W
P
No repeats bound
phyloP
6
1.3
1
0
−1
−1.3
−6

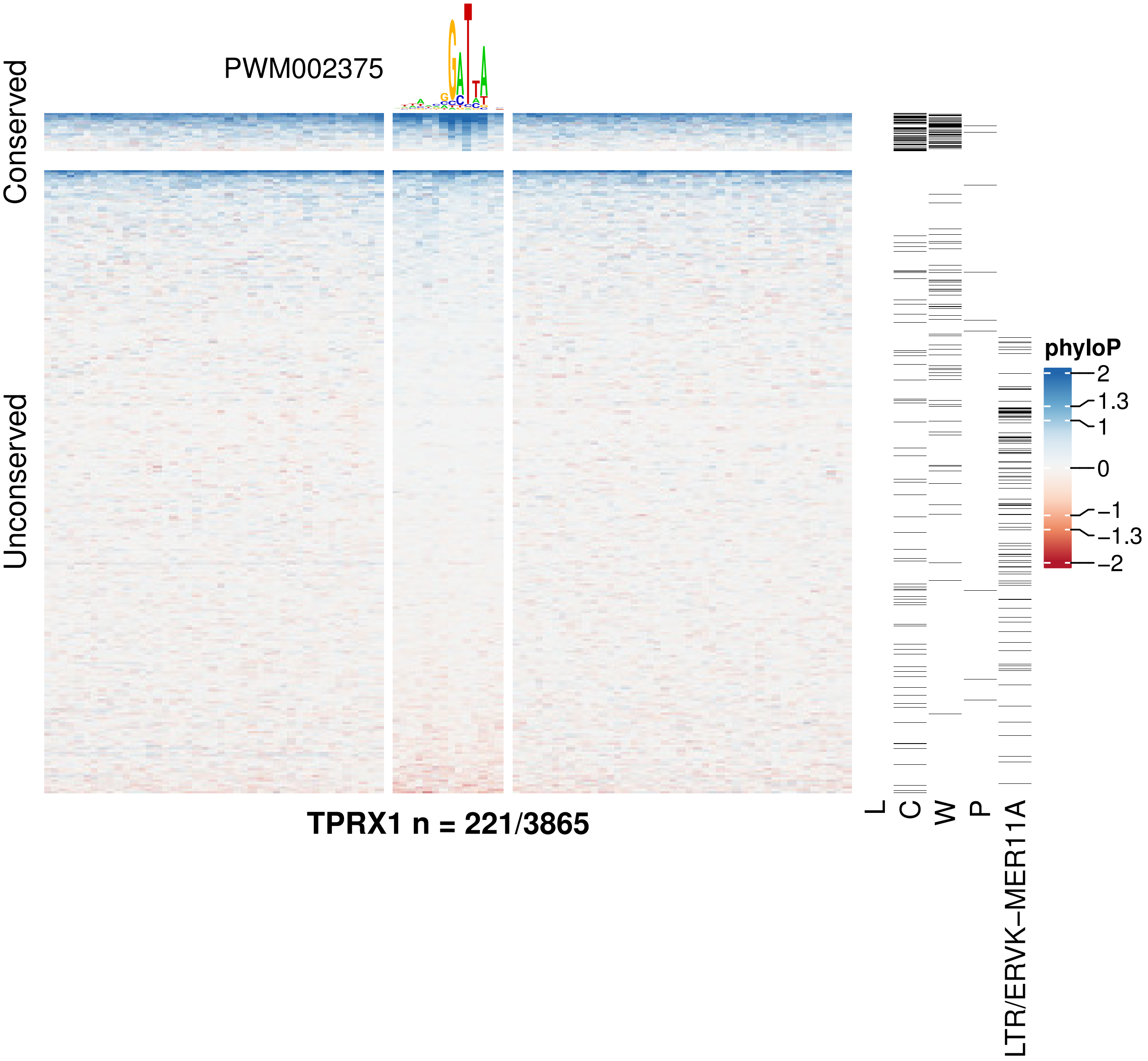
PWM002375
Conserved
Unconserved
TPRX1 n = 221/3865
L
C
W
P
LTR/ERVK−MER11A
phyloP
2
1.3
1
0
−1
−1.3
−2

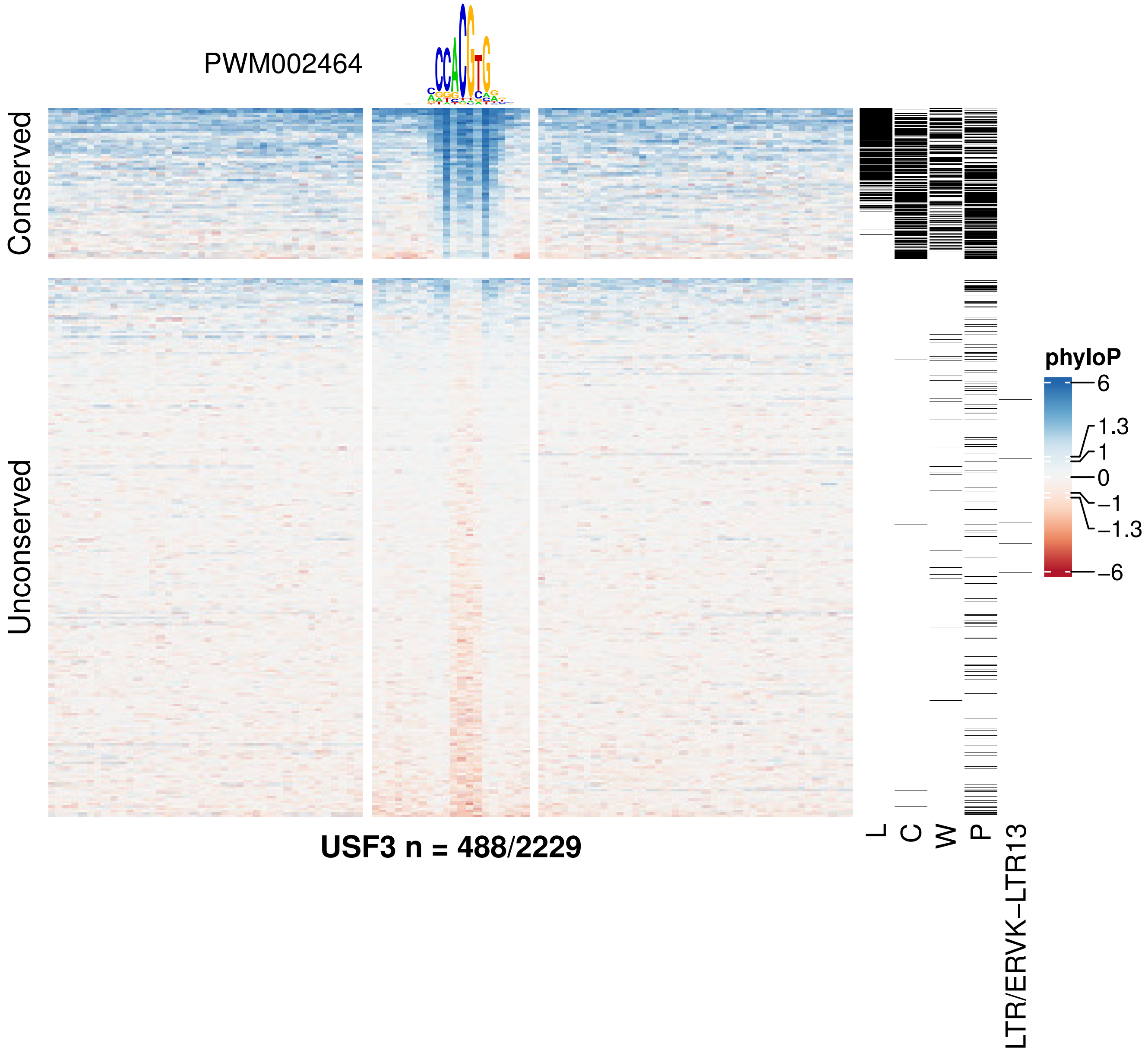
PWM002464
Conserved
Unconserved
USF3 n = 488/2229
L
C
W
P
LTR/ERVK–LTR13
phyloP
6
1.3
1
0
−1
−1.3
−6

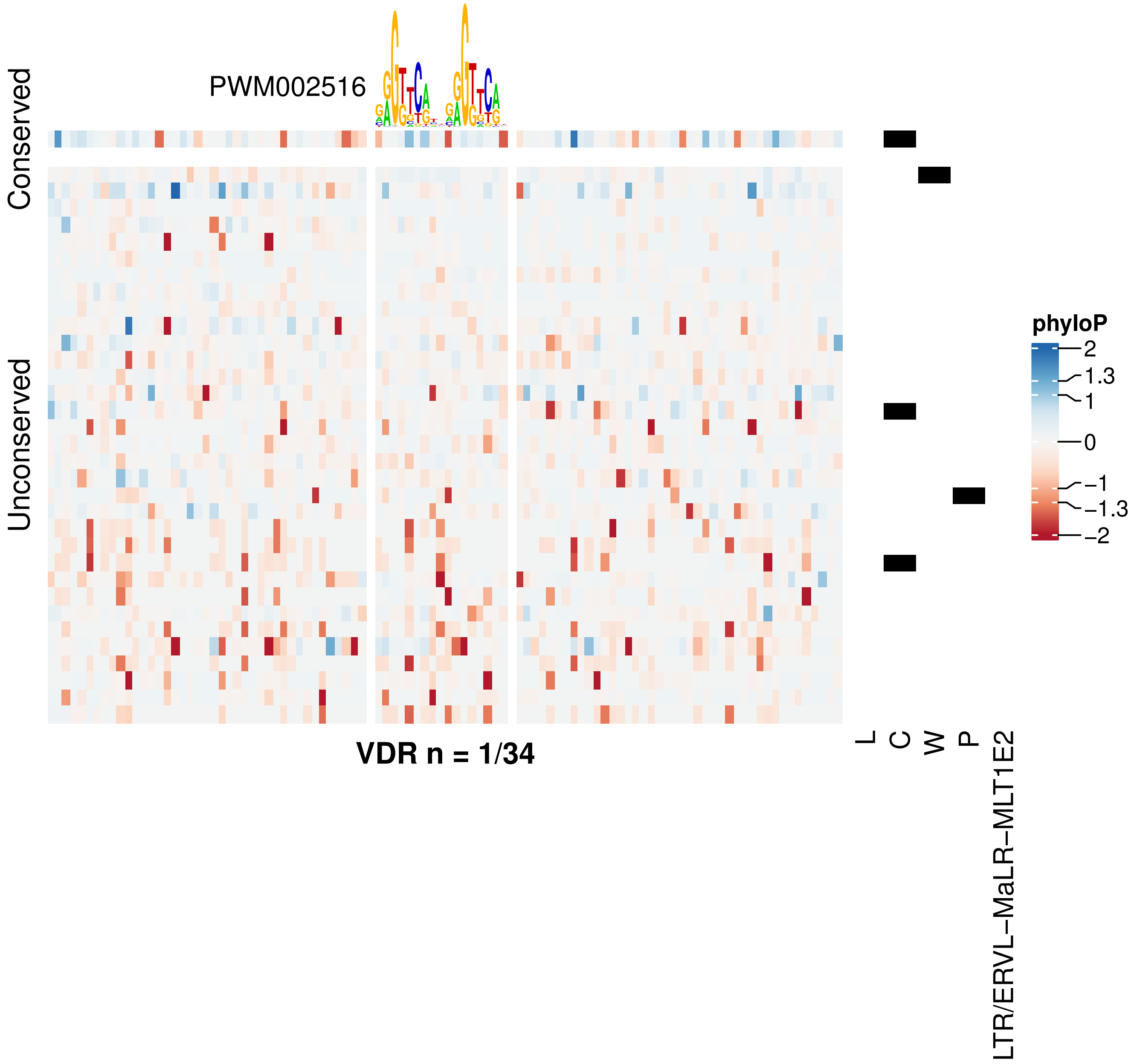
PWM002516
Conserved
Unconserved
VDR n = 1/34
phyloP
2
1.3
1
0
−1
−1.3
−2
L
C
W
P
LTR/ERVL−MaLR−MLT1E2

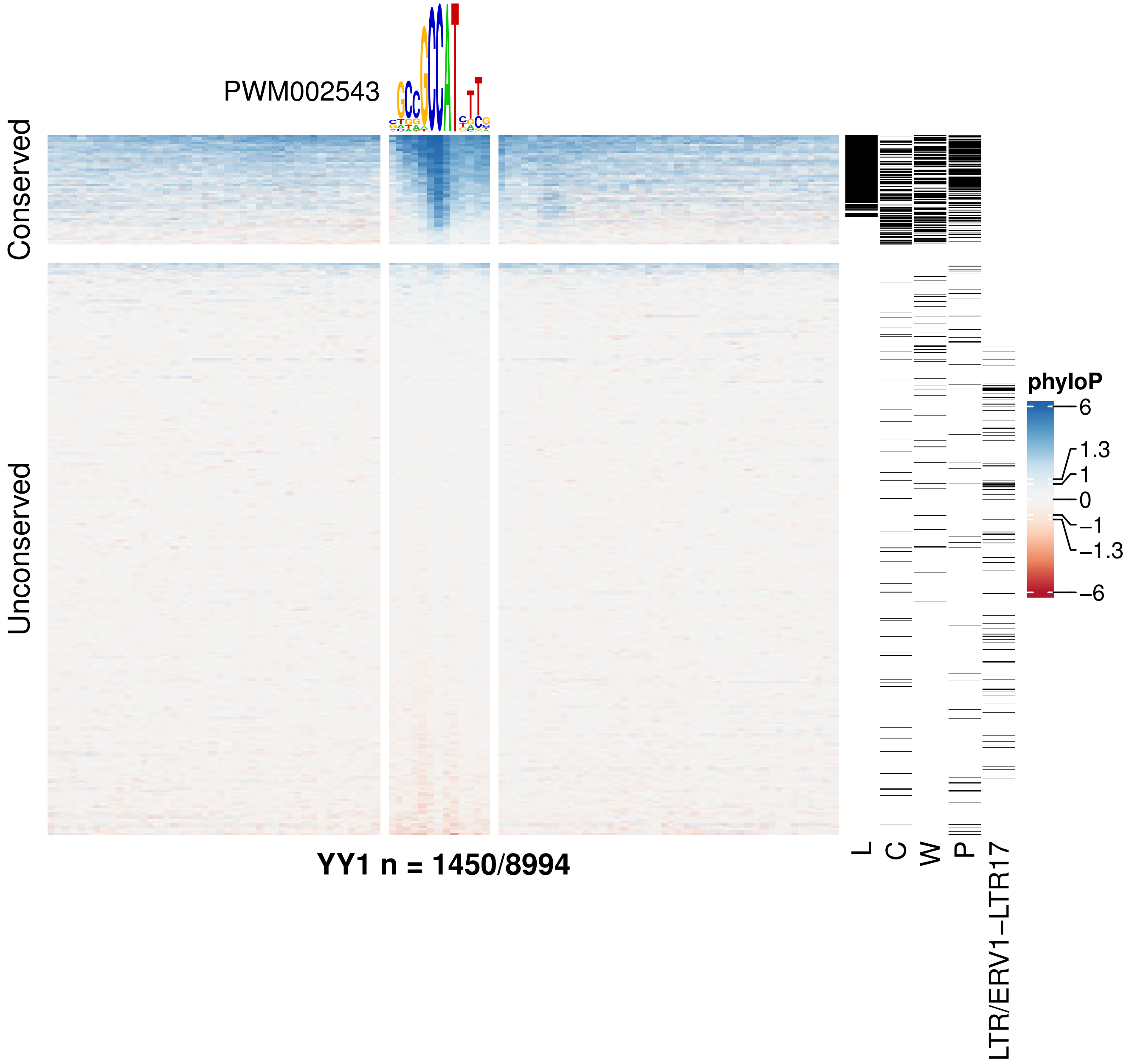
PWM002543
Conserved
Unconserved
YY1 n = 1450/8994
L
C
W
P
LTR/ERV1−LTR17
phyloP
6
1.3
1
0
−1
−1.3
−6

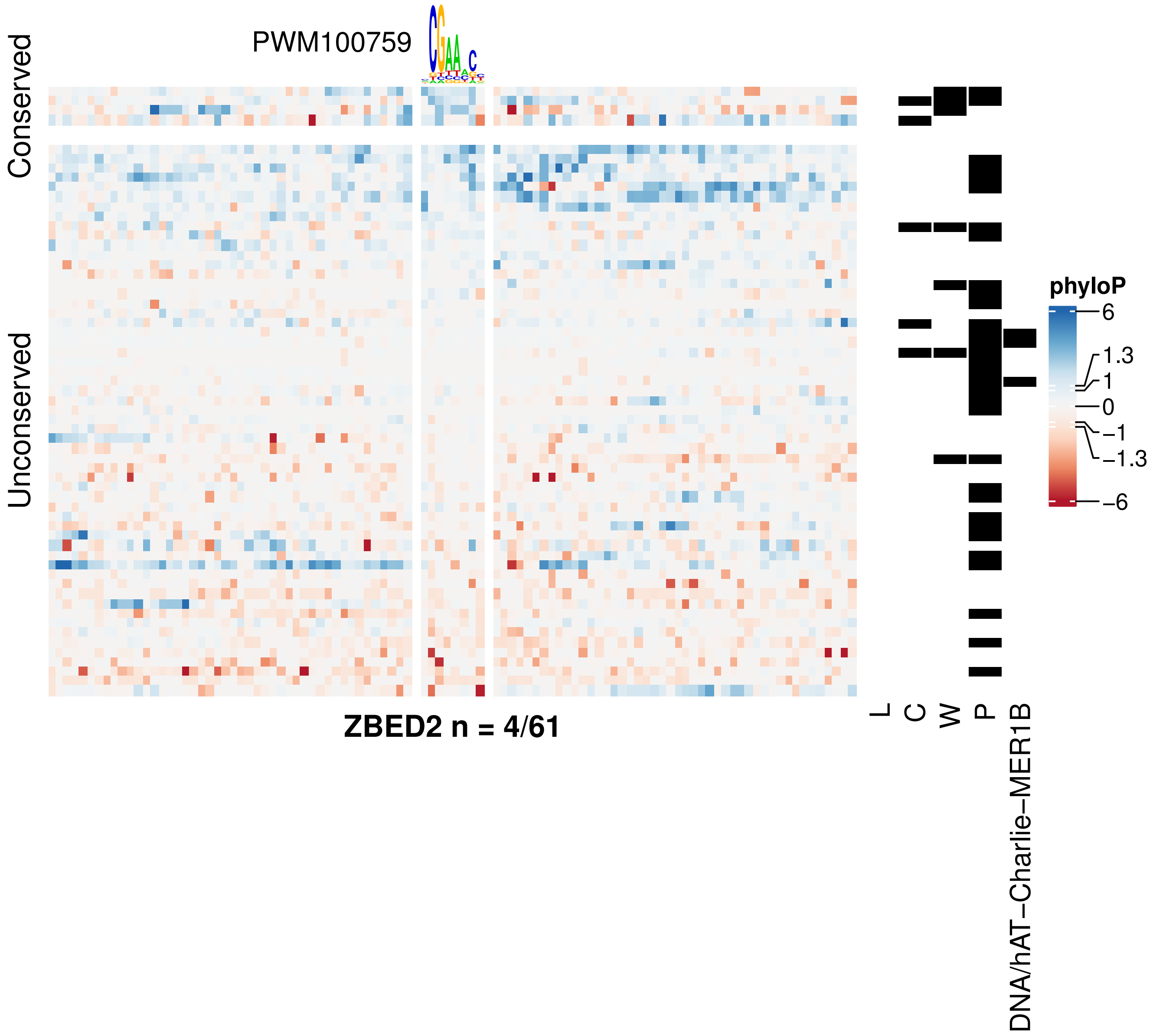
PWM100759
Conserved
Unconserved
ZBED2 n = 4/61
phyloP
6
1.3
1
0
−1
−1.3
−6
L
C
W
P
DNA/hAT−Charlie−MER1B

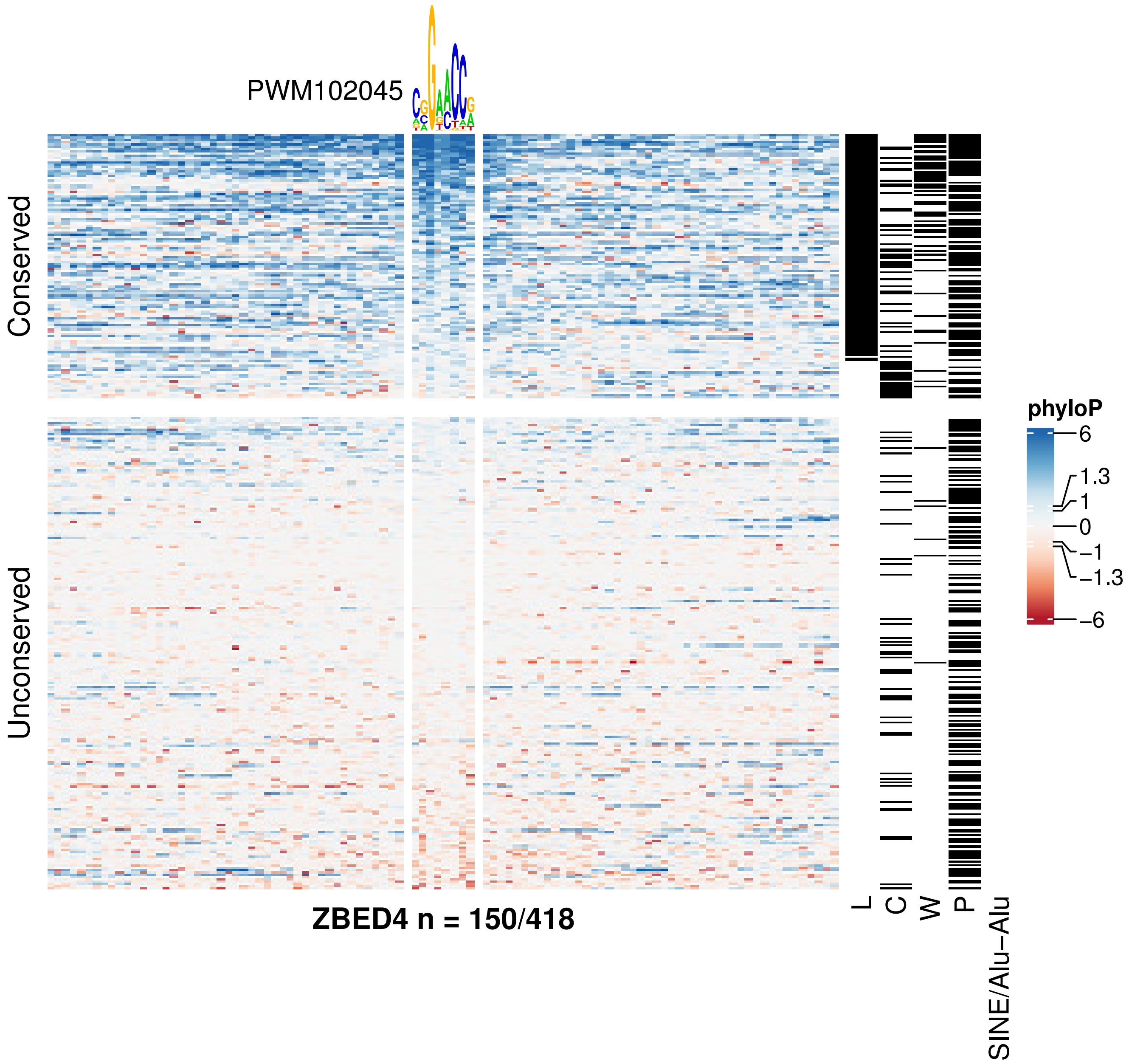
PWM102045
Conserved
Unconserved
ZBED4 n = 150/418
L
C
W
P
SINE/Alu−Alu
phyloP
6
1.3
1
0
−1
−1.3
−6

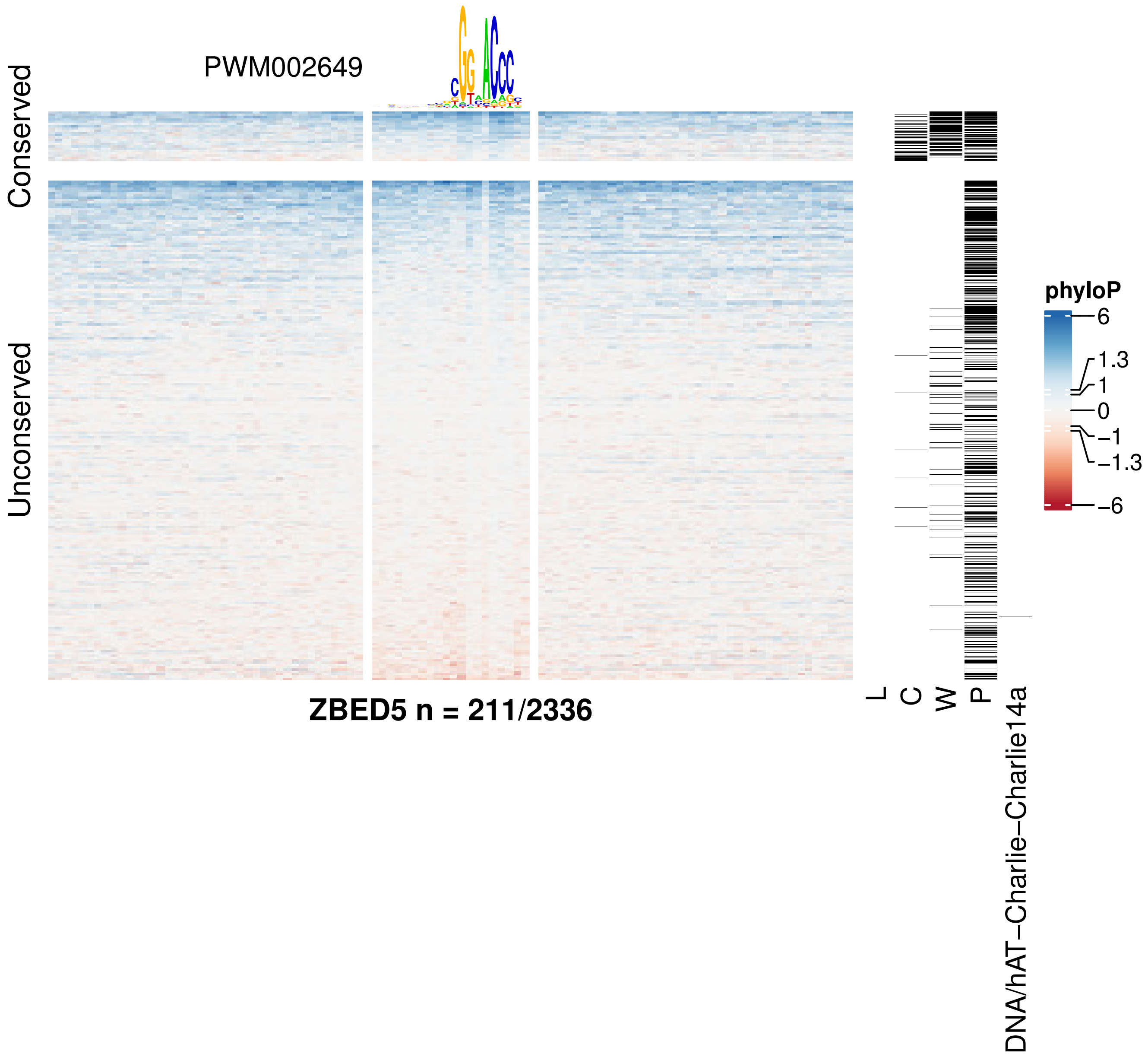
PWM002649
Conserved
Unconserved
ZBED5 n = 211/2336
L
C
W
P
DNA/hAT−Charlie−Charlie14a
phyloP
6
1.3
1
0
−1
−1.3
−6

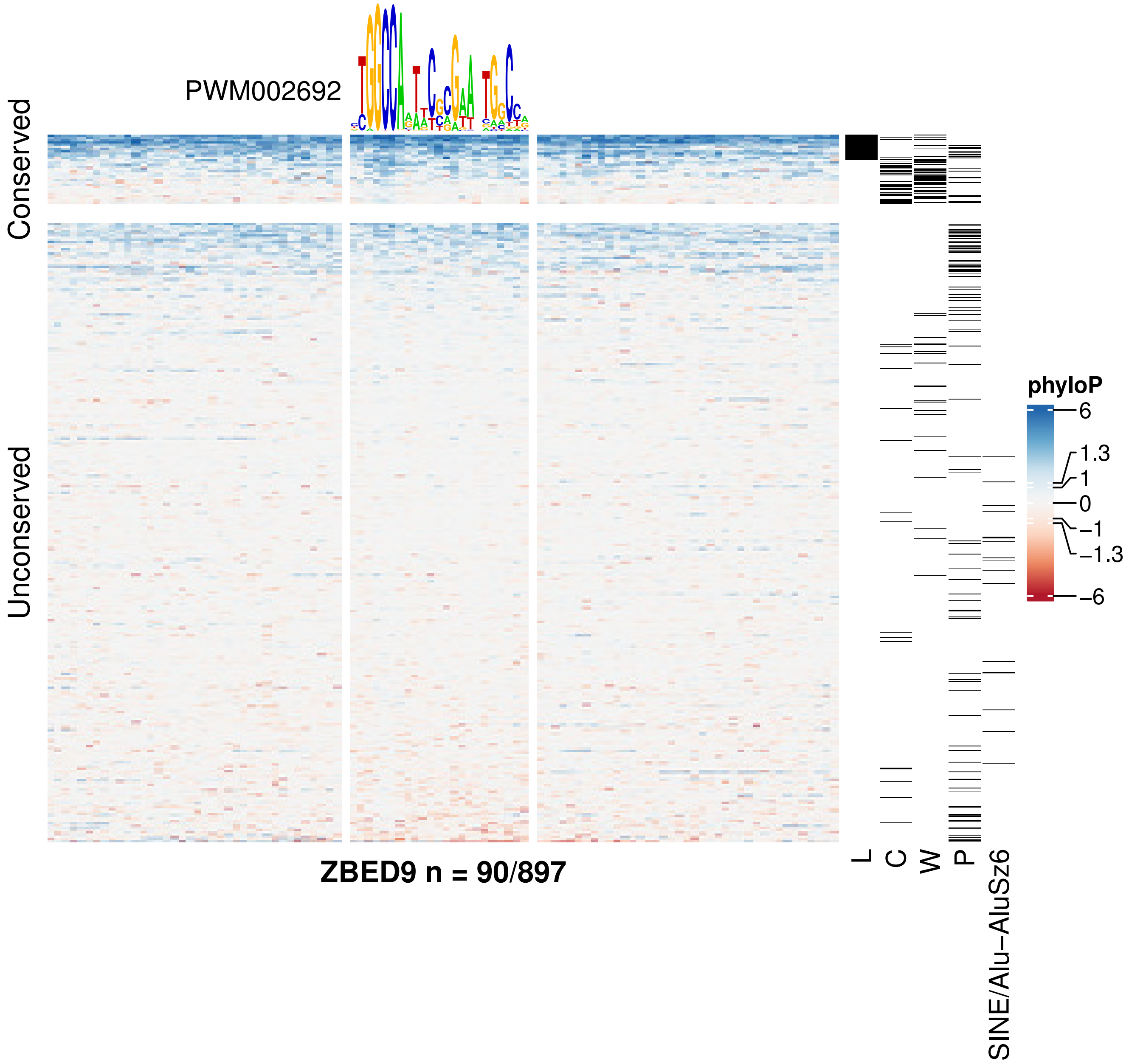
PWM002692
Conserved
Unconserved
ZBED9 n = 90/897
L
C
W
P
SINE/Alu−AluSz6
phyloP
6
1.3
1
0
−1
−1.3
−6

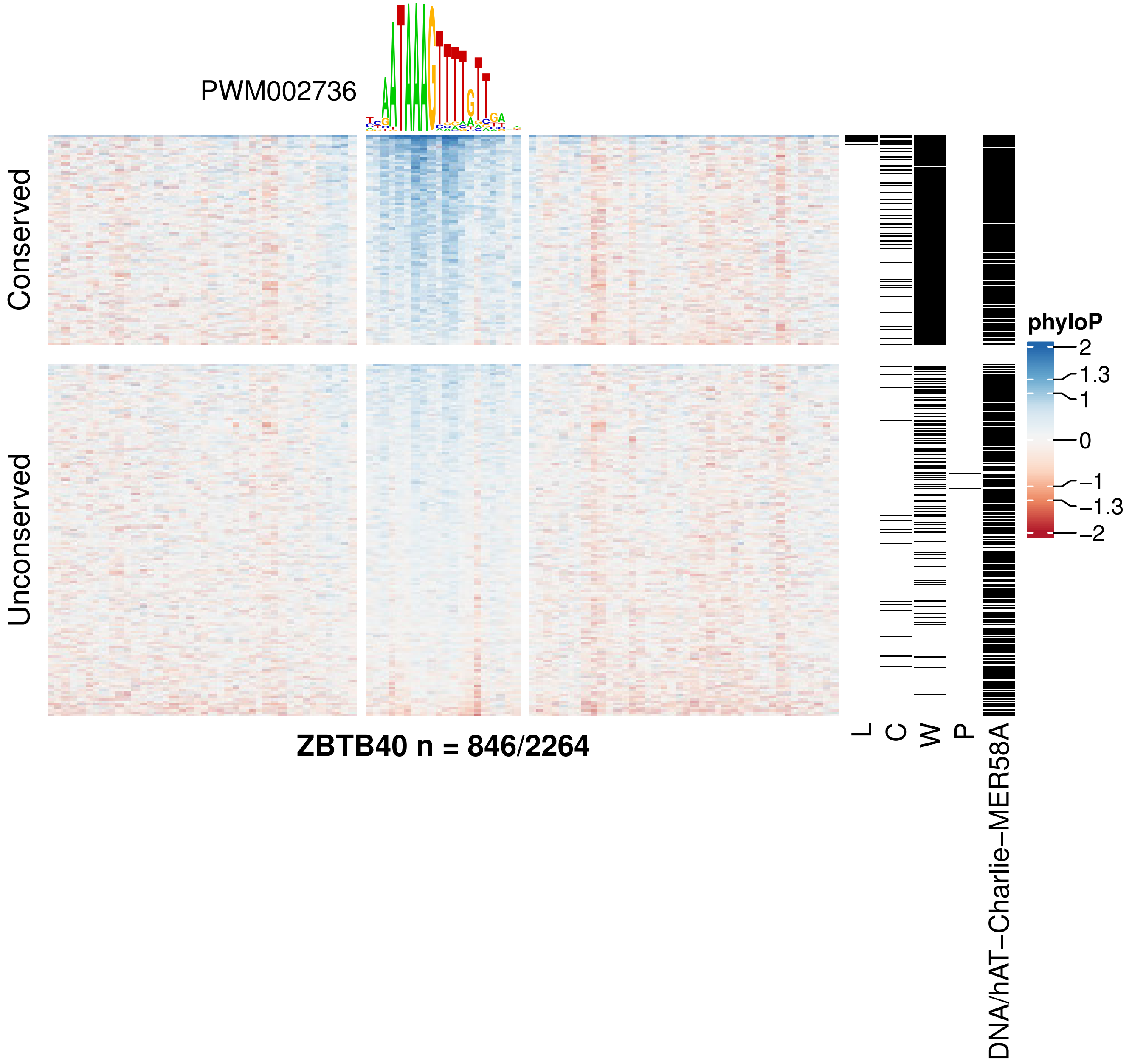

PWM002736
Conserved
Unconserved
phyloP
2
1.3
1
0
−1
−1.3
−2
L
C
W
P
DNA/hAT−Charlie−MER58A
ZBTB40 n = 846/2264

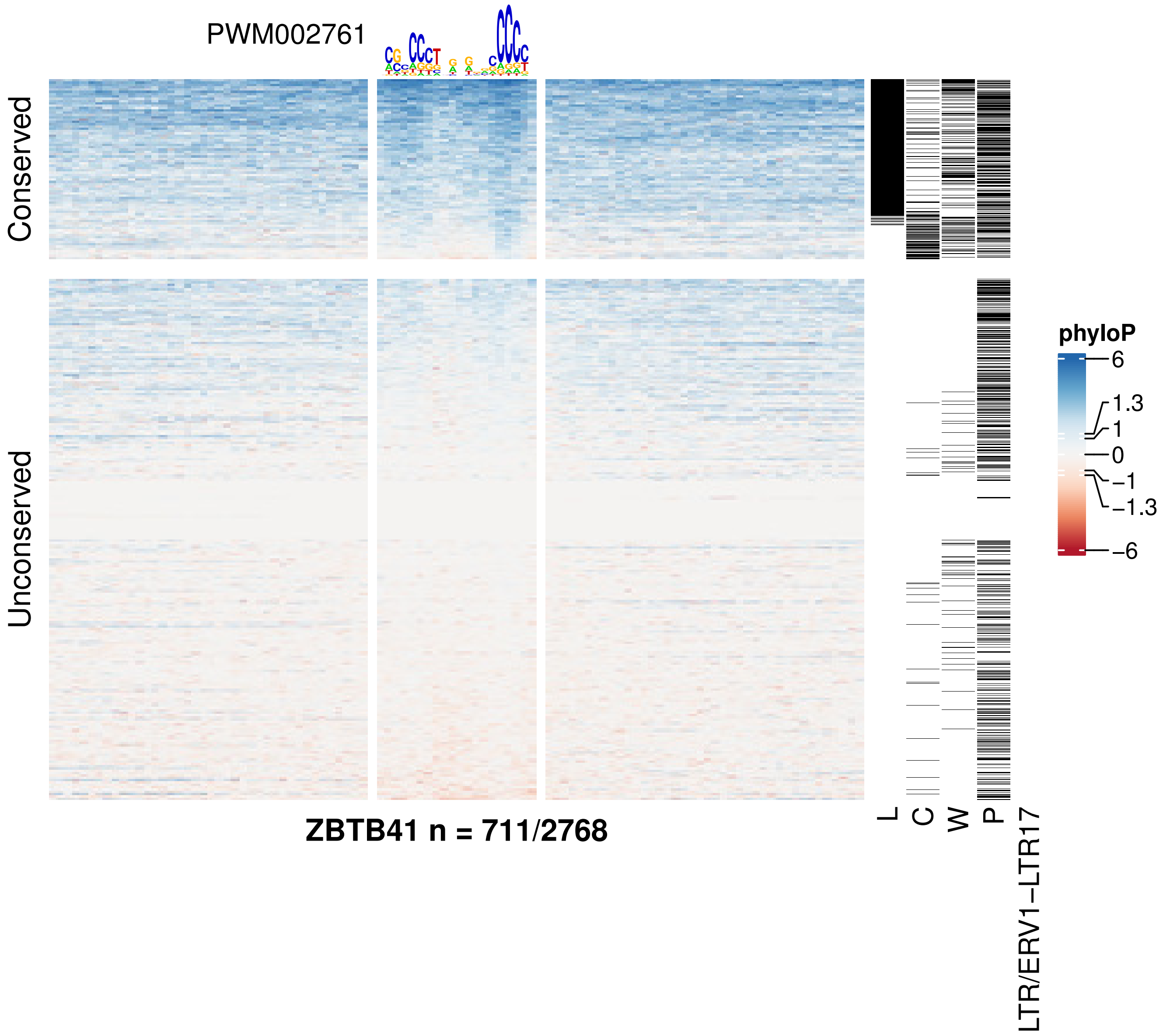
PWM002761
Conserved
Unconserved
ZBTB41 n = 711/2768
L
C
W
P
LTR/ERV1−LTR17
phyloP
6
1.3
1
0
−1
−1.3
−6

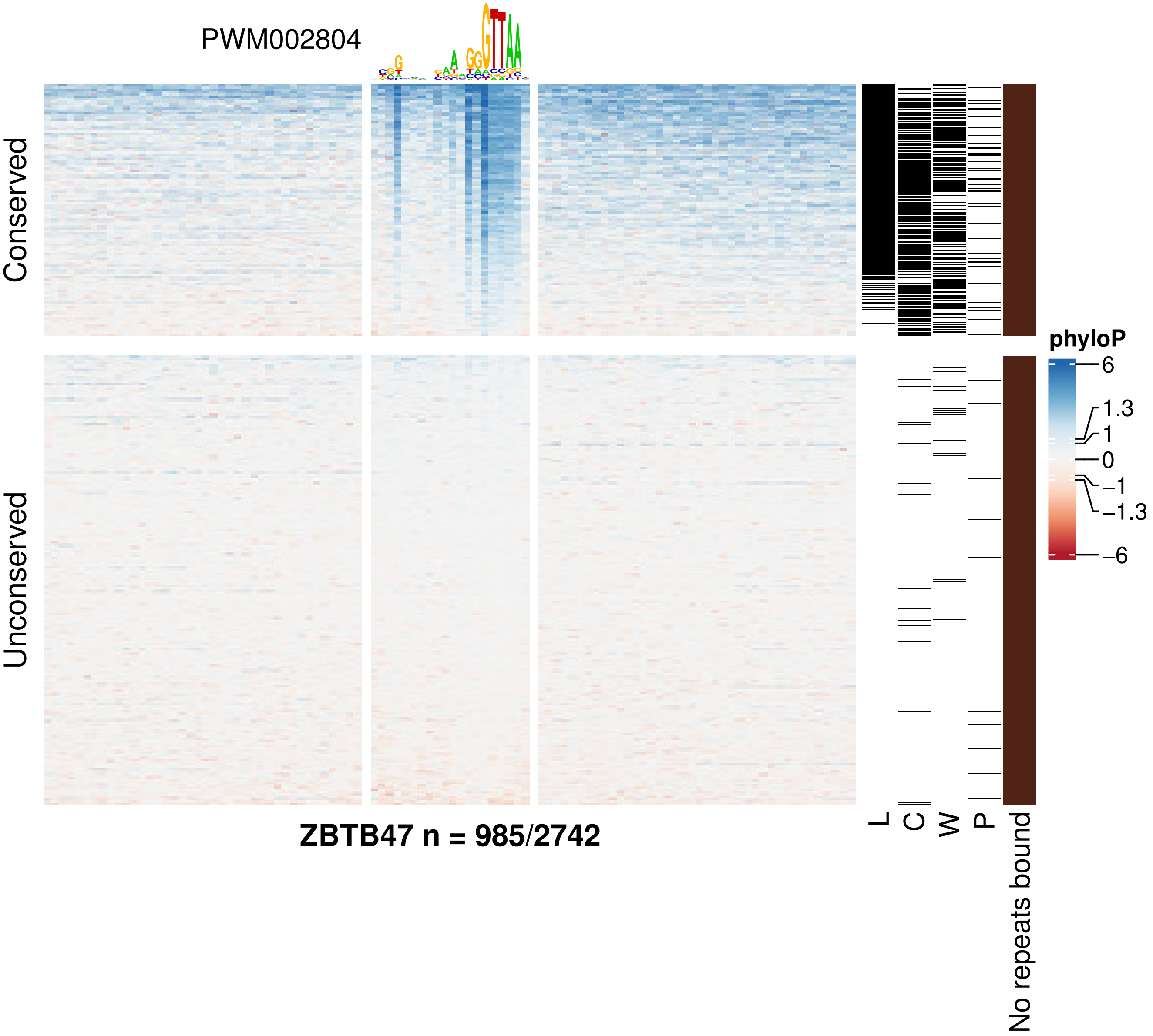
PWM002804
Conserved
Unconserved
phyloP
6
1.3
1
0
−1
−1.3
−6
L
C
W
P
No repeats bound
ZBTB47 n = 985/2742

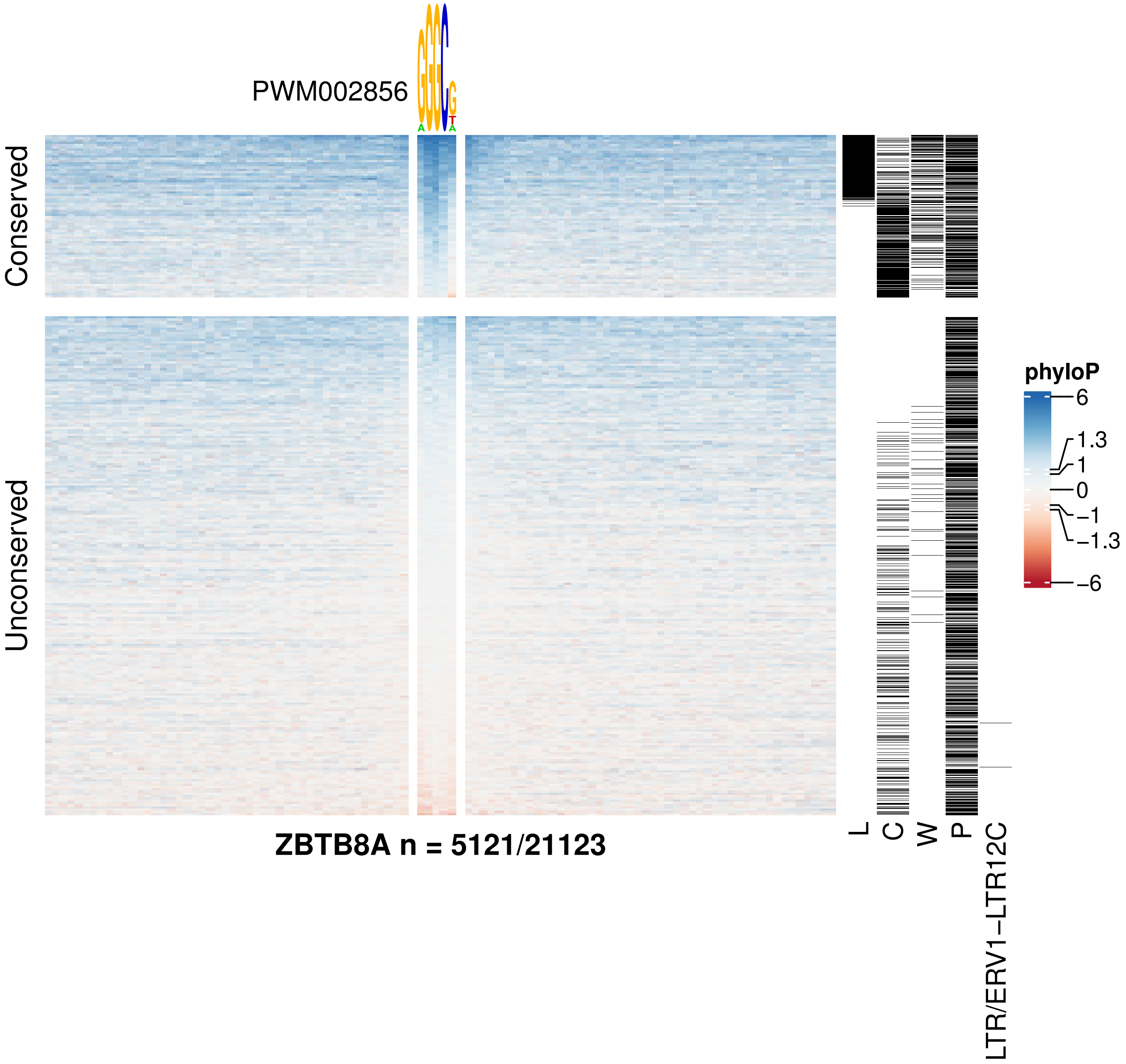

PWM002856
Conserved
Unconserved
ZBTB8A n = 5121/21123
L
C
W
P
LTR/ERV1−LTR12C
phyloP
6
1.3
1
0
−1
−1.3
−6

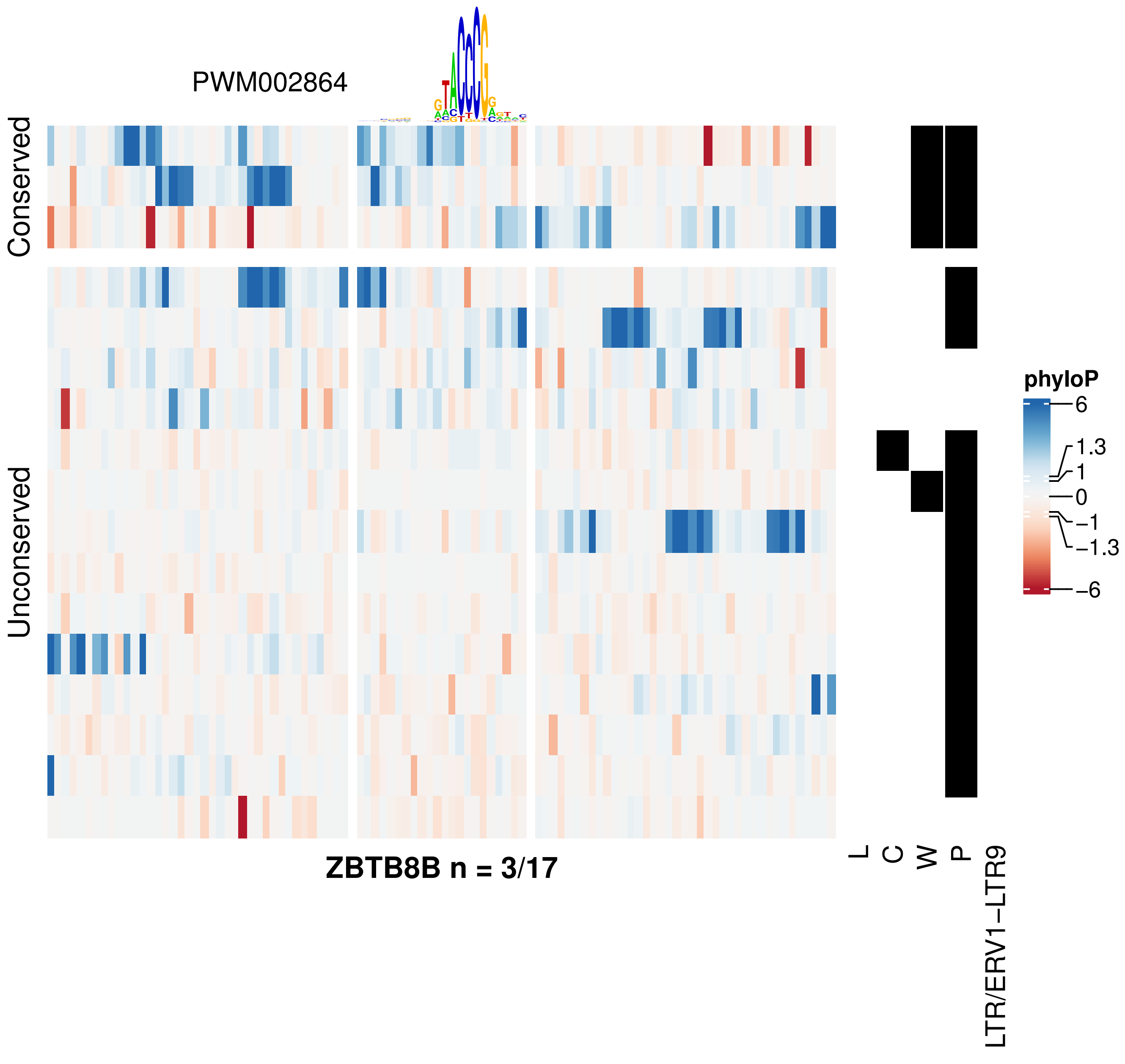
PWM002864
Conserved
Unconserved
ZBTB8B n = 3/17
L
C
W
P
LTR/ERV1−LTR9
phyloP
6
1.3
1
0
−1
−1.3
−6

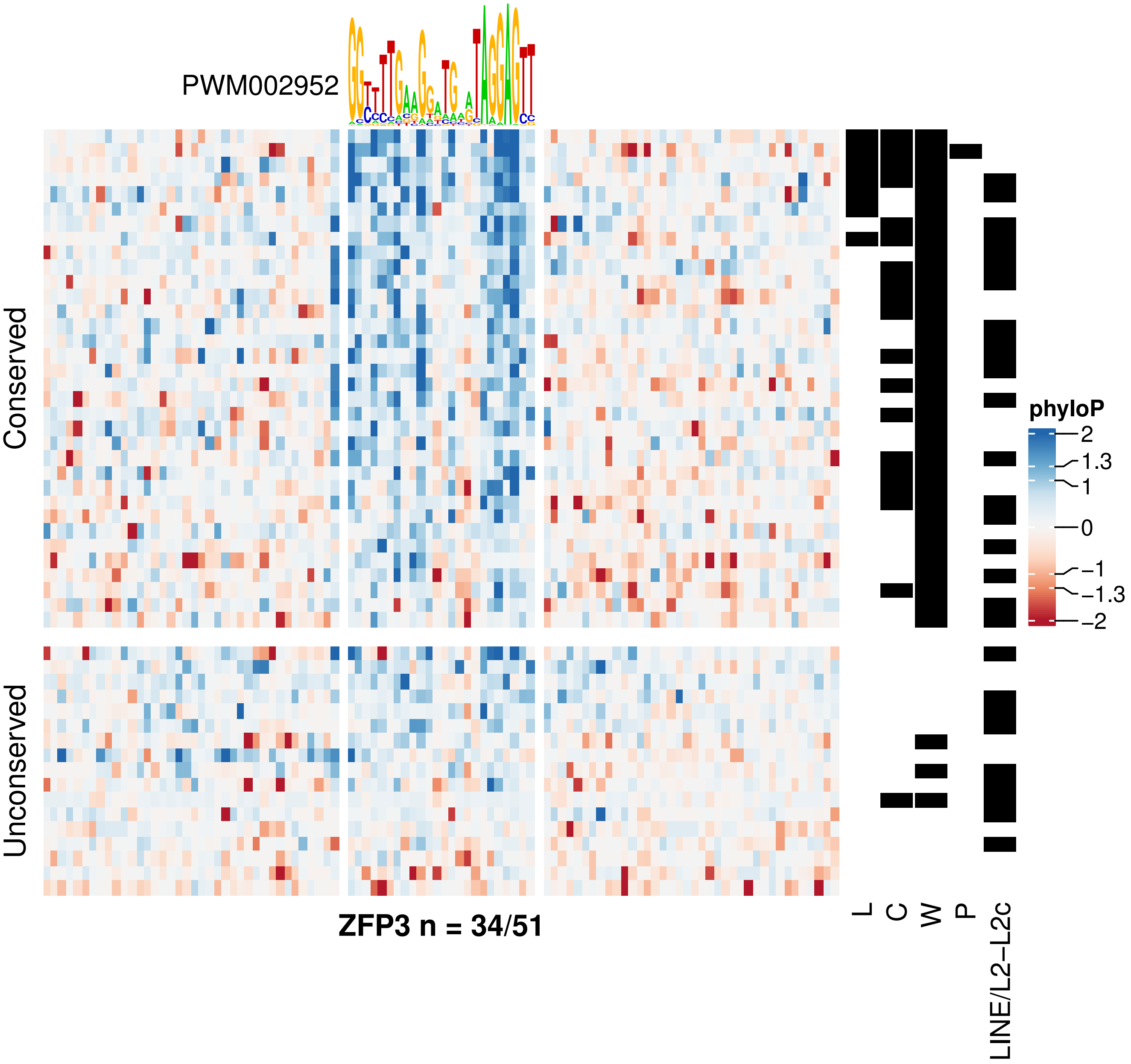
PWM002952
Conserved
Unconserved
ZFP3 n = 34/51
L
C
W
P
LINE/L2−L2c
phyloP
2
1.3
1
0
−1
−1.3
−2

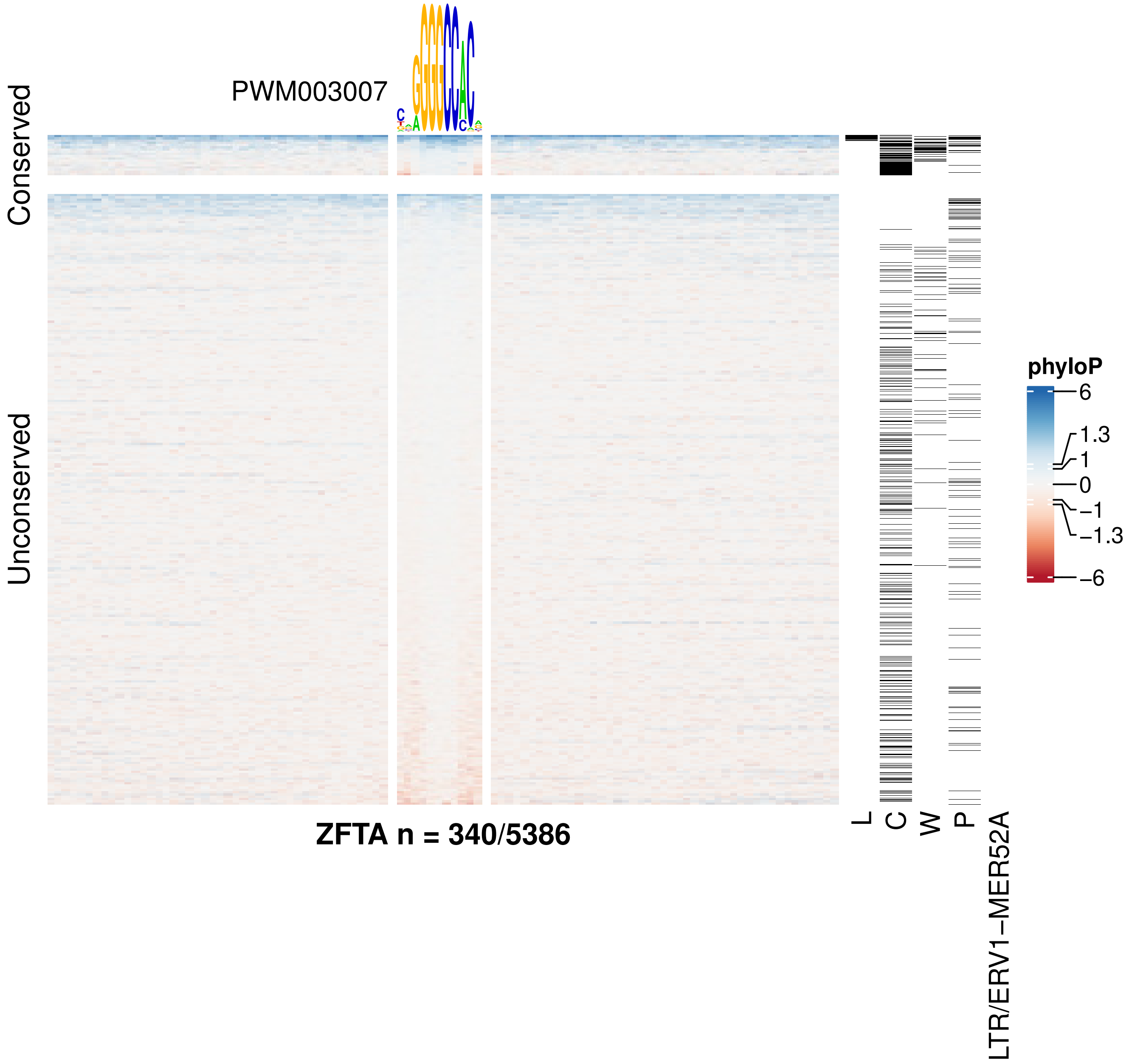
PWM003007
Conserved
Unconserved
ZFTA n = 340/5386
L
C
W
P
LTR/ERV1−MER52A
phyloP
6
1.3
1
0
−1
−1.3
−6

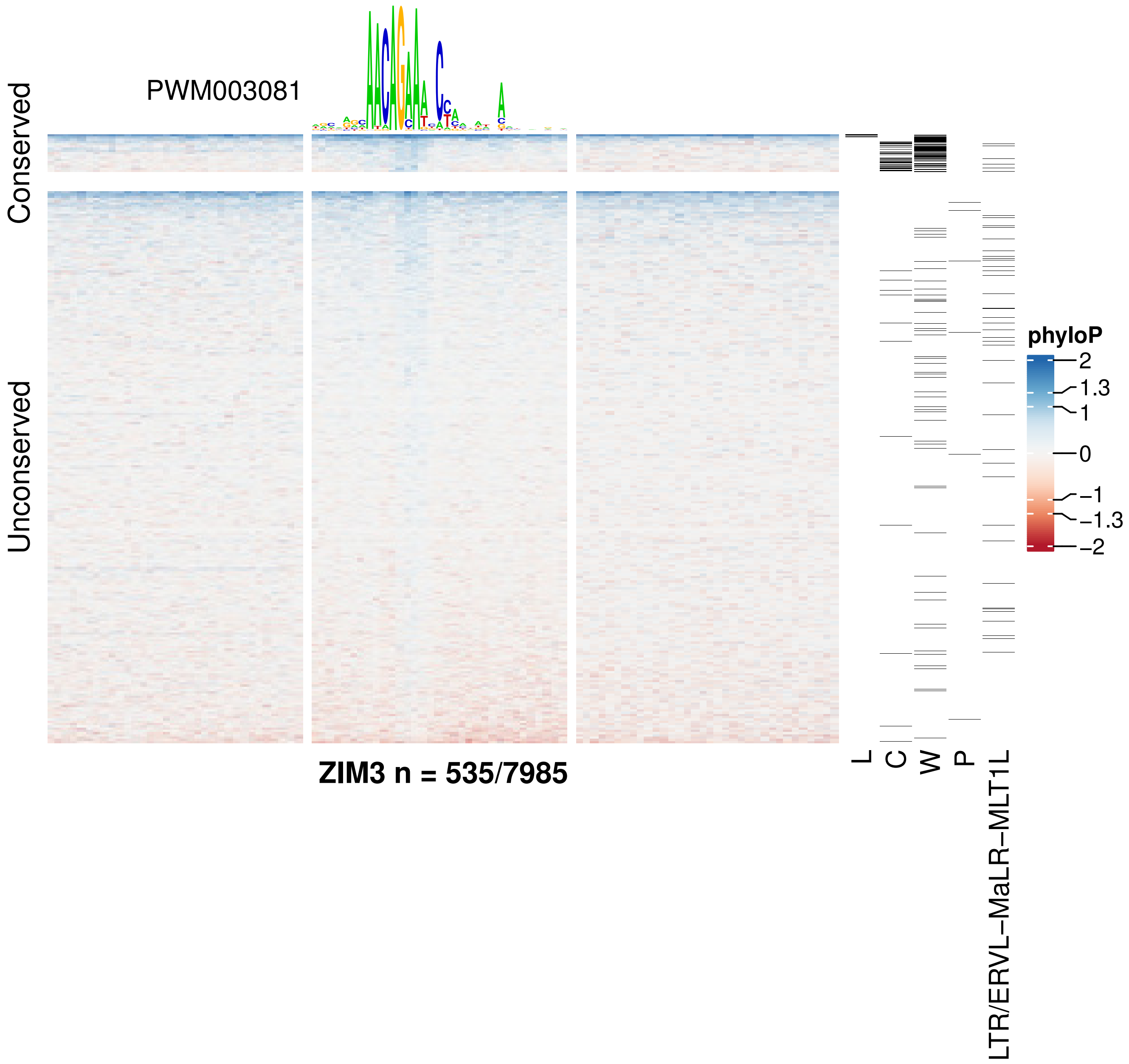

PWM003081
Conserved
Unconserved
ZIM3 n = 535/7985
L
C
W
P
LTR/ERVL–MaLR–MLT1L
phyloP
2
1.3
1
0
−1
−1.3
−2

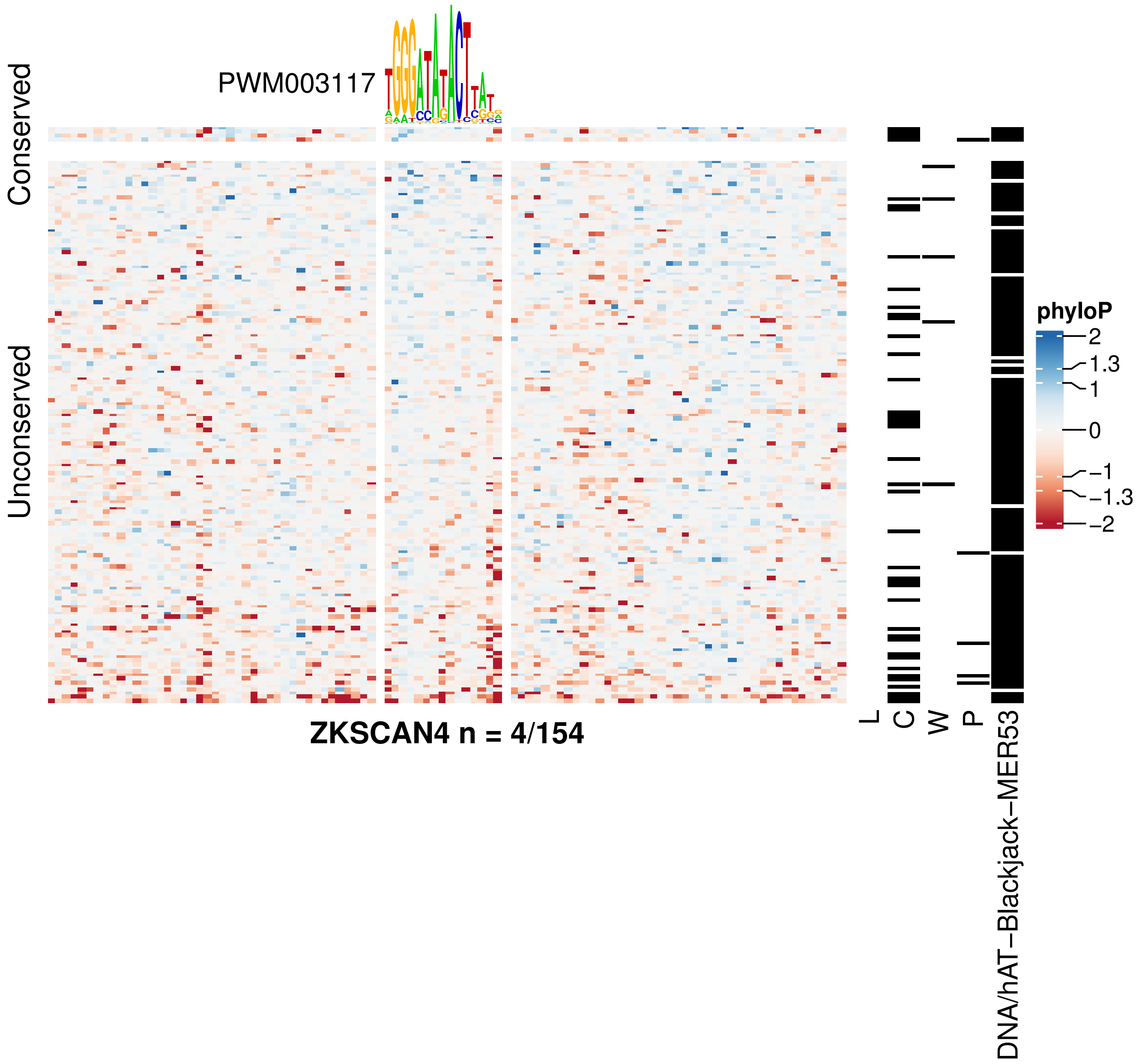

PWM003117
Conserved
Unconserved
ZKSCAN4 n = 4/154
L
C
W
P
DNA/hAT−Blackjack−MER53
phyloP
2
1.3
1
0
−1
−1.3
−2

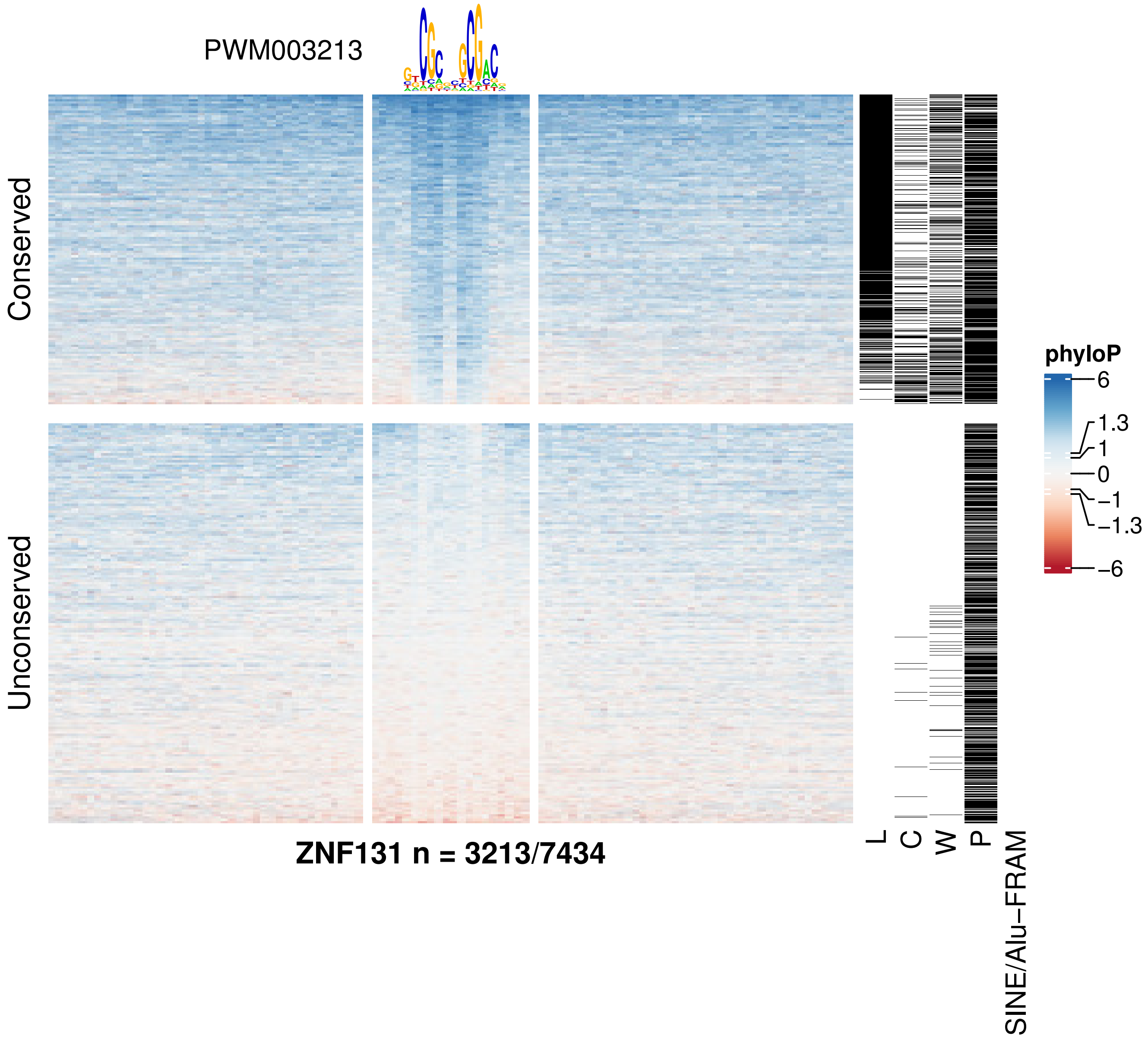

PWM003213
Conserved
Unconserved
ZNF131 n = 3213/7434
L
C
W
P
SINE/Alu−FRAM
phyloP
6
1.3
1
0
−1
−1.3
−6

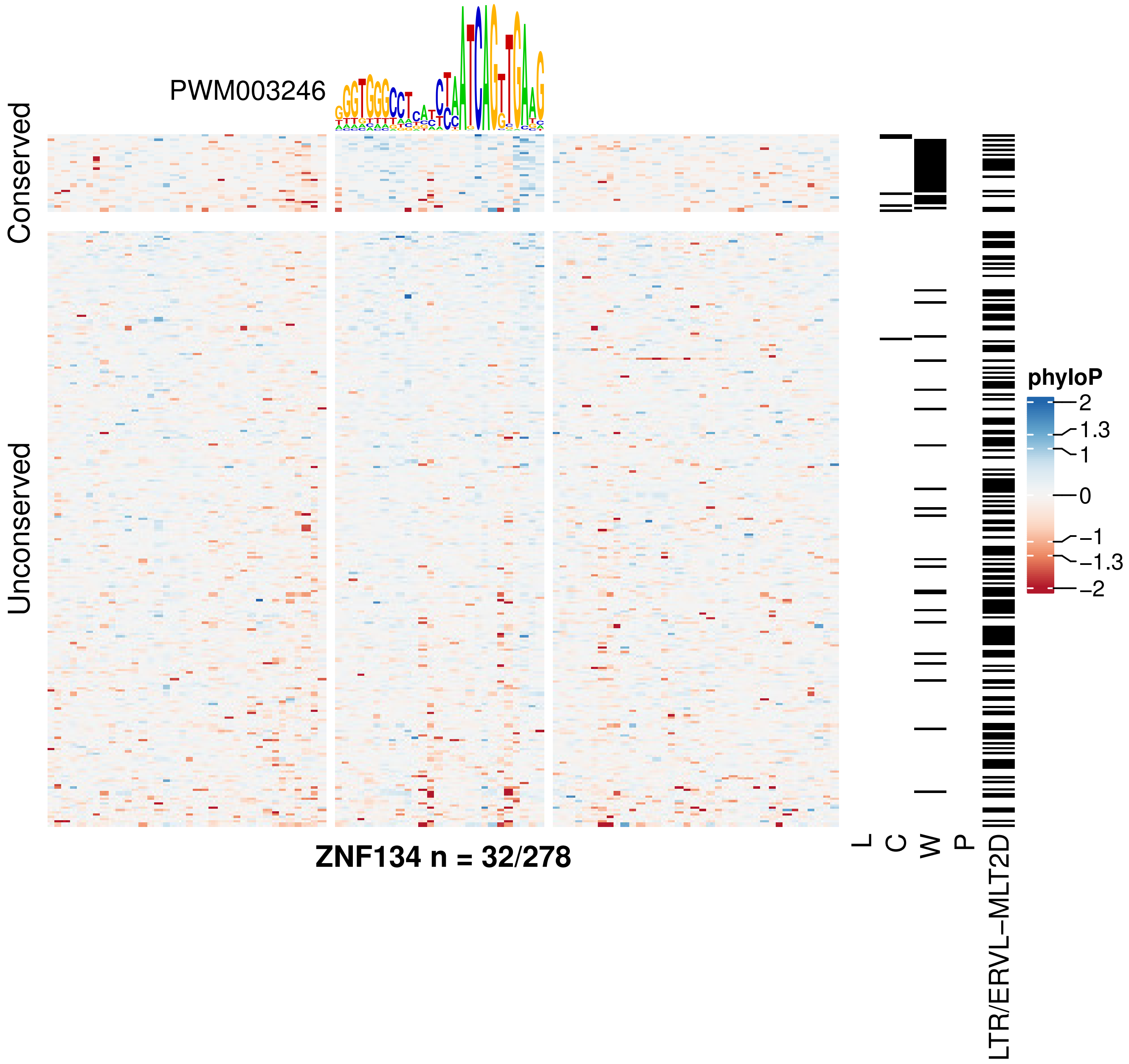
PWM003246
Conserved
Unconserved
ZNF134 n = 32/278
L
C
W
P
LTR/ERVL–MLT2D
phyloP
2
1.3
1
0
−1
−1.3
−2

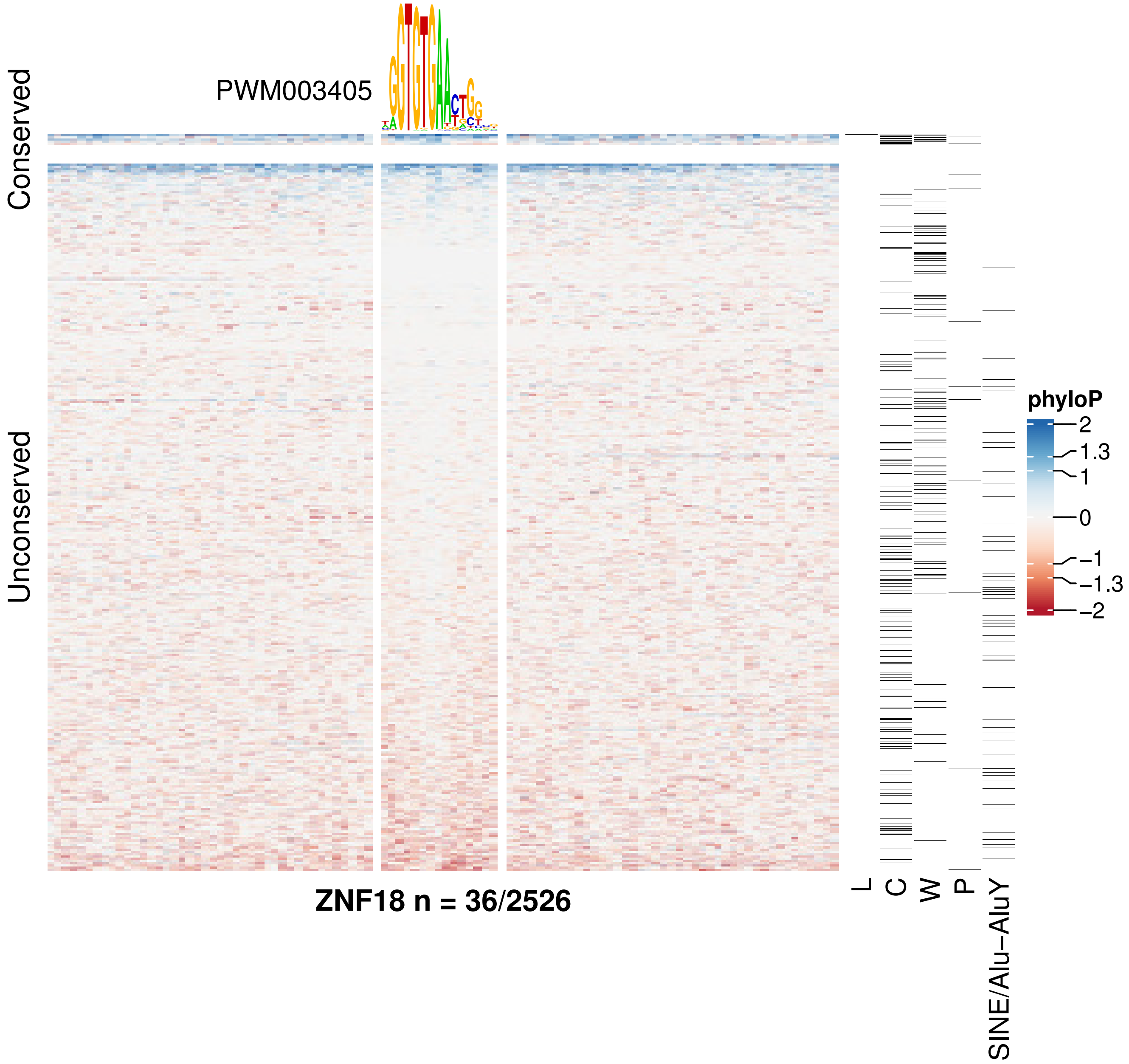

PWM003405
Conserved
Unconserved
ZNF18 n = 36/2526
L
C
W
P
SINE/Alu−AluY
phyloP
2
1.3
1
0
−1
−1.3
−2

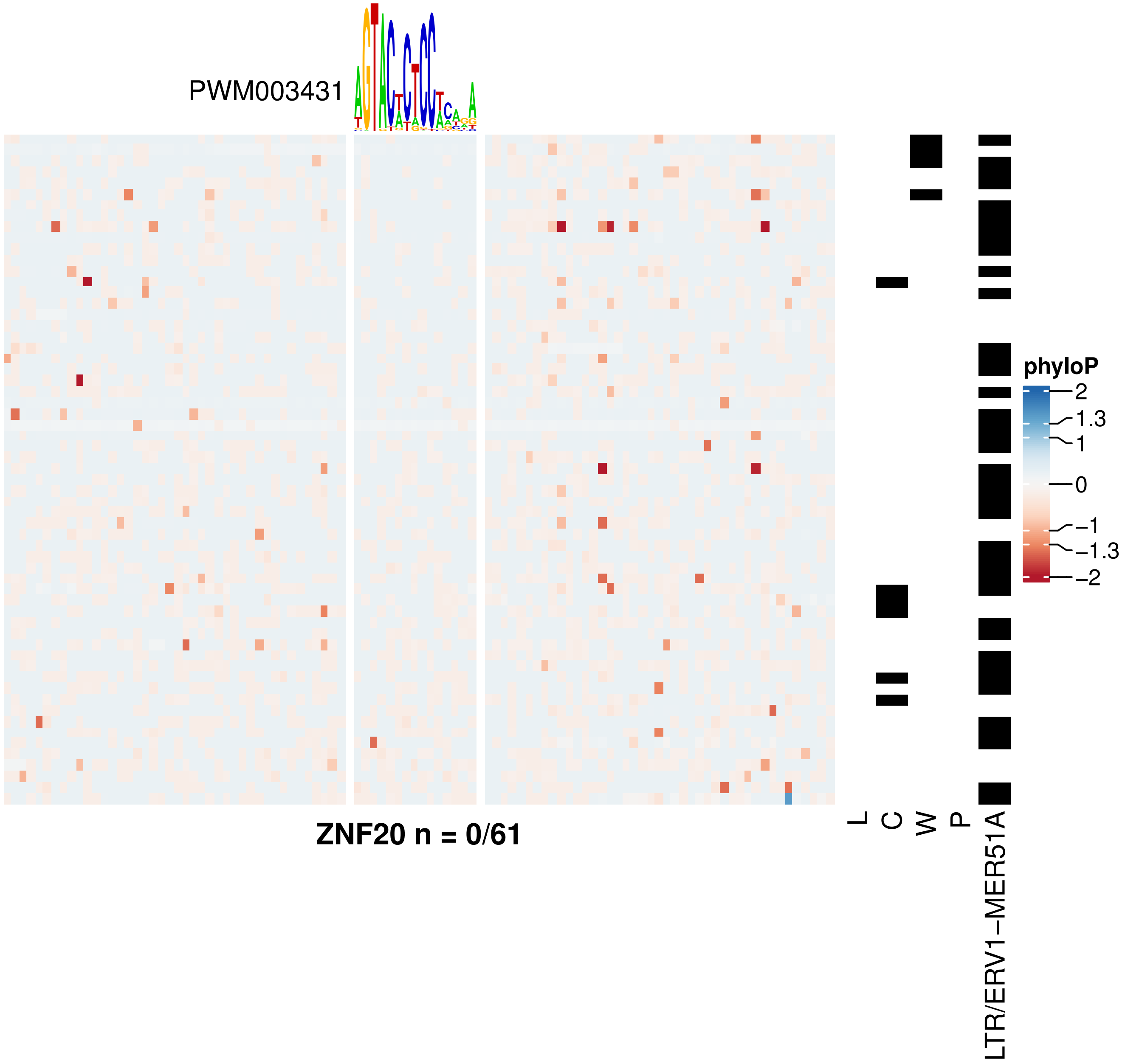
PWM003431
ZNF20 n = 0/61
L
C
W
P
LTR/ERV1−MER51A
phyloP
2
1.3
1
0
−1
−1.3
−2

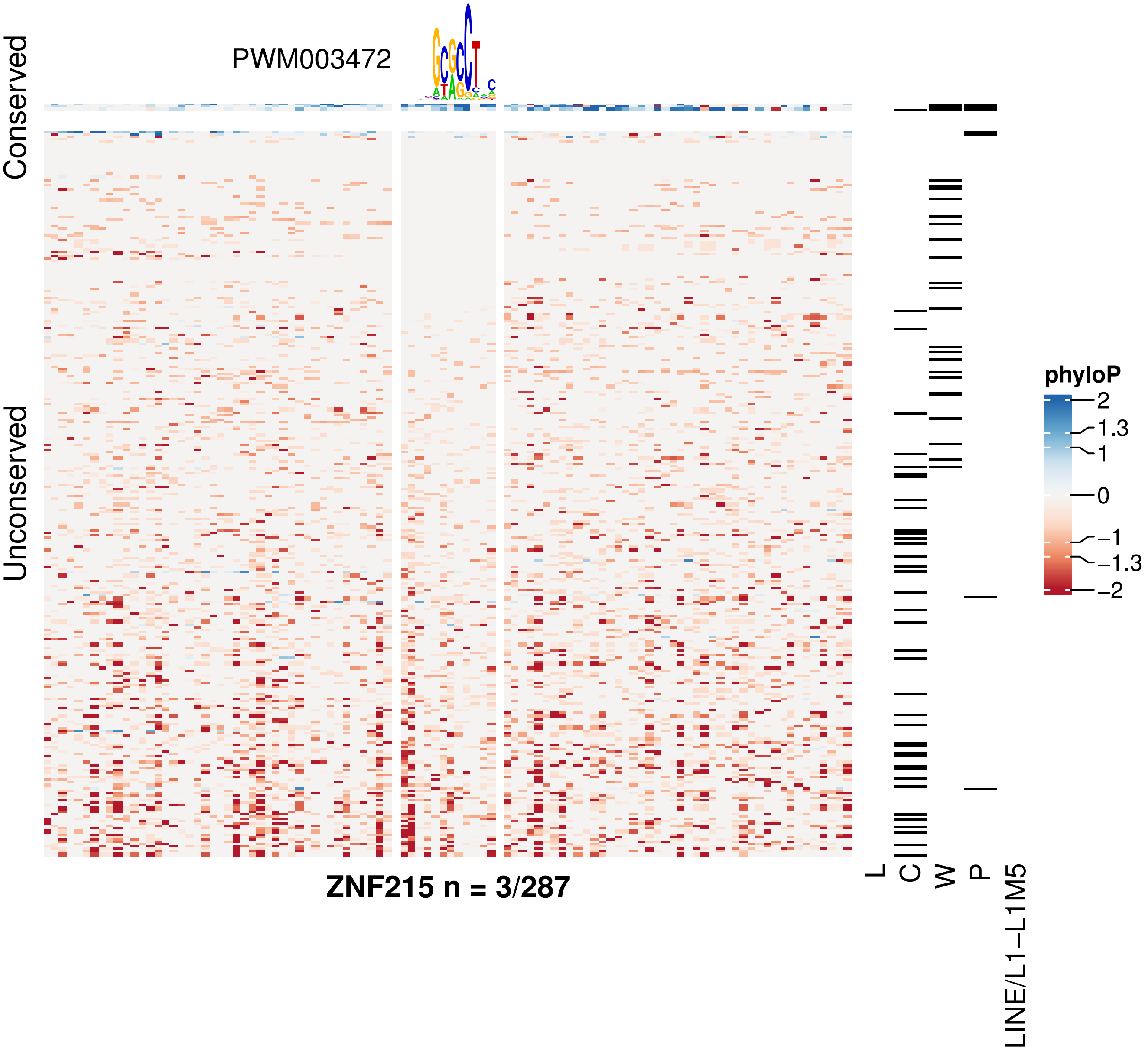
PWM003472
Conserved
Unconserved
ZNF215 n = 3/287
L
C
W
P
LINE/L1−L1M5
phyloP
2
1.3
1
0
−1
−1.3
−2

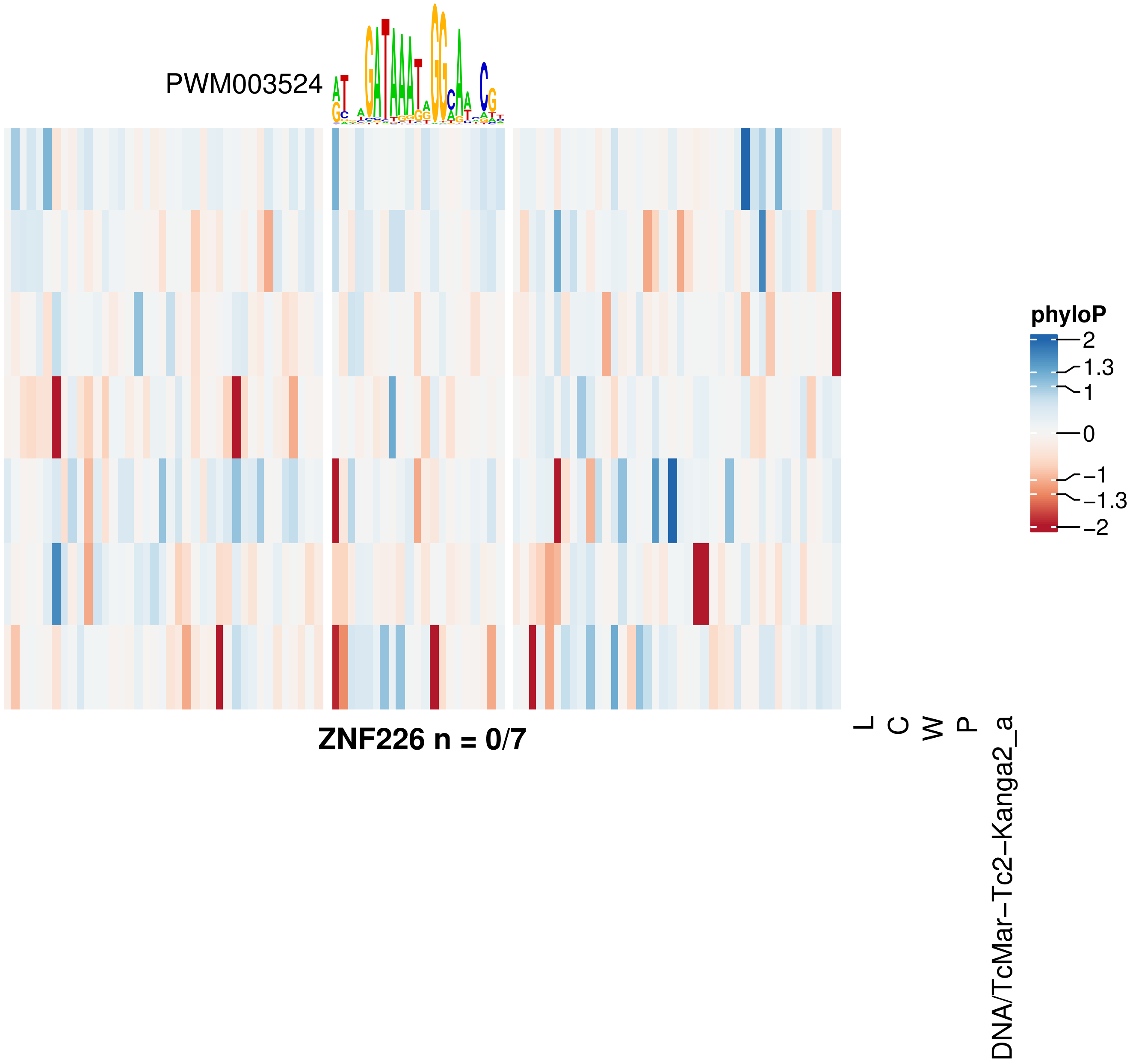


**ZNF226 n = 0/7**

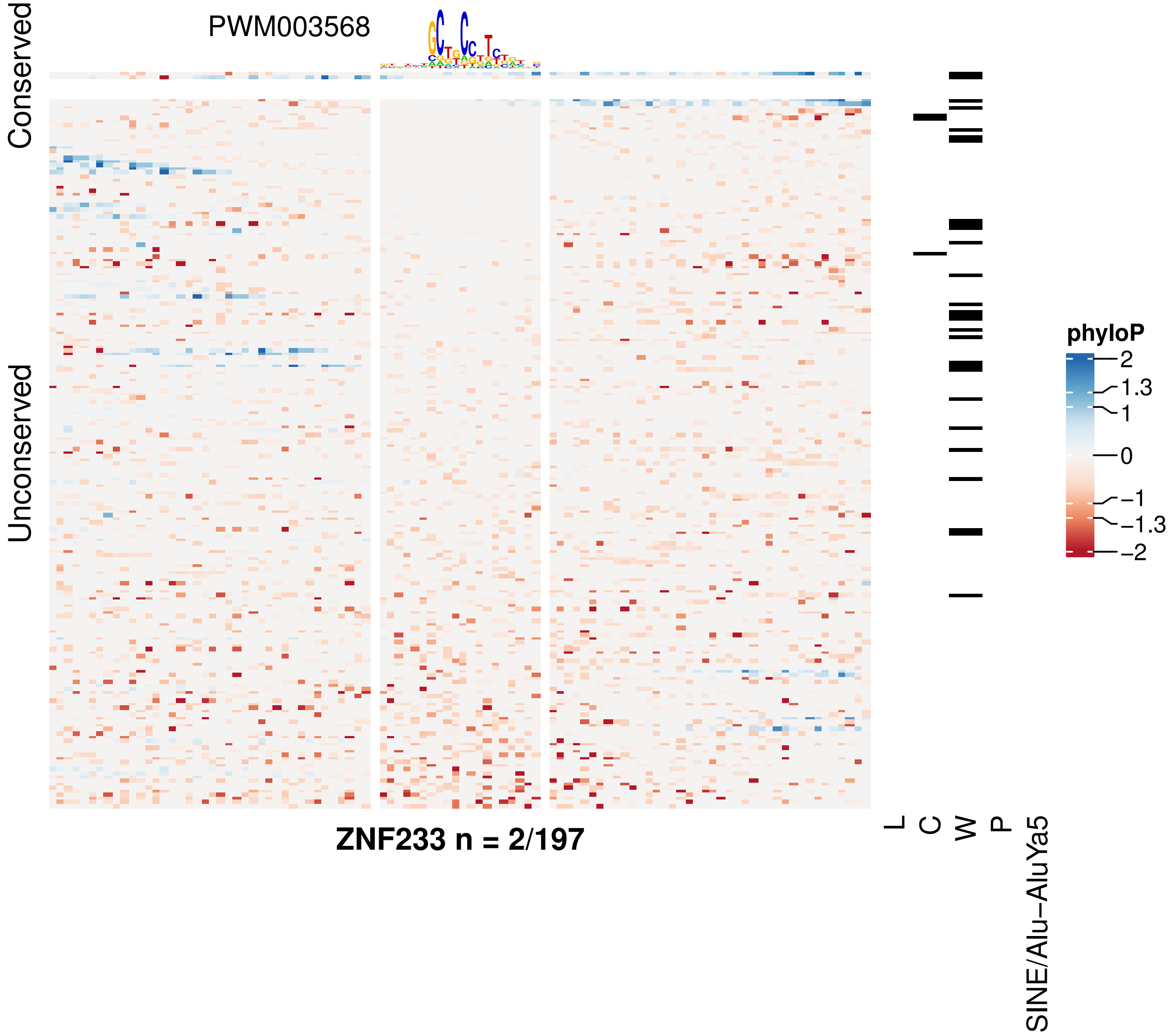

Conserved
Unconserved
PWM003568
ZNF233 n = 2/197
L
C
W
P
SINE/Alu−AluYa5
phyloP
2
1.3
1
0
−1
−1.3
−2

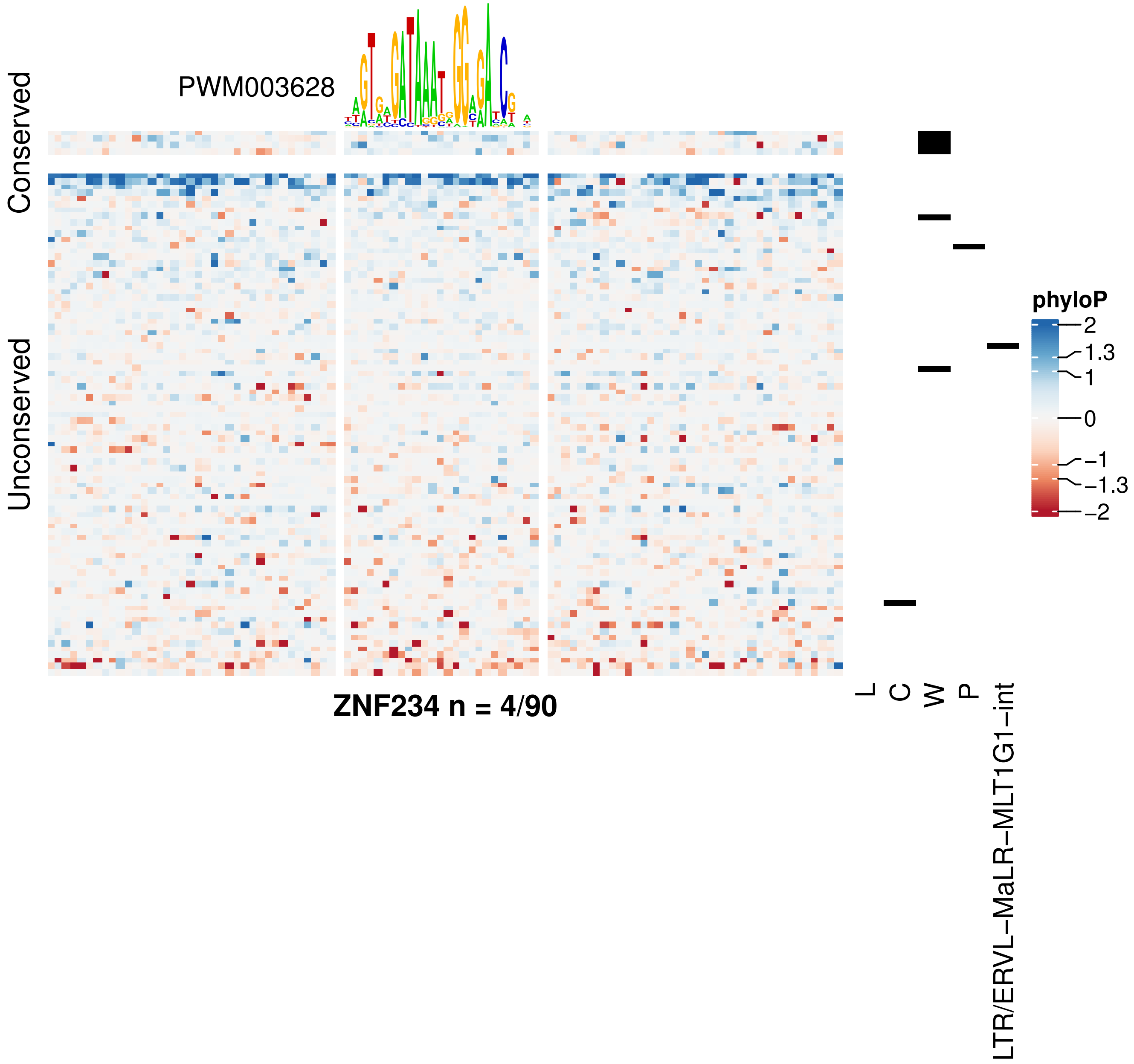

PWM003628
Conserved
Unconserved
phyloP
2
1.3
1
0
−1
−1.3
−2
ZNF234 n = 4/90
L
C
W
P
LTR/ERVL−MaLR−MLT1G1−int

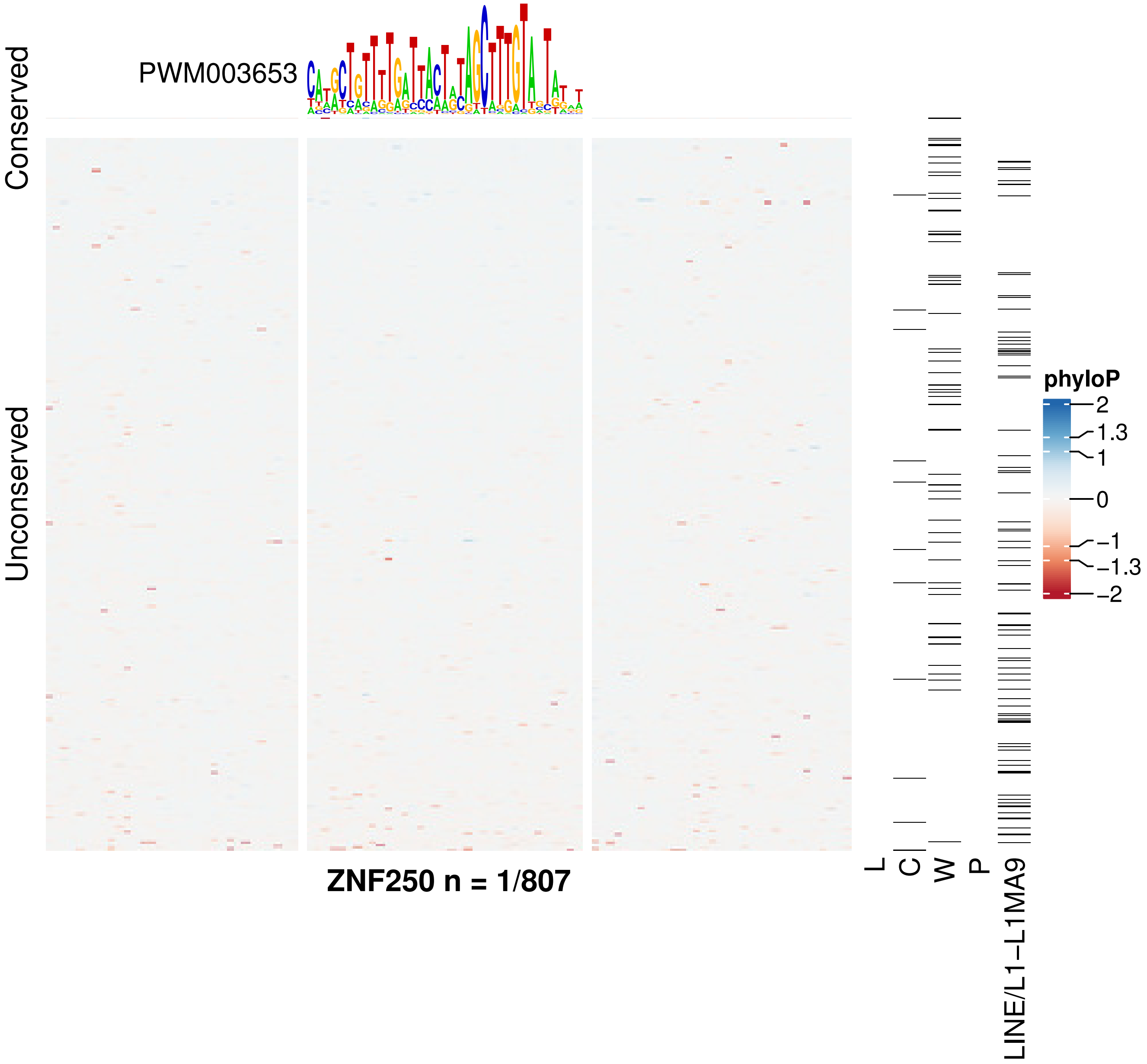

PWM003653
Conserved
Unconserved
ZNF250 n = 1/807
L
C
W
P
LINE/L1−L1MA9
phyloP
2
1.3
1
0
−1
−1.3
−2

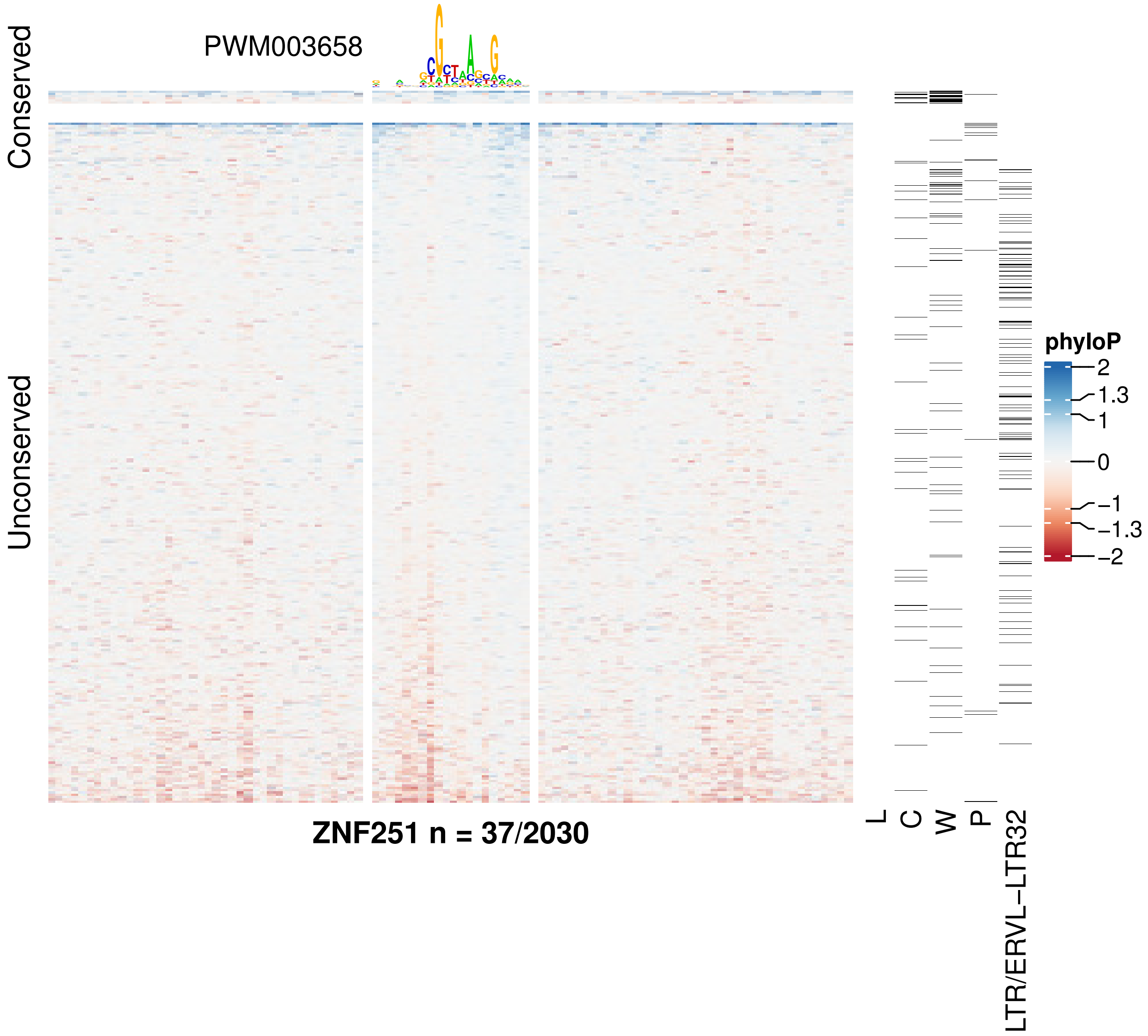

PWM003658
Conserved
Unconserved
ZNF251 n = 37/2030
L
C
W
P
LTR/ERVL–LTR32
phyloP
2
1.3
1
0
−1
−1.3
−2

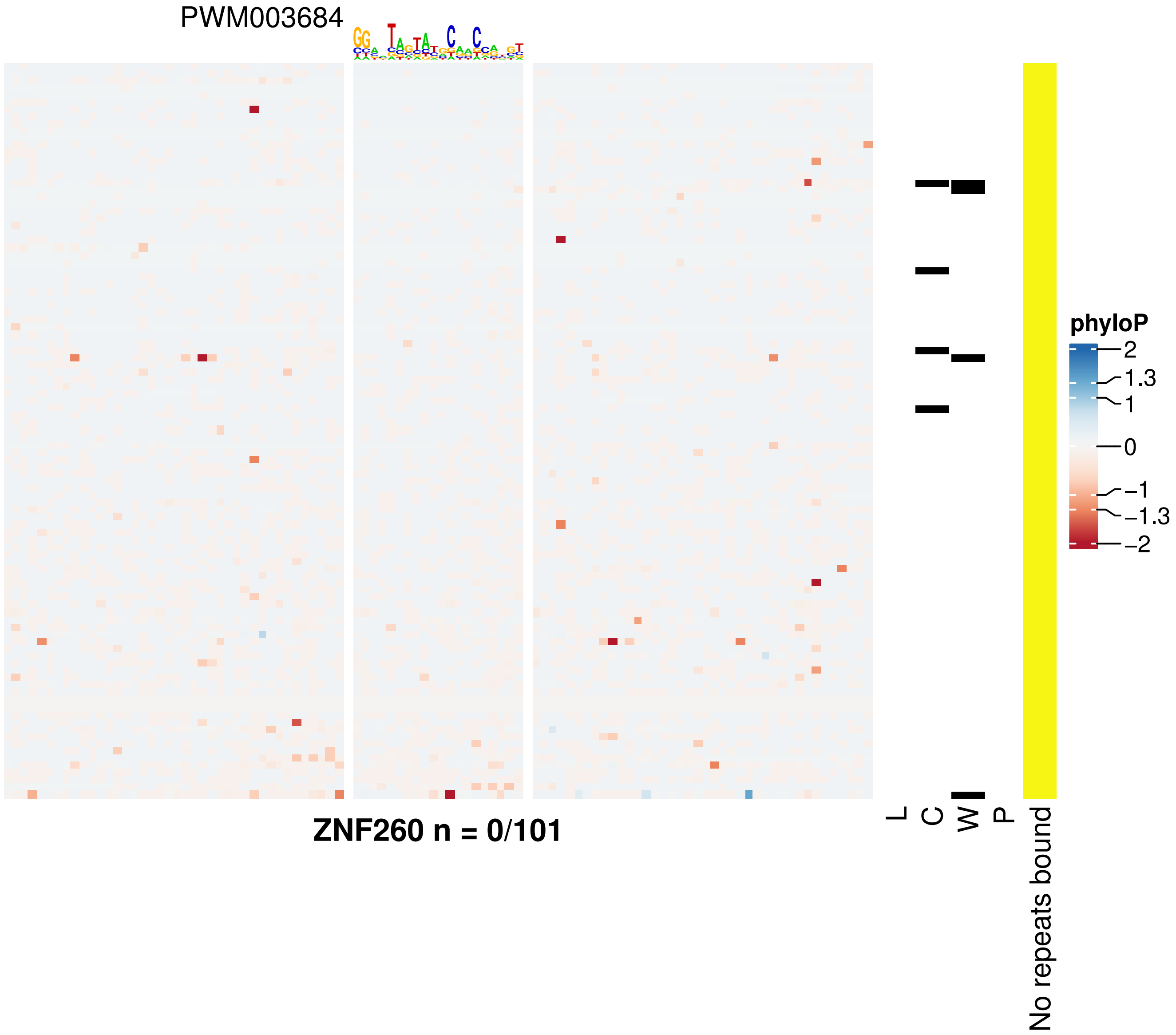
PWM003684
phyloP
2
1.3
1
0
−1
−1.3
−2
L
C
W
P
No repeats bound
ZNF260 n = 0/101

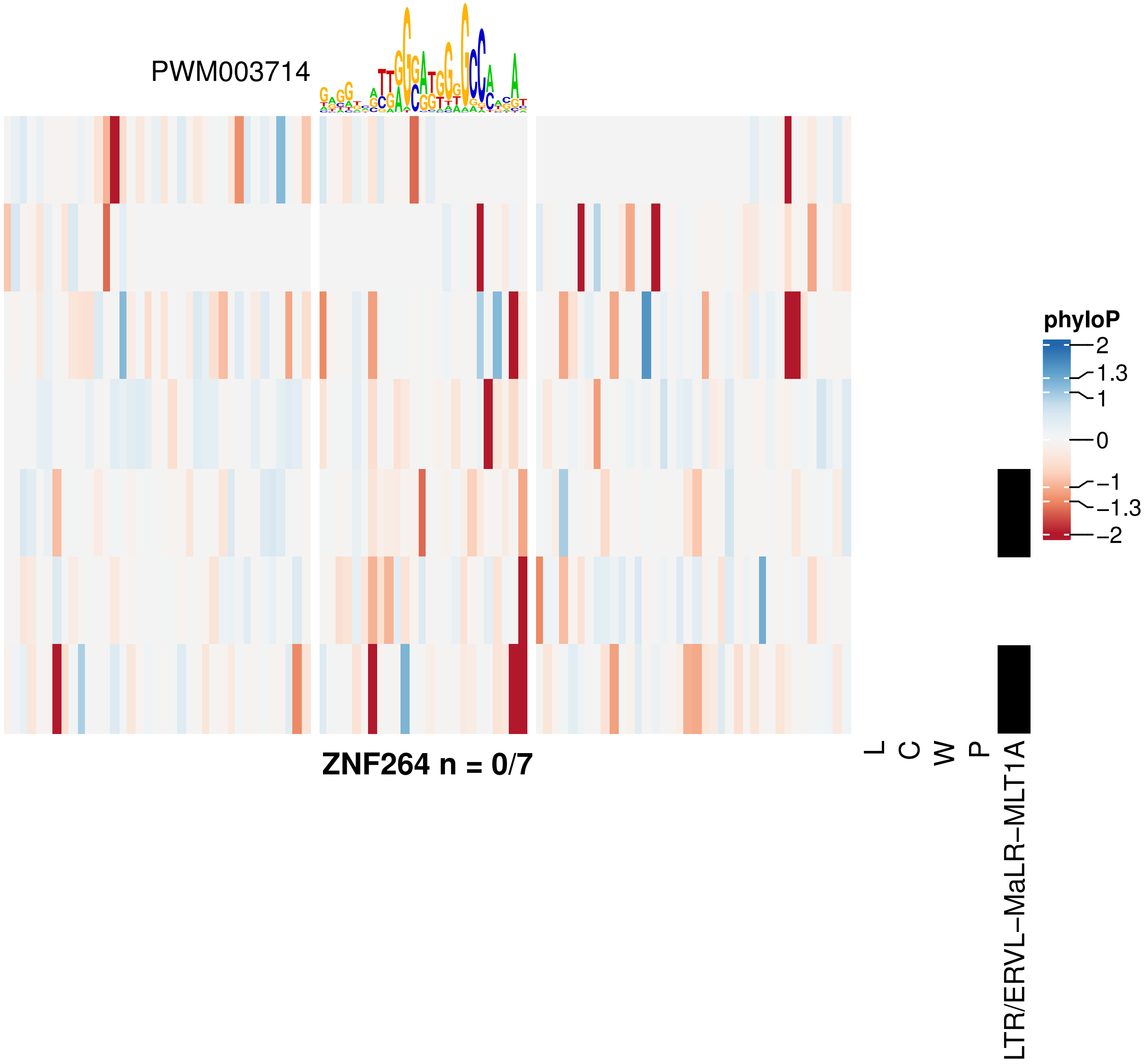
PWM003714
phyloP
2
1.3
1
0
−1
−1.3
−2
ZNF264 n = 0/7
L
C
W
P
LTR/ERVL−MaLR−MLT1A

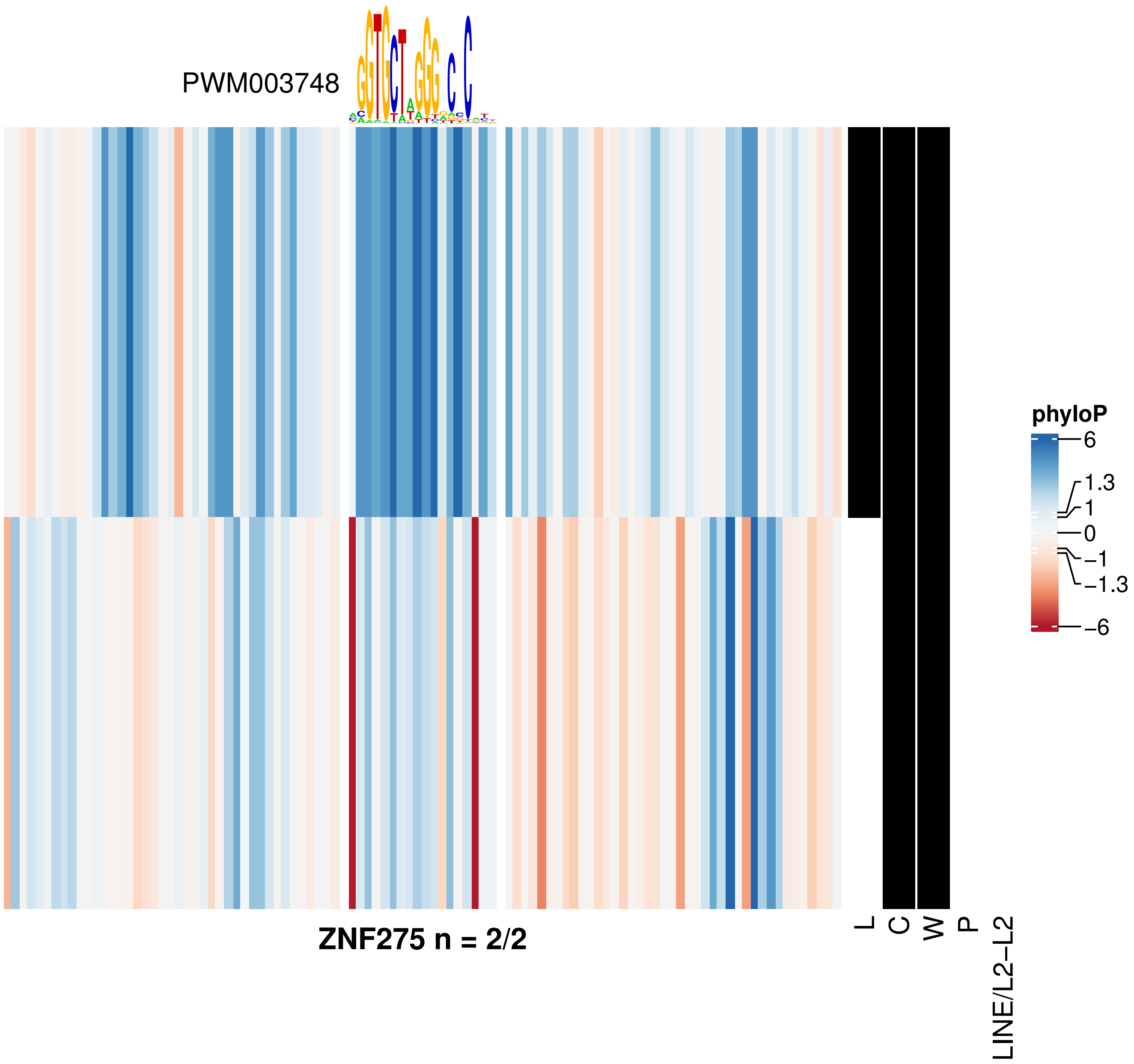
PWM003748
phyloP
6
1.3
1
0
−1
−1.3
−6
ZNF275 n = 2/2
L
C
W
P
LINE/L2−L2

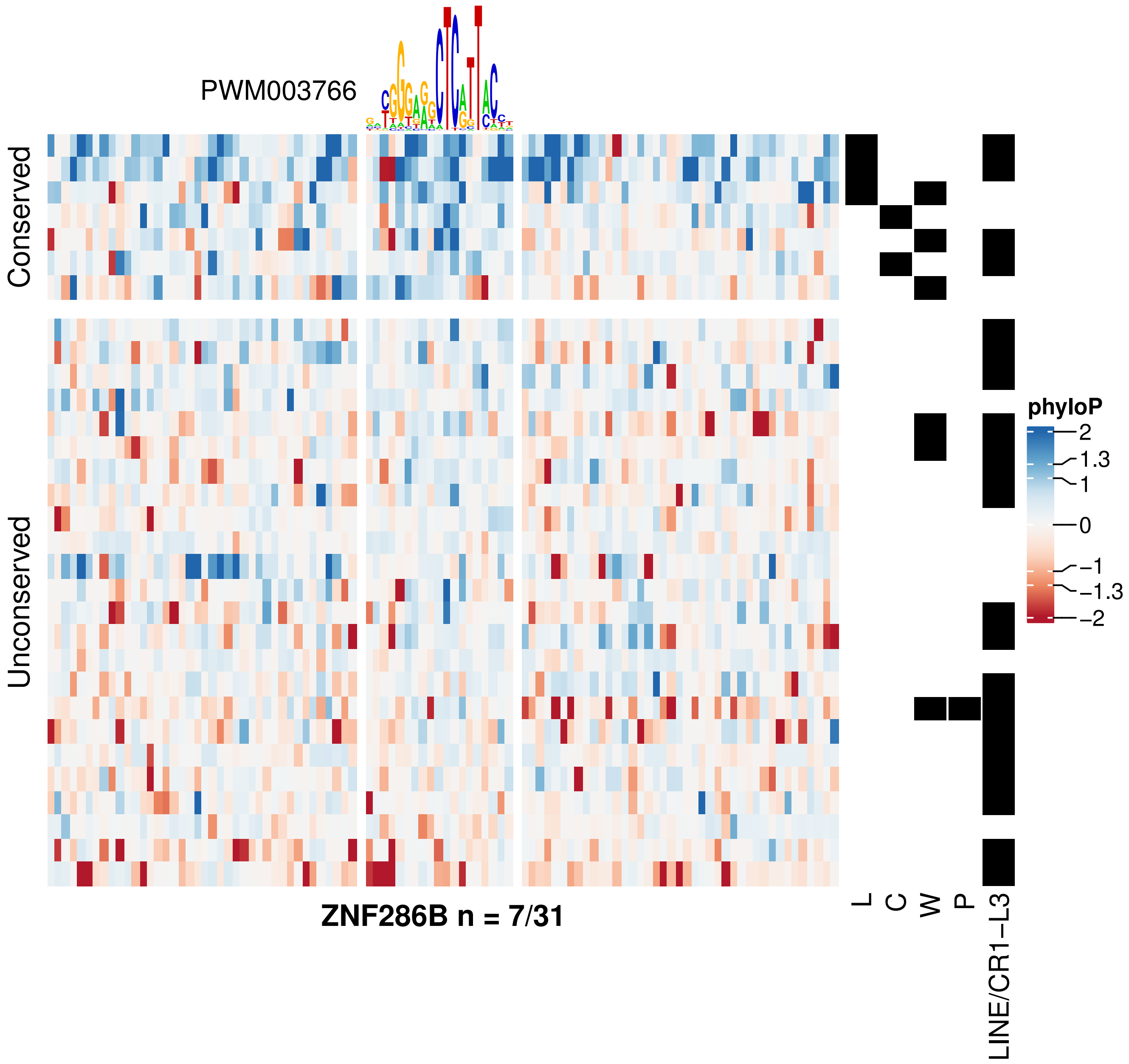
PWM003766
Conserved
Unconserved
ZNF286B n = 7/31
L
C
W
P
LINE/CR1−L3
phyloP
2
1.3
1
0
−1
−1.3
−2

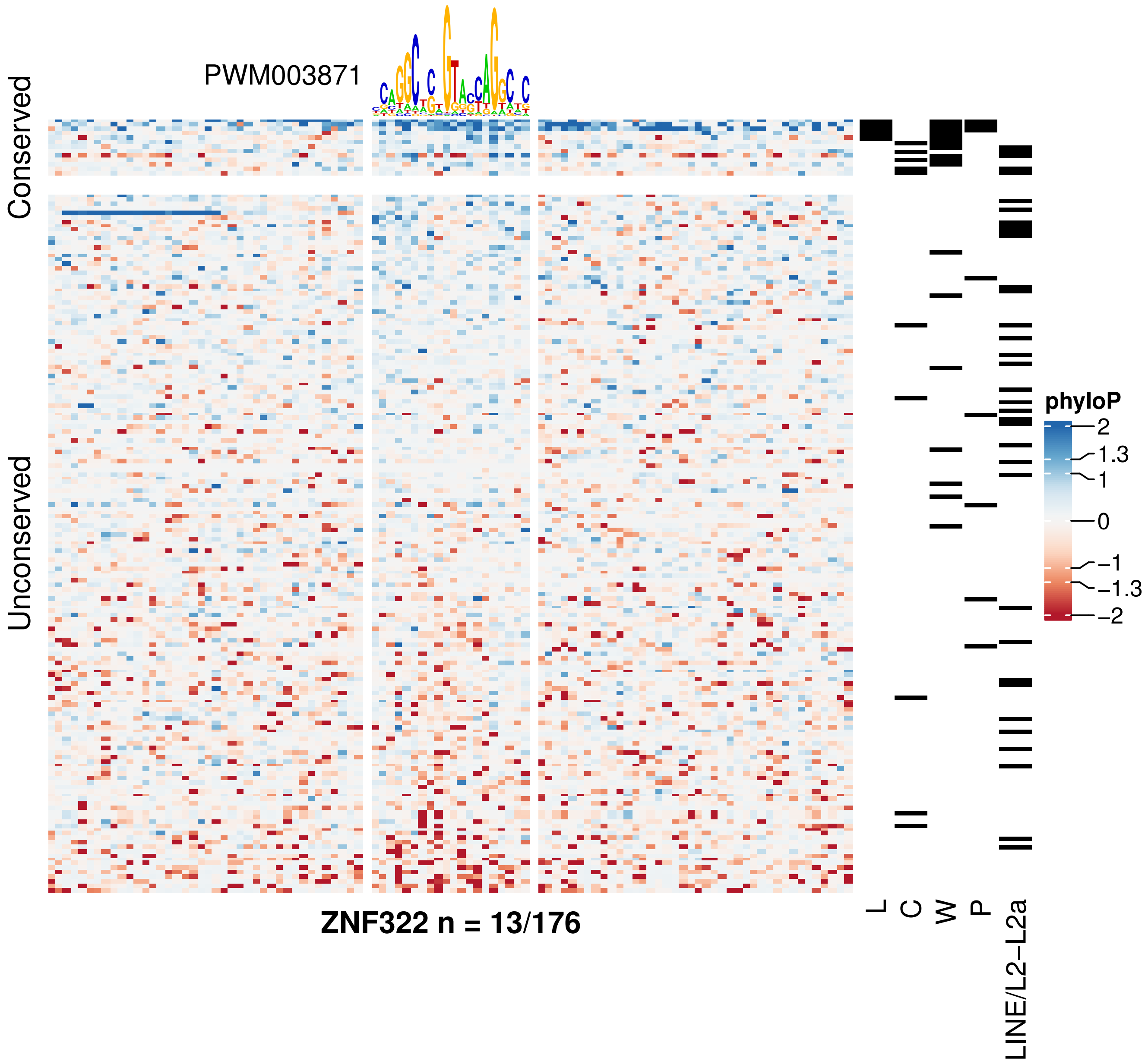
PWM003871
Conserved
Unconserved
phyloP
2
1.3
1
0
−1
−1.3
−2
L
C
W
P
LINE/L2−L2a
ZNF322 n = 13/176

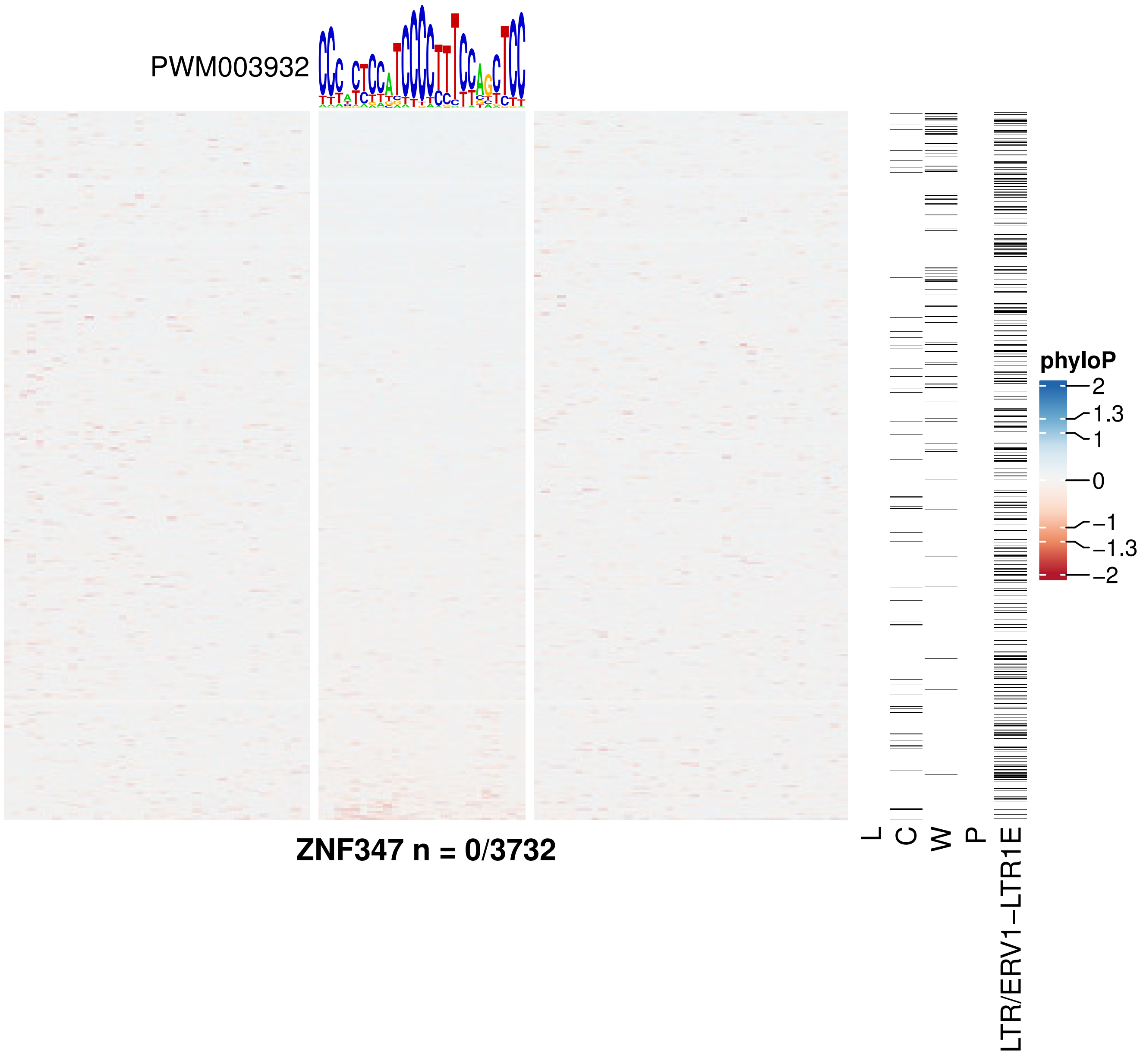
PWM003932
ZNF347 n = 0/3732
L
C
W
P
LTR/ERV1−LTR1E
phyloP
2
1.3
1
0
−1
−1.3
−2

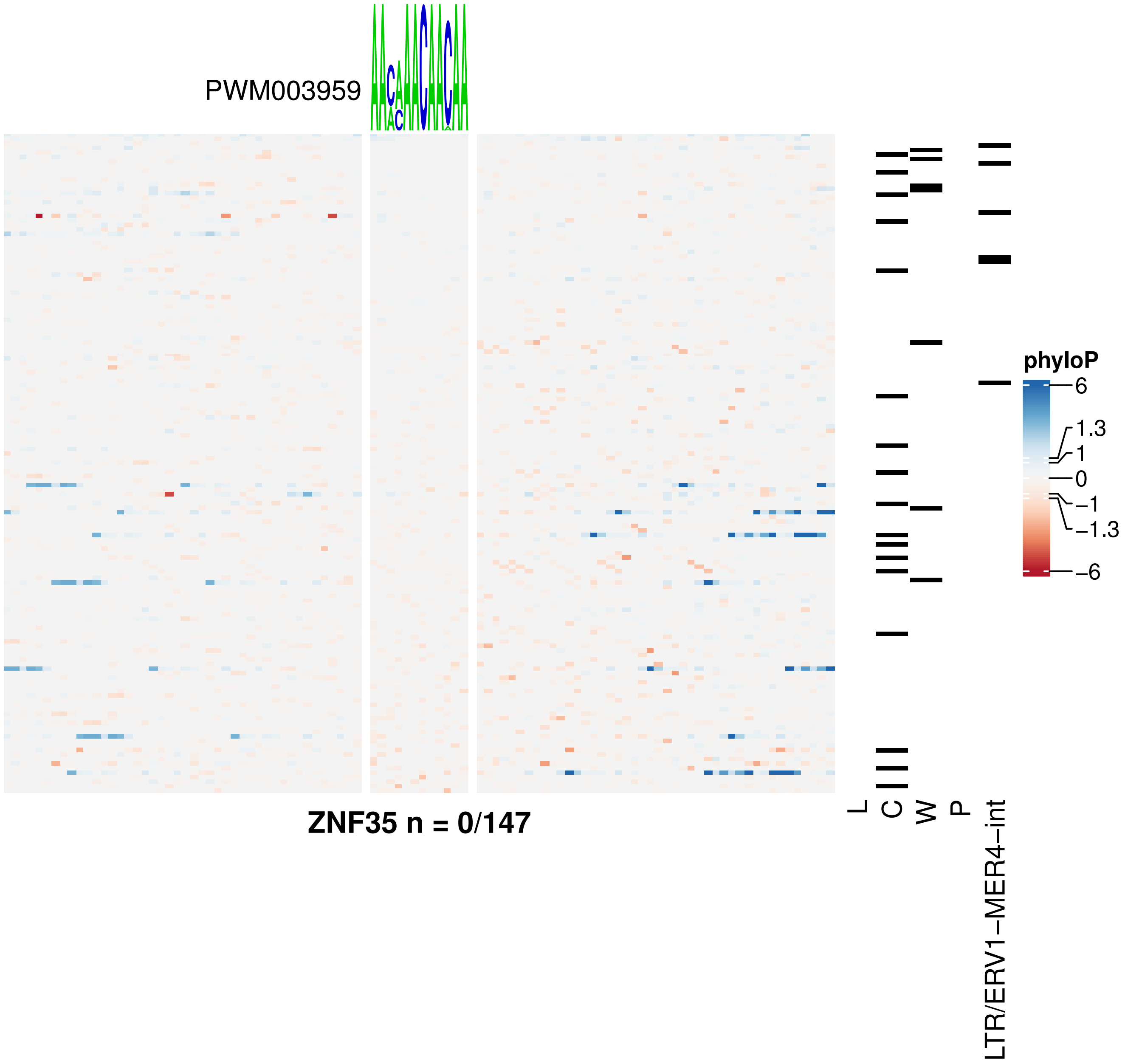
PWM003959
ZNF35 n = 0/147
L
C
W
P
LTR/ERV1−MER4−int
phyloP
6
1.3
1
0
−1
−1.3
−6

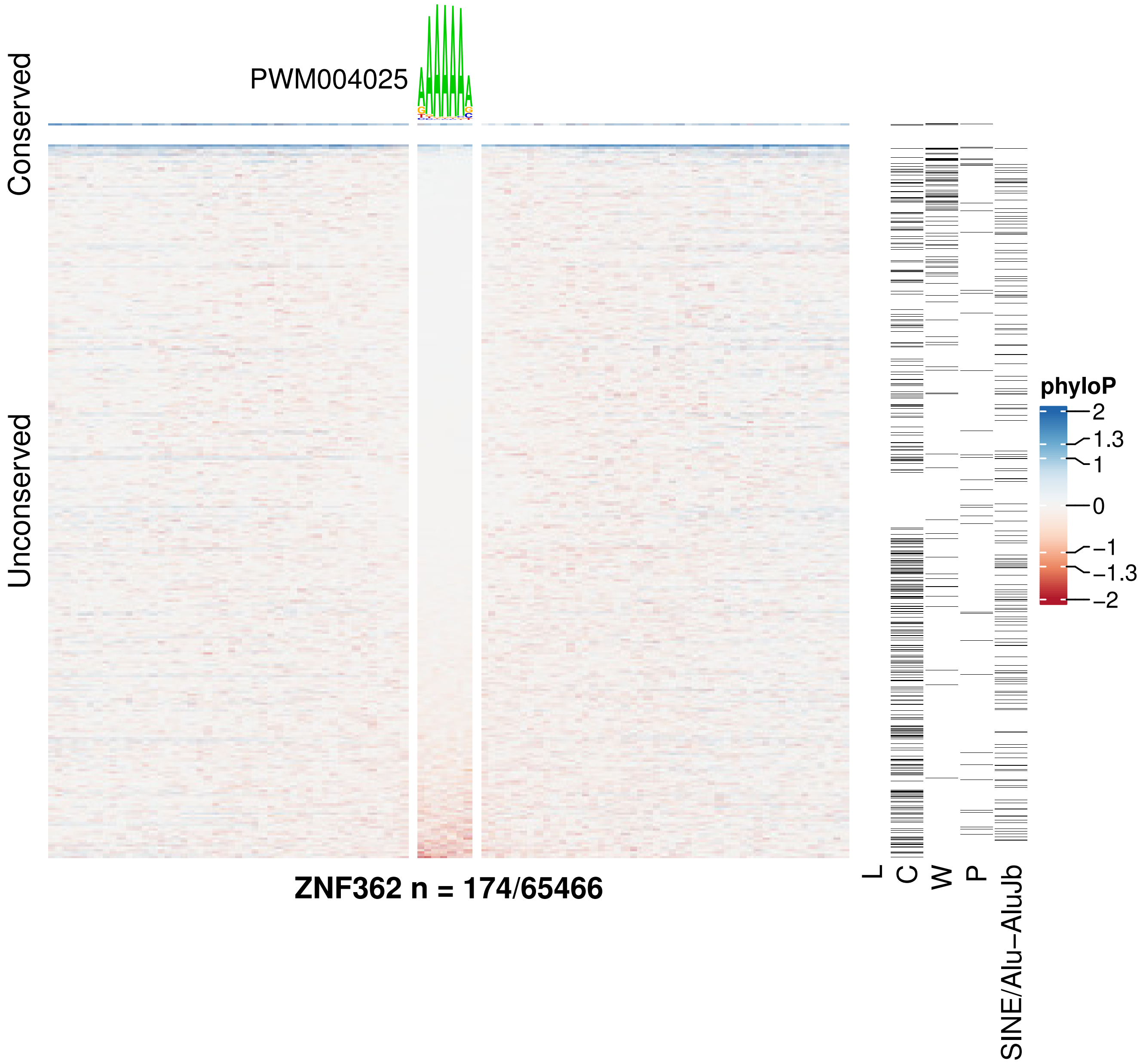
PWM004025
Conserved
Unconserved
ZNF362 n = 174/65466
L
C
W
P
SINE/Alu−AluJb
phyloP
2
1.3
1
0
−1
−1.3
−2

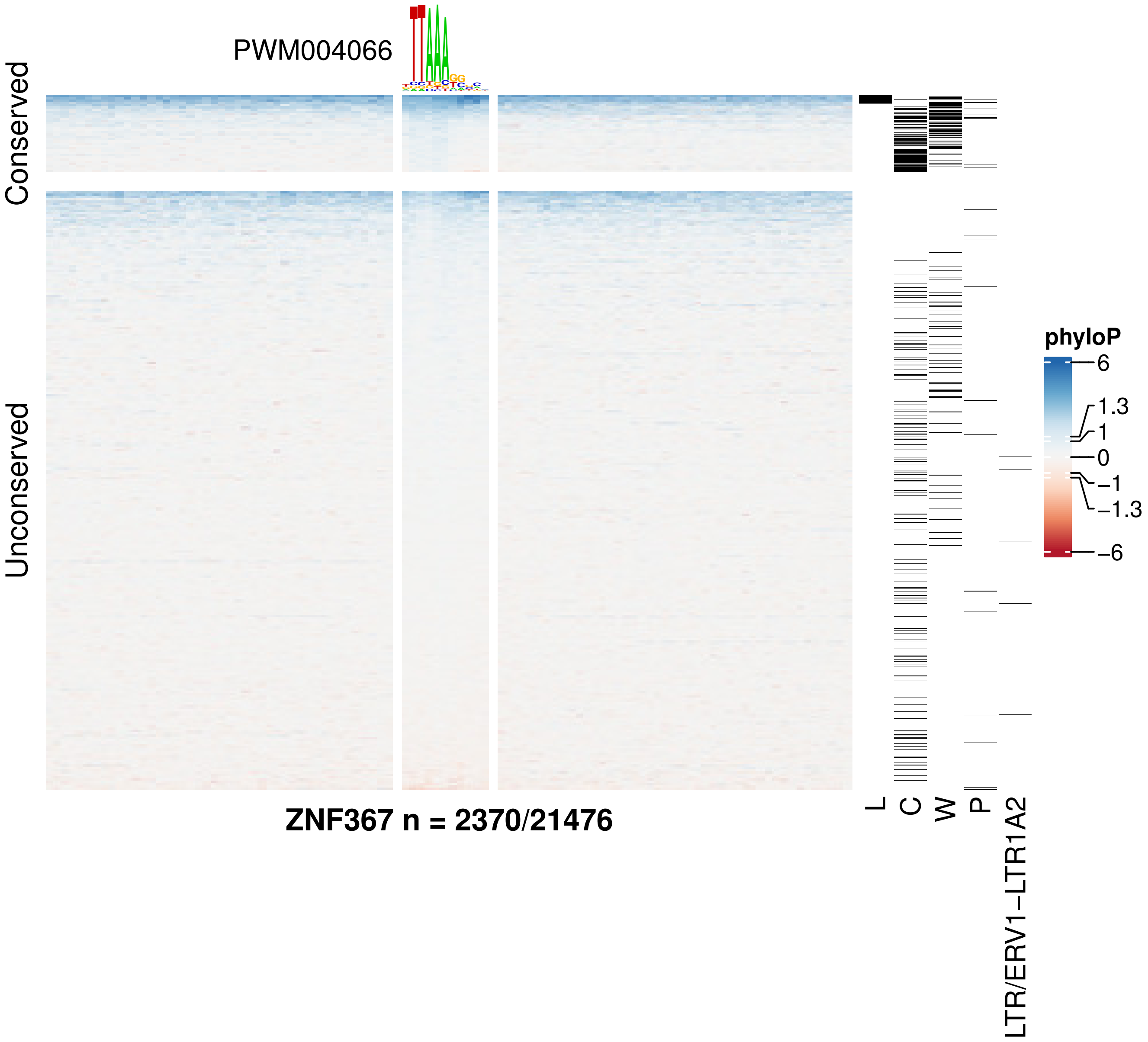
PWM004066
Conserved
Unconserved
ZNF367 n = 2370/21476
L
C
W
P
LTR/ERV1−LTR1A2
phyloP
6
1.3
1
0
−1
−1.3
−6

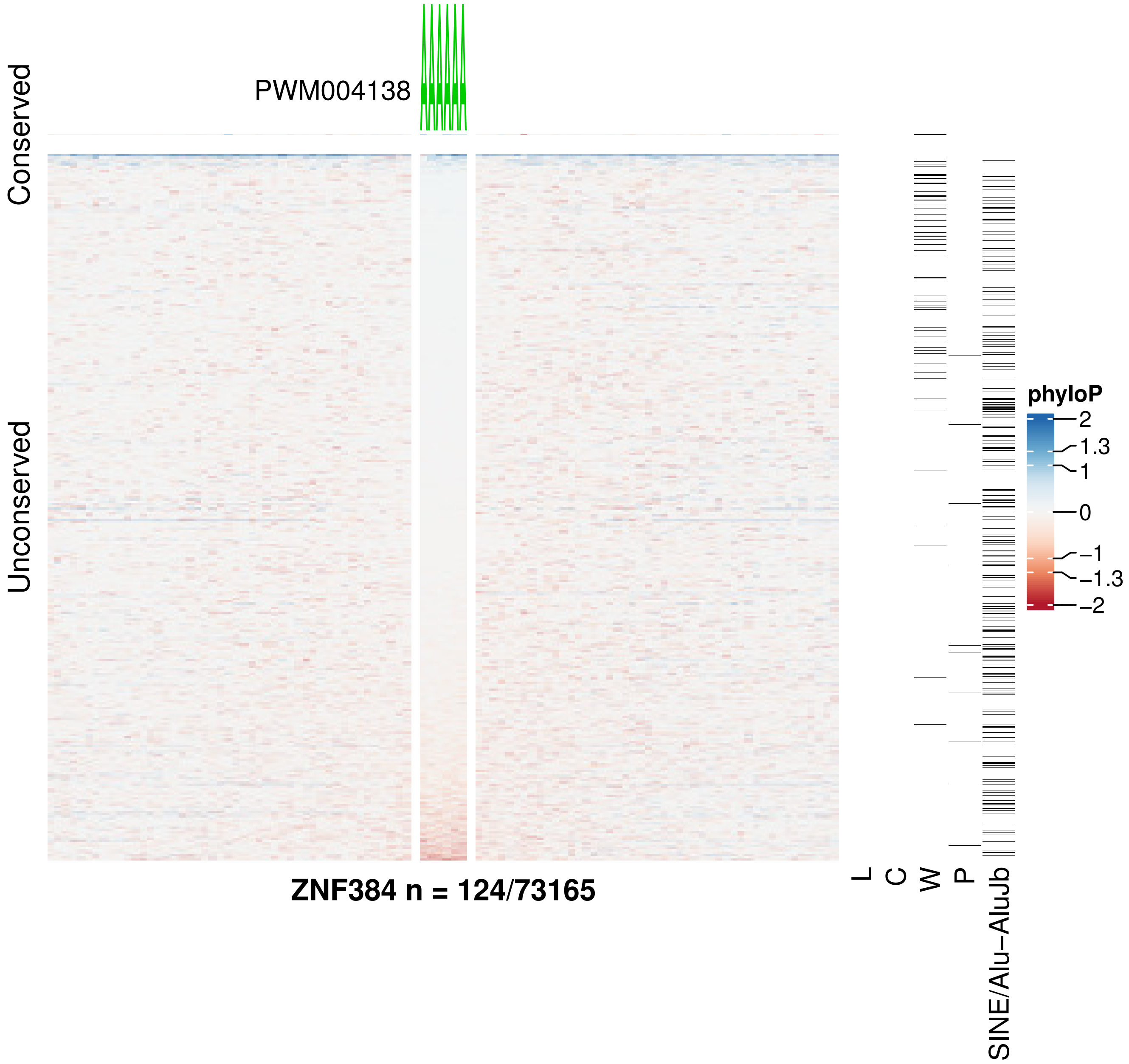
PWM004138
Conserved
Unconserved
ZNF384 n = 124/73165
L
C
W
P
SINE/Alu−AluJb
phyloP
2
1.3
1
0
−1
−1.3
−2

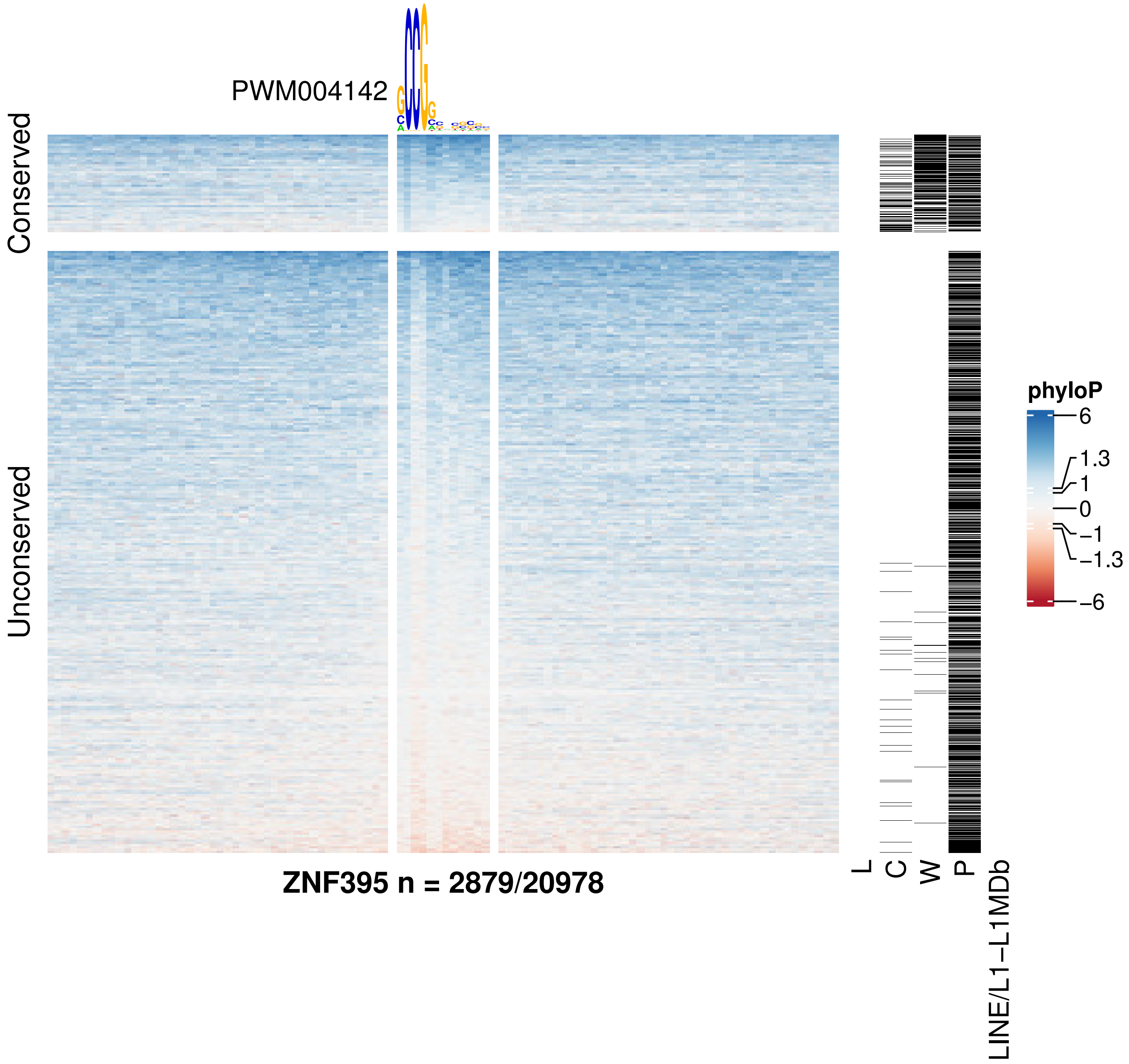
PWM004142
Conserved
Unconserved
ZNF395 n = 2879/20978
L
C
W
P
LINE/L1−L1MDb
phyloP
6
1.3
1
0
−1
−1.3
−6

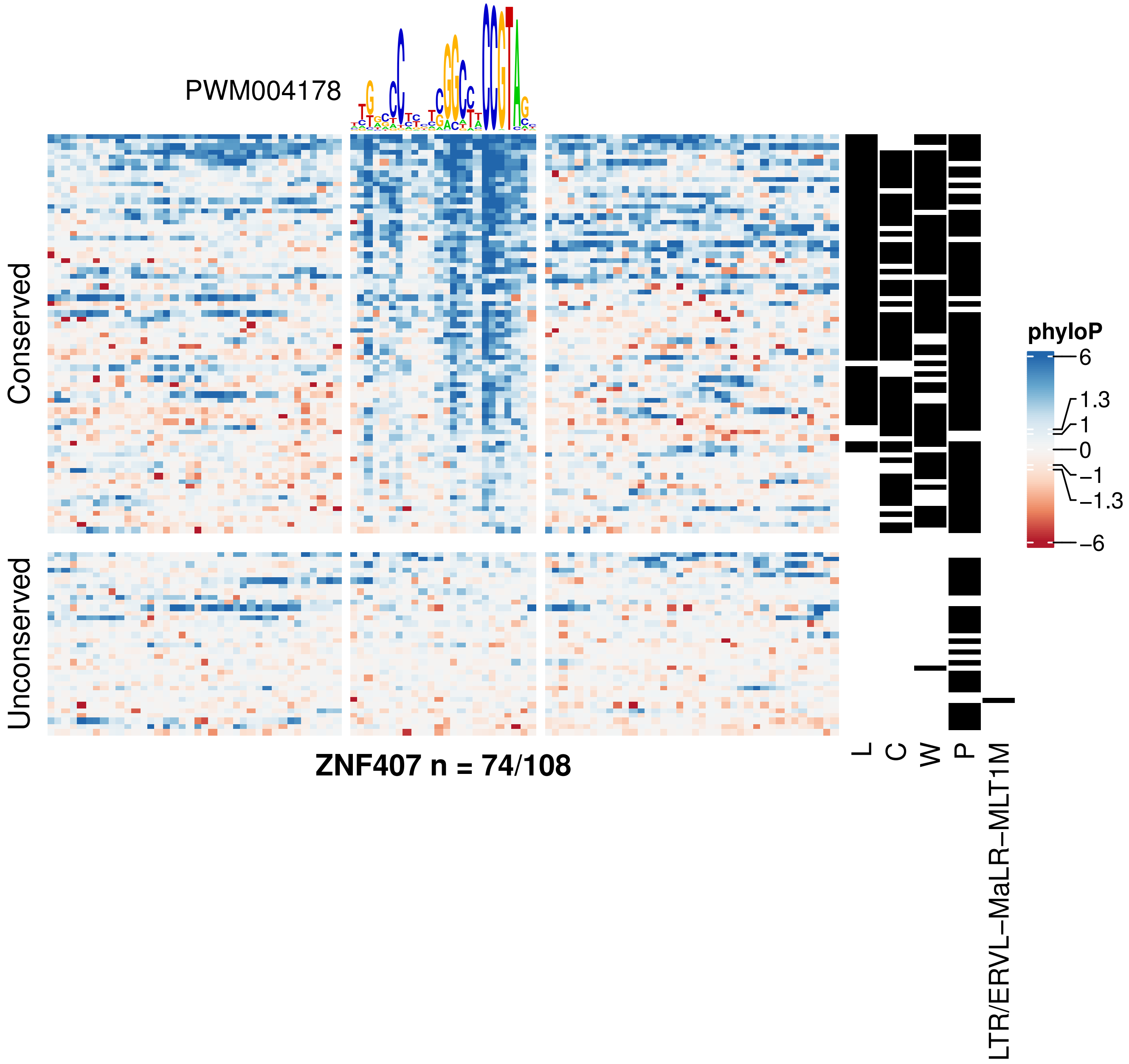
PWM004178
Conserved
Unconserved
ZNF407 n = 74/108
L
C
W
P
LTR/ERVL−MaLR−MLT1M
phyloP
6
1.3
1
0
−1
−1.3
−6

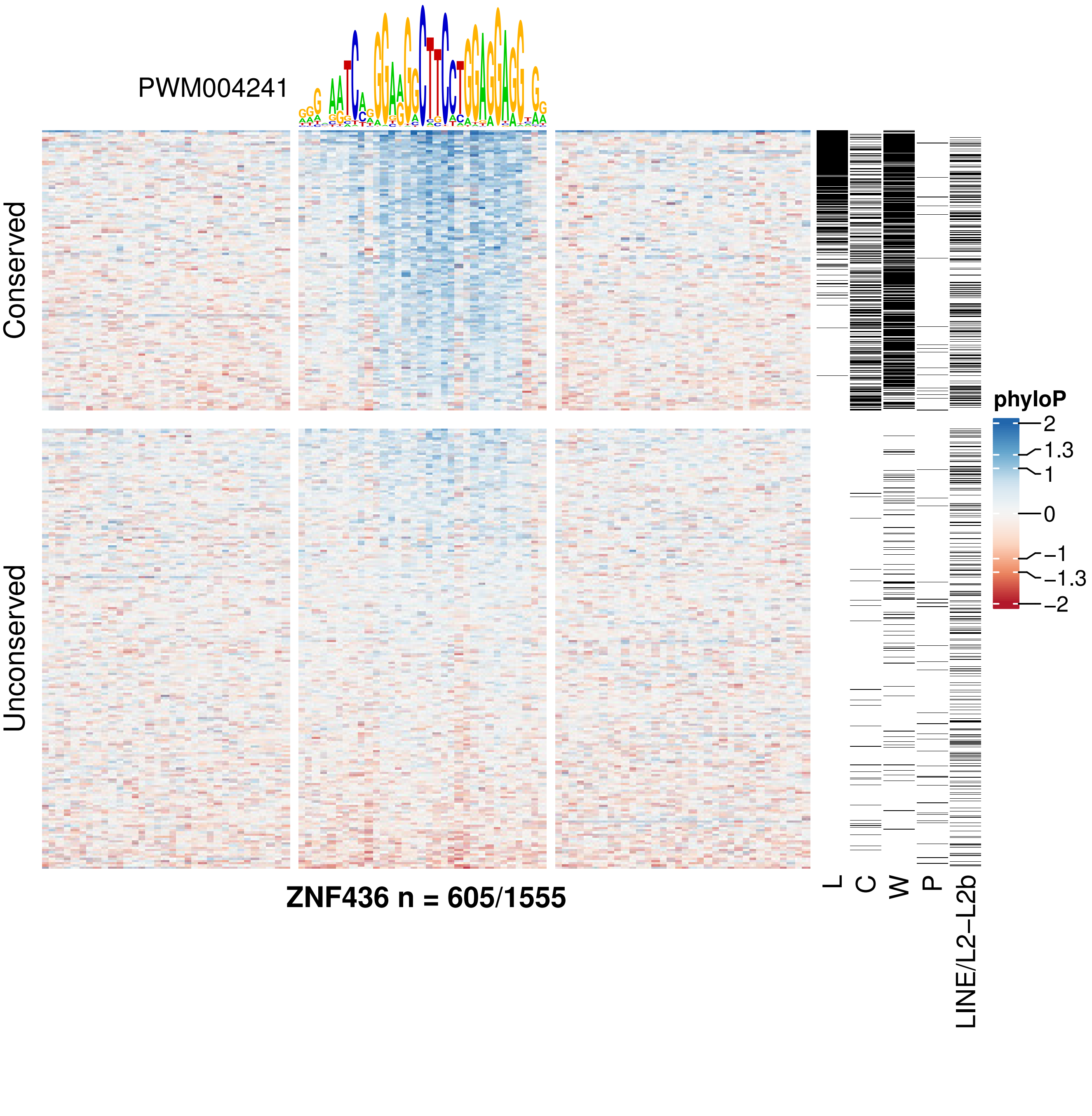
PWM004241
Conserved
Unconserved
phyloP
2
1.3
1
0
−1
−1.3
−2
L
C
W
P
LINE/L2−L2b
ZNF436 n = 605/1555

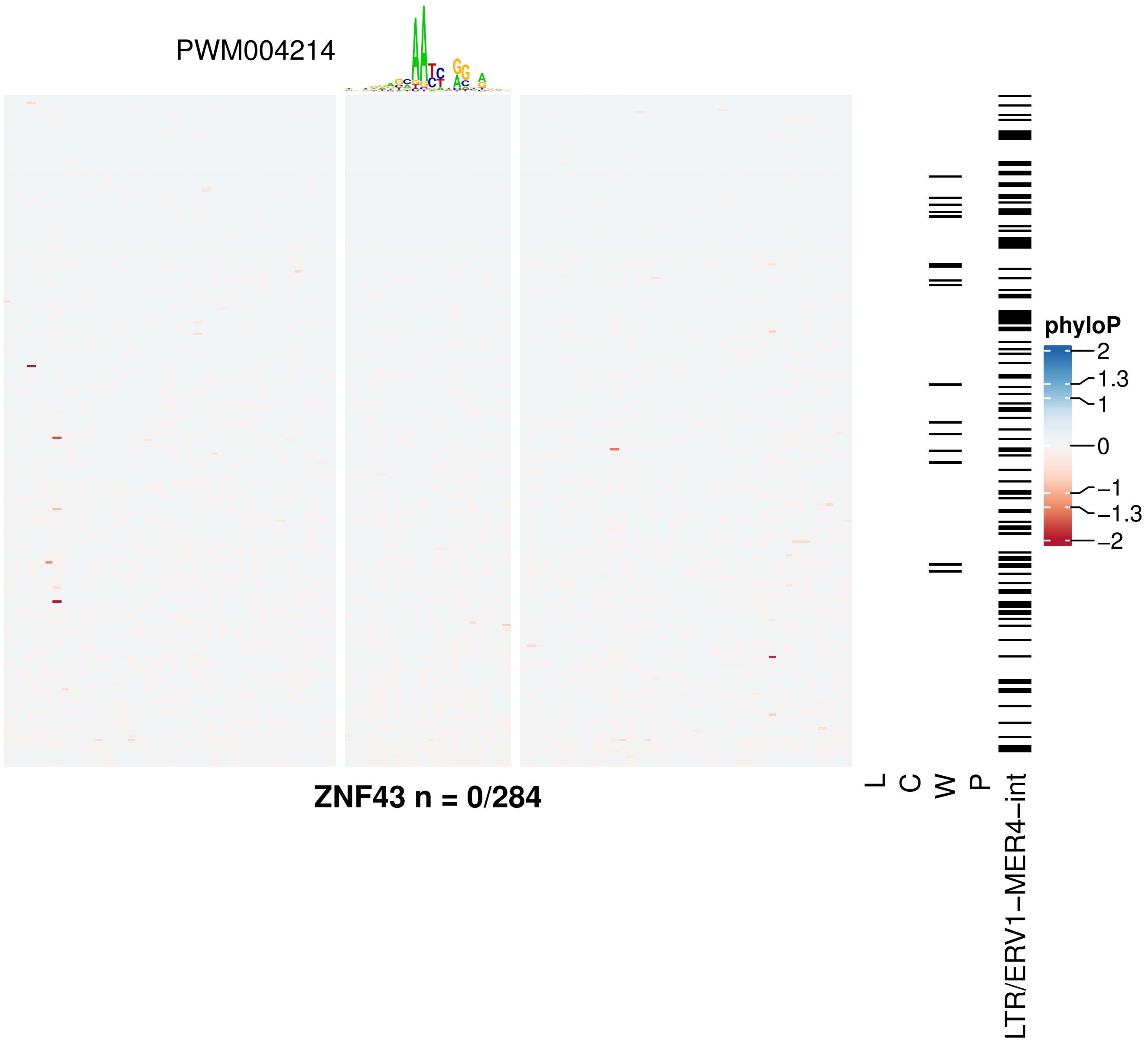
PWM004214
ZNF43 n = 0/284
L
C
W
P
LTR/ERV1−MER4−int
phyloP
2
1.3
1
0
−1
−1.3
−2

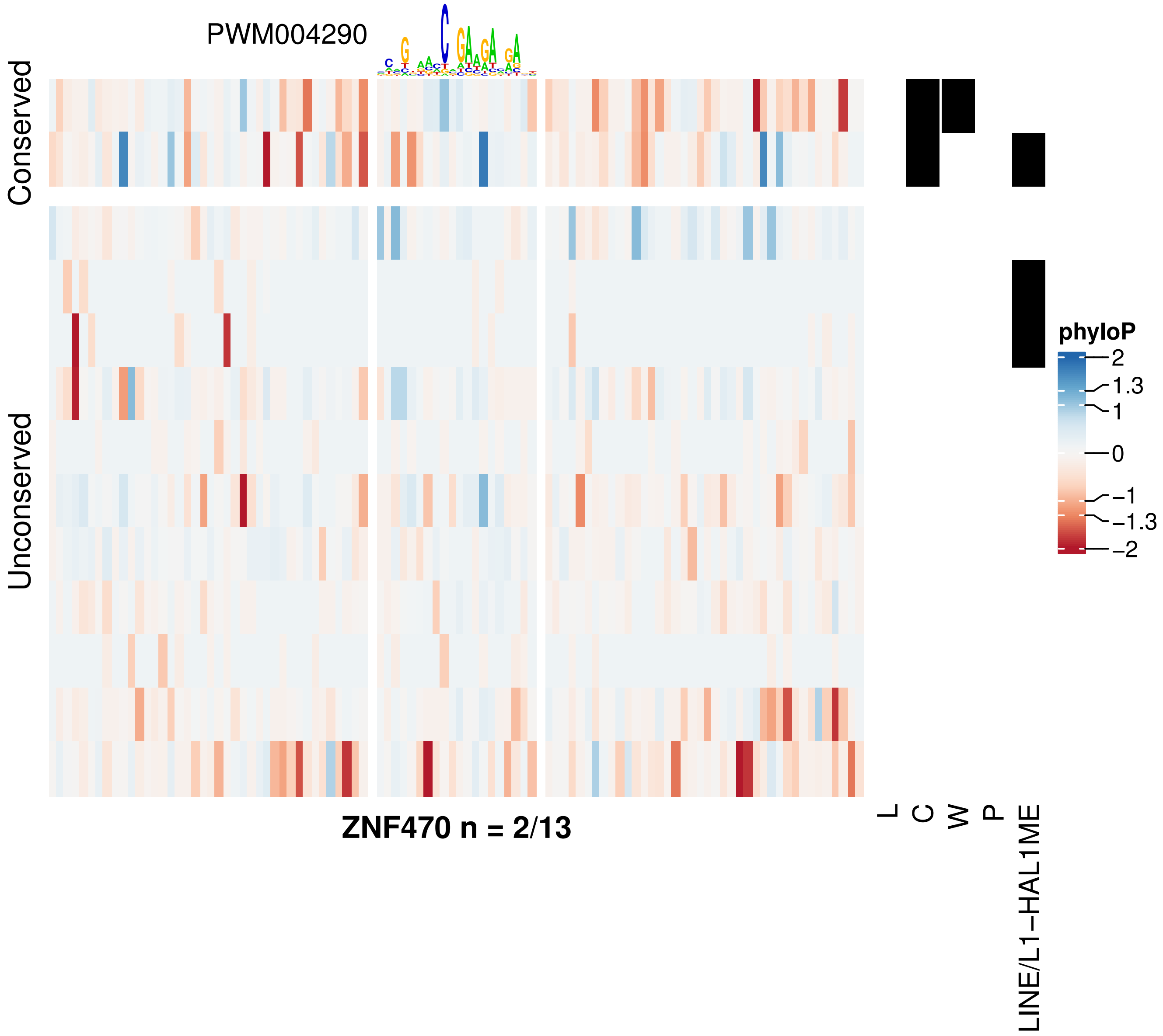

PWM004290
Conserved
Unconserved
ZNF470 n = 2/13
L
C
W
P
LINE/L1−HAL1ME
phyloP
2
1.3
1
0
−1
−1.3
−2

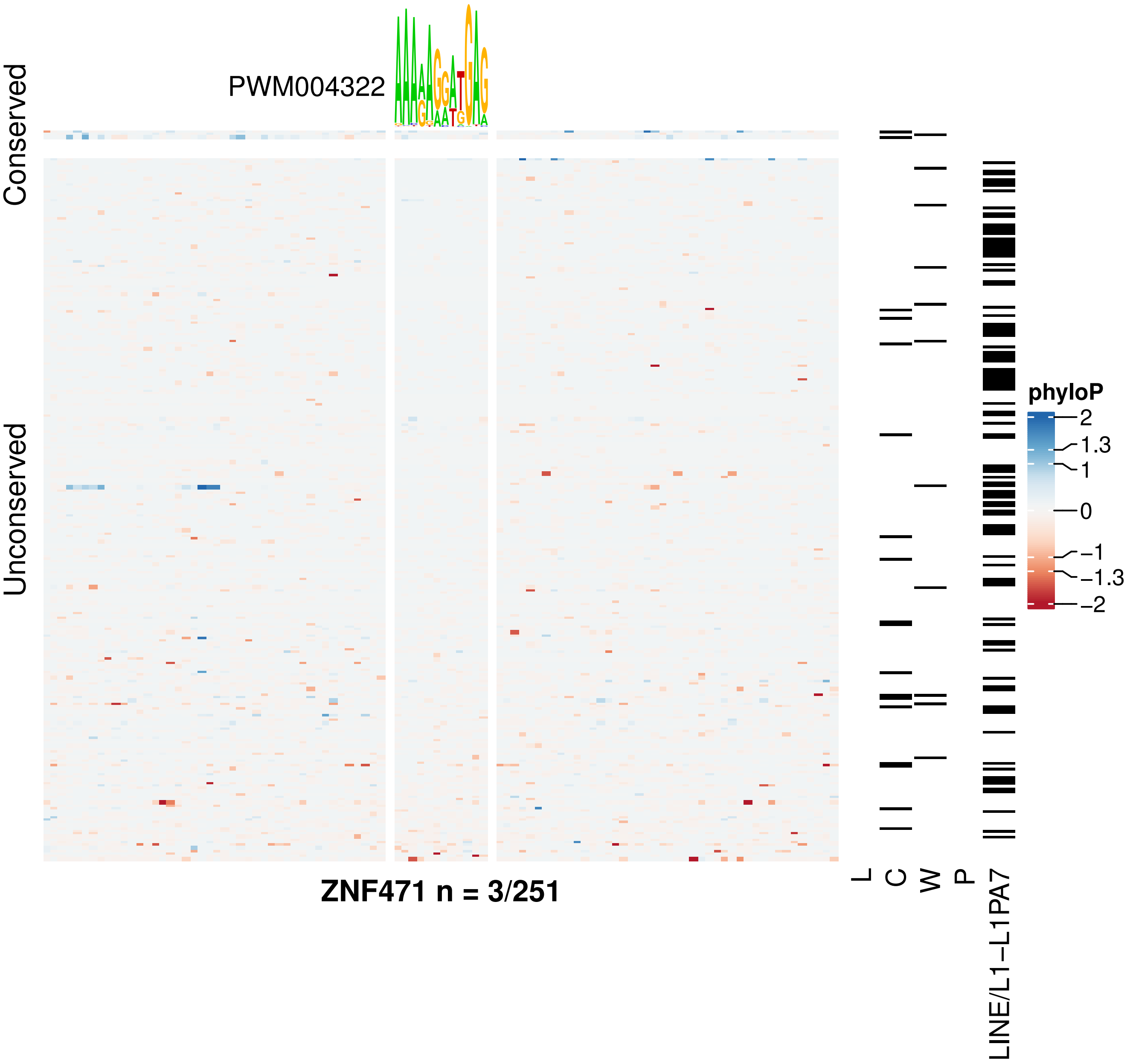

PWM004322
Conserved
Unconserved
ZNF471 n = 3/251
L
C
W
P
LINE/L1−L1PA7
phyloP
2
1.3
1
0
−1
−1.3
−2

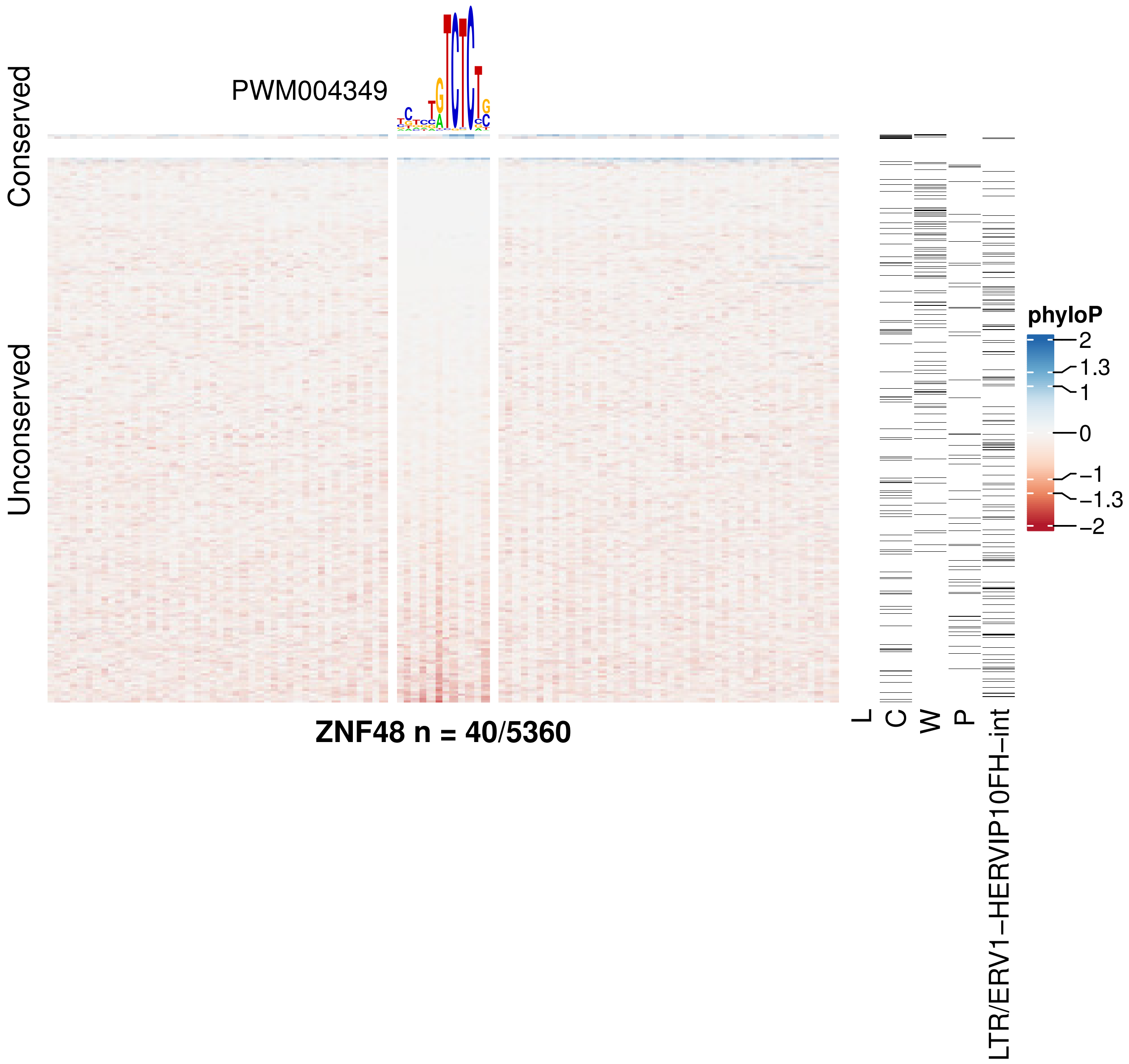

PWM004349
Conserved
Unconserved
ZNF48 n = 40/5360
L
C
W
P
LTR/ERV1−HERVIP10FH−int
phyloP
2
1.3
1
0
−1
−1.3
−2

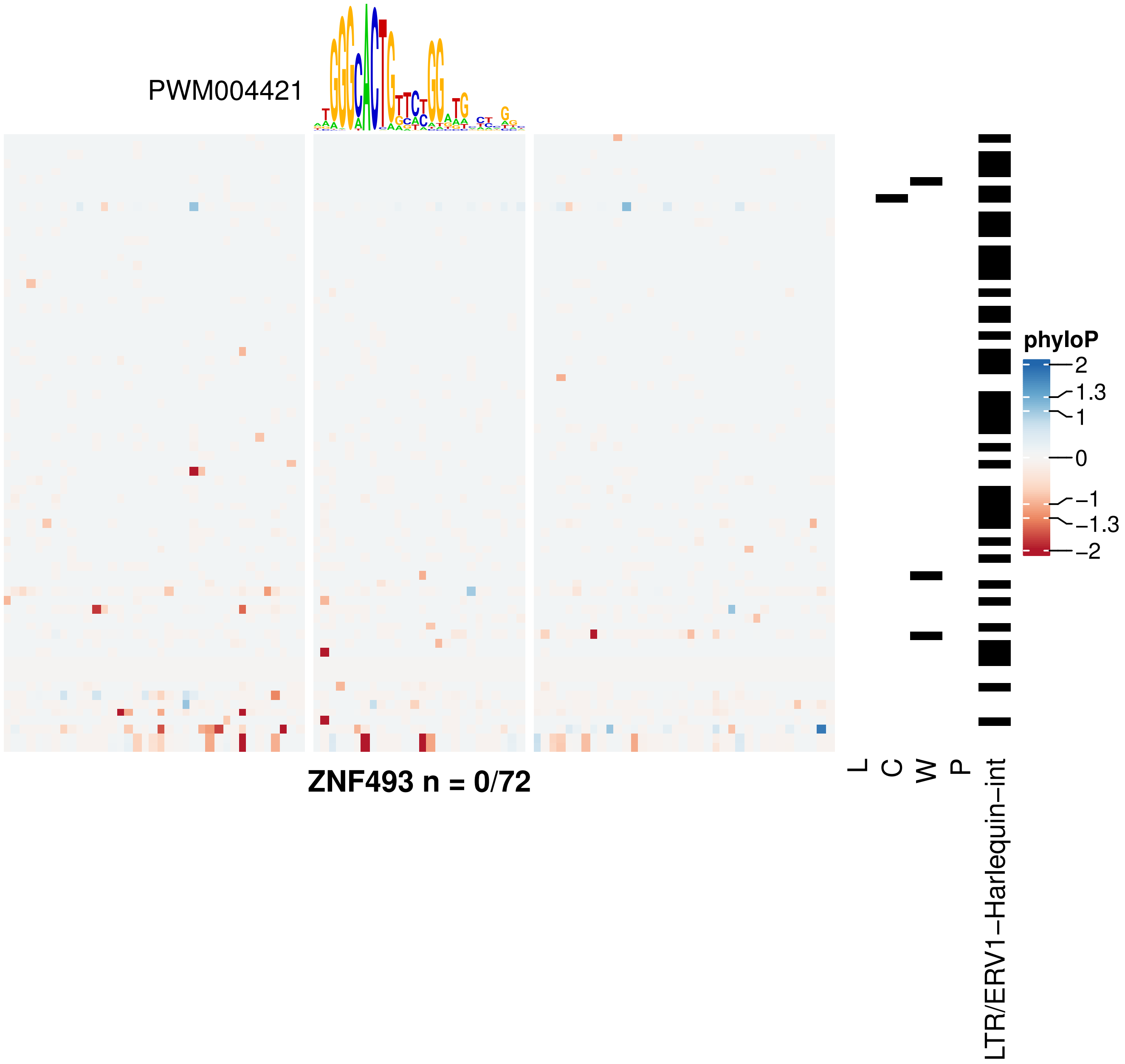

PWM004421
phyloP
2
1.3
1
0
−1
−1.3
−2
L
C
W
P
LTR/ERV1−Harlequin−int
ZNF493 n = 0/72

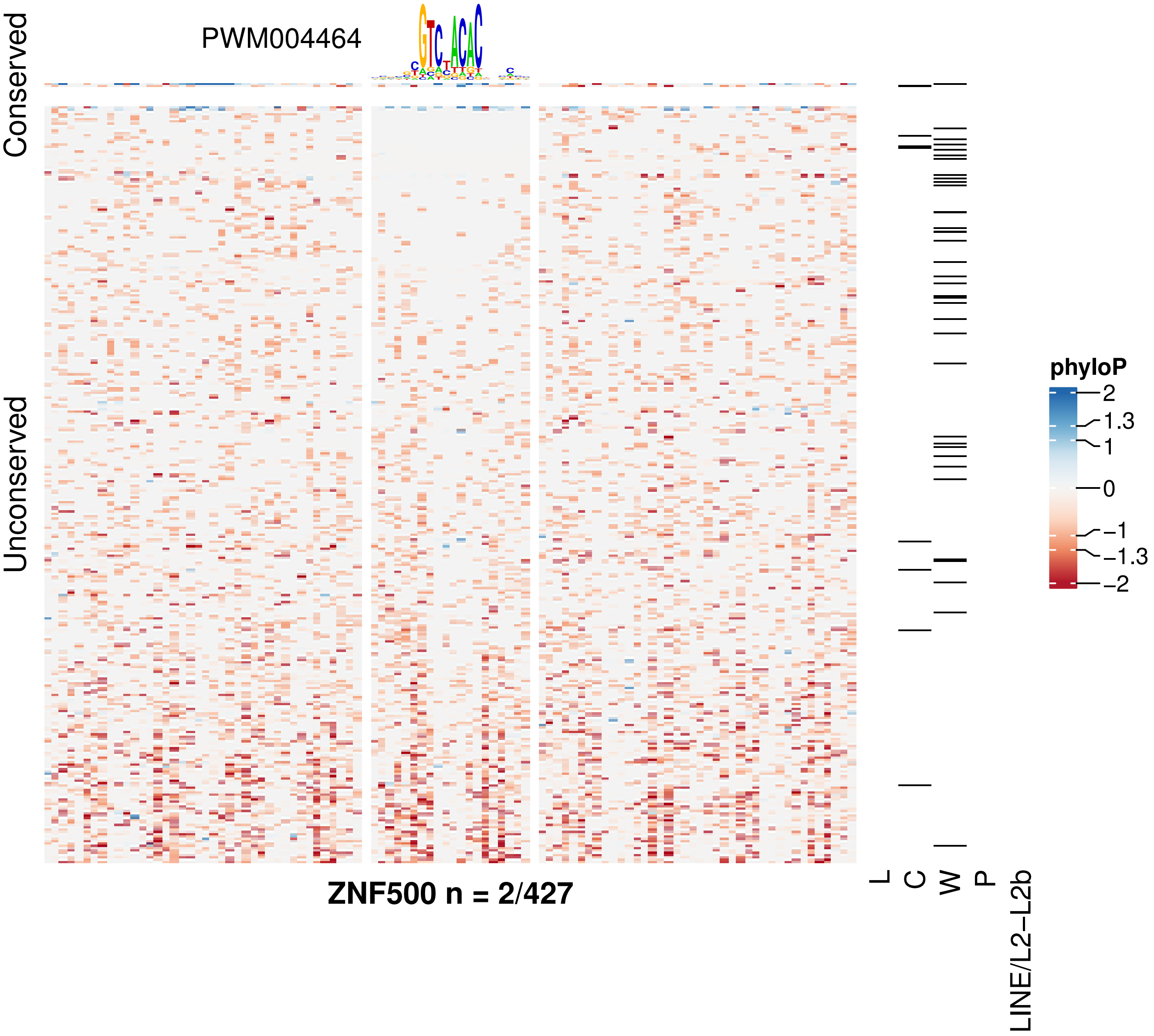

ZNF500 n = 2/427

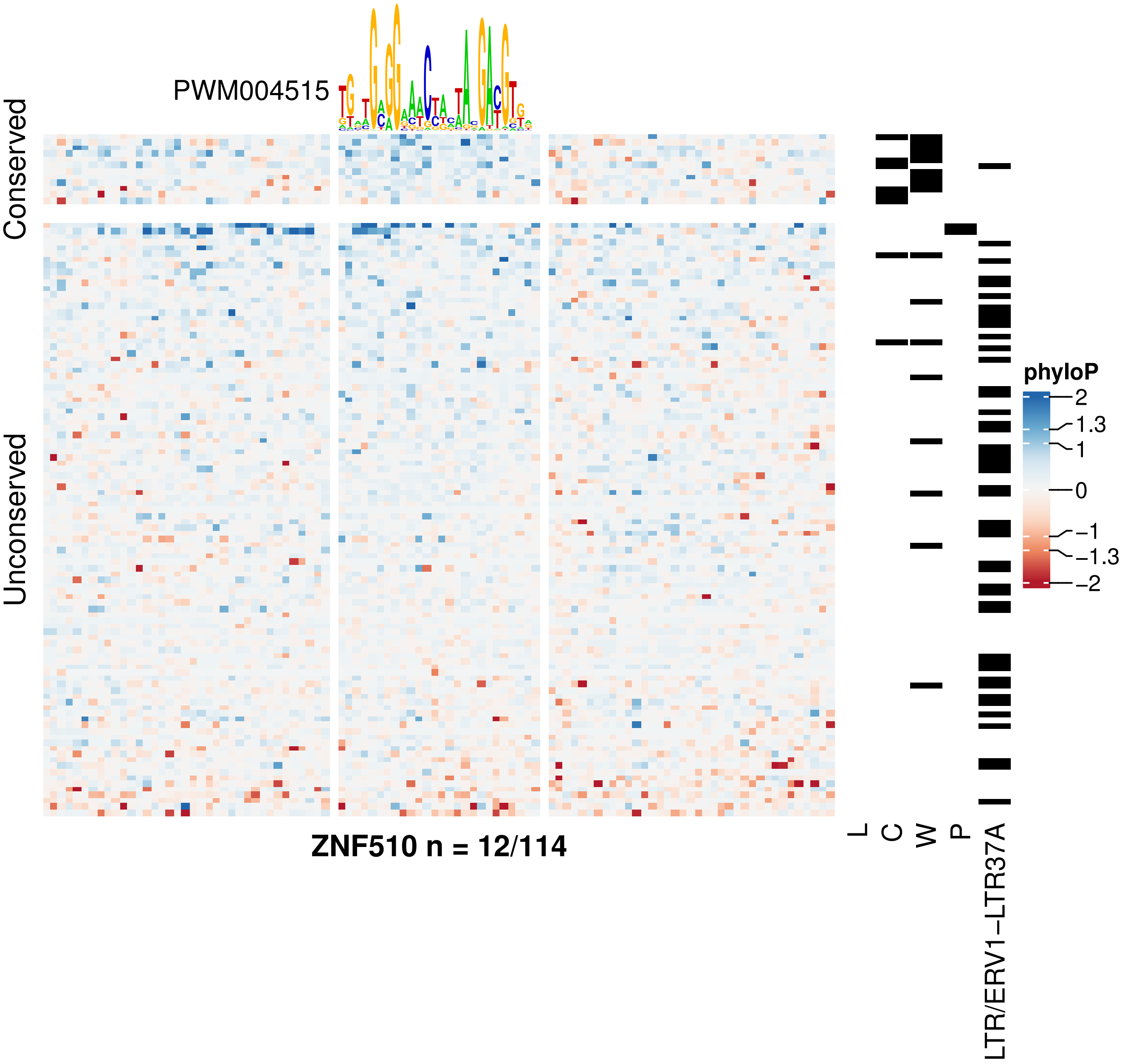

PWM004515
Conserved
Unconserved
ZNF510 n = 12/114
L
C
W
P
LTR/ERV1−LTR37A
phyloP
2
1.3
1
0
−1
−1.3
−2

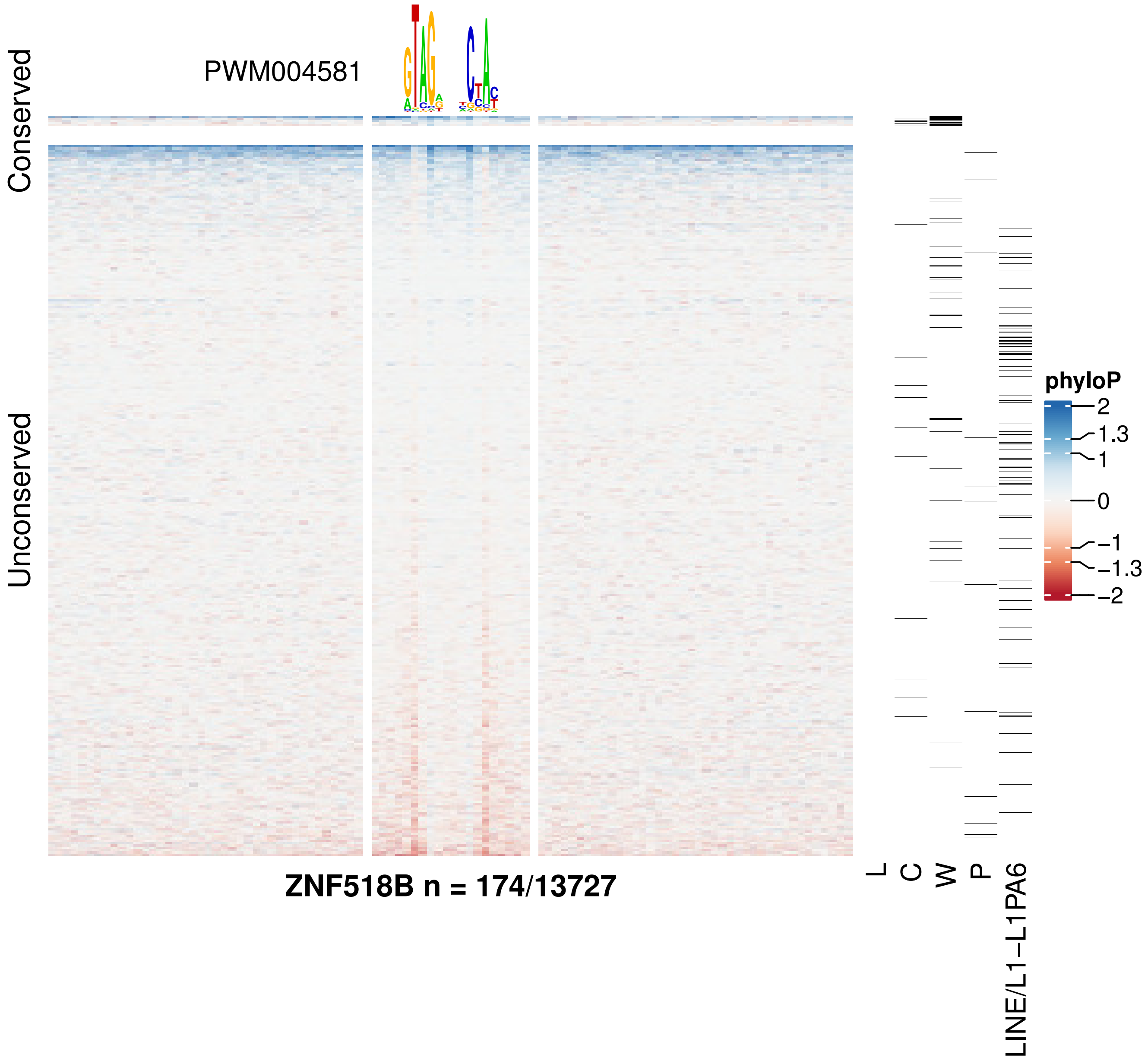

PWM004581
Conserved
Unconserved
ZNF518B n = 174/13727
L
C
W
P
LINE/L1−L1PA6
phyloP
2
1.3
1
0
−1
−1.3
−2

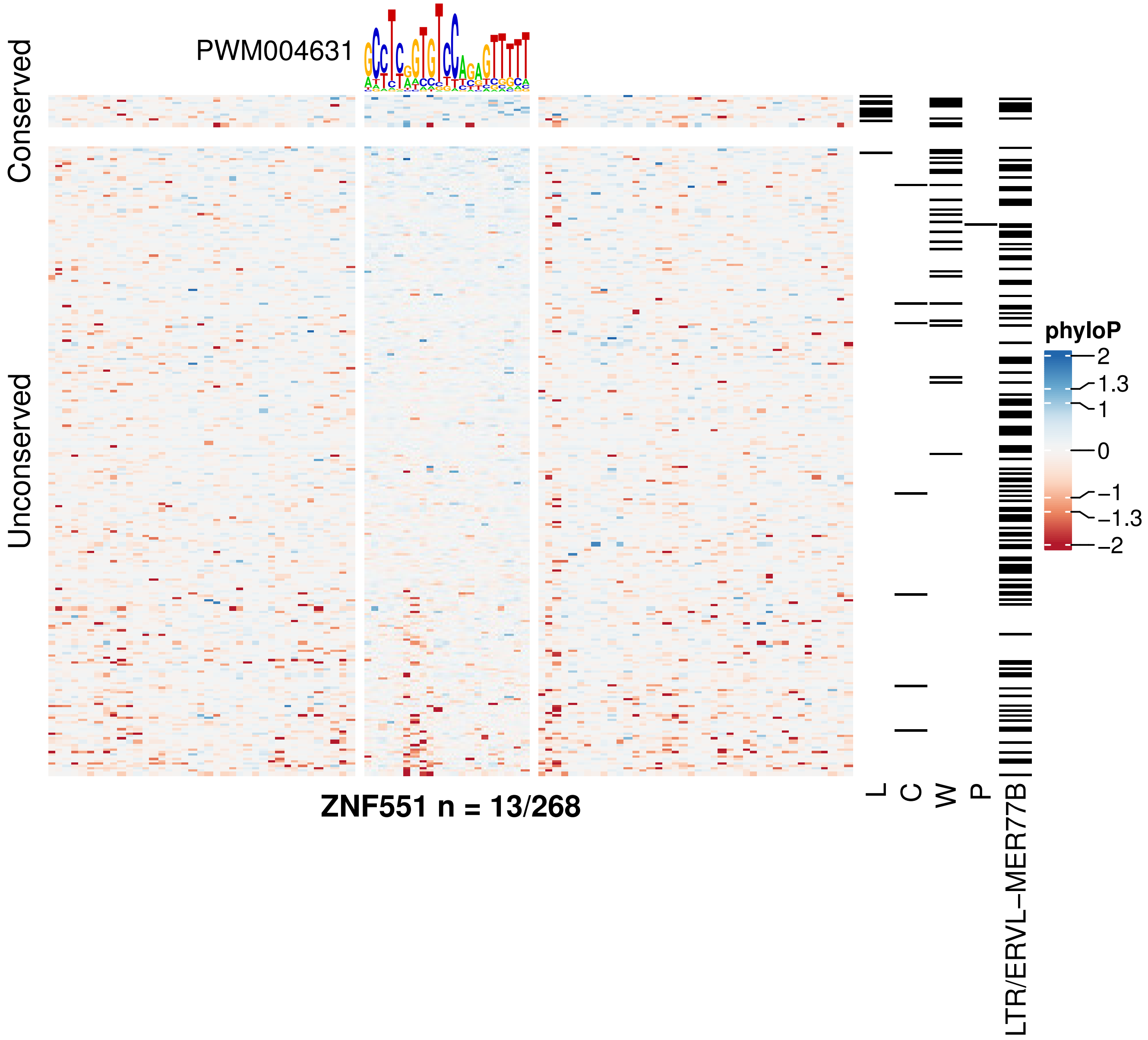
PWM004631
Conserved
Unconserved
ZNF551 n = 13/268
L
C
W
P
LTR/ERVL–MER77B
phyloP
2
1.3
1
0
−1
−1.3
−2

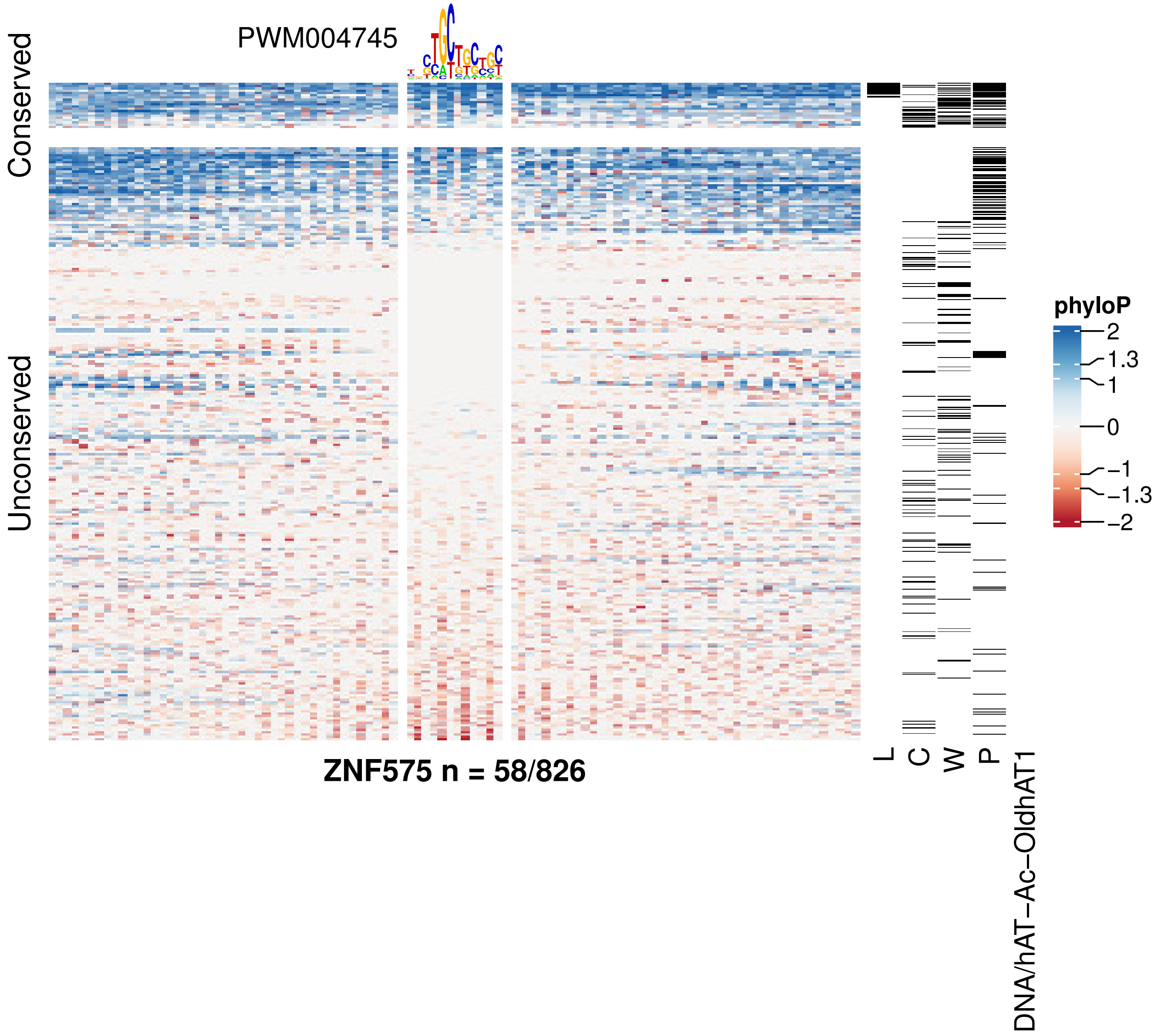

PWM004745
Conserved
Unconserved
ZNF575 n = 58/826
L
C
W
P
DNA/hAT−Ac−OldhAT1
phyloP
2
1.3
1
0
−1
−1.3
−2

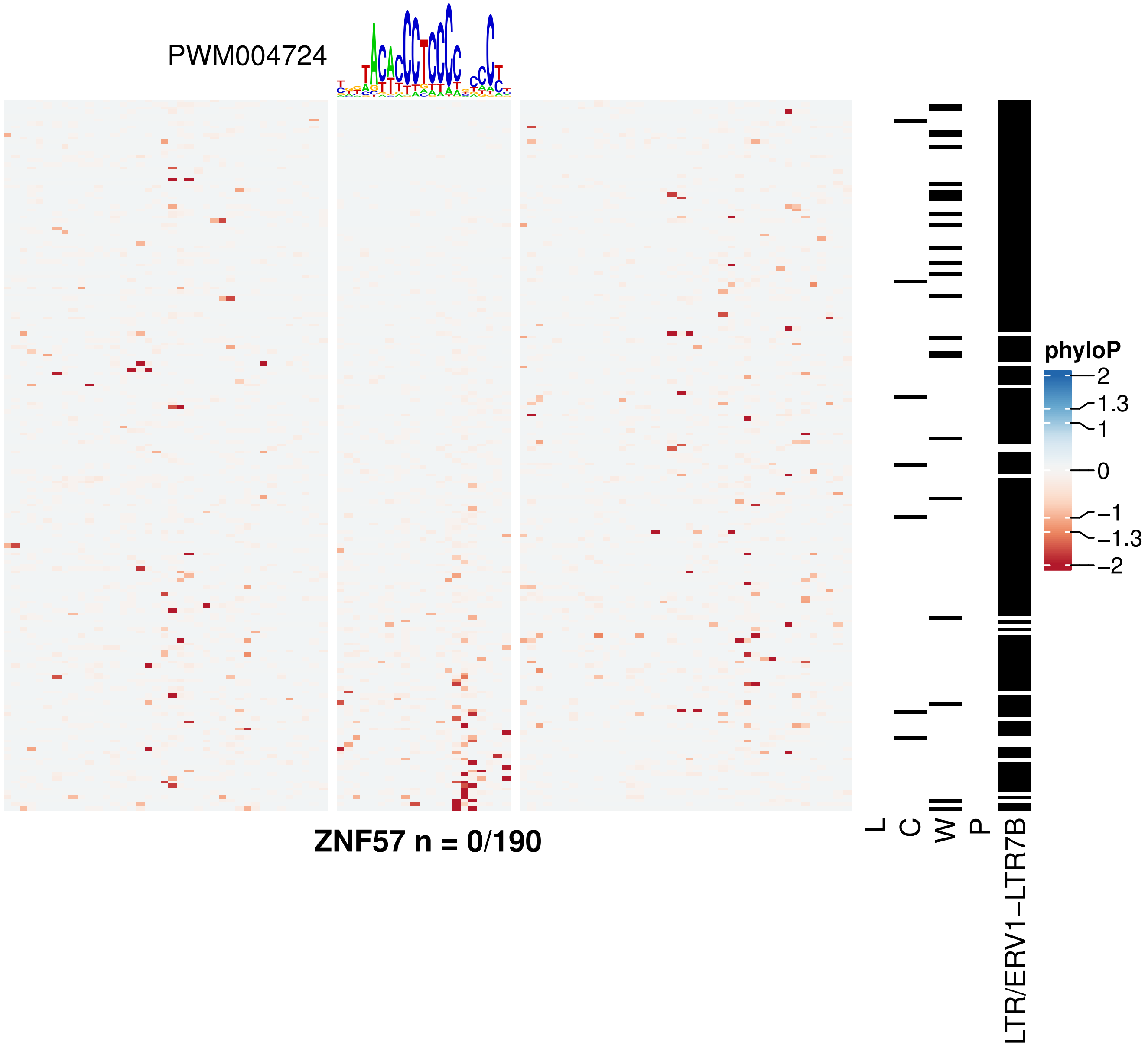

PWM004724
ZNF57 n = 0/190
L
C
W
P
LTR/ERV1−LTR7B
phyloP
2
1.3
1
0
−1
−1.3
−2

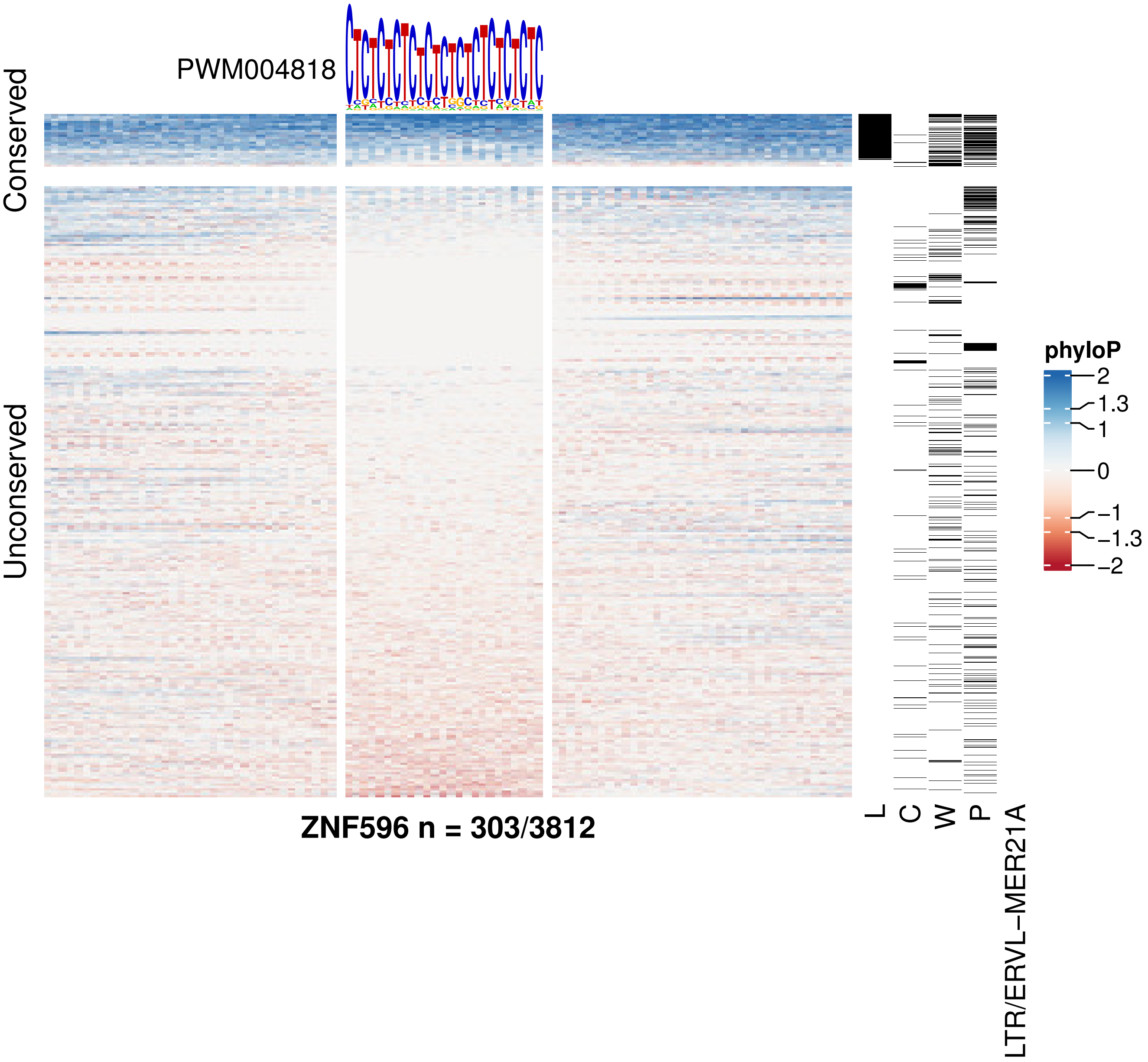

PWM004818
Conserved
Unconserved
phyloP
2
1.3
1
0
−1
−1.3
−2
L
C
W
P
LTR/ERVL−MER21A
ZNF596 n = 303/3812

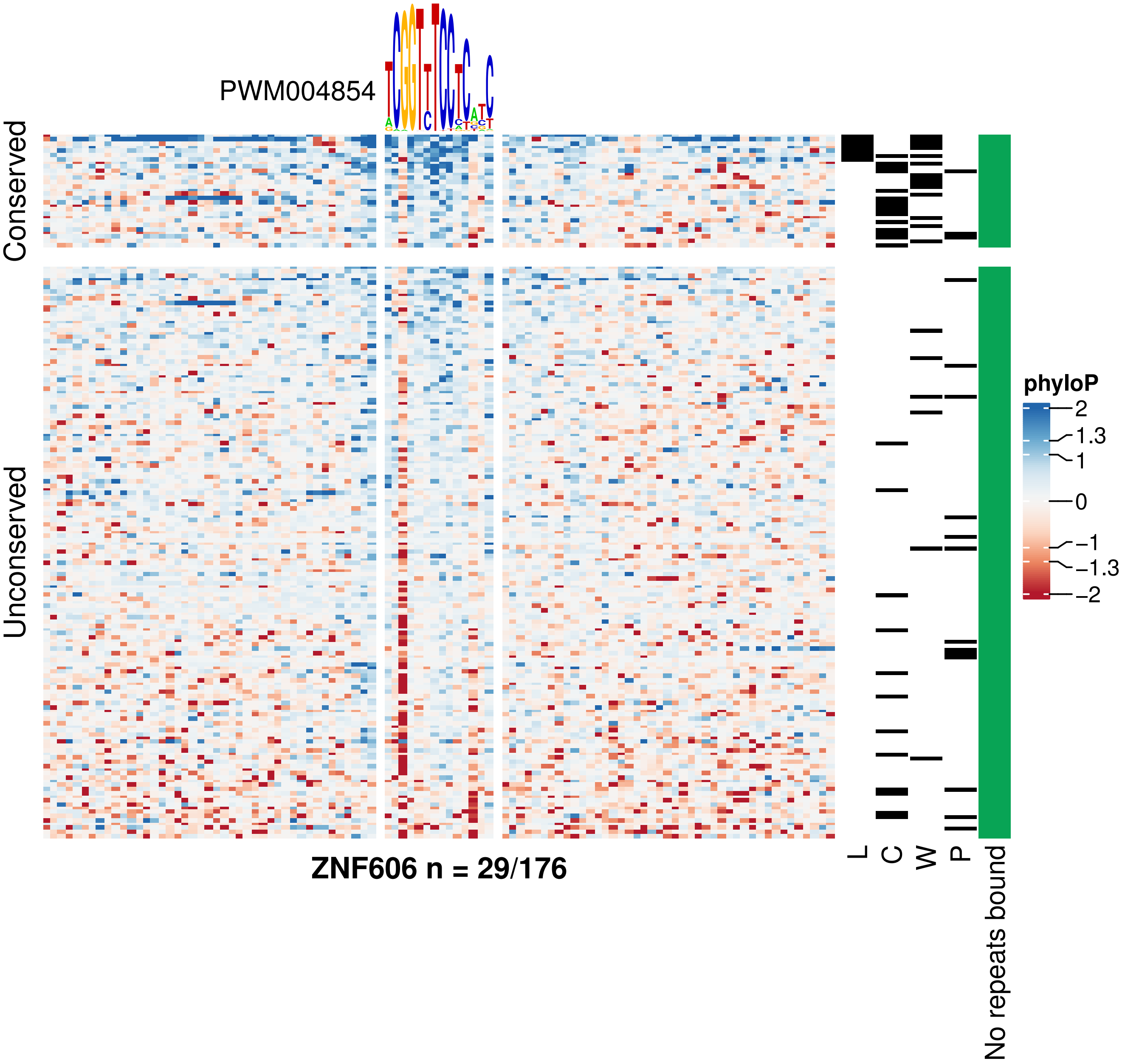
PWM004854
Conserved
Unconserved
ZNF606 n = 29/176
L
C
W
P
No repeats bound
phyloP
2
1.3
1
0
−1
−1.3
−2

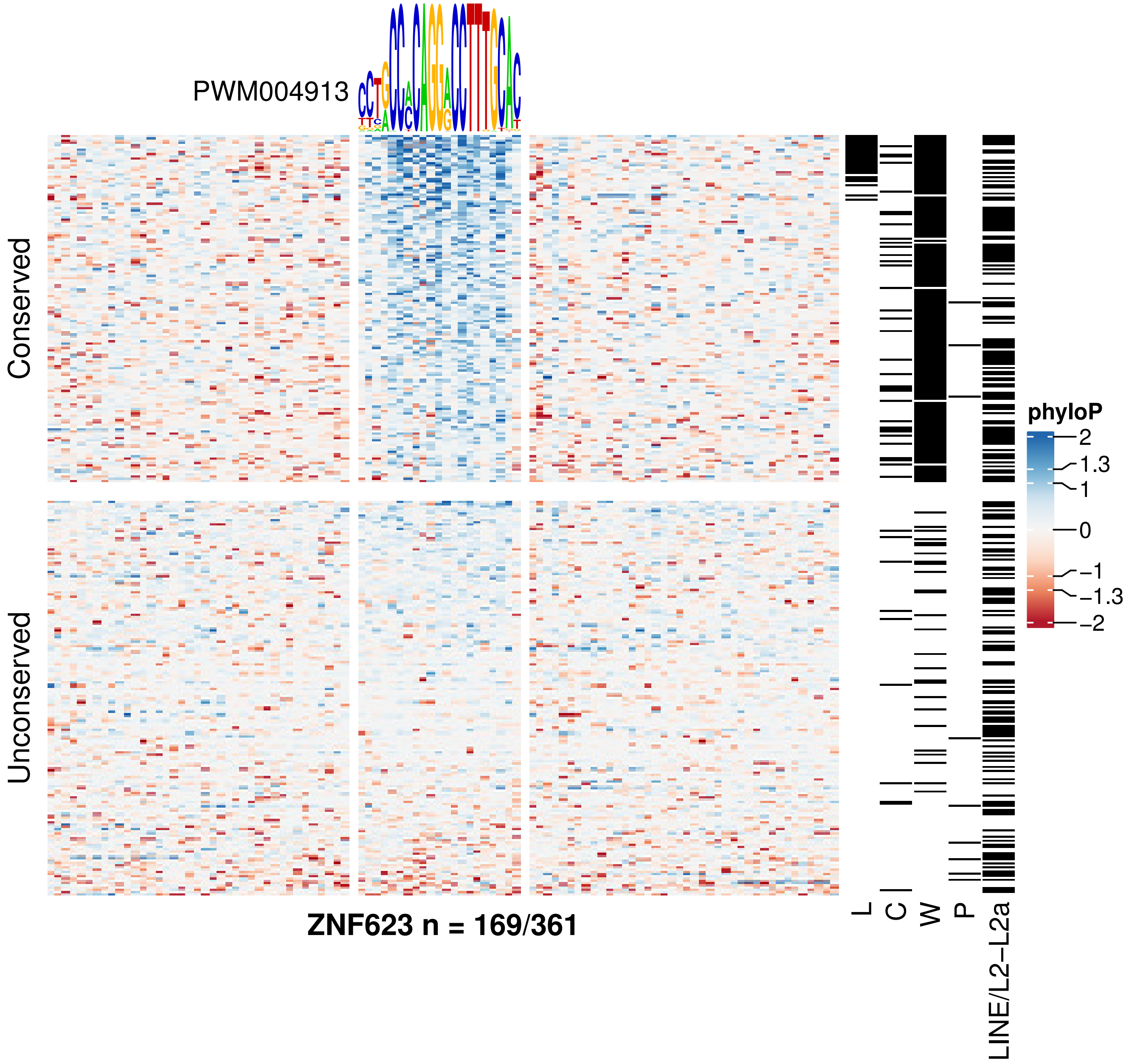
PWM004913
Conserved
Unconserved
ZNF623 n = 169/361
L
C
W
P
LINE/L2−L2a
phyloP
2
1.3
1
0
−1
−1.3
−2

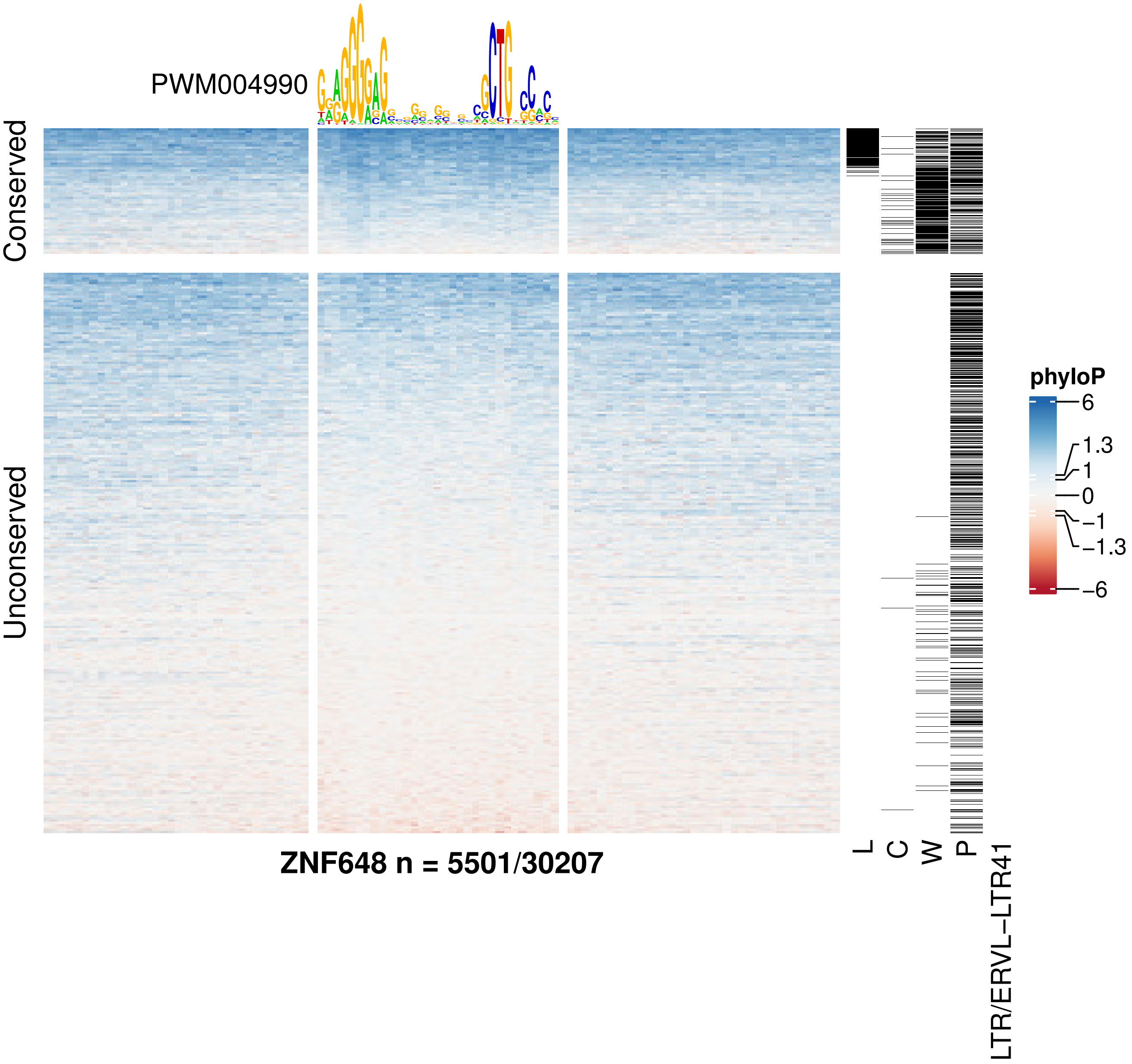
PWM004990
Conserved
Unconserved
ZNF648 n = 5501/30207
L
C
W
P
LTR/ERVL–LTR41
phyloP
6
1.3
1
0
−1
−1.3
−6

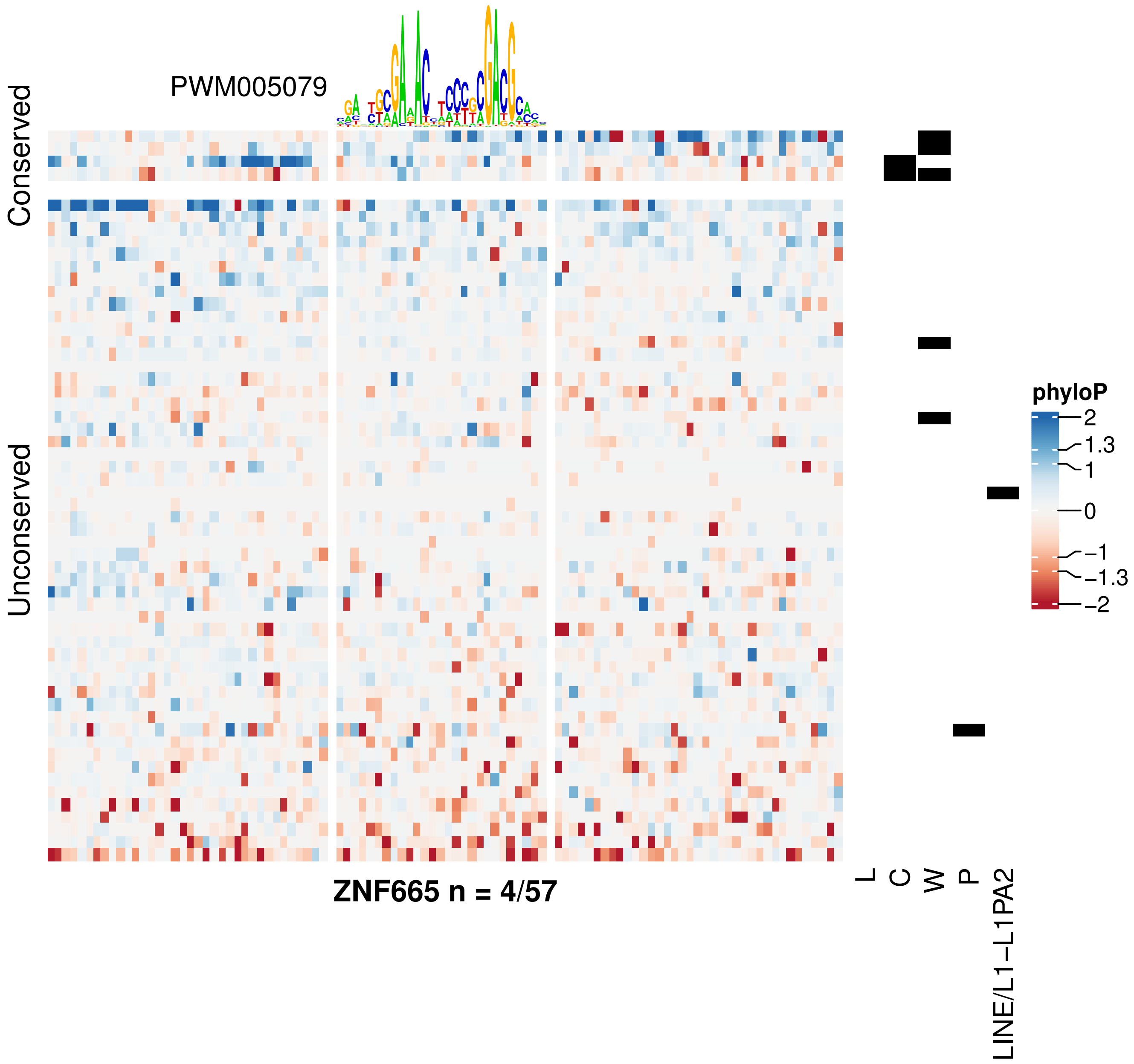
PWM005079
Conserved
Unconserved
ZNF665 n = 4/57
L
C
W
P
LINE/L1−L1PA2
phyloP
2
1.3
1
0
−1
−1.3
−2

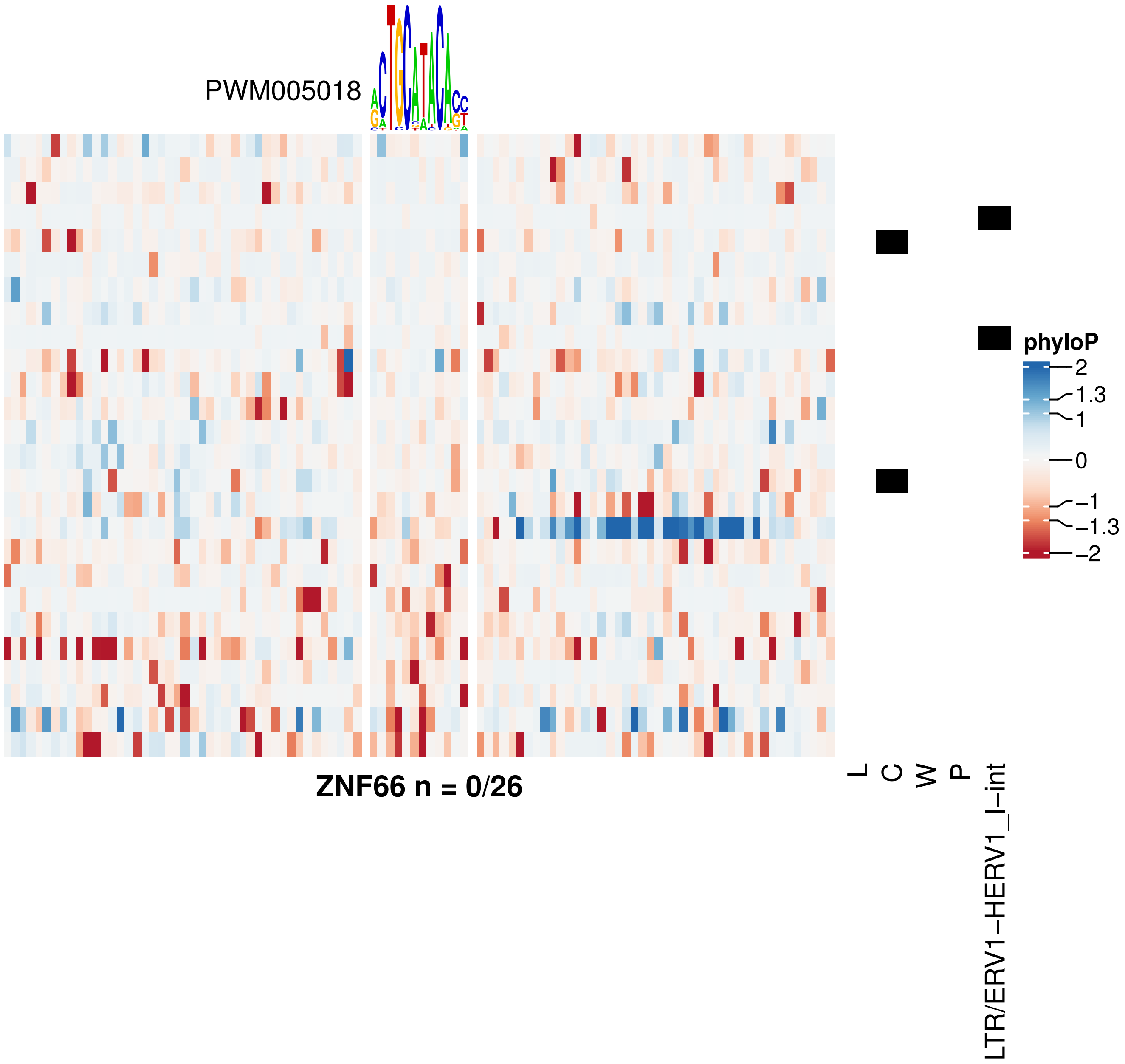
PWM005018
ZNF66 n = 0/26
L
C
W
P
LTR/ERV1−HERV1_I−int
phyloP
2
1.3
1
0
−1
−1.3
−2

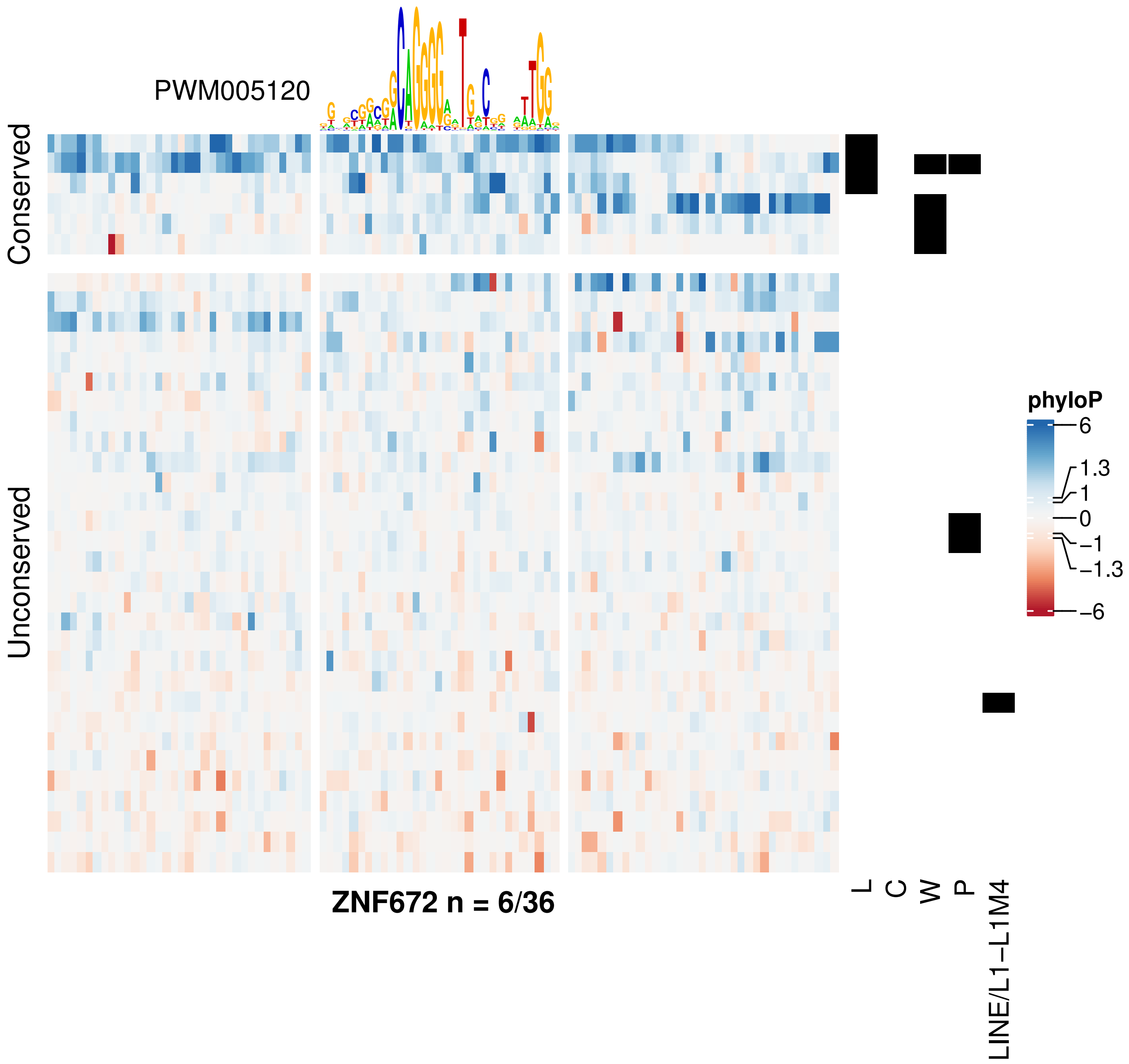

PWM005120
Conserved
Unconserved
phyloP
6
1.3
1
0
−1
−1.3
−6
ZNF672 n = 6/36
L
C
W
P
LINE/L1−L1M4

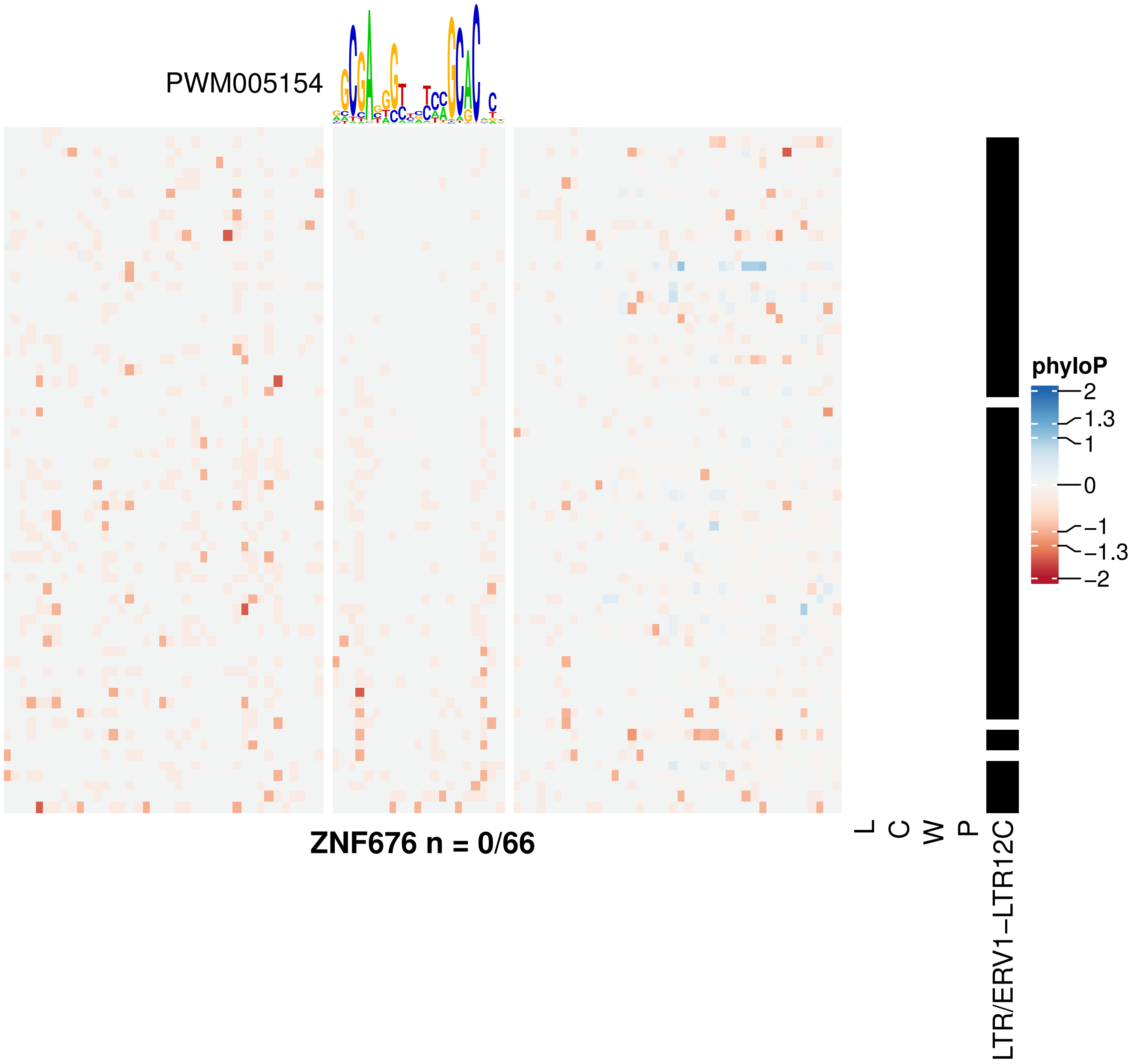
PWM005154
ZNF676 n = 0/66
L
C
W
P
LTR/ERV1–LTR12C
phyloP
2
1.3
1
0
−1
−1.3
−2

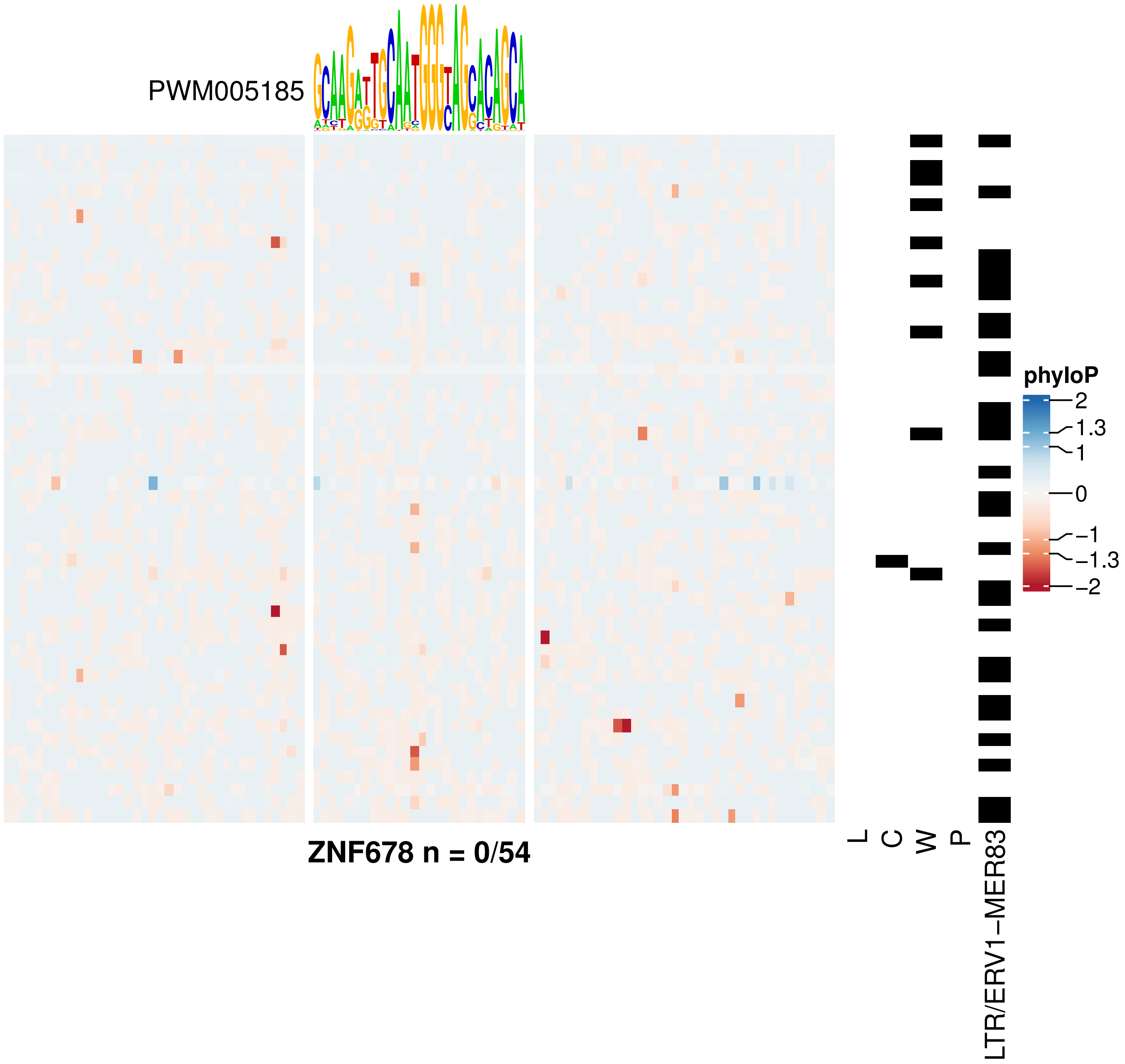
PWM005185
phyloP
2
1.3
1
0
−1
−1.3
−2
ZNF678 n = 0/54
L
C
W
P
LTR/ERV1−MER83

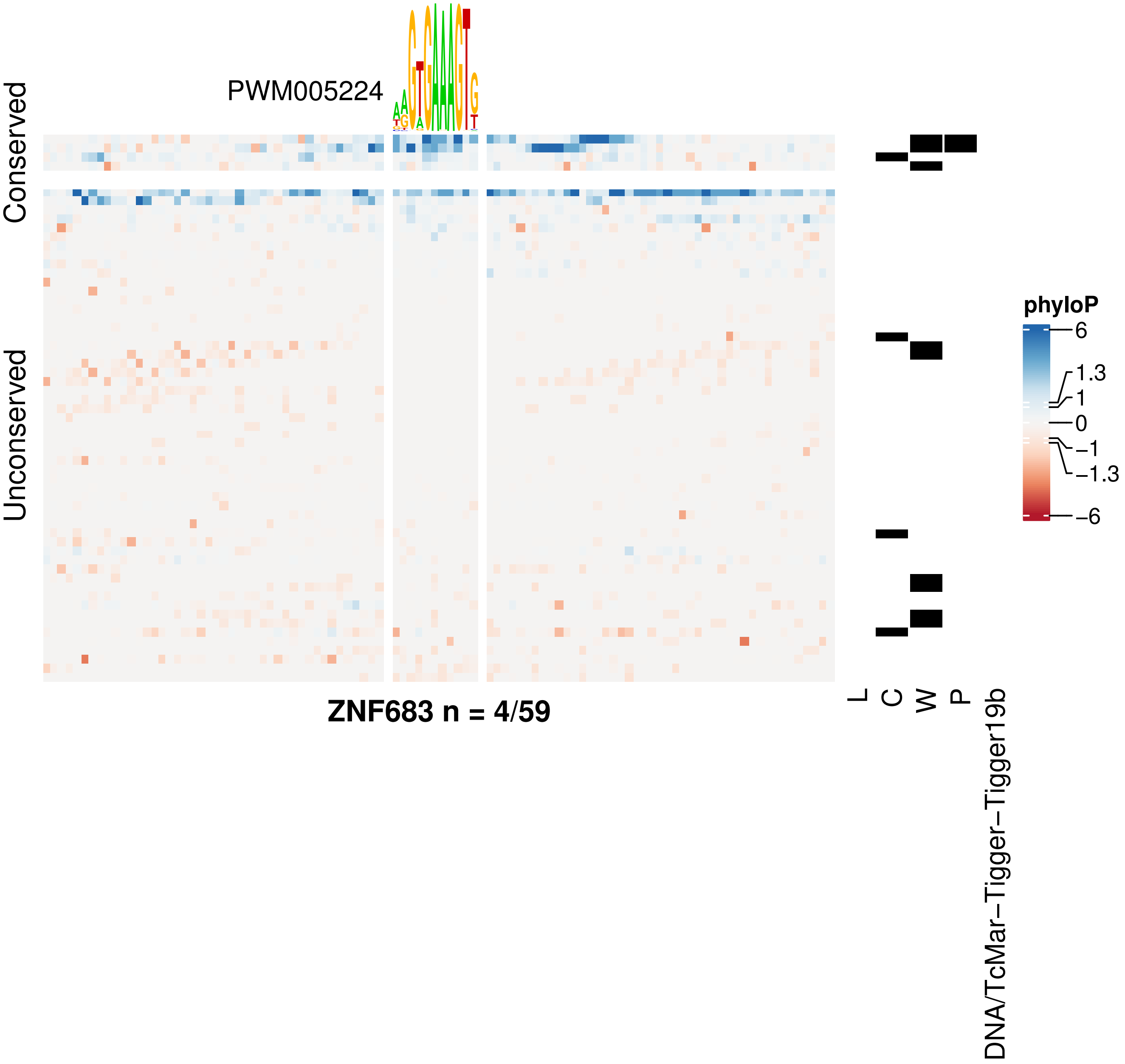

PWM005224
Conserved
Unconserved
phyloP
6
1.3
1
0
−1
−1.3
−6
ZNF683 n = 4/59
L
C
W
P
DNA/TcMar−Tigger−Tigger19b

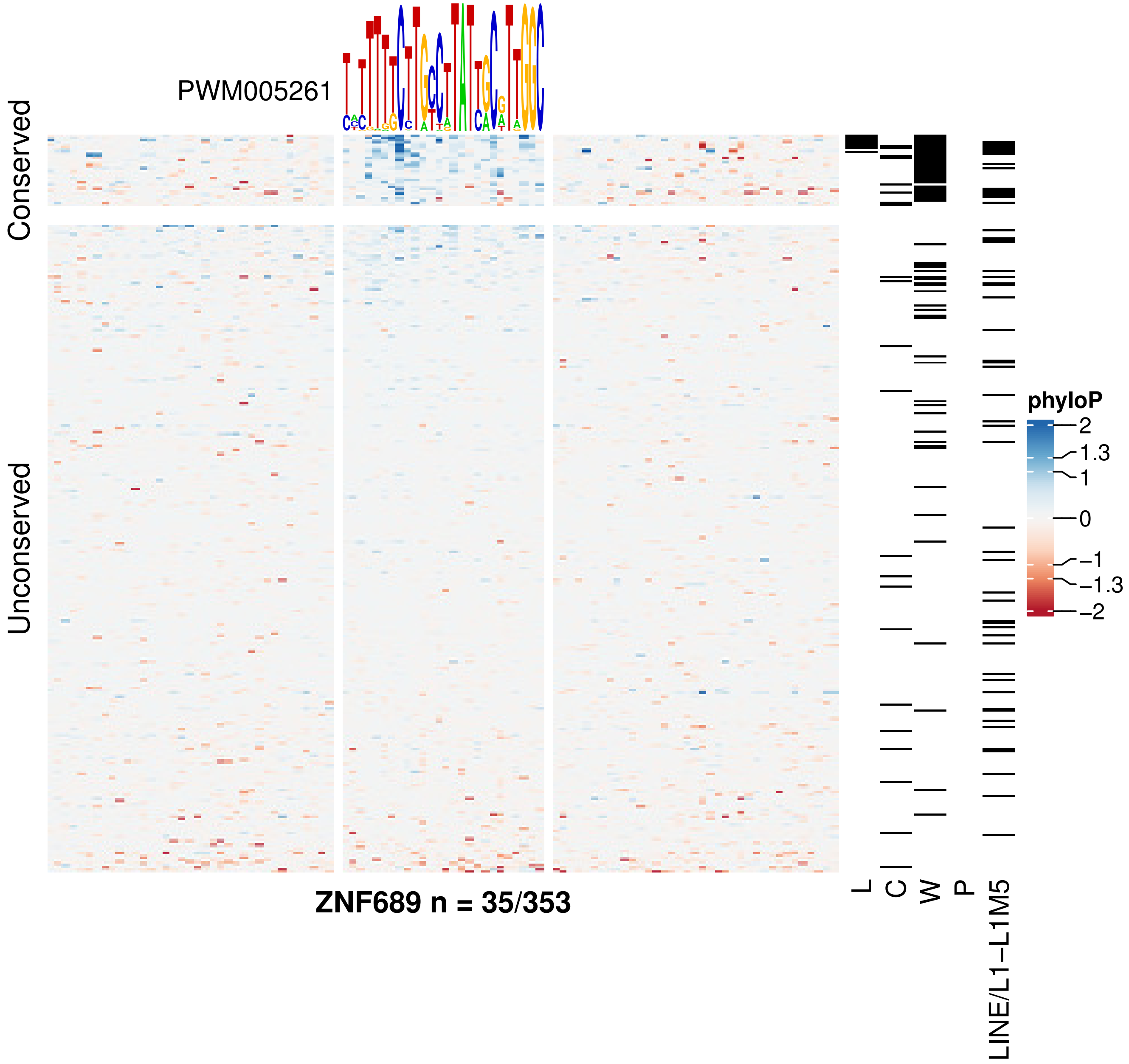

PWM005261
Conserved
Unconserved
ZNF689 n = 35/353
L
C
W
P
LINE/L1−L1M5
phyloP
2
1.3
1
0
−1
−1.3
−2

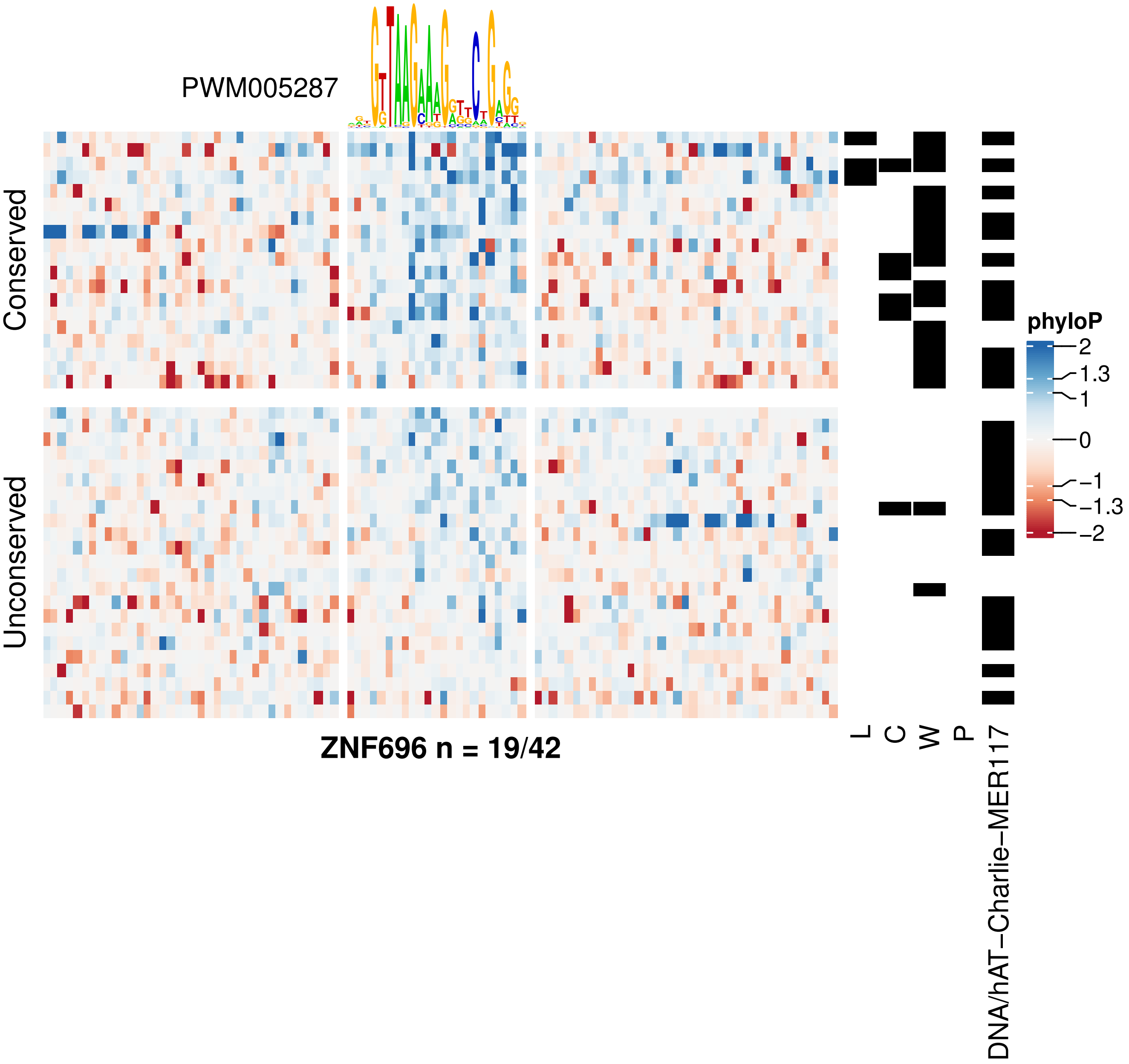
PWM005287
Conserved
Unconserved
ZNF696 n = 19/42
L
C
W
P
DNA/hAT−Charlie−MER117
phyloP
2
1.3
1
0
−1
−1.3
−2

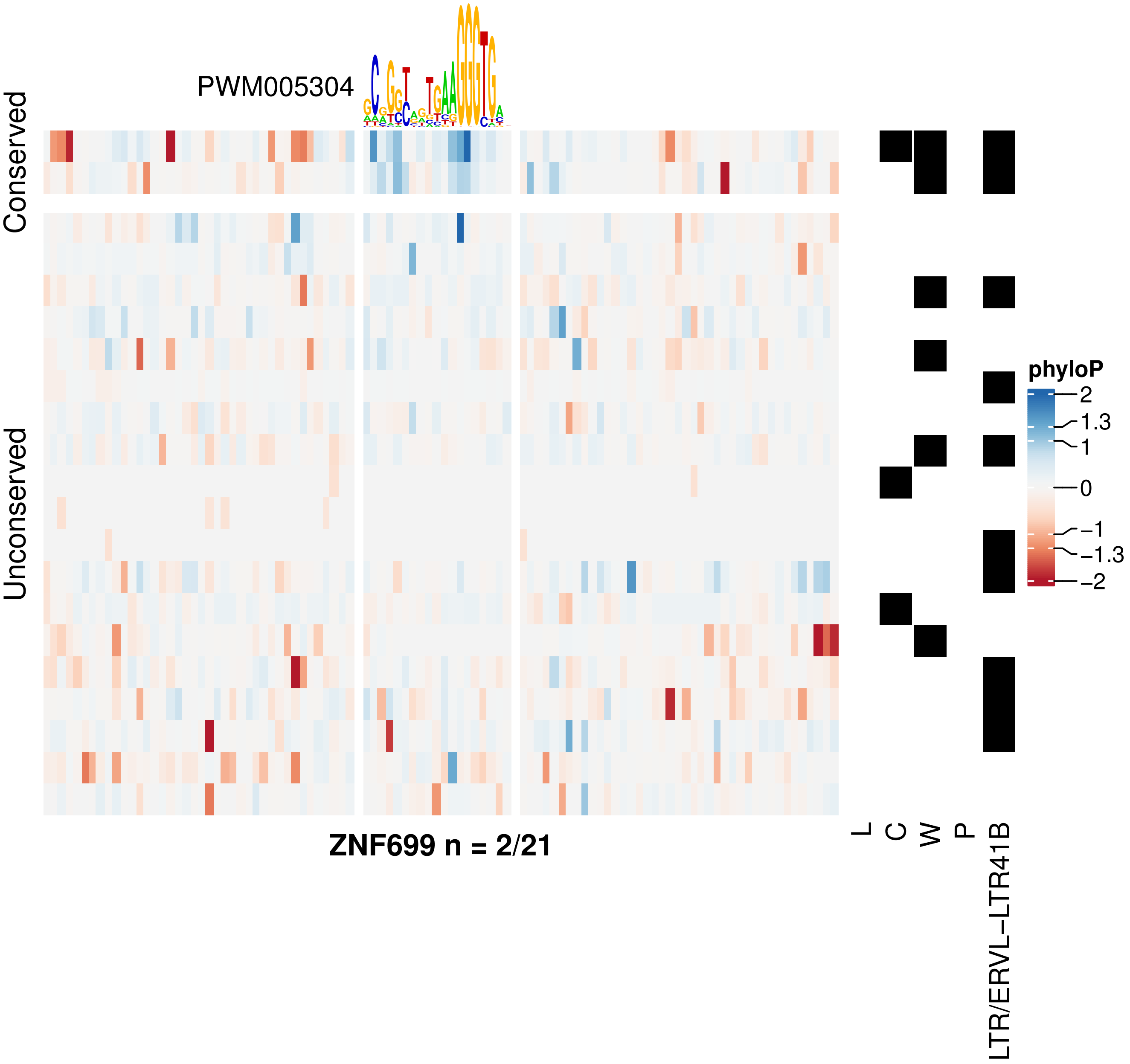
PWM005304
Conserved
Unconserved
ZNF699 n = 2/21
L
C
W
P
LTR/ERVL–LTR41B
phyloP
2
1.3
1
0
−1
−1.3
−2

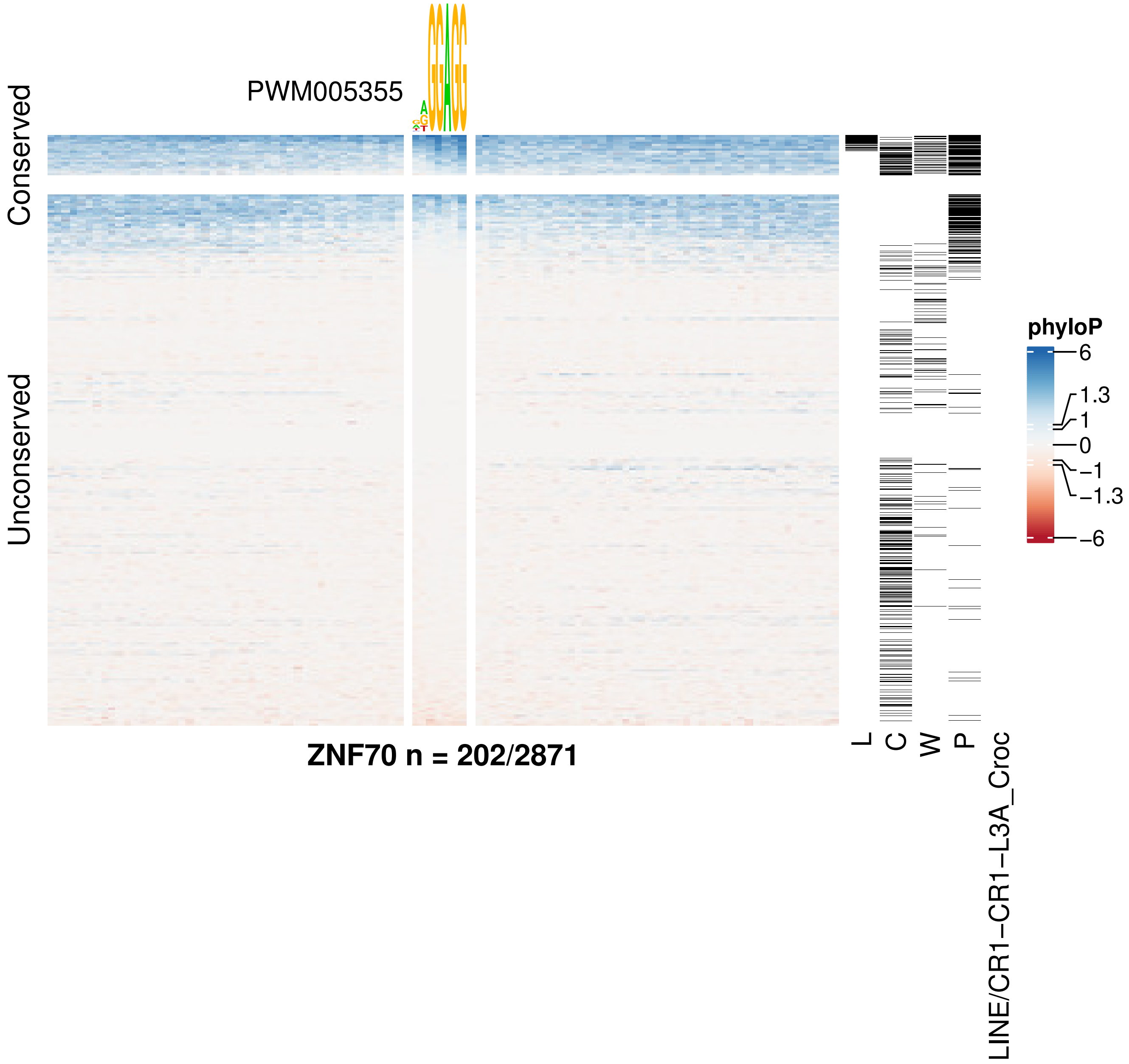
PWM005355
Conserved
Unconserved
ZNF70 n = 202/2871
L
C
W
P
LINE/CR1−CR1−L3A_Croc
phyloP
6
1.3
1
0
−1
−1.3
−6

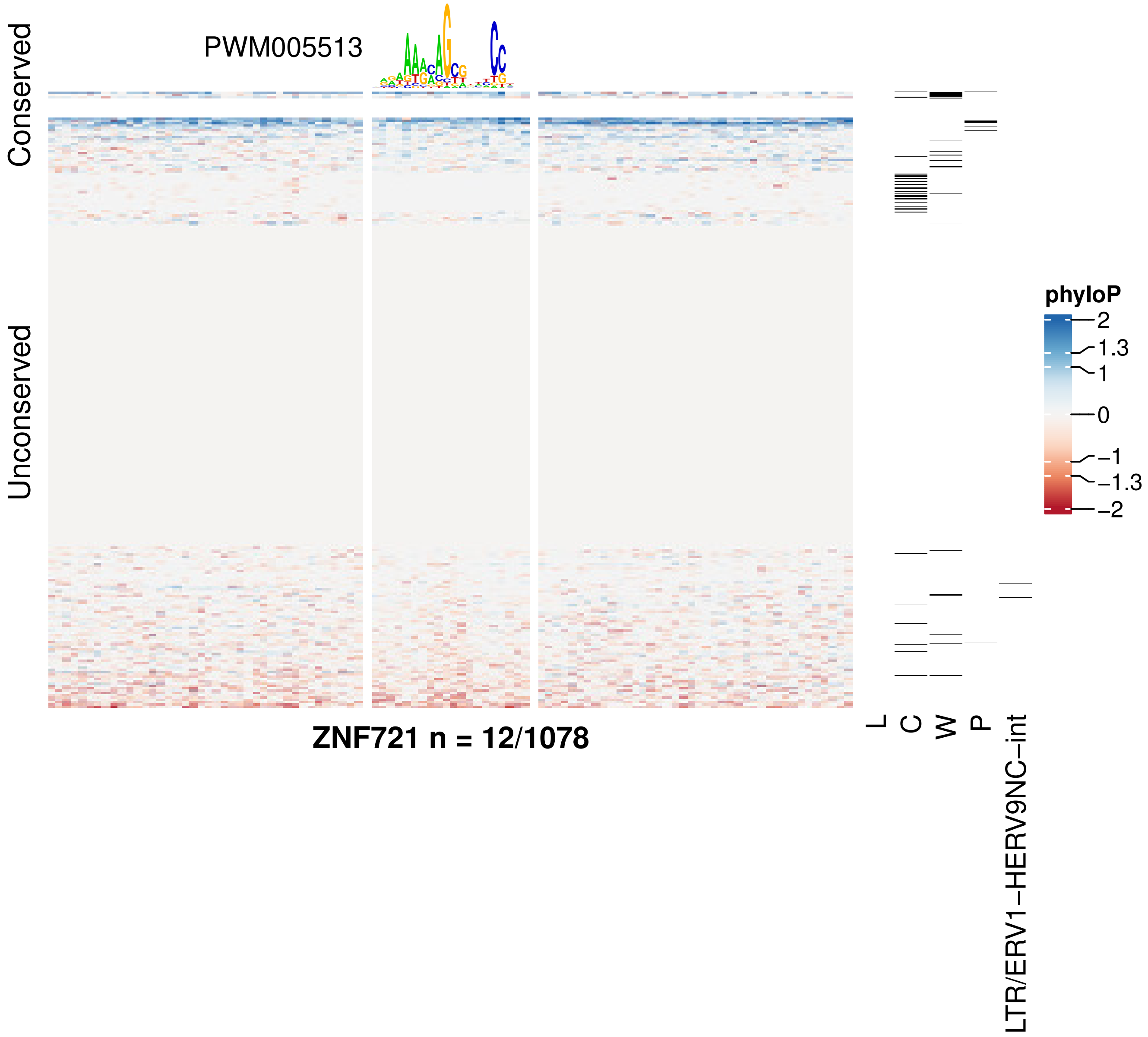
PWM005513
Conserved
Unconserved
phyloP
2
1.3
1
0
−1
−1.3
−2
ZNF721 n = 12/1078
L
C
W
P
LTR/ERV1−HERV9NC−int

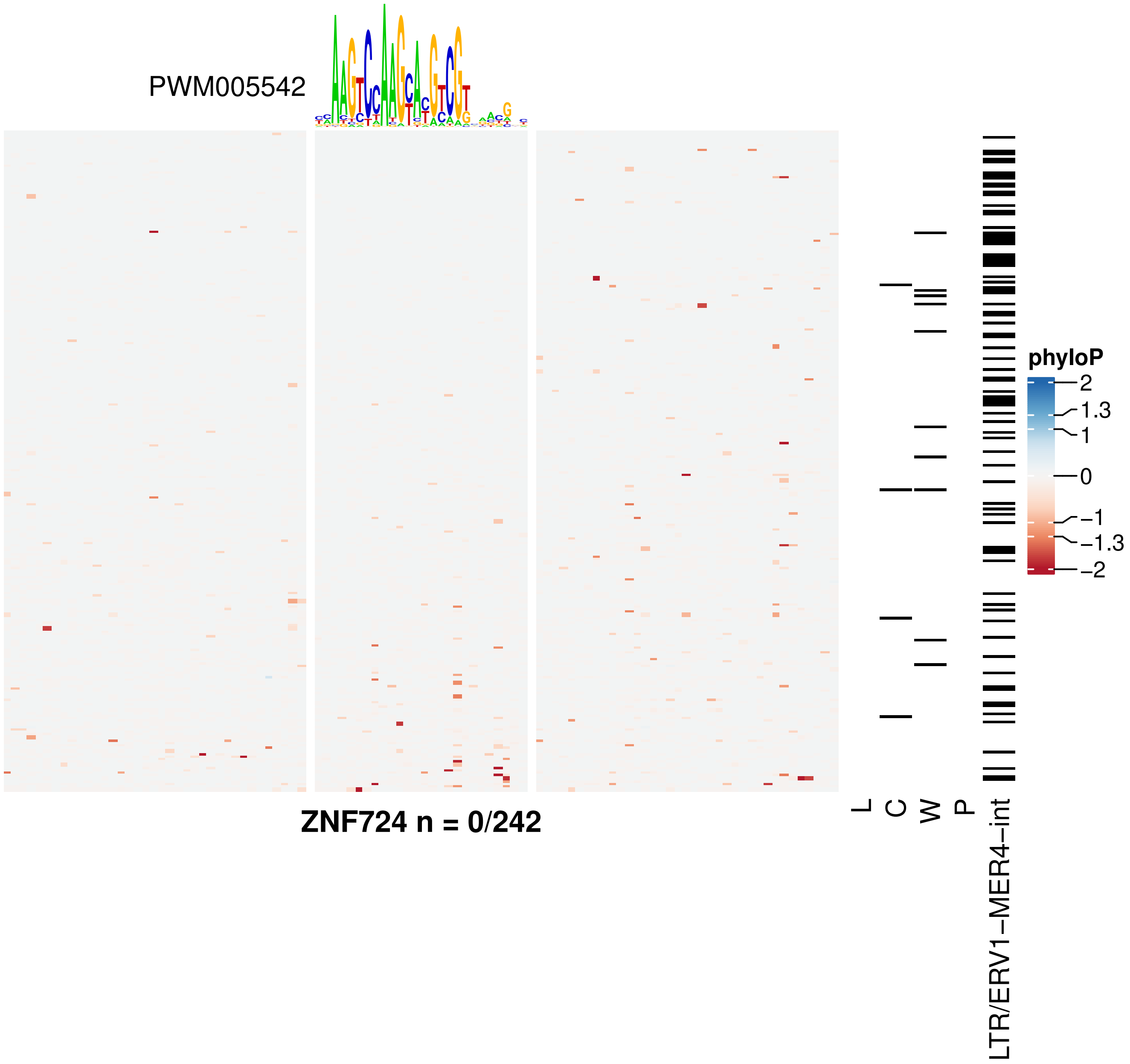
PWM005542
ZNF724 n = 0/242
L
C
W
P
LTR/ERV1−MER4−int
phyloP
2
1.3
1
0
−1
−1.3
−2

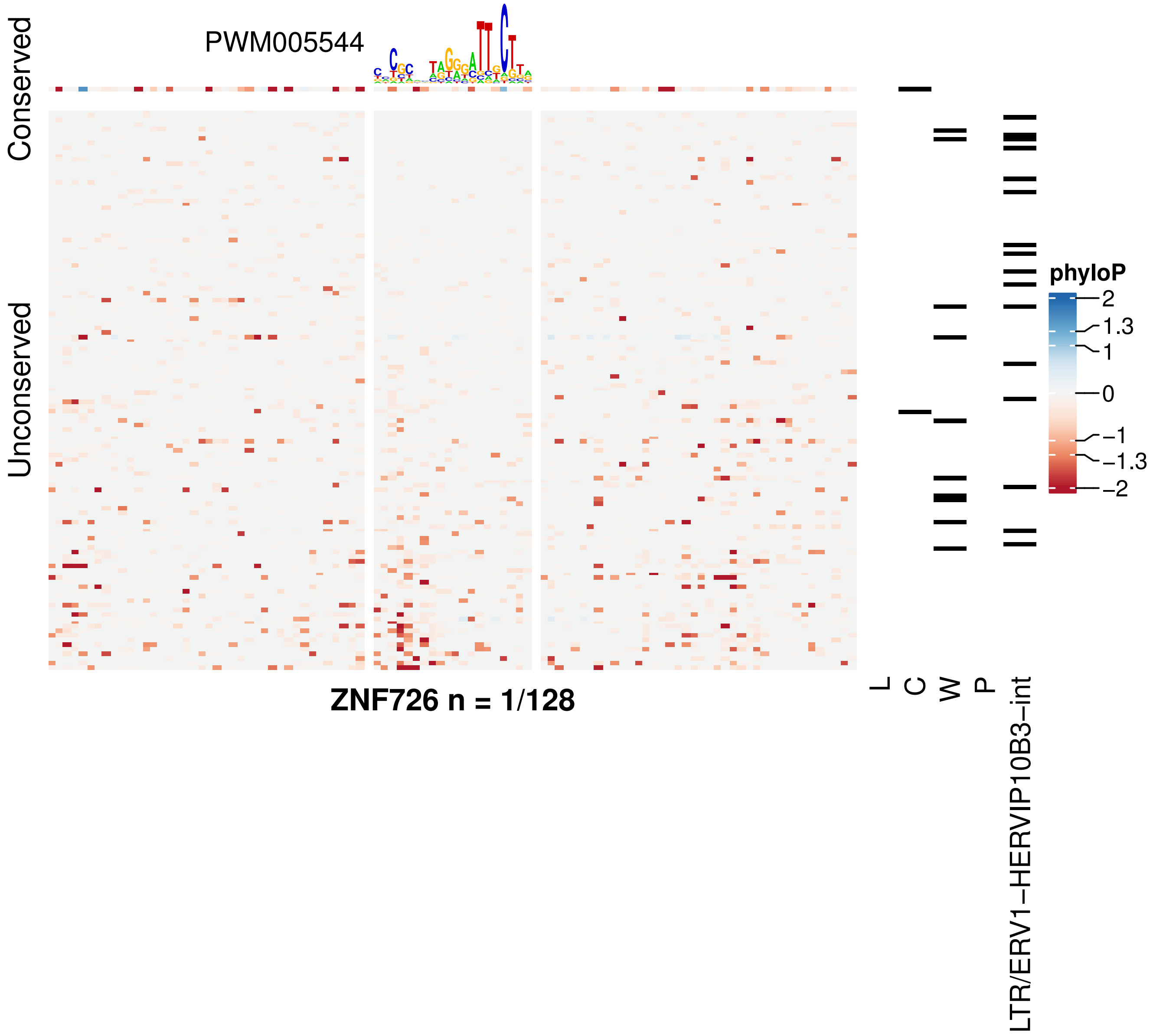

PWM005544
Conserved
Unconserved
phyloP
2
1.3
1
0
−1
−1.3
−2
ZNF726 n = 1/128
L
C
W
P
LTR/ERV1−HERVIP10B3−int

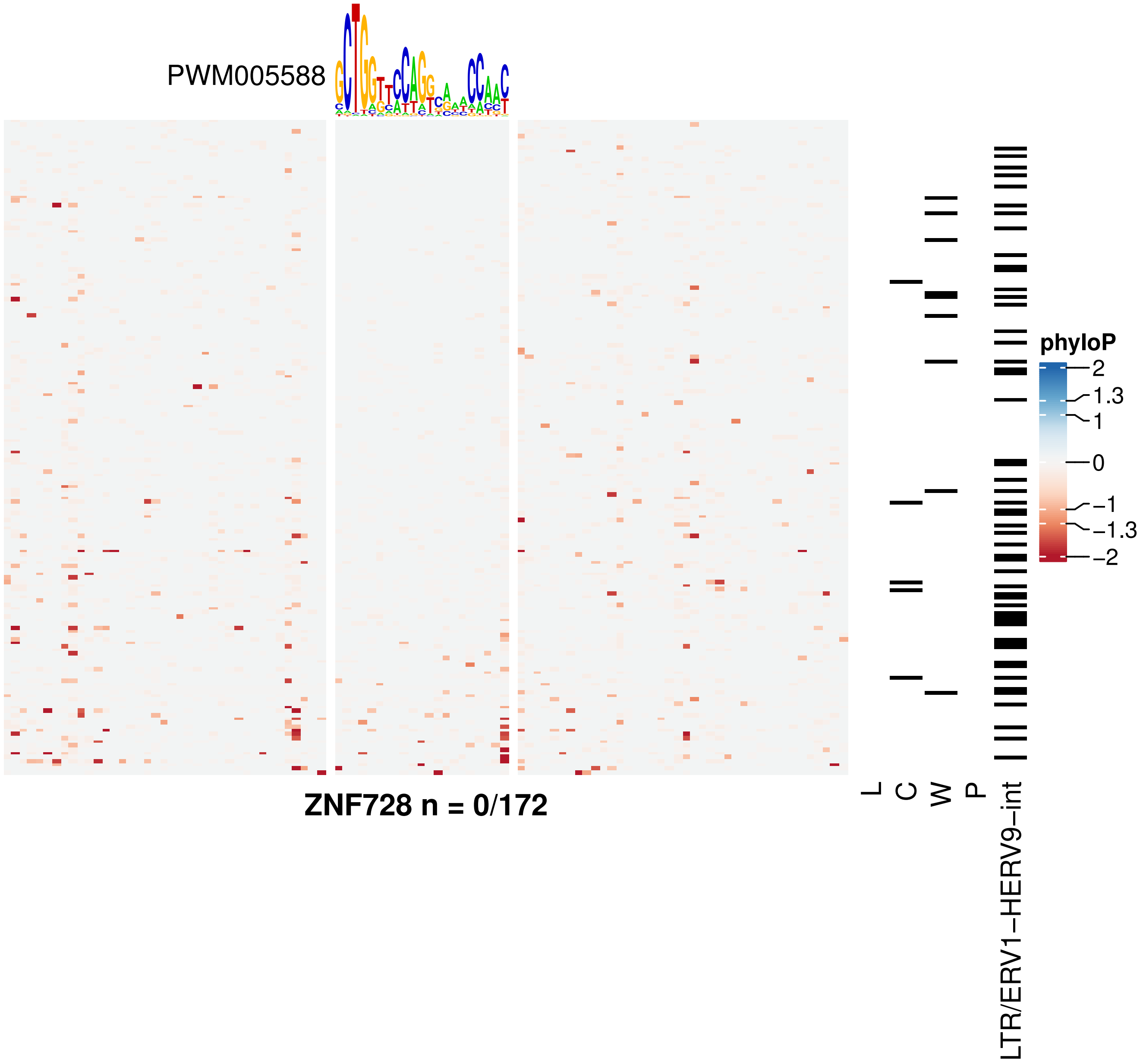
PWM005588
ZNF728 n = 0/172
L
C
W
P
LTR/ERV1–HERV9–int
phyloP
2
1.3
1
0
−1
−1.3
−2

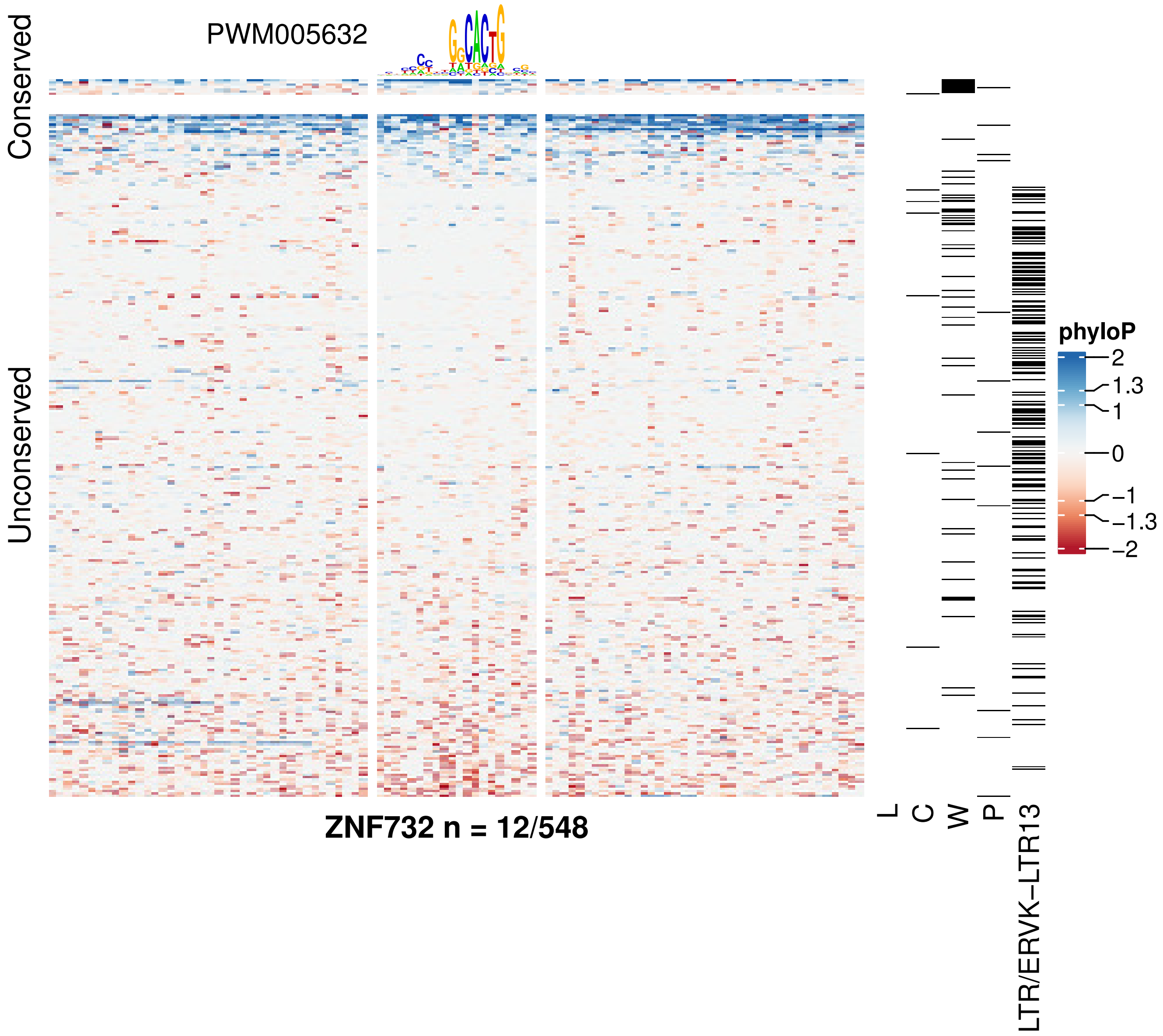
PWM005632
Conserved
Unconserved
phyloP
2
1.3
1
0
−1
−1.3
−2
ZNF732 n = 12/548
L
C
W
P
LTR/ERVK−LTR13

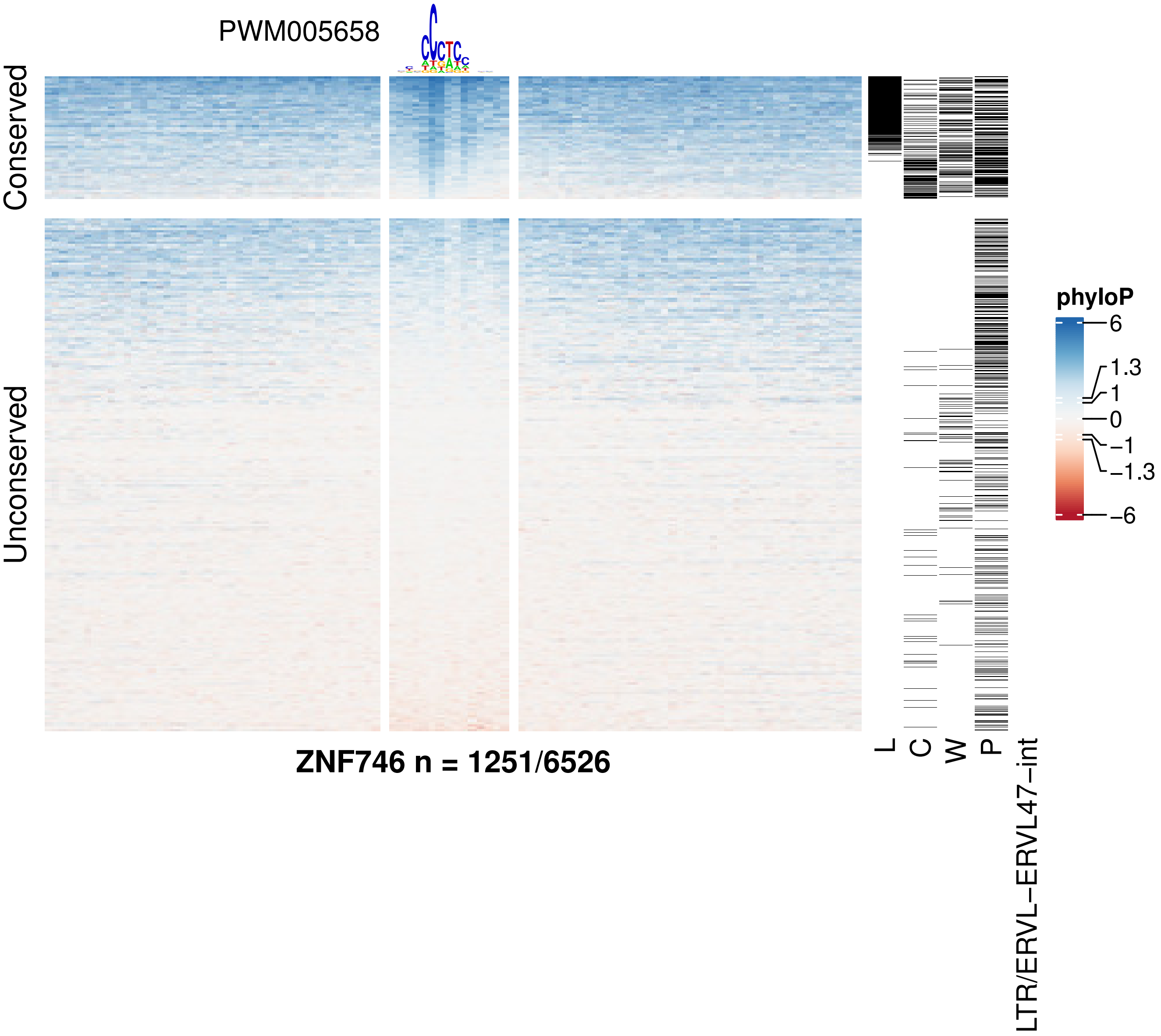
PWM005658
Conserved
Unconserved
phyloP
6
1.3
1
0
−1
−1.3
−6
ZNF746 n = 1251/6526
L
C
W
P
LTR/ERVL−ERVL47−int

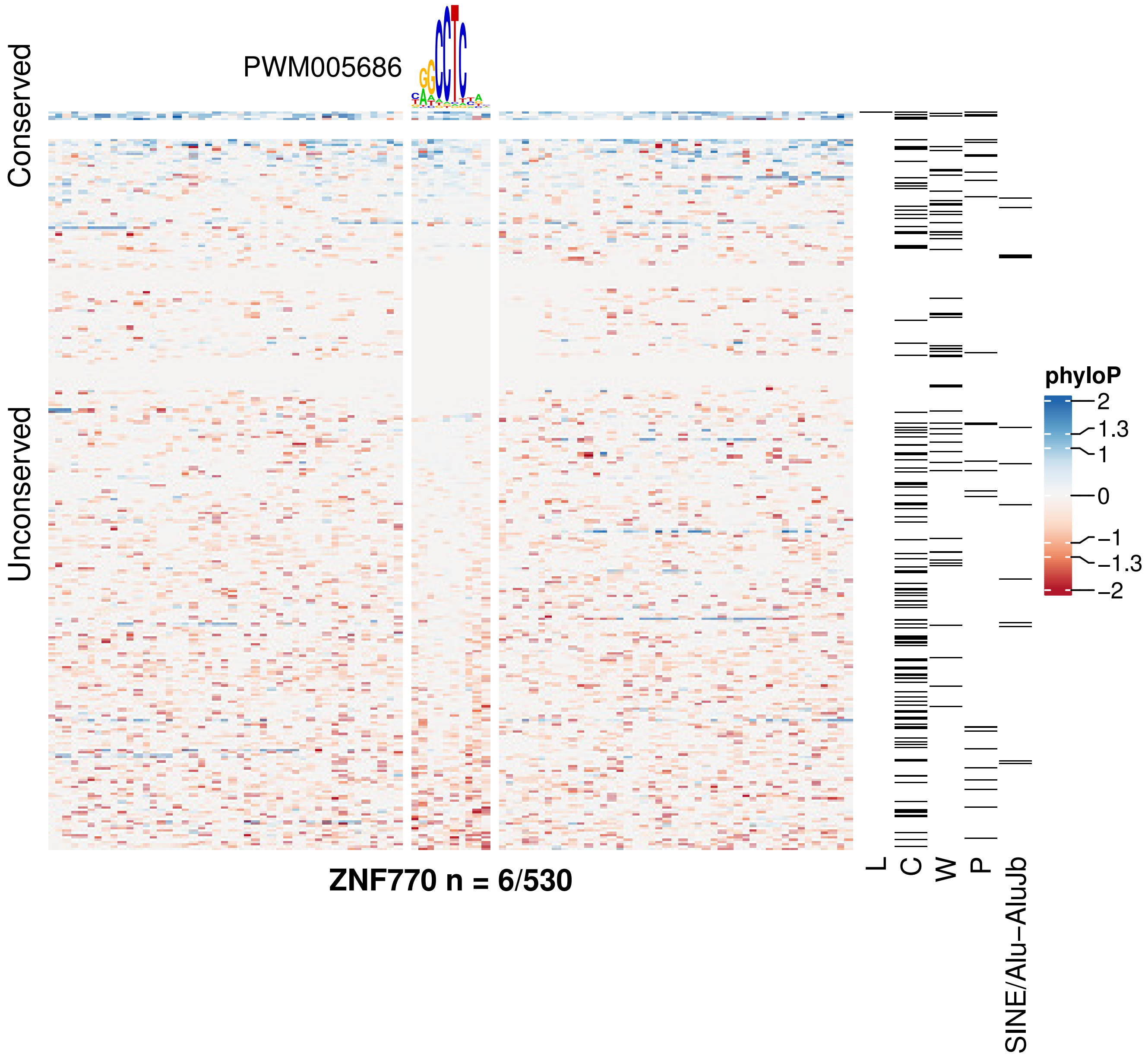

PWM005686
Conserved
Unconserved
ZNF770 n = 6/530
L
C
W
P
SINE/Alu−AluJb
phyloP
2
1.3
1
0
−1
−1.3
−2

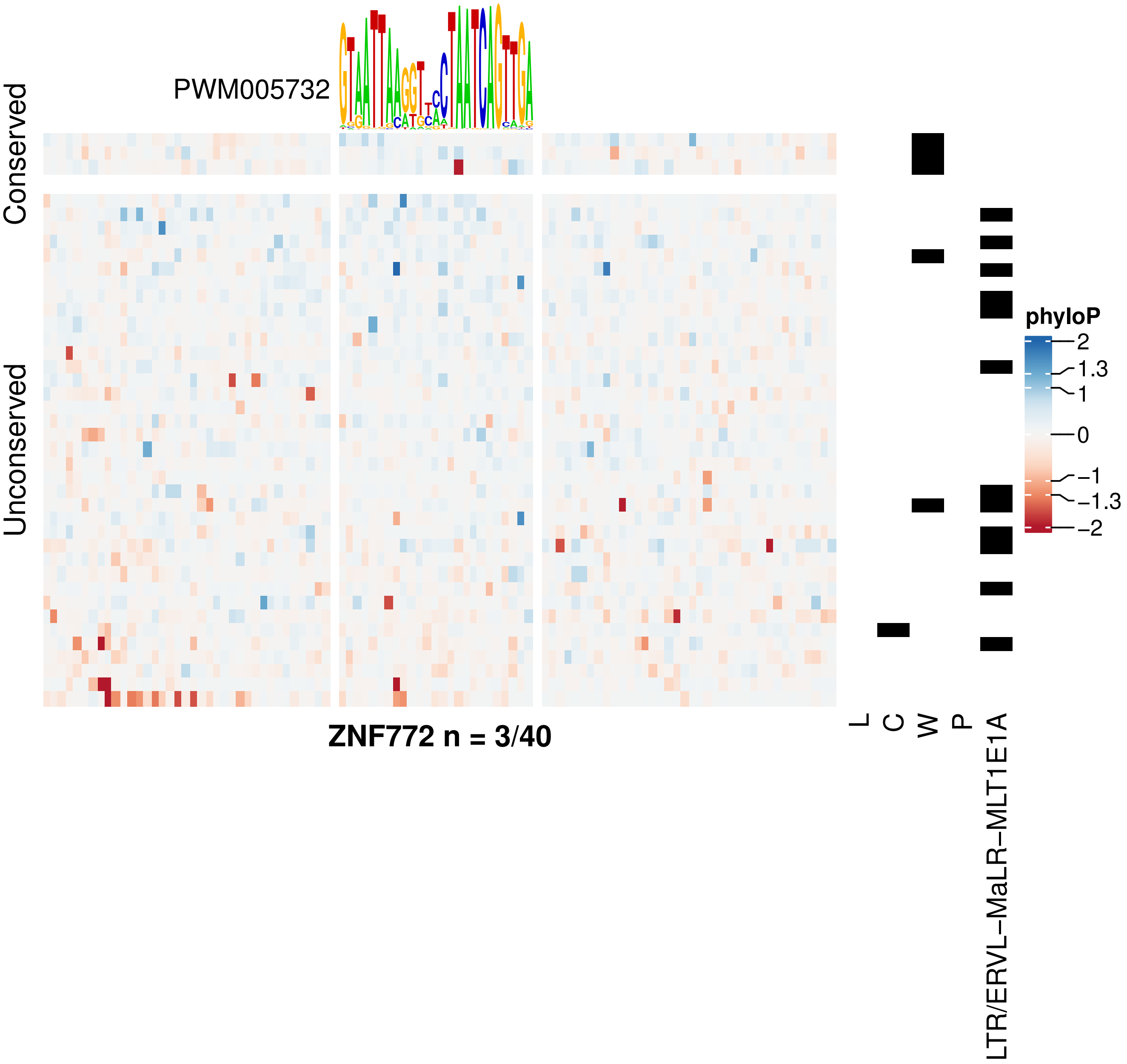

PWM005732
Conserved
Unconserved
phyloP
2
1.3
1
0
−1
−1.3
−2
L
C
W
P
LTR/ERVL−MaLR−MLT1E1A
ZNF772 n = 3/40

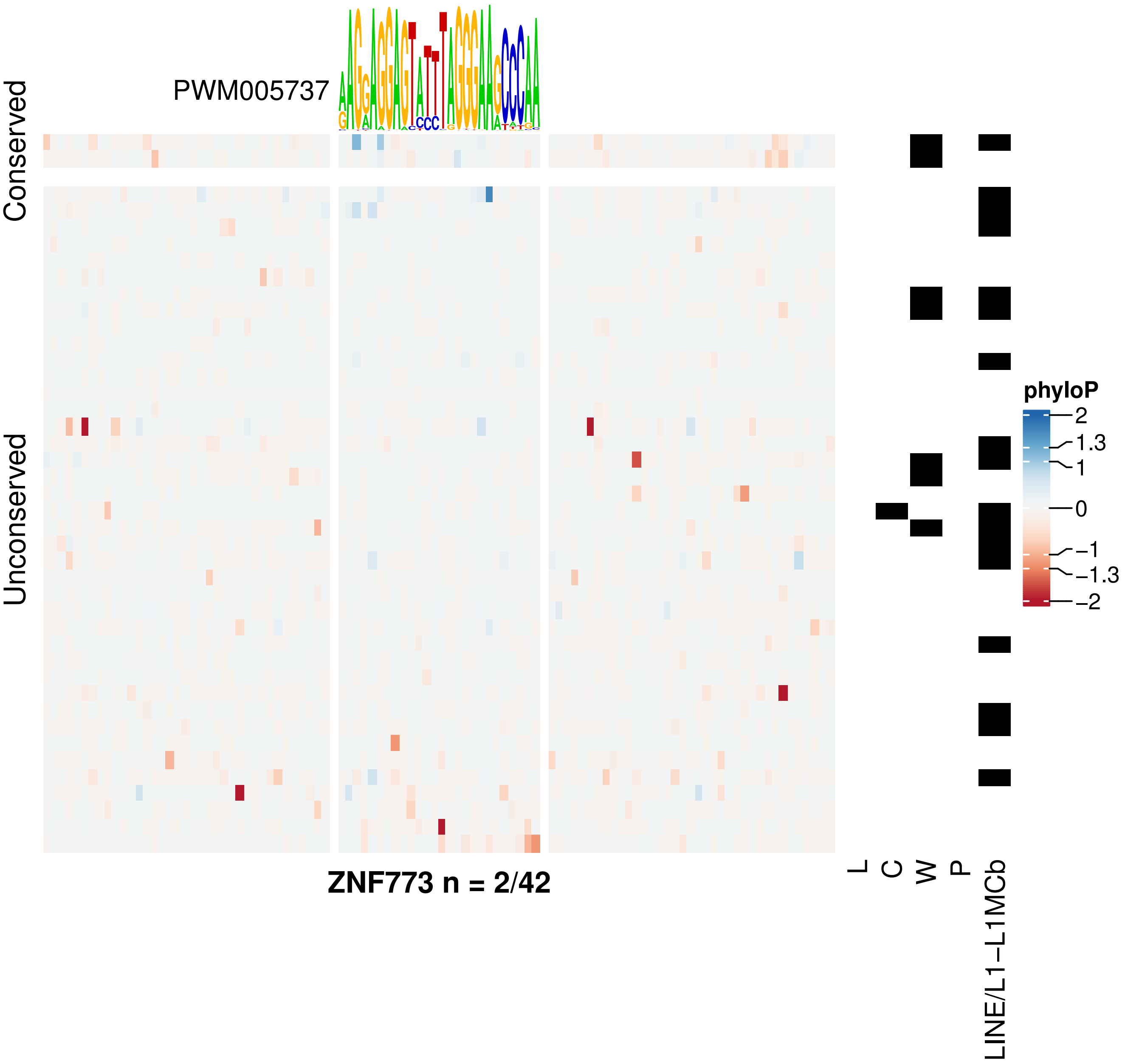
PWM005737
Conserved
Unconserved
ZNF773 n = 2/42
L
C
W
P
LINE/L1−L1MCb
phyloP
2
1.3
1
0
−1
−1.3
−2

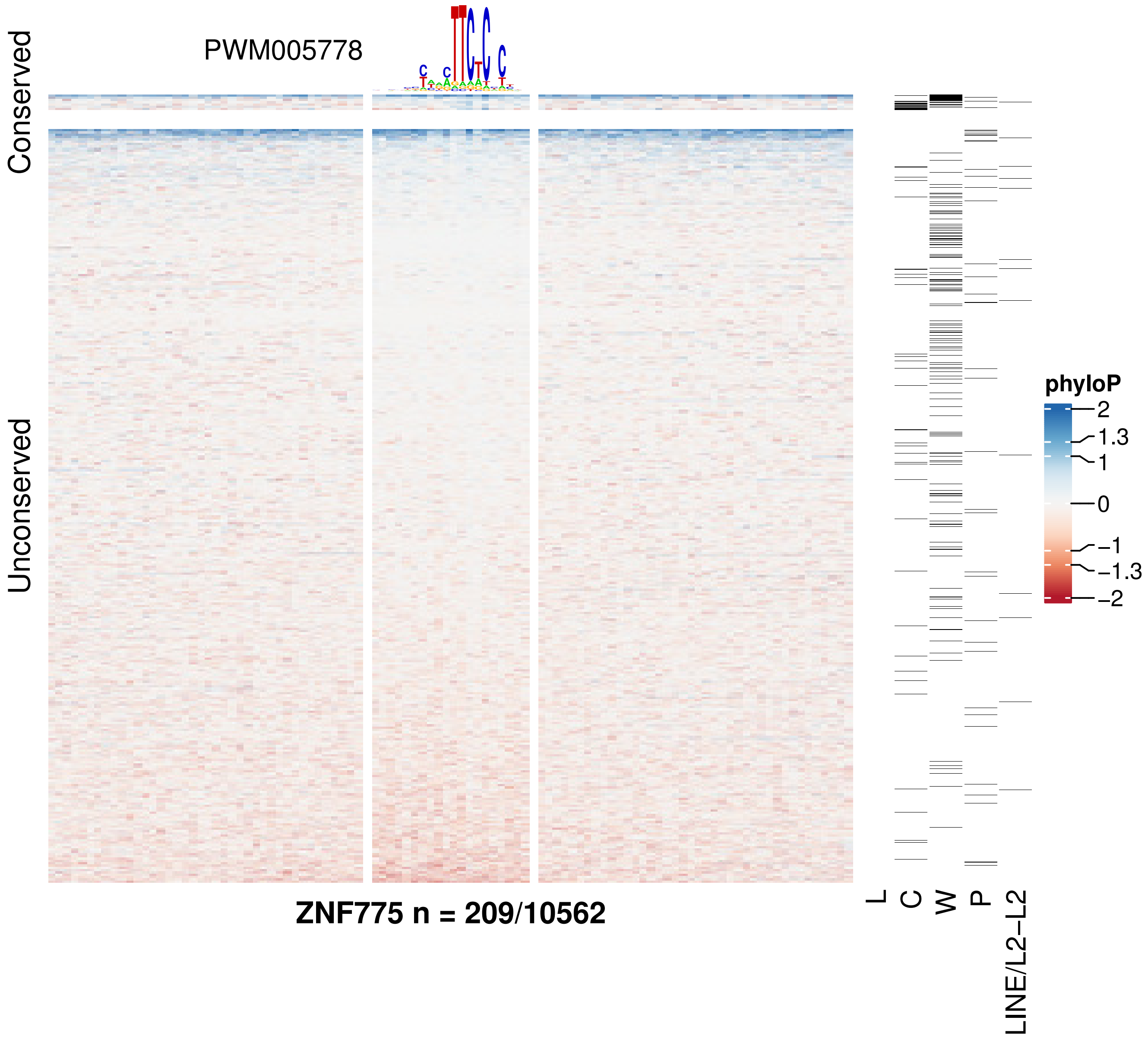
PWM005778
Conserved
Unconserved
ZNF775 n = 209/10562
L
C
W
P
LINE/L2−L2
phyloP
2
1.3
1
0
−1
−1.3
−2

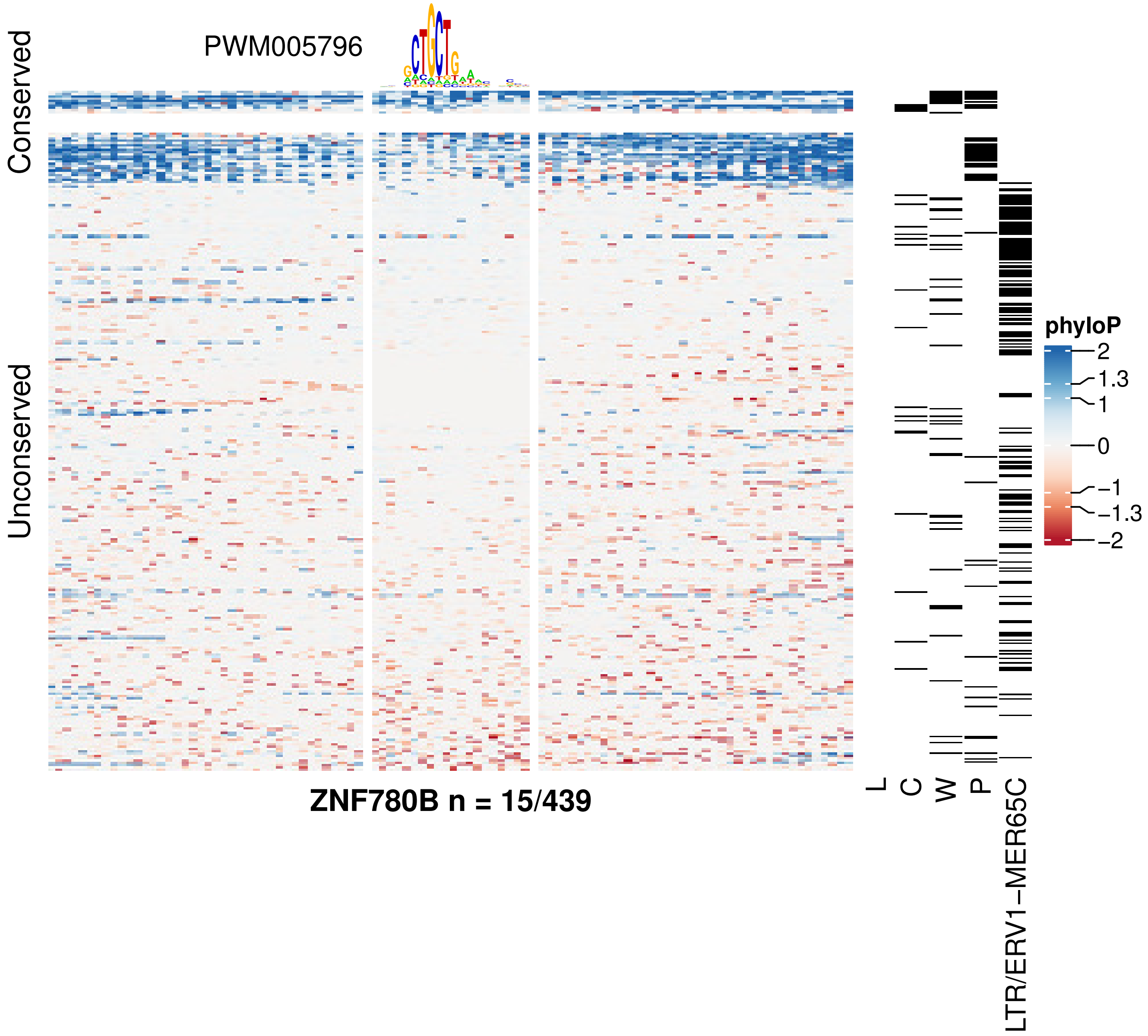

PWM005796
Conserved
Unconserved
ZNF780B n = 15/439
L
C
W
P
LTR/ERV1−MER65C
phyloP
2
1.3
1
0
−1
−1.3
−2

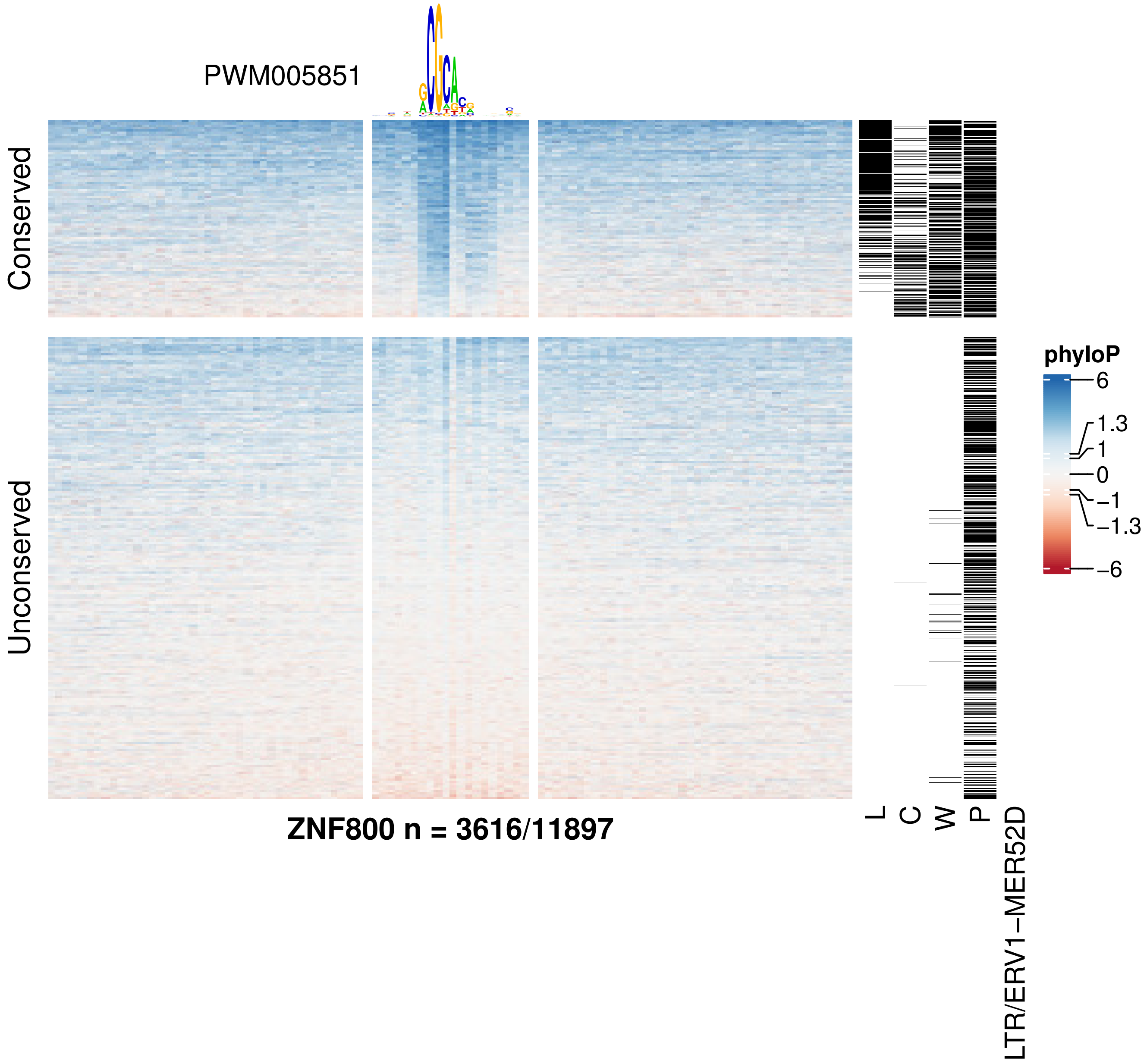

PWM005851
Conserved
Unconserved
ZNF800 n = 3616/11897
L
C
W
P
LTR/ERV1−MER52D
phyloP
6
1.3
1
0
−1
−1.3
−6

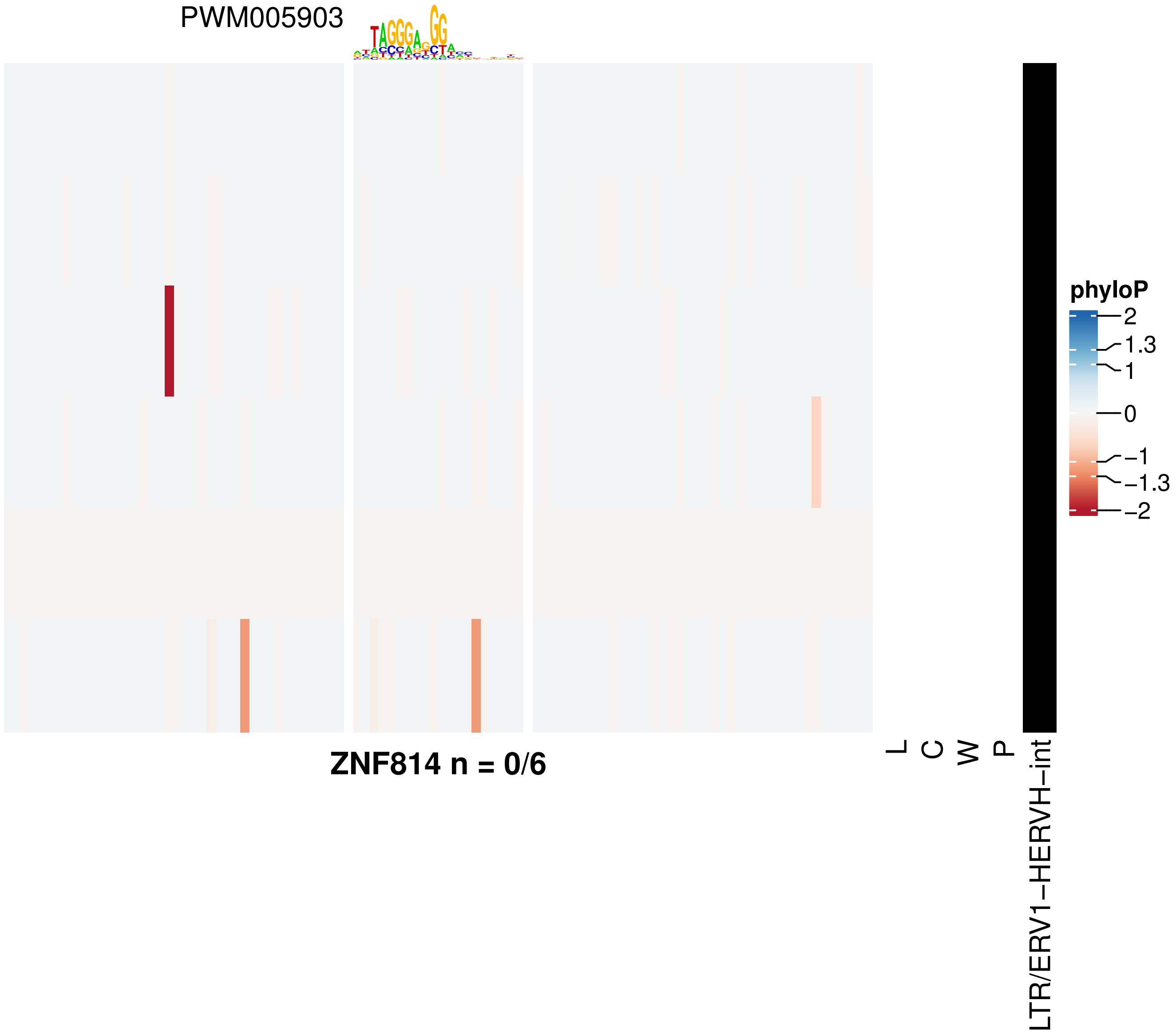
PWM005903
ZNF814 n = 0/6
L
C
W
P
LTR/ERV1−HERVH−int
phyloP
2
1.3
1
0
−1
−1.3
−2

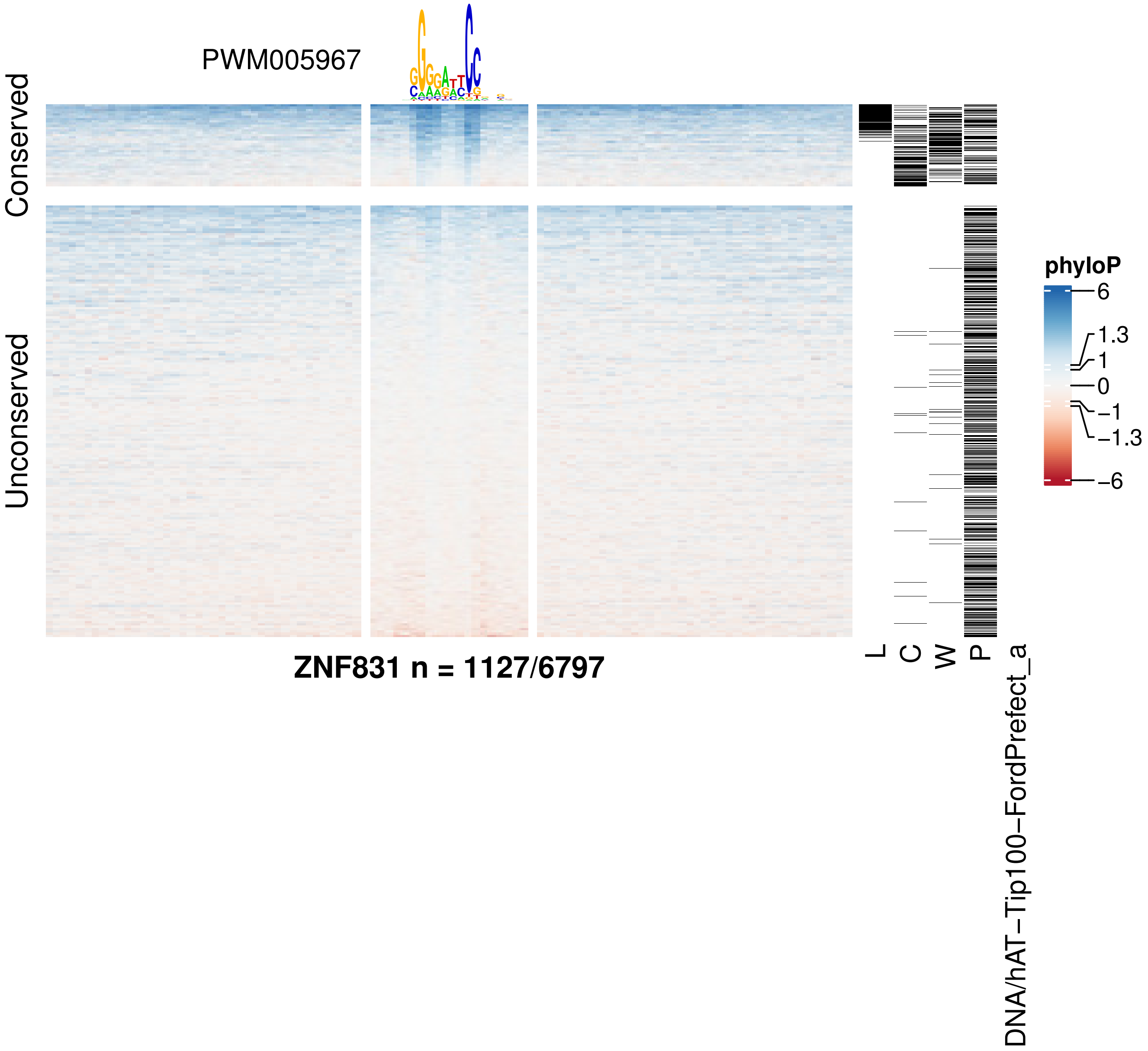
PWM005967
Conserved
Unconserved
ZNF831 n = 1127/6797
L
C
W
P
DNA/hAT−Tip100−FordPrefect_a
phyloP
6
1.3
1
0
−1
−1.3
−6

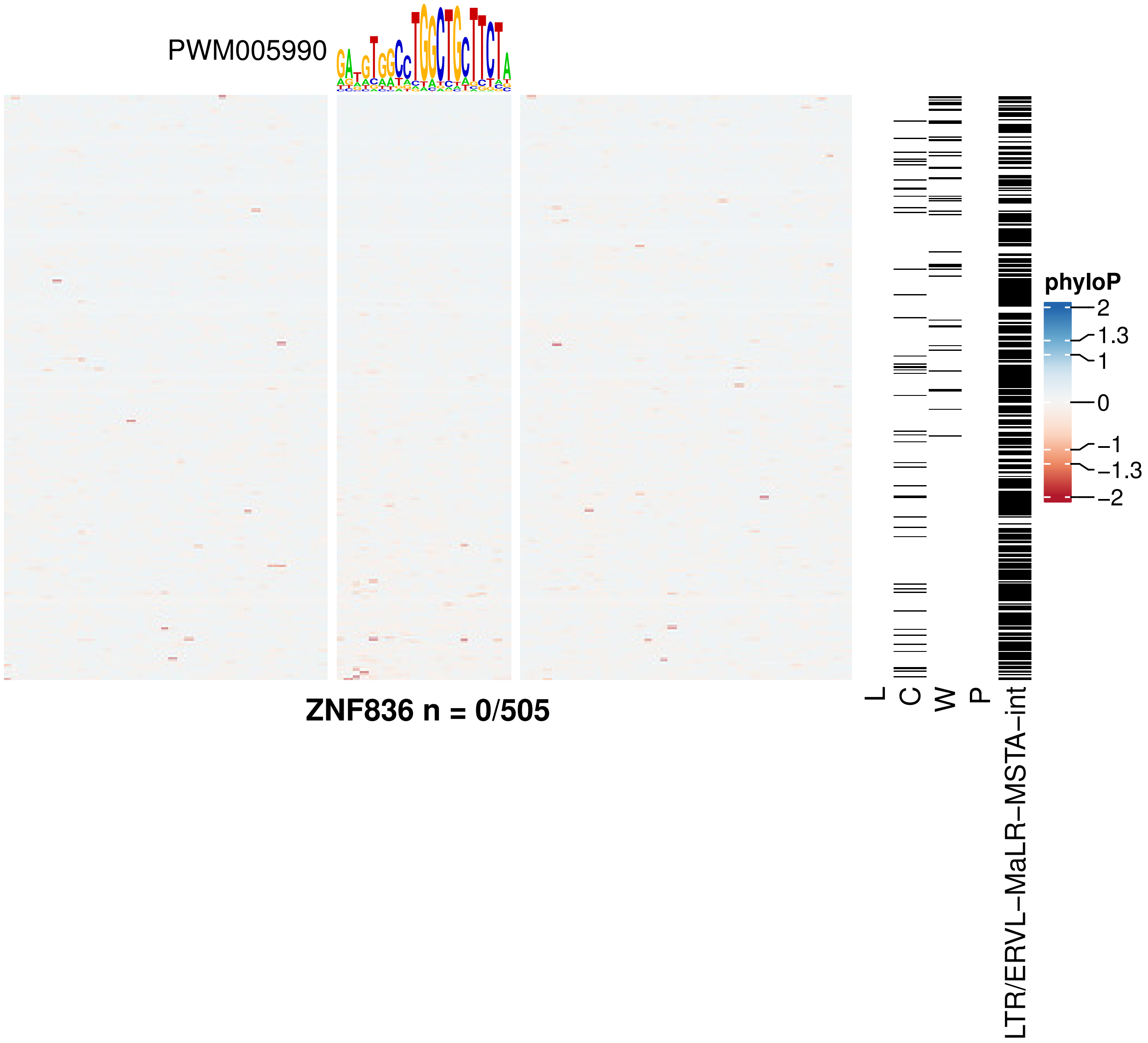
PWM005990
ZNF836 n = 0/505
L
C
W
P
LTR/ERVL−MaLR−MSTA−int
phyloP
2
1.3
1
0
−1
−1.3
−2

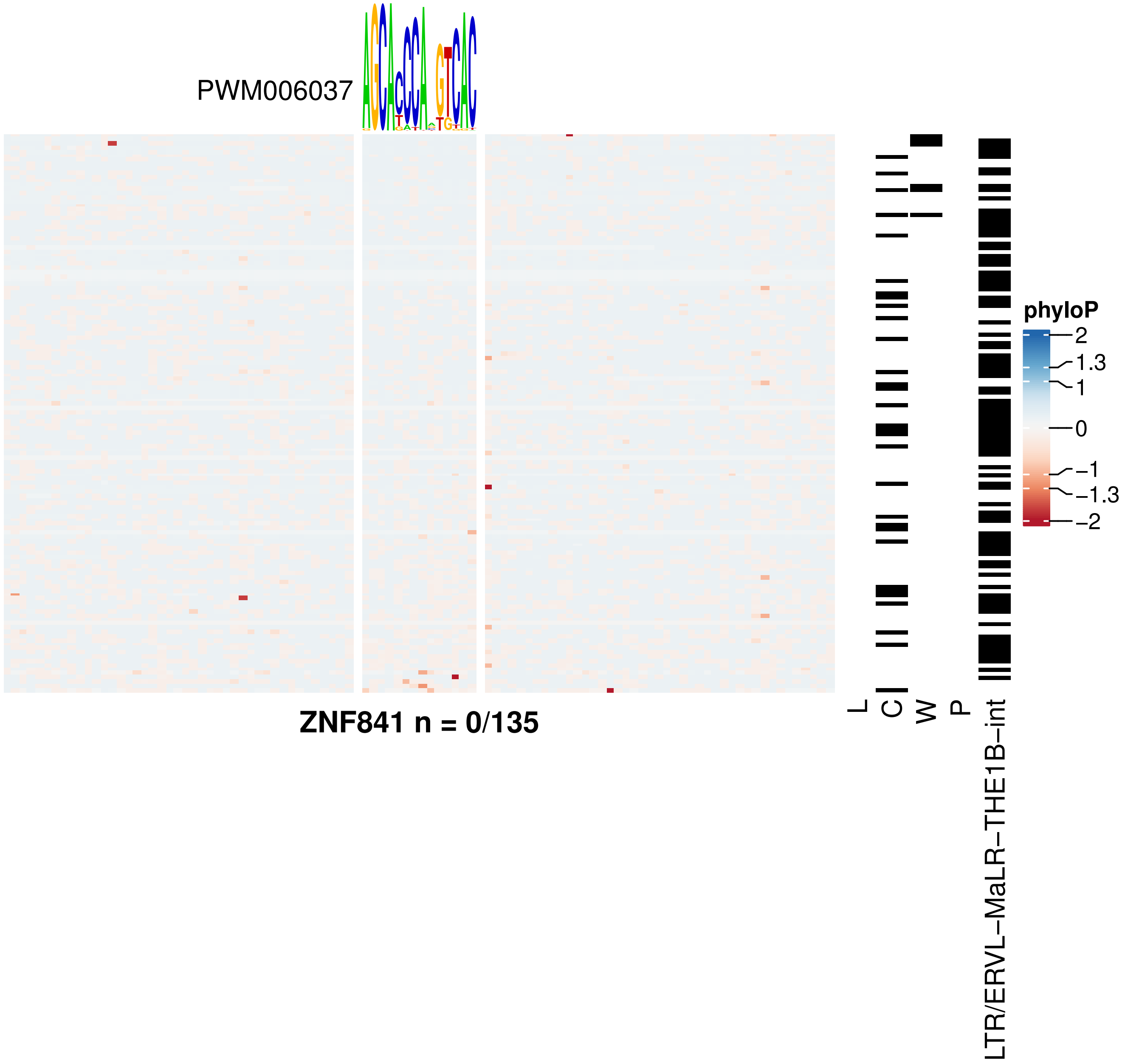
PWM006037
ZNF841 n = 0/135
L
C
W
P
LTR/ERVL−MaLR−THE1B−int
phyloP
2
1.3
1
0
−1
−1.3
−2

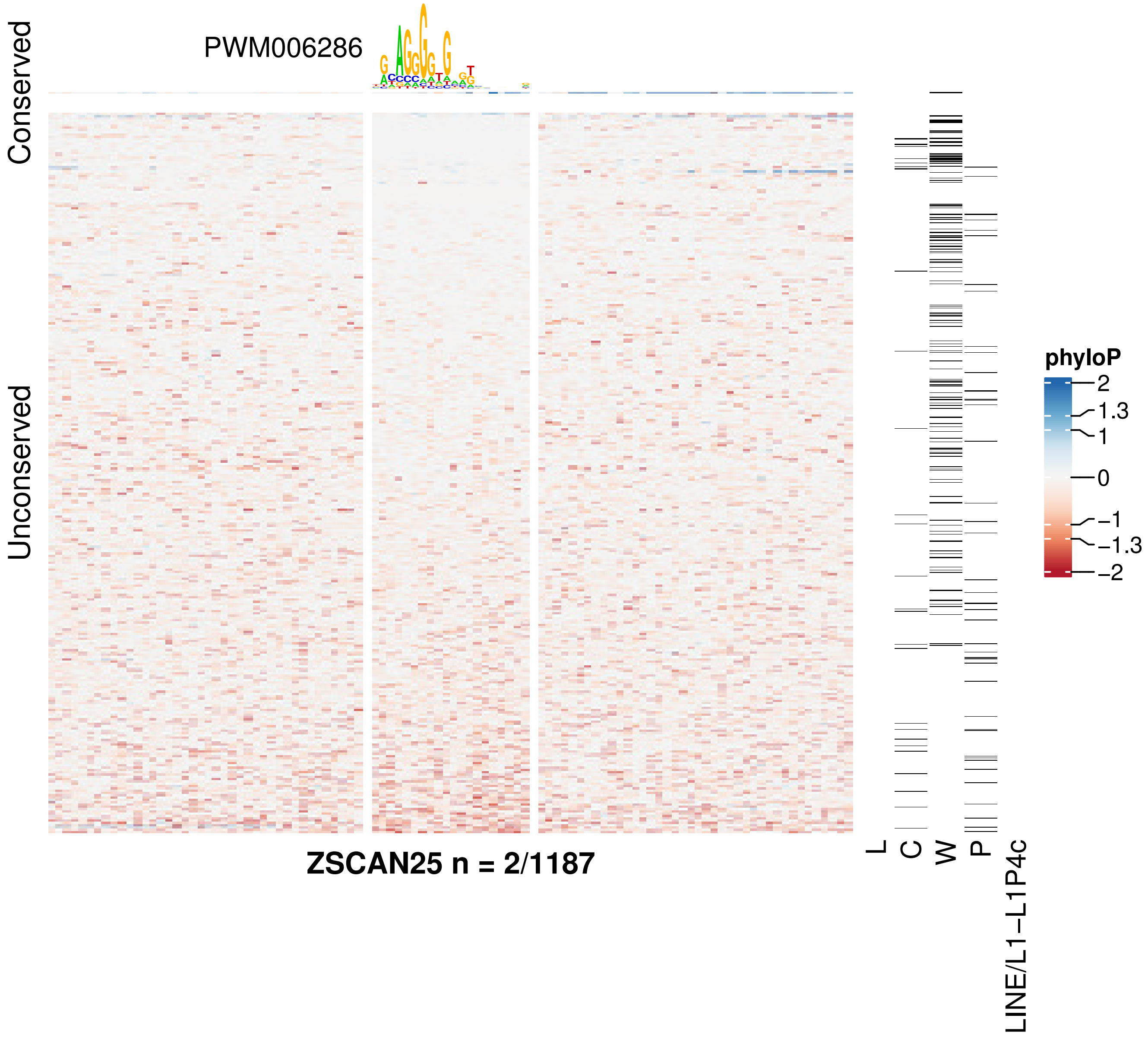

PWM006286
Conserved
Unconserved
ZSCAN25 n = 2/1187
L
C
W
P
LINE/L1−L1P4c
phyloP
2
1.3
1
0
−1
−1.3
−2

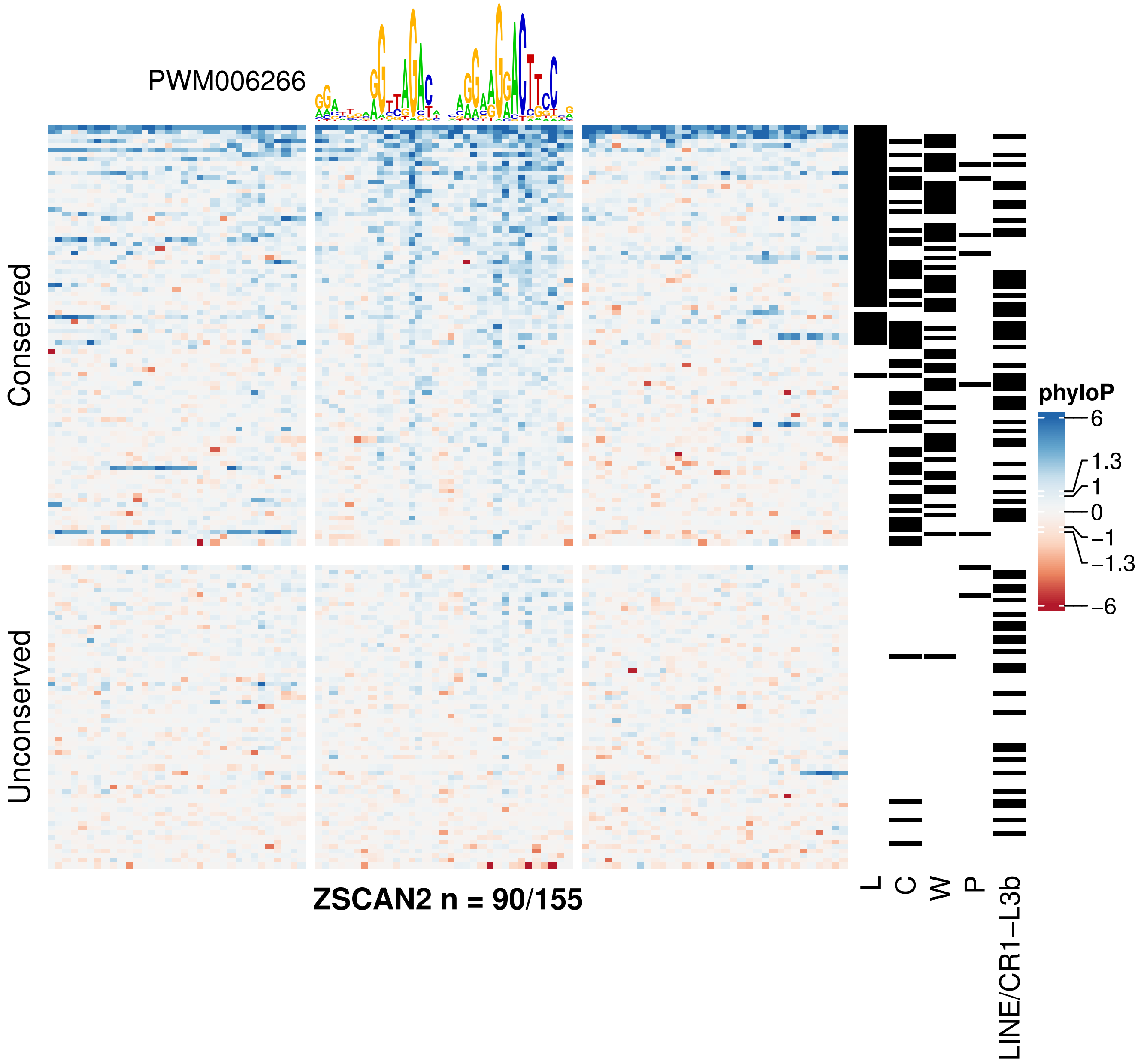

PWM006266
Conserved
Unconserved
ZSCAN2 n = 90/155
L
C
W
P
LINE/CR1−L3b
phyloP
6
1.3
1
0
−1
−1.3
−6
