## Supplementary Document 3 for "Extensive binding of nebulous human transcription factors to genomic dark matter"

### Supplementary Document 3: Overview of Dark TFs' function

We examined existing literature and databases to survey known and potential functions for the Dark TFs, and related it to the data we collected (**Fig. 7**, **Supplementary Table 4**). Most Dark TFs have apparent roles in repression of transcription. 36 out of 54 are KZNFs, and for 18 of them (and one non-KRAB TF, ZNF888), physical association with KAP1 has been verified[1-5]. The KZNFs may also have repressive functions beyond the recruitment of KAP1[1,2]. Five of these Dark TF KZNFs also interact with TRIM39, which itself interacts with numerous ubiquitin conjugating enzymes, H3K4 demethylase KDM1A, and dozens of other KZNFs[3,6]. Two additional Dark TF KZNFs (and two other Dark TFs) interact with TRIM33, a member of the TIF1Y complex that specifically suppresses TGF$\beta$-responsive genes by directly interacting with the histone subunits as well as E3 ubiquitin ligase[7].

Nine of the eleven non-KRAB Dark C2H2-zf proteins also appear to contribute to the formation and maintenance of heterochromatin, by association with chromatin proteins (CBX/HP1) directly, or via recruitment of other C2H2-zf proteins. One of them, ZNF518B, was identified as a partner of both H3K27 methylase EZH2 and H3K9/H3K27 methylase G9A, and to promote H3K9me2[8]. ZNF518B and ZNF280D both associate with multiple CBX/HP1 proteins[3,6]. ZNF518B binds many primate-specific L1 elements, but its most conserved binding sites are in its own promoter, suggesting a critical negative feedback mechanism (**Supplementary Fig. 11**). In another example, ZNF516 associates with the multifunctional CTBP1/KDM1A/RCOR1 corepressor complex, and its repressive function was shown in reporter assays[9]. Intriguingly, 22 of the 47 C2H2-zf proteins, including both KRAB and non-KRAB C2H2-zf proteins (as well as transposon-derived ZBED9, SOX2, and SP100) interact with other C2H2-zf proteins, often extensively[3], suggesting a potentially widespread role in organization of chromosome topology.

Four additional Dark TFs have other potential roles in repression of transcription. Three of them are the paralogous nuclear speckle proteins SP100, SP140, and SP140L. Each contains a SAND domain, which we confirmed binds to unmethylated CG dinucleotides *in vitro*[10], and CG-containing motifs are enriched in their ChIP-seq peaks[11]. These proteins also contain PHD and BRD domains, which typically function as epigenetic readers[12]. In our ChIP-seq data, they are enriched at sites of H3K27me3 methylation (**Fig. 7**, **Supplementary Table 4**). The fourth protein is SCML4, a polycomb group protein that was included in our study because it contains an AT hook, but we did not obtain evidence for its sequence-specific DNA binding. Thus, it may be more properly described as a chromatin protein. SCML4 is reported to associate with H3K4 demethylase KDM5C[13] as well as ubiquitination factors FBXO11 and UBR1[14].

Five of the non-KZNF Dark TFs may have roles other than repression. One of them, SOX2, is a well-known pioneer factor that can bind to motif matches within unmodified closed chromatin, but is inhibited to some extent by H3K9me3[15]. Indeed, in the ChIP-seq data reported here, most (66%) of its TOP sites are in “empty” chromatin, and only 5% overlap with ATAC-seq peaks in unperturbed HEK293 cells, consistent with its pioneer function. Less than 2% of SOX2 peaks overlap with heterochromatin (defined by ChromHMM mainly by H3K9me3 and H327me3), consistent with H3K9me3 being refractory to SOX2 binding. Two additional Dark TFs may also represent pioneers: TPRX1 has recently been described as a master regulator in zygotic genome activation[16], while SALL3 controls the differentiation of hiPSCs into cardiomyocytes vs neural cells[17]. In contrast, two other Dark TFs have been described as impacting DNA metabolism. ZNF384, which we find binds many Alu and Poly-A repeats, as described above, is also known to bind Ku and recruit NHEJ factors to double-strand breaks[18]. ZNF146 binds L1 elements, and its depletion slows the replication fork[19].

Reported physiological consequences for some the Dark TFs perturbations are available and are summarized in **Fig. 7**, right column (**Supplementary Table 4** provides the values and sources). The correspondence between recorded phenotypes and the discovered binding sites is discussed in the main text. Expanding on one Dark TF, ZBTB40, its most conserved TOP resides within the 3’UTR of PRKACA (**Fig. 5g**). PRKACA encodes the catalytic subunit α of protein kinase A, whose deficiency is associated with fertility defects in male mice and humans[20]. This site is also less than 1 kb from the TSS of the chromatin regulator SAMD1, which impacts sperm cells[21].

### REFERENCES


1. Helleboid, P.Y. *et al.* The interactome of KRAB zinc finger proteins reveals the evolutionary history of their functional diversification. *EMBO J* **38**, e101220 (2019).
2. Schmitges, F.W. *et al.* Multiparameter functional diversity of human C2H2 zinc finger proteins. *Genome Res* **26**, 1742–1752 (2016).
3. Huttlin, E.L. *et al.* Dual proteome-scale networks reveal cell-specific remodeling of the human interactome. *Cell* **184**, 3022–3040 e28 (2021).
4. Silva, F.P., Hamamoto, R., Furukawa, Y. & Nakamura, Y. TIPUH1 encodes a novel KRAB zinc-finger protein highly expressed in human hepatocellular carcinomas. *Oncogene* **25**, 5063–70 (2006).
5. Kim, J.J. *et al.* Systematic bromodomain protein screens identify homologous recombination and R-loop suppression pathways involved in genome integrity. *Genes Dev* **33**, 1751–1774 (2019).
6. Oughtred, R. *et al.* The BioGRID database: A comprehensive biomedical resource of curated protein, genetic, and chemical interactions. *Protein Sci* **30**, 187–200 (2021).
7. Agricola, E., Randall, R.A., Gaarenstroom, T., Dupont, S. & Hill, C.S. Recruitment of TIF1gamma to chromatin via its PHD finger-bromodomain activates its ubiquitin ligase and transcriptional repressor activities. *Mol Cell* **43**, 85–96 (2011).
8. Maier, V.K. *et al.* Functional Proteomic Analysis of Repressive Histone Methyltransferase Complexes Reveals ZNF518B as a G9A Regulator. *Mol Cell Proteomics* **14**, 1435–46 (2015).
9. Li, L. *et al.* ZNF516 suppresses EGFR by targeting the CtBP/LSD1/CoREST complex to chromatin. *Nat Commun* **8**, 691 (2017).
10. Huggenvik, J.I. *et al.* Characterization of a nuclear deformed epidermal autoregulatory factor-1 (DEAF-1)-related (NUDR) transcriptional regulator protein. *Mol Endocrinol* **12**, 1619–39 (1998).
11. Vorontsov, I.E. *et al.* Cross-platform DNA motif discovery and benchmarking to explore binding specificities of poorly studied human transcription factors. *bioRxiv*, 2024.11.11.619379 (2024).
12. Fraschilla, I. & Jeffrey, K.L. The Speckled Protein (SP) Family: Immunity's Chromatin Readers. *Trends Immunol* **41**, 572–585 (2020).
13. Tumber, A. *et al.* Potent and Selective KDM5 Inhibitor Stops Cellular Demethylation of H3K4me3 at Transcription Start Sites and Proliferation of MM1S Myeloma Cells. *Cell Chem Biol* **24**, 371–380 (2017).
14. Huttlin, E.L. *et al.* The BioPlex Network: A Systematic Exploration of the Human Interactome. *Cell* **162**, 425–440 (2015).
15. Soufi, A., Donahue, G. & Zaret, K.S. Facilitators and impediments of the pluripotency reprogramming factors' initial engagement with the genome. *Cell* **151**, 994–1004 (2012).
16. Zou, Z. *et al.* Translatome and transcriptome co-profiling reveals a role of TPRXs in human zygotic genome activation. *Science* **378**, abo7923 (2022).
17. Kuroda, T. *et al.* SALL3 expression balance underlies lineage biases in human induced pluripotent stem cell differentiation. *Nat Commun* **10**, 2175 (2019).

18. Singh, J.K. *et al.* Zinc finger protein ZNF384 is an adaptor of Ku to DNA during classical non-homologous end-joining. *Nat Commun* **12**, 6560 (2021).
19. Feu, S. *et al.* OZF is a Claspin-interacting protein essential to maintain the replication fork progression rate under replication stress. *FASEB J* **34**, 6907–6919 (2020).
20. Burton, K.A. *et al.* Haploinsufficiency at the protein kinase A RI alpha gene locus leads to fertility defects in male mice and men. *Mol Endocrinol* **20**, 2504–13 (2006).
21. Stielow, B. *et al.* The SAM domain-containing protein 1 (SAMD1) acts as a repressive chromatin regulator at unmethylated CpG islands. *Sci Adv* **7**(2021).
