## Supplementary Document 4 for "Extensive binding of nebulous human transcription factors to genomic dark matter"

### Supplementary Document 4: Analysis of binding site ages for TFs

In addition to insertions such as TEs, new TF binding sites can emerge from random mutations in pre-existing sequences. This mechanism is thought to be dominant for traditional enhancer-binding TFs[1]. To determine whether binding sites for Codebook TFs evolved from ancestral DNA, we estimated the age of each TOP site for each TF.

We gauged binding site age as that of the oldest ancestral genome that contains the entire site, using halLiftover[2] to map TOP PWM hits to syntenic loci in all other genomes in the Zoonomia 240-mammal alignment[3], including 9 reconstructed genomes ancestral to human, and calculated each sequence's % identity to the aligned human TOP.

We then assigned an age using multiple criteria, identifying the oldest ancestral genome with a gapless alignment of any % identity to the human PWM hit (**Supplementary Fig. 9a).** This is a simple heuristic, but we obtained a qualitatively similar conclusions using other approaches to estimate binding site age (**Supplementary Fig. 12**), including the oldest extant *species* with a gapless alignment to a human TOP (at various threshold % identities), or the oldest *clade* wherein 60% of species have a gapless alignment to the human TOP (at various threshold % identities). We acquired the age of each clade from TimeTree[4].

As expected, TOP sites for the Dark TFs (most of which correspond to TEs) are estimated to be considerably younger on average than those of Promoter TFs (median ages of 19.5 and >=99.2 MYA, respectively, **Supplementary Fig. 9a),** but there is a large overlap of age distributions between the two TF groups. Both groups also contain TFs with binding sites at both extremes (i.e. very old or very young binding sites). Thus, average age of the binding site is not a discriminating characteristic of Promoter vs. Dark TFs.

We also estimated the ages of the TFs, by first acquiring all vertebrate ortholog annotations and ortholog quality statistics for each TF from Ensembl[5], and ages of each pair of species from TimeTree[4] (**Supplementary Fig. 9c**, **Supplementary Table 3**). The age of a TF was taken as the oldest ortholog annotated as having a 1-1 relationship with human, or having a gene order conservation (GOC, a metric of synteny) score >= 50 and classification as a high-confidence ortholog by Ensembl.

Overall, TFs in both classes tend to be older than the sites they bind: Promoter TFs have a median age of 429 MYA, while Dark TFs have a median age of 99.19 MYA, which is approximately as old as a typical binding site even for Promoter TFs. These results are consistent with the established phenomenon of TF binding site turnover[1]. They are also reminiscent of previous observations with KZNFs showing weak correlation between the age of the KZNFs and the age of their binding sites, and inconsistent with a model in which the KZNFs evolve only to silence TEs[6]. Together with the retention of many KZNFs that bind extinct TEs, this finding supports the notion that KZNFs must frequently take on additional roles, e.g., in regulation of host genes.

### REFERENCES


1. Villar, D. *et al.* Enhancer evolution across 20 mammalian species. *Cell* **160**, 554–66 (2015).
2. Hickey, G., Paten, B., Earl, D., Zerbino, D. & Haussler, D. HAL: a hierarchical format for storing and analyzing multiple genome alignments. *Bioinformatics* **29**, 1341–2 (2013).
3. Armstrong, J. *et al.* Progressive Cactus is a multiple-genome aligner for the thousand-genome era. *Nature* **587**, 246–251 (2020).
4. Kumar, S. *et al.* TimeTree 5: An Expanded Resource for Species Divergence Times. *Mol Biol Evol* **39**(2022).
5. Dyer, S.C. *et al.* Ensembl 2025. *Nucleic Acids Res* **53**, D948–D957 (2025).
6. Imbeault, M., Helleboid, P.Y. & Trono, D. KRAB zinc-finger proteins contribute to the evolution of gene regulatory networks. *Nature* **543**, 550–554 (2017).
